# A User-Friendly Guide to Using Distance Measures to Compare Time Series in Ecology

**DOI:** 10.1101/2022.05.11.491333

**Authors:** Shawn Dove, Monika Böhm, Robin Freeman, Sean Jellesmark, David Murrell

## Abstract

1. Time series are a critical component of ecological analysis, used to track changes in biotic and abiotic variables. Information can be extracted from the properties of time series for tasks such as classification, clustering, prediction, and anomaly detection. These common tasks in ecological research rely on the notion of (dis-) similarity which can be determined by using distance measures. A plethora of distance measures have been described in the scientific literature, but many of them have not been introduced to ecologists. Furthermore, little is known about how to select appropriate distance measures and the properties they focus on for time-series related tasks.
2. Here we describe 16 potentially desirable properties of distance measures, test 42 distance measures for each property, and present an objective method to select appropriate distance measures for any task and ecological dataset. We then demonstrate our selection method by applying it to a set of real-world data on breeding bird populations in the UK. We also discuss ways to overcome some of the difficulties involved in using distance measures to compare time series.
3. Our real-world population trends exhibit a common challenge for time series comparison: a high level of stochasticity. We demonstrate two different ways of overcoming this challenge, first by selecting distance measures with properties that make them well-suited to comparing noisy time series, and second by applying a smoothing algorithm before selecting appropriate distance measures. In both cases, the distance measures chosen through our selection method are not only fit-for-purpose but are consistent in their rankings of the population trends.
4. The results of our study should lead to an improved understanding of, and greater scope for, the use of distance measures for comparing time series within ecology, and allow for the answering of new ecological questions.

## 1. Introduction

Time series are a critical component of ecological analysis: ecologists use time series to track changes in biotic variables, such as population sizes and mean growth rates of individuals, as well as abiotic variables, such as temperature and atmospheric carbon dioxide. Time series provide insight into food web and ecosystem function and the causes and effects of environmental change, and are vital to any scientific approach to environmental management (Boero *et al*., 2015). Time series datasets may contain thousands or even millions of time series (e.g., The Living Planet Index – WWF, 2020; BioTIME - Dornelas *et al*., 2018; the North American Breeding Bird Survey - Pardieck *et al*., 2019; the British Trust for Ornithology Breeding Bird Survey - Harris *et al*., 2020; and the Continuous Plankton Recorder Survey - Edwards *et al*., 2012). Ecologists make inferences through time series comparisons. For example, one might look for similarities or differences in climate change response between populations within or across geographic or taxonomic groups. However, examining and analysing each time series by hand is unwieldy.

Data mining of time series is the process of extracting information from the properties of time series for tasks such as classification, clustering, prediction, and anomaly detection (Esling and Agon, 2012). These tasks are common in ecology, e.g., clustering time series of parasite counts to identify infection patterns (Marques *et al*., 2018); predicting the emergence of fruiting bodies by classifying time series of environmental drivers (Capinha, 2019); identifying insect species by classifying wingbeat frequency signals (Potamitis *et al*., 2015); surveying bird population sizes by classifying recorded calls (Priyadarshani *et al*., 2020); and predicting species distributions based on time series of environmental variables (Capinha *et al*., 2020). These tasks all rely on the notion of (dis-) similarity. Clustering involves grouping similar time series together by maximizing the similarity within groups and minimizing the similarity between groups (Liao, 2005; Esling and Agon, 2012; Aghabozorgi *et al*., 2015). Classification is like clustering, except labels are predefined and new time series are assigned to existing clusters to which they are most similar (Keogh and Kasetty, 2003; Esling and Agon, 2012). Prediction may rely on similarity to determine accuracy by comparing predicted time series against the originals (Capinha, 2019; Esling and Agon, 2012). Finally, anomaly detection involves comparing time series against an anomaly-free model to determine if they fall outside of a similarity threshold (Teng, 2010; Esling and Agon, 2012).

Similarity between time series can be determined by using distance measures to measure its inverse: dissimilarity. Dissimilarity is more intuitive as a measurement because a value of zero occurs when two time series are identical (while similarity is at a scale-dependent maximum value). Distance measures can be broadly categorized into four different types: shape-based, feature-based, model-based, and compression-based. Shape-based distances compare the shapes of time series by measuring differences in the raw data values (Aghabozorgi *et al*., 2015; Esling and Agon, 2012) and can be further divided into lock-step measures and elastic measures. Lock-step measures compare each time point of one time series to the corresponding time point of another time series, while elastic measures allow a single point to be matched with multiple points or no points (Wang *et al*., 2013). Elastic measures fall into two groups. The first, Dynamic Time Warping (DTW), computes an optimal match between two time series by allowing single points to be matched with multiple points, thus allowing local distortion or “warping” of the time dimension (Esling and Agon, 2012). The second comprises edit distances, which compare the minimum number of “edits”, or changes, required to transform one time series into another (Esling and Agon, 2012). They are based on the concept of transforming one string into another by changing one letter at a time, with each “edit” being an insertion, deletion, or substitution. Feature-based distances compute some feature of time series, such as Discrete Fourier Transforms or autocorrelation coefficients, and use either a specialized or common distance function (e.g., the Euclidean distance) to determine the distance between the computed features (Mori *et al*., 2016). Model-based distances compare the parameters of models fitted to the time series, such as autoregressive moving average (ARMA) models, with the advantage that they can incorporate knowledge about the process used to generate the time series data (Esling and Agon, 2012). Finally, compression-based distances assess the similarity of two digital objects according to how well they can be “compressed” when connected (Esling and Agon, 2012; Cilibrasi and Vitanyi, 2005); the more similar the objects, the better they compress when joined in series (Esling and Agon, 2012). Although there are comparatively few model-based and compression-based distance measures, there are many shape-based and feature-based measures available.

The choice of distance measure for any task should depend on the properties of the data to be analysed and the nature of the task (Esling and Agon, 2012). In practice, choosing a distance measure often becomes a matter of convenience. For example, the well-known and easy to use Euclidean distance is among the most widely used distance measures, although there are often better choices (Wang *et al*., 2012; Paparrizos *et al*., 2020). When investigating the performance of five distance measures for comparing animal movement trajectories, Cleasby *et al*. (2019) found that the most used measure was the least appropriate choice. One problem is that many distance measures originate within computer science, information science, systems science, and mathematics, and few are in common use within ecology. Another problem is that information on the strengths, weaknesses, and appropriate uses of distance measures is limited and often difficult to find. Some reviews of distance measures have been published (Liao, 2005; Lhermitte *et al*., 2011; Esling and Agon, 2012; Montero and Vilar, 2014; Mori *et al*., 2016), but are not generally aimed at ecologists (but see Lhermitte *et al*., 2011); analysis of the properties of distance measures is limited, and guidance of how to choose an appropriate distance measure is either missing or very general. Other studies have analysed the classification accuracy of multiple distance measures across a variety of datasets (Wang *et al*., 2013; Pree *et al*., 2014; Bagnall *et al*., 2017; Paparrizos *et al*., 2020), but pooled the results to give overall performance scores. This ignores the fact that different distance measures perform better on different datasets and for different tasks. Kocher and Savoy (2017) tested 24 distance measures for six properties, then compared their effectiveness in classification on 13 real-world datasets. However, the study focused on a single task (author profiling, i.e., determining demographic information about the author of a document based on the document itself) and did not present a general method for selecting distance measures for other tasks. Furthermore, the distance measures that demonstrated all proposed properties did not perform best on real-world datasets. Mori *et al*. (2015) developed an automated process for selecting distance measures based on nine quantifiable properties of datasets. However, their classifier is limited to clustering tasks, and only includes five common distance measures. We are not aware of any more generalized method of distance measure selection.

In this study, we present a generalized, objective, user-driven method of choosing fit-for-purpose distance measures for time-series comparison tasks (see Figs 5-6 and Table 1). We evaluate 42 distance measures for 16 properties related to time series comparison. We then demonstrate our selection method by applying it to a set of real-world UK bird population trends from a study of the effectiveness of conservation measures (Jellesmark *et al*., 2021). Finally, we discuss how to select appropriate distance measure(s) for any dataset and task.

**Table 1.** Solutions to potential issues in the data. Note that choice of invariance or sensitivity as a solution should depend on whether the difference in question is important.

| Problem | Pre-processing solution | Properties-based solution |
| --- | --- | --- |
| Missing data points | Interpolate missing values. | Choose an elastic distance. They handle gaps through one-to-none or one-to-many point matching. |
| Different starting values but similar value scales | Apply a translation shift. | Choose a distance measure invariant (or sensitive) to translation. |
| Different value scales | Normalize or standardize data. |  |
| Zeros or negative values | Transform data to obtain positive values. | Choose a distance with non-positive value handling. |
| Noise | Apply a smoothing algorithm. | Choose a distance measure robust (or sensitive) to the type of noise that is of concern. |
| Out of phase |  | Choose a phase invariant (or phase sensitive) distance measure. |
| Unequal lengths | Cut all time series to the same length. | Choose an elastic, model-based or compression-based distance measure. |
| Different time scales |  | Choose a distance measure invariant (or sensitive) to uniform time scaling. |

## 2. Methods

We selected 42 distance measures from the literature (see supplementary materials Table S1). We chose measures that had already been implemented in publicly accessible R packages, and that represented each of the categories we defined in the introduction, as well as a variety of potential use cases. Eighteen of the distance measures we selected are implemented in the R package ‘TSclust’ and have been studied for use in clustering time series (Montero and Vilar, 2014). The other twenty-four are implemented in the R package ‘philentropy’ (Drost, 2018).

We defined a set of 16 properties of distance measures that may be of interest in time series comparison: four metric properties, six value-based properties, five time-based properties, and one uncategorized property. Metric properties define whether dissimilarity is measured in metric space (a space that has real physical meaning). Distance measures that do not demonstrate all the metric properties (semi-metrics and non-metrics; McCune et al., 2002) are useful, but less intuitive (e.g., negative distances, or distances between identical objects may be non-zero). Value-based properties focus on dissimilarities on the y-axis (differences in values; Figs 1-2), while time-based properties focus on dissimilarities on the x-axis (differences in time; Fig. 1).

**Figure 1.**
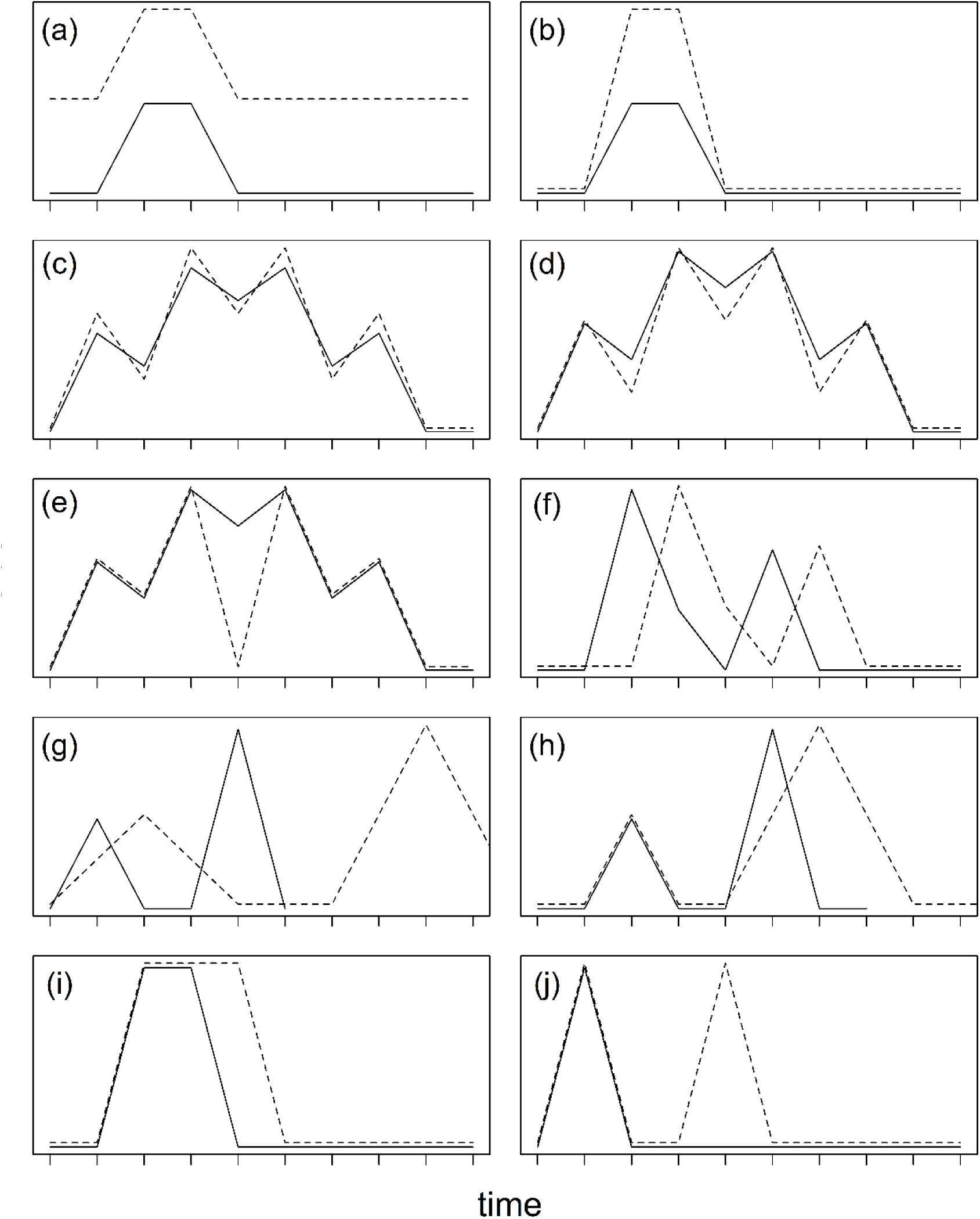
Illustration of time series distortions used to demonstrate sensitivities or invariances of distance measures to: a) translation; b) amplitude; c) white noise; d) biased noise; e) outliers; f) phase; g) time scaling; h) warping; i) duration; and j) frequency. A dissimilarity value of zero (or equivalent, for any distance measure not demonstrating uniqueness) between any of the illustrated pairs of time series would indicate an invariance to that type of distortion.

### 2.1. Metric properties (adapted from McCune *et al*., 2002)

M1. Zero distance. d(X, X) = 0. Identical time series should have a dissimilarity value of zero.

M2. Symmetry. d(X, Y) = d(Y, X). The dissimilarity value should be the same regardless of the order in which time series are compared, X to Y or Y to X. A distance measure without symmetry might, for example, cluster a collection of time series differently depending on how the time series are ordered.

M3. Triangle inequality. d(X, Y) ≤ d(X, Z) + d(Y, Z). Given three time series, the distance between any pair of them should never be larger than the sum of the distances between the other two pairs of time series. This property is related to Euclidean geometry (one side of a triangle cannot be longer than the other two combined). A distance measure that does not obey the triangle inequality is less intuitive to interpret.

M4. Non-negativity. d(X, Y) ≥ 0. The dissimilarity value should never be less than zero.

### 2.2. Value-based properties

V1. Translation invariance (also called amplitude shifting invariance or offset invariance; Fig. 1a). d(X + q, Y) = d(X, Y), where q is any real number (Batyrshin *et al*., 2016). If we increase the value of all observations of one time series by the same amount q, the dissimilarity value should not change. We can further define translation *sensitivity*, where the dissimilarity between X and Y increases relative to the value of q, and translation *insensitivity*, where the dissimilarity between X and Y increases by an amount that is independent of q. Translation sensitivity can be measured in relative terms, allowing comparison between distance measures.

V2. Amplitude sensitivity (Fig. 1b). Translation sensitivity can be defined on a local scale (sensitivity to translation of a section of a time series) and in that case will be referred to as amplitude sensitivity.

V3. White noise invariance (invariance against random noise; Fig. 1c). d(X + f(X), Y) ≈ d(X, Y), where f(X) is a function that adds a small pseudo-random value from a normal distribution with a mean of zero and standard deviation q to each observation of time series X (adapted from Lhermitte *et al*., 2011). Adding a random noise term to one time series from a pair should have an inconsequential effect on the dissimilarity value between them. A distance measure sensitive to white noise will show an increase in dissimilarity values relative to q, allowing us to obtain a relative measure of robustness against white noise. Robustness against white noise might be desirable, e.g., when comparing trends of stochastic processes, such as population growth.

V4. Biased noise invariance (invariance against non-random noise, i.e., noise in a single direction; Fig. 1d). d(X + g(X), Y) ≈ d(X, Y), where g(X) is a function that adds a small non-random value q to half of the observations (randomly chosen) of time series X (adapted from Lhermitte *et al*., 2011). Biased noise is different from random noise in that it is in a single direction and therefore more likely to be systematic or have important meaning.

V5. Outlier invariance (Fig. 1e). d(X + h(X), Y) ≈ d(X, Y), where h(X) is a function that adds a large pseudo-random value q to a single randomly chosen observation of time series X. Outlier sensitivity is thus defined as the dissimilarity value increasing with q, and is a specific case of amplitude sensitivity limited to a single time point. Sensitivity to outliers is useful for detecting anomalies or disruptive events, but robustness may be preferred where outliers represent measurement errors or irrelevant anomalies.

V6. Antiparallelism bias (see Fig. 2). Antiparallelism refers to line segments or trends which have slopes with the same value but opposite signs, while parallelism refers to those which have identical slopes in both value and sign. A distance measure with positive antiparallelism bias ignores the sign of the slope and treats antiparallel and parallel trend curves the same. A distance measure with negative antiparallelism bias treats trend curves with opposite signs as more dissimilar than those with identical signs. Distance measures with no antiparallelism bias (neutral) measure absolute differences on the y-axis, without respect to slope or direction. Whether and which kind of antiparallelism bias is desirable depends on the application. For example, a negative antiparallelism bias might be desirable if one is more concerned with the direction of population trends than their slope.

**Figure 2.**
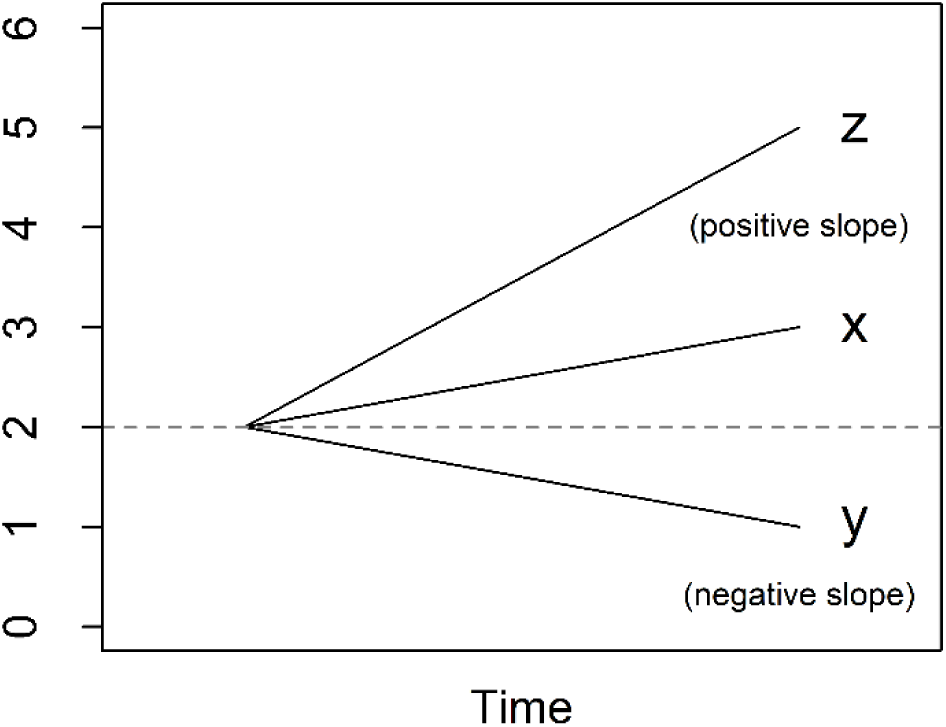
Illustration of antiparallelism bias. Time series x and y are antiparallel (y has the same slope as x, but in the opposite direction), while z has a different slope than x, but in the same direction. The total difference in values between x and z is the same as that between x and y. Distance measures with positive antiparallelism bias rate time series x as more dissimilar to time series z than to time series y, while the opposite is true for those with negative antiparallelism bias. Distance measures with neutral antiparallelism bias rate the time series pairs as equally dissimilar.

### 2.3. Time-based properties

T1. Phase invariance (Fig. 1f). d(X_i+p_, Y_i_) = d(X_i_, Y_i_) (adapted from Lhermitte *et al*., 2011). Phase invariance is the x-axis equivalent of translation invariance. If all observations of X are shifted horizontally by the same value p, it should not affect the dissimilarity value. Phase invariance may be a desirable property to detect similarities that occur separated in time. For example, when matching audio recordings of bird songs, it is likely that similar songs occur at different time points in different recordings. Conversely, when comparing population trends of different species within a community or geographical area to see which ones responded similarly to a disruptive event occurring at time t, phase invariance is not a desirable property as responses should match in time.

T2. Time scaling invariance (Fig. 1g). d(X_pi_, Y_i_) = d(X_i_, Y_i_) (adapted from Esling and Agon, 2012). If one time series is expanded or compressed along its time axis, the dissimilarity value should not change. This property is useful for certain applications, such as comparing animal behaviour patterns occurring at different speeds.

T3. Warping invariance (Fig. 1h). Time scaling invariance can be defined locally, i.e., involving the expansion or compression of one or more sections of a time series, rather than the entire series (Batista *et al*., 2011). Warping invariance is particularly useful when matching similar time series which have plateaus or valleys of uneven lengths.

T4. Frequency sensitivity (Fig. 1i). If time series Y is obtained by applying the same transformation j(t) to one or more observations t of time series X, such that d(X, Y) > d(X, X), then the dissimilarity value will depend on the number of observations to which the transformation j(t) is applied. In other words, if a distance measure is sensitive to frequency, increasing the number of differences between two time series should increase the dissimilarity value.

T5. Duration sensitivity (Fig. 1j). If time series Y is obtained by applying the same transformation k(t) to one or more consecutive observations of time series X, such that d(X, Y) > d(X, X), then the dissimilarity value will depend on the number of consecutive observations to which the transformation k(t) is applied. This property is a special case of frequency sensitivity. Distance measures which are sensitive to duration must be sensitive to frequency, but the converse is not true.

### 2.4. Other properties

N1. Non-positive value handling. Some distance measures will not return results if the data contains negative values or zeros. This has implications e.g., for tasks such as classification, where it is common to first perform min-max normalization to rescale time series values to [-1,1].

### 2.5. Metric properties tests

The metric properties of some distance measures are specified in the literature, but for others it is unclear. Therefore, we devised a set of tests for metric properties (see supplementary materials for details). We confirmed the robustness of our tests by comparing our results to the literature for distance measures with known metric properties.

### 2.6. Time-based and value-based properties tests

We performed two types of testing for non-metric properties in this study. Controlled testing was performed on sets of short, simple time series to clearly demonstrate specific properties. However, the demonstrated properties may not translate as clearly onto real-world datasets, and the behaviour of distance measures may vary depending on the types of time series involved (see Lhermitte *et al*., 2011). Therefore, we employed uncontrolled testing by applying functions to real-world time series to induce differences, then comparing the altered time series to their unaltered counterparts. We applied the functions over a range of parameters, then plotted the resulting curves to show how responses of distance measures vary with magnitude. For full details, see supplementary materials.

### 2.7. Controlled testing

We created sets of short time series to demonstrate each property. We devised tests for all value-based and time-based properties (see supplementary materials for details) and applied the tests to all distance measures. For V1-V5, T4, and T5, we separated the resulting values into five bins, which we designated as “very low,” “low,” “medium,” “high,” or “very high.” For T1-T3, results were not binned. Distance measures were designated “sensitive” for a given property if the distance was directly dependent on the phase difference or degree of scaling or warping. For all sensitivities and invariances, distance measures were classified as “invariant” if they returned zero values for all time-series pairs, “insensitive” if the same non-zero value was returned for all time-series pairs, or “unpredictable” if distance values varied but did not show a clear relationship. All measures that were unable to handle unequal-length time series were designated “n/a” for uniform time scaling invariance and warping invariance.

Antiparallelism bias was tested by comparing pairs of time series that differed by the same relative amount in different directions. Distance measures were designated as “positive” bias if they gave a greater dissimilarity value to pairs of time series differing in opposite directions than to pairs differing in the same direction, “negative” bias if they gave a greater dissimilarity value to those differing in the same direction, or “neutral” if they assigned each pair of time series the same dissimilarity value.

### 2.8. Uncontrolled testing

We created a function for each property to be tested, which applies a transformation to one or more time points of a real-world time series. Each function accepts a value q, the purpose of which varies depending on the function (see supplementary materials for details). For example, the translation function adds a real number q to every value of a time series. The transformed time series is returned as output and compared against its unaltered counterpart. We applied the functions to a range of q in increments, then graphed the results as response curves (see Figs S5-S8 in supplementary materials). We did not compare them against a reference or assign sensitivity ratings, as they were intended only as a confirmatory check against the results of controlled testing.

### 2.9. Selection process

We devised a selection process to guide researchers through determining the most appropriate distance measure(s) for their intended application. First, use the decision tree (Figs 5-6) to select a general category of distance measures. Next, use Table 1 to determine which pre-processing steps might be necessary to prepare the dataset and/or to further narrow the choice of distance measures. Finally, determine which properties will be most important to achieve the desired outcome and use Figs S1-S3 (see supplementary materials) to narrow the selection to the distance measures which exhibit these properties. We demonstrate the selection process on a real-world dataset.

### 2.10. Example datasets

We used a dataset from a study of conservation impact of wet grassland reserves on breeding birds in the UK (Jellesmark *et al*., 2021). The dataset consists of 25 years of breeding pair count data for five wading bird species, from within and outside of reserves. The within-reserves data came from 47 RSPB lowland wet grassland reserves, while the counterfactual (outside of reserves) data was selected from the UK Breeding Bird Survey data. Data were matched to select sites that represent how reserve land would look in the absence of conservation measures. The reserve and counterfactual count data were aggregated into species trends, then converted to indices by dividing each annual species count total by the first-year species count total. Thus, each of the five bird species was represented with a reserve trend index and a matched counterfactual trend index. Jellesmark *et al*. (2021) compared each pair of indices to determine the effects of conservation efforts on each bird species, by calculating the percentage improvement of reserve indices over counterfactual indices and performing t-tests to determine significance and effect size of the difference. We ranked the results of Jellesmark *et al*. (2021) according to both percentage improvement and effect size. We then applied our selection method to select appropriate distance measures, ranked the dissimilarity results returned by each selected distance measure, and examined the rankings with respect to Jellesmark *et al*. (2021). We also ranked the results returned by rejected distance measures as a reference (see supplementary materials).

## 3. Results

### 3.1. Metric test results

Fourteen out of 42 distance measures were identified as full metrics, meaning they passed the metric tests for uniqueness, symmetry, non-negativity, and the triangle inequality (see Fig. S1). Sixteen distance measures were identified as semi-metrics (failed the triangle inequality test but passed the other three tests) and 12 were identified as non-metrics (failed at least one of the tests for uniqueness, symmetry, or non-negativity; Fig. S1). However, in some cases results depended on settings or input values (some distance measures passed the triangle inequality and/or non-negativity tests only when inputs were constrained to non-negative real numbers). All tested feature-based and model-based distances were full metrics, while all tested compression-based distances were non-metrics. Shape-based measures showed mixed results, even within families and groups.

### 3.2. Sensitivity test results

Lock-step shaped-based measures varied in the strength of responses to the sensitivity tests, but none tested as unpredictable and only two (the Chebyshev distance and the Short Time Series, or STS, distance) showed any invariances or insensitivities. There were no clear differences between families of distance measures, with responses seeming to vary as much within families as between them. Elastic, feature-based and model-based distances showed greater variation in responses, with insensitivities, invariances, and unpredictability being common. The two compression-based distances we tested responded unpredictably to all controlled tests except translation and outliers; they responded unpredictably to *all* uncontrolled tests without exception. See supplementary materials for more detailed results.

### 3.3. Time-based invariances and other test results

All distance measures except the Time Alignment Measurement (TAM) distance responded unpredictably to phase invariance testing. TAM was sensitive to phase changes, however the response curve in uncontrolled testing was not smooth, suggesting some level of unpredictability. The Edit Distance with Real Penalty (ERP) distance was sensitive to uniform time scaling, while all other distances either responded unpredictably or were unable to be tested due to an inability to handle unequal-length time series. Warping sensitivity was more common, occurring in three elastic distance measures. DTW tested as invariant to warping and was thus the only distance measure we tested with any time-based invariances. Elastic measures were the only group of distance measures that showed any predictable time-based sensitivities or time-based invariances.

Two distance measures in the Shannon’s entropy family were unable to deal with zeros, while the entire family was unable to deal with negative values. Three other lock-step shape-based measures also showed an inability to deal with negative values. Antiparallelism bias showed no obvious group-based patterns, but negative antiparallelism bias was most common and positive bias was least common.

### 3.4. Selection process

We began by examining our wading bird dataset in context of the decision trees in Figs 5-6. It consisted exclusively of short (25 data points), non-stationary time series. Following Fig. 5, we focused on shape-based distance measures, which compare raw data values. As the time series were of equal-length, in phase, using the same time scale, and without any missing data points, both lock-step and elastic measures would be appropriate (Fig. 6).

Next, we worked through Table 1. As our wading bird trends were indexed to a starting value of one (Fig. 3), they had the same starting value and the same value scale. There were no negative values because the trends were indexed and based on wetland bird counts; nor were there any zeroes. However, we did notice that some of our time series were noisy (Fig. 3), which could obscure the trends. Noise is a common characteristic of population data, largely due to the stochasticity of population dynamics and the environmental variables they depend on (Vasseur and Yodzis, 2004). While this noise is often white (random, uncorrelated), biased ‘red’ noise (positively autocorrelated, tending toward a single direction) is also common, e.g., when environmental conditions are above or below average for an extended period (Vasseur and Yodzis, 2004; van de Pol *et al*., 2011). Biased noise is therefore more likely to represent a legitimate difference in trends. There are multiple ways to deal with noisy time series (Table 1). We first tried the properties-based solution (Table 1; see below for the pre-processing solution). Using Fig. S2, we filtered out all shape-based distance measures with a white noise sensitivity category of medium or higher (a sensitivity value of 0.7 or more). Next, we required biased noise to be at least two categories higher in sensitivity than white noise (Fig. S2; e.g., if white noise sensitivity was very low, biased noise sensitivity must be at least medium). Our choices here were based on practicality; sensitivity categories are arbitrary (we categorized them for convenience), so we wanted to avoid being too specific while ensuring that any chosen distance measure exhibited a non-trivial difference in sensitivity between white noise and biased noise.

**Figure 3.**
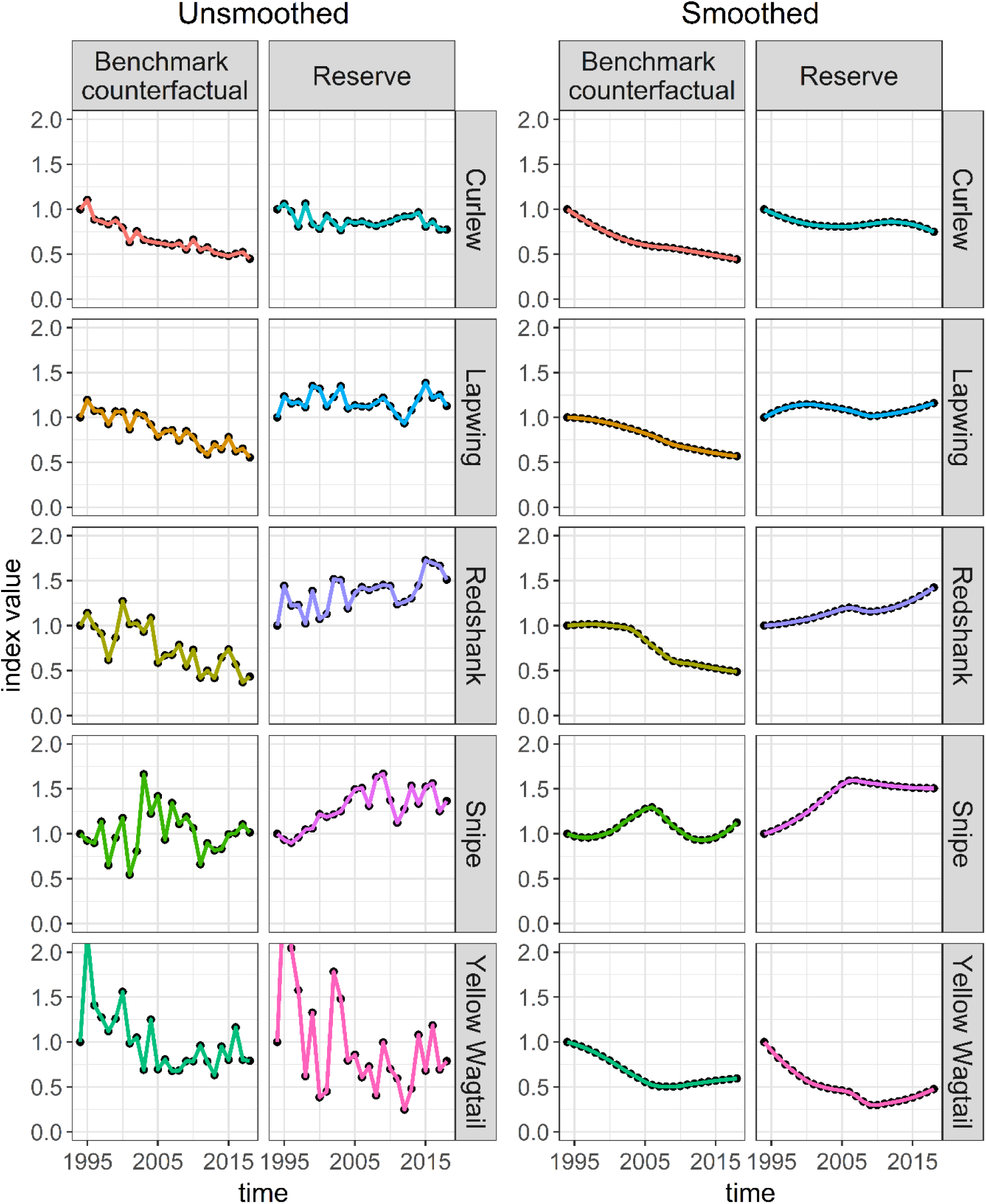
Reserve and counterfactual trends for five wading bird species that breed on RSPB lowland wet grassland reserves in the UK. Left: Unsmoothed trends based on original data presented in Jellesmark et al. (2021). Right: LOESS smoothed trends with a span setting of 0.75.

Finally, we considered the remaining properties in the context of our intended task and desired outcome. We deemed amplitude sensitivity to be important, as we were interested in the overall divergence between population indices within and outside reserves. Duration sensitivity was also important, as we would consider population indices which diverge more steeply or for a longer period to be more different, i.e., that conservation measures had a stronger effect on these species. Therefore, both amplitude and duration sensitivity had to be at least low (a sensitivity value of 0.2 or higher; Fig. S2). Again, we could have chosen a different (higher) category, but we were more concerned with making sure the distance measures exhibited *some* sensitivity to these properties than the exact degree of sensitivity. We did not filter for antiparallelism bias, as the high stochasticity in some of our time series (Fig. 3) would dilute the signal too much for it to matter.

This selection process left us with two distance measures: the K-Divergence (KDiv) and the Kullback-Leibler distance (Kullback), both of which returned the same rankings that Jellesmark *et al*. (2021) obtained using percent improvement (Fig. 4). Only one of the 40 unselected distance measures returned the same rankings. Results from unselected distance measures are in supplementary materials S10 and S15.

**Figure 4.**
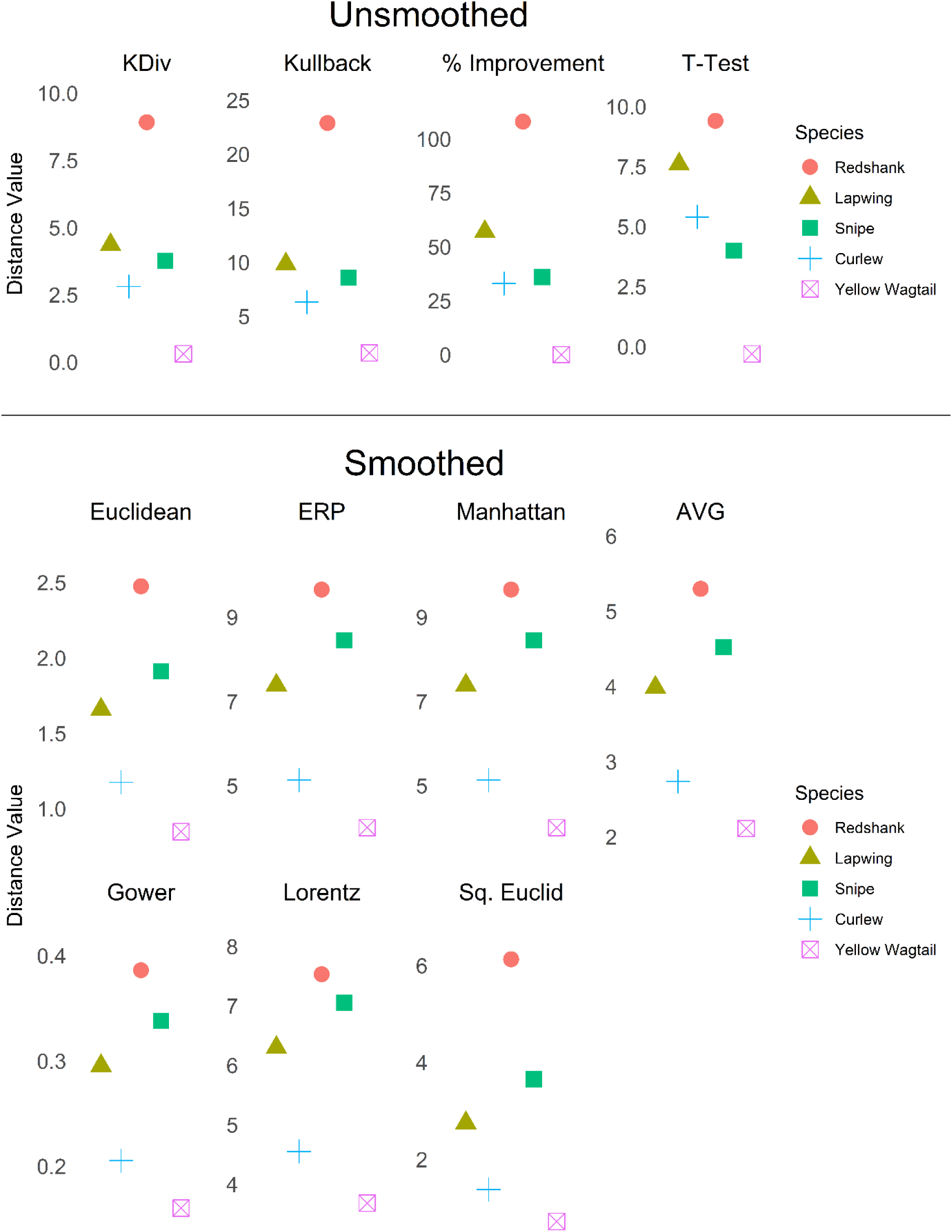
Comparative rankings of conservation impact on five wading bird species. Values on the y-axis represent the distance between unsmoothed (top) or LOESS smoothed (bottom) reserve and counterfactual trends for each species. Results are from the distance measures chosen by our selection process, as well as the percent improvement and t-test methods (top) used by Jellesmark *et al*. (2021).

Another way of dealing with noisy time series is by applying a smoothing algorithm (Table 1). We applied a LOESS smoothing algorithm (span = 0.75) to all time series in the dataset to remove the noise and reveal the trends (Fig. 3). We then re-ran the selection process using the same settings, except that we did not filter for noise sensitivity, and we added a filter for antiparallelism bias. Antiparallelism bias is not very important when dealing with highly stochastic time series because the signals for slope and direction are muddied by noise; however, smoothing introduces strong positive autocorrelation, making the slope and direction signals clear. We selected neutral for antiparallelism bias (Fig. S3) because we were more interested in relative differences in the population indices than the direction of change.

We were left with seven distance measures: ERP, the Euclidean distance, the Manhattan distance, the Gower distance, the Lorentzian distance (Lorentz), the Average distance (AVG), and the Squared Euclidean distance (Sq. Euclid). All seven selected distance measures agreed on the following order: Redshank, Snipe, Lapwing, Curlew, Yellow Wagtail (Fig. 4). Four of the 35 unselected distance measures returned the same results. See supplementary materials S10 and S15 for complete results from unselected distance measures.

## 4. Discussion

The aim of this study was to provide enough information to make informed, objective decisions about which distance measures to use. We tested 42 distance measures for 16 properties and presented an objective method of selecting distance measures for any task based on those properties. We demonstrated the viability of the method on a real-world dataset by selecting distance measures to rank differences between pairs of wading bird population trends (within and outside of reserves) and showing that the distance measures we selected were fit-for-purpose and consistent in their rankings. The method is user-directed; therefore, success depends on an understanding of the dataset, the task to be performed, and the hoped-for outcome.

Time series length and stationarity inform what category of distance measures the user should focus on (Fig. 5). Shape-based distances are best for short time series with differences that are easy to visualize, while longer, stationary time series may be better suited to feature-based, model-based, or compression-based distance measures (Esling and Agon, 2012).

**Figure 5.**
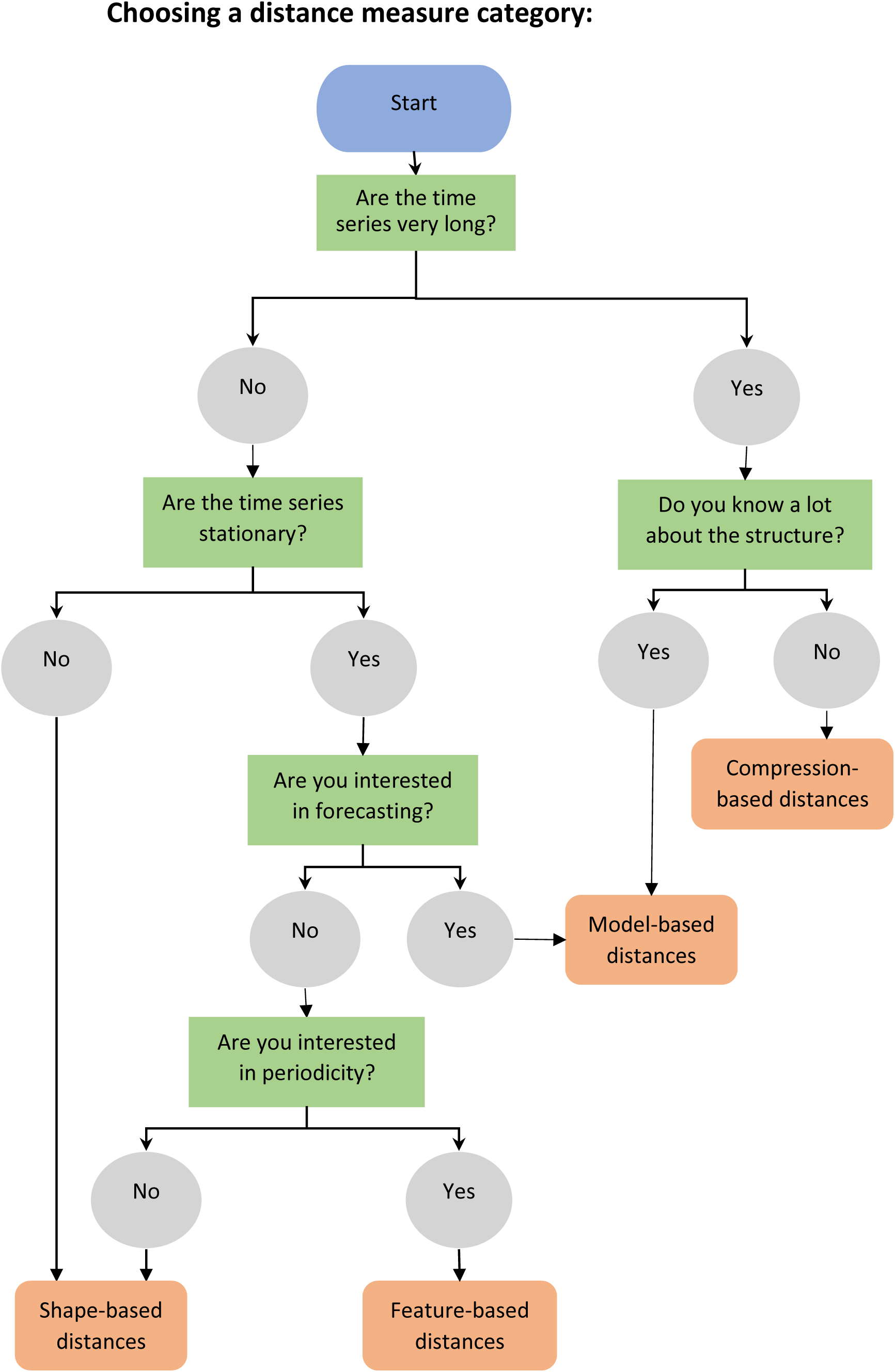
Decision tree to aid in choosing a distance measure category.

The results of our properties tests showed a variation in strength of sensitivity to different properties in different distance measures (Fig. S2), although most distance measures were highly sensitive to outliers (Fig. S2). Invariances were uncommon among the distance measures we tested (Fig. S2 and S3), although several distance measures did demonstrate invariance to translation (Fig. S2). Some distance measures, such as the Edit Distance for Real Sequences (EDR) and ERP, have settings that may affect their behaviour. In the case of ERP, settings can determine whether and how sensitive it is to missing values, while in the case of EDR, the threshold setting determines how far apart values must be to be considered different, and therefore serves to toggle responses to multiple properties between invariance and sensitivity.

When dealing with time series of unequal length or missing data points, distance measures that allow unequal matching (e.g., matching multiple points to one point), such as DTW, or that allow gaps, such as ERP, may be the solution. Alternatively, pre-processing of data may remove such concerns. For example, missing data points can be filled in by interpolation, or longer time series can be cut to the same length as shorter ones (only attempt such solutions if they make sense for the data).

Elastic measures, such as DTW, EDR, and ERP, are the most versatile distance measures, able to handle many common complications of datasets with little or no pre-processing. For general tasks, they are often a good option (see our decision tree: Figs 5-6). However, for tasks involving large datasets containing thousands of time series, some elastic measures may be impractical due to processing speed. Much of the research into speeding up time series comparisons for large datasets has focused on a select few distance measures, especially the Euclidean Distance and DTW. While the Euclidean Distance is faster, better known, and still widely used in some fields, an extensive body of research has shown DTW to be more accurate (Zhu *et al*., 2012; Dau *et al*., 2019; Paparrizos *et al*., 2020) and it is considered the *de facto* standard for accuracy in classification (note that it is still important to consider the properties of DTW in relation to the data, as it does not perform well in every case). Despite this, it is rarely used in ecology (Hegg and Kennedy, 2021). Note, however, that DTW is computationally expensive and therefore can be slow for large datasets (for discussion on ways to speed up DTW, see supplementary materials S11).

**Figure 6.**
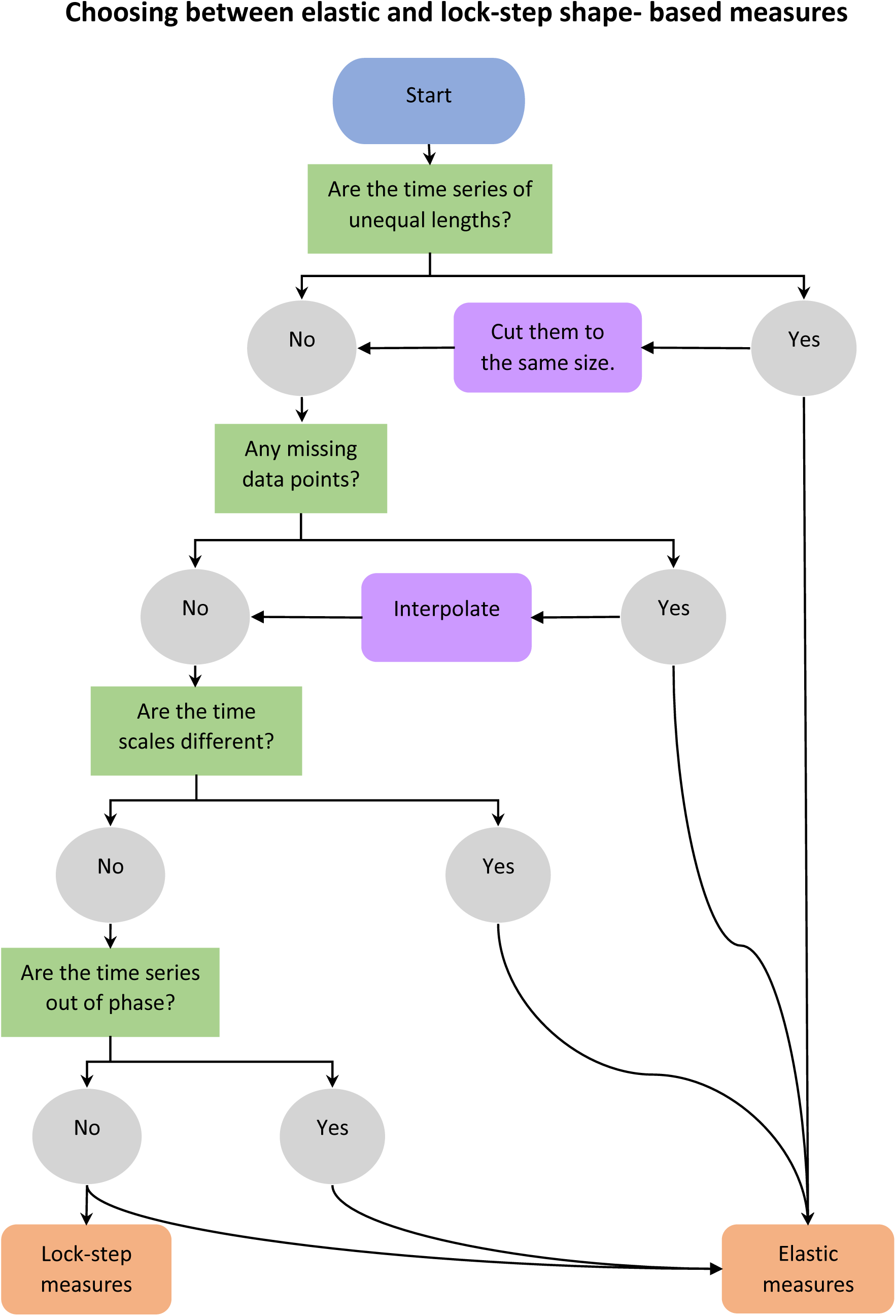
Decision tree to aid in choosing a sub-category of shape-based distance measures.

For many analyses involving distance measures, researchers may first want to normalize or standardize their data or translate it along the y-axis. This may be an important step if the time series use different scales or have different starting values. For example, when performing classification or clustering tasks, it is common to apply z-normalization to rescale time series to a mean of zero and standard deviation of one (Rakthanmanon *et al*., 2013). Min-max normalization to a scale of [0,1] or [-1,1] is also common for datasets that are not normally distributed. Be aware, however, that these transformations may affect the subsequent choice of distance measures, as some cannot handle zeros or negative values and some metrics are non-metric when there are negative values present (see Fig. S1).

Although we ignored the metric properties of distance measures for our real-world example, they are very important for some tasks. For example, many algorithms for classification and clustering are designed to work only in metric space and may return unexpected results for non-metric distances (Weinshall *et al*., 1999).

Noise is a common aspect of ecological time series, as environmental and population dynamics are stochastic. There are several potential ways to deal with noisy time series. Some distance measures, such as EDR, have threshold settings; any difference between time series that falls below the threshold will be ignored. If the noise is relatively uniform in amplitude, this may be a simple solution if the distance measure in question meets all other requirements. Other distance measures, such as KDiv, are relatively robust against white noise although lacking a sensitivity setting, and may be more appropriate if the noise is less uniform. A more drastic solution is to apply a smoothing algorithm as a pre-processing step, though this should be approached with caution. Smoothing will remove noise and outliers but may distort the time series in the process. Therefore, it is important to avoid over-smoothing. Smoothing time series that have sudden and/or drastic value changes may also be problematic, particularly if these changes are an important aspect of differentiation between time series.

Our demonstration using data from Jellesmark *et al*. (2021) served to illustrate both the potential benefits and complications introduced by smoothing. When we filtered by noise sensitivity, we were left with two distance measures; both returned the same results as the percentage difference calculations by Jellesmark *et al*. (2021). When we ran the method after applying a smoothing algorithm, we were left with a larger choice of seven distance measures. Although the ordering differed slightly from Jellesmark *et al*. (2021), all seven distance measures agreed. The slight difference in ordering (Snipe vs Lapwing, ambiguous from visual inspection of the trends; Figs 3-4) is unsurprising given that the smoothing algorithm removed all noise from the trends, while the distance measures we selected using noise filtering, although demonstrating very low sensitivity to white noise, were not invariant to it. Smoothing in this case gave us more distance measures to choose from, but with the added complication of not knowing whether we had improved or distorted our results.

While in both cases (smoothed and unsmoothed trends) there were distance measures that gave the same rankings as Jellesmark *et al*. (2021) despite not matching our selection criteria (see supplementary materials S10), the distance measures we selected were all in agreement. Had we been less specific when choosing important properties, we would have risked including measures that were not fit-for-purpose. A single suitable distance measure is better than any number of ill-suited measures.

## 5. Conclusion

Distance measures are widely used in ecology, but the selection of distance measures described in the ecological literature is limited and their use is often poorly understood, leading to misuse. In the wider literature, there are hundreds of distance measures, with new ones frequently described. This study introduces a selection of 42 distance measures for the purpose of ecological time series analysis and describes an objective method for choosing an appropriate distance measure for any task involving time series. This should lead to an improved understanding of, and greater scope for, the use of distance measures for comparing time series within the field of ecology. Nonetheless, it is up to the user to think their way through the process. There are hundreds of potential cases for using distance measures to compare time series in ecology, and as many potential issues that may arise in the process. Most of them are beyond the scope of this study. However, we hope that we have covered the basics and provided enough data and theory on distance measures and their properties to help select one that is appropriate for the task. There is not always a right choice of distance measure, but there are wrong ones, and our main goal is to help avoid those.

## Authors’ Contributions

SD, DM, RF, and MB conceived the ideas; SJ produced the wading bird indices; SD designed the methodology, wrote the code, produced the simulated data, analysed the data, and wrote the manuscript, with input from all authors; all authors contributed critically to the drafts and gave first approval for publication.

## Acknowledgements

We thank Gonzalo Albaladejo-Robles and Bouwe Reijenga for their support. This project has received funding from the European Union’s Horizon 2020 research and innovation programme under the Marie Skłodowska-Curie grant agreement No 766417.

## Data availability

We used data from multiple sources, as well as simulated data, for this study. R scripts to recreate all simulated data and reproduce all results are available on github at https://github.com/shawndove/Trend_compare, and will be versioned and archived at Zenodo upon acceptance of the manuscript. Wading bird indices produced from data provided by the RSPB and UK Breeding Bird Survey will be archived at Zenodo upon acceptance of the manuscript. Datasets from the UCR Time Series Classification Archive are available at https://www.cs.ucr.edu/~eamonn/time_series_data_2018/.

