## Supplementary Materials for "A User-Friendly Guide to Using Distance Measures to Compare Time Series in Ecology"

### Contents

1. Distance measures included in our study ... 2
2. Descriptions and formulas for selected distance measures ... 5
3. Distance measure properties ... 10
4. Metric testing ... 12
5. Controlled testing ... 14
6. Uncontrolled testing ... 17
7. Metric test results ... 19
8. Controlled test results ... 21
   8.1. Sensitivity tests ... 21
   8.2. Time-based invariances and other tests ... 21
9. Uncontrolled test results ... 24
10. Example dataset results ... 30
11. Speeding up DTW ... 33
12. References ... 34
13. Plots of controlled test results for all distance measures ... 37
14. Tables of controlled test results for all distance measures ... 79
15. Plots of wading bird rankings for all distance measures ... 121

### 1. Distance measures included in our study

Table S1: Distance measures we tested (names are not necessarily given by the original authors, as some authors did not name their distance measures), with abbreviated names, type or family, category, important and/or interesting characteristics, and number of parameters that can or must be set, as well as sources for additional information. Parameters are provided as a range if one or more parameters are optional (e.g., some can be set to “NULL”, while others are only sometimes relevant, depending on the input data). Note that we may list one or more parameters for distance measures that are considered parameter-free if they require the user to choose a linked component, e.g., a compression algorithm.

| Distance Measure | Abbreviated Name | Type/Family | Category | Characteristics | Parameters | Source |
|---|---|---|---|---|---|---|
| Euclidean Distance[1,2] | Euclidean | Norm distance ($L_p$ Minkowski family) | Lock-step Shape-based | Shortest distance between points in Euclidean space | 0 | (Cha, 2007) |
| Manhattan Distance[1,2] | Manhattan | Norm distance ($L_p$ Minkowski family) | Lock-step Shape-based | Shortest distance between points on a grid | 0 | (Cha, 2007) |
| Chebyshev Distance[1,2] | Chebyshev | Norm distance ($L_p$ Minkowski family) | Lock-step Shape-based | Takes maximum distance between point pairs (all other point pairs are ignored) | 0 | (Cha, 2007) |
| Complexity-Invariant Distance[1] | CID | Correction factor | Lock-step Shape-based | Invariant to complexity | 1 | (Batista et al., 2011) |
| Dynamic Time Warping Distance[1] | DTW | Dog-man Distance | Elastic Shape-based | Warping path | 0-4 | (Sakoe and Chiba, 1978) |
| Time Alignment Measurement Distance[1] | TAM | Dog-man Distance | Elastic Shape-based | Warping path<br>Time distortion penalty | 0 | (Folgado et al., 2018) |
| Normalized Compression Distance[1] | NCD | Compression distance | Compression-based | Normalization of differences<br>Quasi-universality<br>Choice of compression algorithm | 1 | (Cilibrasi and Vitanyi, 2005) |
| Compression-Based Dissimilarity Measure[1] | CDM | Compression distance | Compression-based | Compatible with symbolic representation<br>Choice of compression algorithm | 1 | (Keogh et al., 2004) |
| Edit Distance with Real Penalty[1] | ERP | Edit distance | Elastic Shape-based | Gap-length penalty | 1-2 | (Chen and Ng, 2004) |
| Edit Distance on Real Sequences[1] | EDR | Edit distance | Elastic Shape-based | Threshold parameter<br>Gap-length penalty | 1-2 | (Chen et al., 2005) |
| Fourier Coefficient Based Distance[1] | Fourier | | Feature-based | Frequency domain | 1 | (Agrawal et al., 1993) |
| Autocorrelation-Based Dissimilarity[1] | ACF | | Feature-based | Compares autocorrelation coefficients | 1-2 | (D’Urso and Maharaj, 2009) |
| Partial Autocorrelation-Based Dissimilarity[1] | PACF | | Feature-based | Compares partial autocorrelation coefficients | 1-2 | (Montero and Vilar, 2014) |
| Periodogram Based Dissimilarity[1] | Per | | Feature-based | Frequency domain | 2 | (Caiado et al., 2005) |

| Distance Measure | Abbreviated Name | Type/Family | Category | Characteristics | Parameters | Source |
|---|---|---|---|---|---|---|
| Integrated Periodogram Based Dissimilarity[1] | IntPer | | Feature-based | Frequency domain | 1 | (Casado de Lucas, 2010) |
| Piccolo Distance[1] | Piccolo | | Model-based | ARIMA models | 0-3 | (Piccolo, 1990) |
| Short Time Series Distance[1] | STS | | Lock-step Shape-based | Captures temporal information | 0-2 | (Möller-Levet et al., 2003) |
| Dissimilarity Index Combining Temporal Correlation and Raw Value Behaviour[1] | Cort | Correction factor | Lock-step Shape-based & Feature-based | Temporal correlation coefficient Adaptive tuning function | 2 | (Chouakria and Nagabhushan, 2007) |
| Gower distance[2] | Gower | $L_1$ family | Lock-step Shape-based | | 0 | (Cha, 2007) |
| Soergel distance[2] | Soergel | $L_1$ family | Lock-step Shape-based | | 0 | (Cha, 2007) |
| Kulczynski distance[2] | Kulcz | $L_1$ family | Lock-step Shape-based | | 0 | (Cha, 2007) |
| Canberra distance[2] | Canberra | $L_1$ family | Lock-step Shape-based | Normalizes | 0 | (Cha, 2007) |
| Lorentzian distance[2] | Lorentz | $L_1$ family | Lock-step Shape-based | | 1 | (Cha, 2007) |
| Wave-Hedges distance[2] | Wave | Intersection family | Lock-step Shape-based | Normalizes | 0 | (Cha, 2007) |
| Czekanowski distance[2] | Czek | Intersection family | Lock-step Shape-based | Normalizes | 0 | (Cha, 2007) |
| Jaccard distance[2] | Jaccard | Inner Product family | Lock-step Shape-based | Normalizes | 0 | (Cha, 2007) |
| Dice dissimilarity[2] | Dice | Inner Product family | Lock-step Shape-based | Normalizes | 0 | (Cha, 2007) |
| Squared-chord distance[2] | SqChord | Fidelity family | Lock-step Shape-based | | 0 | (Cha, 2007) |
| Squared Euclidean distance[2] | SqEuclid | Squared $L_2$ family | Lock-step Shape-based | | 0 | (Cha, 2007) |
| Squared Chi-squared distance[2] | SqChi | Squared $L_2$ family | Lock-step Shape-based | | 0 | (Cha, 2007) |
| Probabilistic Symmetric Chi-squared distance[2] | ProbSymm | Squared $L_2$ family | Lock-step Shape-based | | 0 | (Cha, 2007) |
| Divergence squared distance[2] | Diverge | Squared $L_2$ family | Lock-step Shape-based | Normalizes | 0 | (Cha, 2007) |
| Clark squared distance[2] | Clark | Squared $L_2$ family | Lock-step Shape-based | Normalizes | 0 | (Cha, 2007) |
| Additive Symmetric Chi-squared distance[2] | Additive | Squared $L_2$ family | Lock-step Shape-based | | 0 | (Cha, 2007) |
| Kullback-Leibler (KL) divergence[2] | Kullback | Shannon's entropy family | Lock-step Shape-based | Non-symmetric | 1 | (Cha, 2007) |
| Jeffreys divergence[2] | Jeffreys | Shannon's entropy family | Lock-step Shape-based | Symmetric form of KL divergence | 1 | (Cha, 2007) |
| K divergence[2] | KDiv | Shannon's entropy family | Lock-step Shape-based | Non-symmetric | 1 | (Cha, 2007) |

| Distance Measure | Abbreviated Name | Type/Family | Category | Characteristics | Parameters | Source |
|---|---|---|---|---|---|---|
| Topsoe distance[2] | Topsoe | Shannon's entropy family | Lock-step Shape-based | Symmetric form of K divergence | 1 | (Cha, 2007) |
| Jensen difference[2] | Jensen | Shannon's entropy family | Lock-step Shape-based | | 1 | (Cha, 2007) |
| Taneja difference[2] | Taneja | | Lock-step Shape-based | Uses both arithmetic and geometric means | 1 | (Cha, 2007) |
| Kumar-Johnson distance[2] | Kumar | | Lock-step Shape-based | Uses symmetric chi-squared, arithmetic, and geometric means. | 0 | (Cha, 2007) |
| Average($L_1$,$L_{inf}$)[2] | AVG | | Lock-step Shape-based | Average of the Manhattan and Chebyshev distances | 0 | (Cha, 2007) |

[1]Available in the TSdist R package (Mori *et al.*, 2016).
[2]Available in the philentropy R package (Drost, 2018).

#### 2. Descriptions and formulas for selected distance measures

The **Euclidean distance** (Euclidean), also known as the $L^2$ distance, is the straight-line distance between a pair of points. It also forms the basis for some of the more complicated transformation-based and model-based metrics presented here. It is defined as:

$$d_{Euc} = \sqrt{\sum_{i=1}^{d} | P_i - Q_i |^2} \tag{3}$$

where P and Q are (time series) vectors and $d$ is the length of the vectors.

The **Manhattan distance** (Manhattan), or $L^1$ distance, is the shortest distance between two points on a grid. Because it is not based on Euclidean geometry, there can be multiple paths with the same shortest distance. It is defined as:

$$d_{CB} = \sum_{i=1}^{d} | P_i - Q_i | \tag{4}$$

The **Chebyshev distance** (Chebyshev), or $L^\infty$ distance, is the greatest of the differences between two points or vectors along any coordinate dimension. For example, if two points had the x,y coordinates (0,0) and (3,5), the Chebyshev distance would be 5, the difference between the y coordinates of the two points, as this is greater than 3, the difference between the x coordinates. The Chebyshev distance is defined as:

$$d_{Cheb} = \max_i | P_i - Q_i | \tag{5}$$

The **Complexity-Invariant Distance** (CID) applies a complexity correction factor to the Euclidean distance to increase the dissimilarity value between time series with different complexities (where complexity is the length of a time series if stretched into a straight line—more and greater peaks and valleys means more complexity). It is defined as:

$$d_{CID}\left(\boldsymbol{X}_T, \boldsymbol{Y}_T\right) = CF\left(\boldsymbol{X}_T, \boldsymbol{Y}_T\right) \cdot d\left(\boldsymbol{X}_T, \boldsymbol{Y}_T\right) \tag{6}$$

where d is the Euclidean distance and CF is a complexity correction factor

$$CF\left(\boldsymbol{X}_T, \boldsymbol{Y}_T\right) = \frac{\max\left\{CE\left(\boldsymbol{X}_T\right), CE\left(\boldsymbol{Y}_T\right)\right\}}{\min\left\{CE\left(\boldsymbol{X}_T\right), CE\left(\boldsymbol{Y}_T\right)\right\}} \tag{7}$$

where CE is a complexity estimator

$$CE\left(\boldsymbol{X}_T\right) = \sqrt{\sum_{t=1}^{T-1}\left(X_t - X_{t+1}\right)^2} \tag{8}$$

The **Dynamic Time Warping distance** (DTW) computes a warping path between two time series to align them in time. It can be defined as a man-dog distance, but instead of the shortest leash length, it measures the average leash length. This makes it more robust than the Fréchet distance, as it is less sensitive to outliers and short divergences. The DTW distance is defined as:

$$d_{DTW}\left(\boldsymbol{X}_T, \boldsymbol{Y}_T\right) = \min_{r \in M}\left(\sum_{i=1,..,m}\left|X_{a_i} - Y_{b_i}\right|\right) \tag{9}$$

The **Time Alignment Measurement distance** (TAM) is a derivative of the DTW distance that measures how well two time series align in time. Segments not in phase are penalized, while amplitude differences are not. A dissimilarity value of zero can occur for non-identical series that are perfectly aligned in time.

The **Normalized Compression Distance** (NCD) is based on the concept of Kolmogorov complexity, which is the minimum information needed to generate a string using an algorithm. The Kolmogorov complexity is a measure of randomness of the string. The smaller the value, the less randomness. The NCD applies the concept to a relationship between objects (time series) when a compression algorithm is applied. The greater the advantage in compression (reduction in randomness) gained by multiplying two time series together, the more closely they are related, and therefore the smaller the dissimilarity between them. NCD is defined as:

$$d_{NCD}\left(\boldsymbol{X}_T, \boldsymbol{Y}_T\right) = \frac{C\left(\boldsymbol{X}_T \boldsymbol{Y}_T\right) - \min\left\{C(\boldsymbol{X}_T), C(\boldsymbol{Y}_T)\right\}}{\max\left\{C(\boldsymbol{X}_T), C(\boldsymbol{Y}_T)\right\}} \quad (10)$$

where C represents the compressed size.

The **Compression-based Dissimilarity Measure** (CDM) is a simplified version of the NCD, defined as:

$$d_{CDM}\left(\boldsymbol{X}_T, \boldsymbol{Y}_T\right) = \frac{C\left(\boldsymbol{X}_T \boldsymbol{Y}_T\right)}{C(\boldsymbol{X}_T) C(\boldsymbol{Y}_T)} \quad (11)$$

The **Edit Distance with Real Penalty** (ERP) is an edit distance, meaning it quantifies the number of insert, delete, or replace operations required to turn one string (time series) into another. ERP includes a penalty for gaps between matched substrings based on the gap length. ERP is defined as:

The **Edit Distance for Real Sequences** (EDR) is an edit distance refined for trajectories. It includes a quantization feature as well as the length-based gap penalty of ERP. EDR is defined as:

The threshold feature prevents it from qualifying as a metric (it fails the triangular inequality test).

The **Fourier Coefficient-based distance** (Fourier) calculates the Euclidean distance between Discrete Fourier Transforms of a pair of time series. Fourier transforms extract frequency information by decomposing a signal (time series) into its frequency components (sine and cosine functions). While a time series is visualized as a single graph of amplitude vs time, its Fourier Transform consists of multiple sinusoidal waves, each with a specific, constant amplitude and frequency. Time information is lost. The Fourier Transform works well for stationary time series, as they have periodic repeating signals. However, the loss of time information presents problems for deconstructing non-stationary series, as they change randomly over time.

The **Autocorrelation-based dissimilarity** (ACF) calculates the Euclidean distance between estimated autocorrelation functions of time series. An autocorrelation function of a time series describes the correlation between two values of the time series at different times with a specified lag (delay between the two values). In other words, it describes the correlation of a time series with a time-offset version of itself. It is defined as:

$$d_{ACF}\left(\boldsymbol{X}_T, \boldsymbol{Y}_T\right) = \sqrt{\left(\hat{\boldsymbol{\rho}}_{X_T} - \hat{\boldsymbol{\rho}}_{Y_T}\right)^\top \boldsymbol{\Omega} \left(\hat{\boldsymbol{\rho}}_{X_T} - \hat{\boldsymbol{\rho}}_{Y_T}\right)} \quad (12)$$

where Ω is a matrix of weights and $\rho$-hat refers to estimated autocorrelation vectors.

The **Partial Autocorrelation-based dissimilarity** (PACF) is identical to the ACF except that it uses the partial autocorrelation functions.

The **Periodogram-based dissimilarity** (Per) calculates the Euclidean distance between the periodograms of time series. A periodogram is a method of estimating the power spectrum of a time series, which is equivalent to the Fourier transform of the autocorrelation function. It describes how power is distributed over the frequency components of a time series.

The **Piccolo distance** (Piccolo) calculates the Euclidean distance between the AR(∞) operators, or autoregressive expansions, of invertible ARIMA models of time series. ARIMA is a time series forecasting method. ARIMA models work by describing autoregressive (AR) and moving average (MA) parameters. An autoregressive model explains a value in a time series by one or more previous values plus random error. It is generally written as AR(p), where p is the order of the model. An autoregressive expansion, AR(∞), is thus an AR model of infinite order. A moving average model—written as MA(q), where q is the order—explains a value in a time series by one or more past random errors as well as its own random error term. Invertible ARIMA models are those which can be written simply as autoregressive (AR) models. This is a necessary property to be able to forecast the dependent variable, and is important for the Piccolo distance, since only the AR aspect is used. ARIMA models can be applied to non-stationary time series, but they must first be converted to stationary time series by one or more differencing operations (subtracting each value from the one before it to remove stochastic trends).

The **Short Time Series distance** (STS) measures the difference between the slopes of time series defined as piecewise linear functions. It is intended to incorporate temporal information while ignoring absolute values, to overcome a weakness of many other distances, including the Euclidean distance, which ignore the temporal order of points and the length of sampling intervals. The STS distance is defined as:

$$d_{STS}(X,Y) = \sqrt{\sum_{k=0}^{N-1}\left(\frac{y_{k+1}-y_k}{t_{k+1}-t_k} - \frac{x_{k+1}-x_k}{t'_{k+1}-t'_k}\right)} \quad (13)$$

Equations 3, 4, and 5 are copied from Cha (2007), equations 6 through 12 are copied from Montero and Vilar (2014), and equation 13 is copied from Mori *et al.* (2016).

#### 3. Distance measure properties

Here we've included some additional explanation for translation invariance, amplitude sensitivity and duration sensitivity.

**Translation invariance**: Translation invariance is a shape-preserving property, meaning that a distance measure with this property would treat two time series with identical shapes as equal, even if the mean values were different. The same effect can be achieved by a vertical shift transformation. For example, time series X can be transformed by adding the same real number q to each observation, X' = X + q, such that time series X and Y have the same starting value (if they already have the same starting value there is no need for transformation). It is a simple matter to apply this transformation to thousands of time series. Note, however, that translation invariance can be problematic. Consider two populations, with population A having a starting size of 100 and population B a starting size of 10,000. If both populations increase by 10 every year for 10 years, population A would now be 200, which means it doubled to twice its original size, while population B would be 10,100, an increase of only 1%. A distance measure with the property of translation invariance (or any distance measure after applying a vertical shift transformation to equalize the starting values) would treat these trends as equal, which makes little sense.

A better way to deal with such comparisons would be a scale transformation, X' = X * q, multiplying each observation of time series x by the same real number q, such that time series X and Y have the same starting value. A scale transformation allows for shape deformation while preserving percentage change. If populations A and B both doubled by increasing linearly for 10 years from 100 to 200 and 10,000 to 20,000 respectively, a scale transformation would result in identical trends, although they did not originally have the same shape. Likewise, in the previous example where two populations of different sizes increase by the same amount but different percentages, a scale transformation would result in trends with very different shapes (slopes). Scale invariance, which is defined as d(X*q, Y) = d(X, Y), with q greater than 0 (Batyrshin *et al.*, 2016), is a rare property of distance measures.

Note that translation invariance is a special case of translation insensitivity, where d(X + q, Y) is independent of q. If we define its opposite, translation sensitivity, we get d(X + q, Y) > d(X, Y), with d(X + q, Y) increasing with q. In other words, if we add a number to all values of X, the dissimilarity between X and Y will increase, and the greater the number we add to X, the greater the increase in dissimilarity. This is a useful property and one that we can measure in relative terms.

**Amplitude sensitivity**: Another variant of translation invariance to consider is local translation invariance/sensitivity (amplitude sensitivity), where if the same vertical shift transformation, $x' = x + q$, is applied to one or more time points x of time series X to form time series Y, either $d(X, Y) = d(X, X)$ (invariance) or $d(X, Y) > d(X, Y)$ with $d(X, Y)$ increasing with q (sensitivity). Amplitude sensitivity is particularly relevant when comparing time series which have been scale transformed or vertical shift transformed to have the same starting value. It is a common property of distance measures and generally desirable. But some distance measures, especially among edit distances, are insensitive to amplitude. For example, the Edit Distance for Real Sequences (EDR) has a threshold value that can be set. Only differences that exceed the threshold are counted. This can be useful when looking for aberrations. For example, time series of maximum daily temperatures could be ranked according to how often they exceed a baseline by a set number of degrees.

**Duration sensitivity**: Some distance measures, such as Dynamic Time Warping (DTW) or the Short Time Series Distance (STS), may rank time series with more differences as more dissimilar *only if those differences are separated by similarities*. Consider a time series $T_1$, with 5 points, $t_0, t_1, \ldots, t_4$. Some transformation k(t) is applied only to point $t_1$ to form time series $T_2$ ($T_2$ thus differs from $T_1$ by a single point), to both $t_1$ and $t_2$ to form time series $T_3$, and to both $t_1$ and $t_3$ to form time series $T_4$ (thus $T_3$ and $T_4$ each differ from $T_1$ by the same value at two points, but in $T_3$ those points are consecutive while in $T_4$ they are not). For distance measures which are sensitive to *both* frequency and duration, $d(T_4, T_1) > d(T_3, T_1) > d(T_2, T_1)$, but for distance measures which are sensitive to frequency but *not* duration, $d(T_4, T_1) > d(T_2, T_1)$, while $d(T_3, T_1) = d(T_2, T_1)$. This is because a distance that is invariant to duration will treat a difference that occurs over multiple consecutive time points as a single difference. This could be especially useful, for example, to cluster a set of time series according to the number of anomalies detected without respect to the lengths of those anomalies.

#### 4. Metric testing

The test for uniqueness was conducted by comparing a time series first to itself, and then to a similar time series with a value difference at a single point. For distance measures with threshold settings (e.g., EDR), we set the threshold to zero to ensure they would recognize the difference. Any distance measure that returned a value of zero when comparing the time series against itself, and any non-zero value when comparing it against a time series with a value difference at a single point, was considered to demonstrate uniqueness.

Symmetry was tested by comparing a pair of different time series, X and Y, in both forward order, d(X, Y), and reverse order, d(Y, X). If the two values returned were identical, the distance measure was considered to demonstrate symmetry. We ensured that the time series were different enough that no distance measure returned 0 for both forward and reverse order.

The triangle inequality and nonnegativity properties were tested by comparing thousands of short, randomized time series generated by a stochastic exponential model. We found that shorter time series were better at detecting violations, so we set the length to five. We generated 300,000 time series and divided them into 100,000 sets of three. Within each set of three, we considered each time series to represent one corner of a triangle and compared them pairwise, with the resulting distances representing the sides of the triangle. We then subtracted the two shorter sides from the longest side. If the difference was greater than zero for any of the 100,000 sets, then the distance measure was considered to violate the triangle inequality. Additionally, if any of the 300,000 time series comparisons produced a negative value, the distance measure was considered to violate nonnegativity. We set the time series generator such that zeros and negative values were included in some time series, as some distance measures satisfy the triangle inequality and/or non-negativity only when all input values are positive or non-negative.

Distance measures were classified as “Full” for full metric if they passed all metric tests, “Semi” for semi-metric if they passed all tests except the triangle inequality, or “Non” for non-metric if they failed one or more of the other tests.

Settings for adaptive distance measures (distance measures with settings that can be changed to alter their behaviour) were set at defaults given in examples from the documentation of the TSdist R package (Mori *et al.*, 2016). For triangle inequality and nonnegativity tests, we kept the same settings for initial testing. If they passed the tests at those settings, we tested them over a range of settings. If they failed at default settings, there was no need for further testing.

#### 5. Controlled testing

We used the Manhattan distance as a basis for devising controlled sensitivity tests for translation, amplitude, duration, frequency, white noise, biased noise, and outliers. The Manhattan distance is the summed absolute difference between each pair of points in a time series. It is a simple-to-calculate metric and demonstrates all the sensitivities we tested for. Furthermore, it responds to sensitivity tests in a linear manner. These properties make the Manhattan distance an ideal basis for comparison of other distance measures.

For each sensitivity test, we constructed a series of n time series with linearly increasing differences, $T_1$, $T_2$, ..., $T_n$, such that the differences in absolute value between point pairs of any consecutive pair of time series summed to 1. Thus, the Manhattan distance between any pair of consecutive time series, $T_i$ and $T_{i+1}$, was 1, and between any non-consecutive pair of time series, $T_i$ and $T_{i+j}$, is j. For example, the Manhattan distance between $T_1$ and $T_2$ would be 1, between $T_2$ and $T_3$ would be 1, and between $T_1$ and $T_5$ would be 5.

Sensitivity tests were conducted for each distance measure by comparing each time series $T_i$ in the set $T_1$, $T_2$, ..., $T_{n,}$ to $T_1$. Any distance measure returning a dissimilarity value of 0 for *every* pair of time series for a given sensitivity test would be considered as invariant for that property, while a distance measure returning the same *non-zero* value for *every* time series pair would be considered as insensitive (note that invariance implies insensitivity, but insensitivity is not the same as invariance. Distance measures that demonstrate insensitivity to a property register differences as binary—different or not different—while those demonstrating invariance do not register differences at all).

Sensitivity is calculated as the mean of all distances between consecutive time series,

$$s = \frac{\sum_{i=1}^{n-1} d(T_i, T_{i+1})}{n} \quad , \tag{1}$$

where s is sensitivity, $d(T_i,T_{i+1})$ is the distance between a pair of consecutive time series $T_i$ and $T_{i+1}$, and n is the total number of time series being compared.

Given that s is an absolute sensitivity value, its interpretation is dependent on the scale of the distance measure. A scale-independent relative sensitivity is obtained by

$$rs_x = \frac{s_x}{s_\mu} \ , \tag{2}$$

where $rs_x$ is the relative sensitivity to property x, $s_x$ is the absolute sensitivity to that property, and $s_\mu$ is the mean of absolute sensitivities to all tested properties.

The sensitivity values for all distance measures are separated into 5 bins and designated as “Very Low,” “Low,” “Medium,” “High,” or “Very High.” The sensitivity value for the Manhattan distance is 1 for every property and serves as the median value for the bins, which are: less than 0.2, 0.2 to 0.75, 0.75 to 1.25, 1.25 to 2.5, and greater than 2.5, respectively. Note, however, that the equation for sensitivity is derived from the linear slope equation, but the sensitivity for many distance measures is non-linear. The calculated sensitivity is a linear approximation along the tested range.

Phase invariance testing was conducted in a similar way to sensitivity testing, with $T_1, T_2, \ldots, T_n$ representing a set of time series, with the difference in phase increasing with i in $T_i$. However, the Manhattan distance could not be used as a basis for comparison. This is because lock-step distance measures (those that match every time point 1-to-1), including the Manhattan distance, do not respond to time translation in a way that can be interpreted by a function. Distance measures were designated as “Inv” (meaning they demonstrated phase invariance) when the dissimilarity between *every* pair of time series was 0, “Ins” (insensitive) when *every* pair of time series returned the same *non-zero* dissimilarity value, “Sens” (sensitive) when the dissimilarity value was dependent on i, or “Unp” (unpredictable) when dissimilarity values differed but did not depend on i. For those distances with window size settings (e.g., some distance measures that act stepwise along time series have a setting to control how many time points are considered in each step), we set the window large enough to cover the maximum difference in phase (that between $T_1$ and $T_n$).

Uniform time scaling invariance was tested using a set of time series in which $T_{i+1}$ was stretched compared to $T_i$. This involved lengthening the time series $T_i$, keeping the first and last time points the same while altering the values at each time point in between to fit the shape change. Warping was tested by stretching only one horizontal section of a time series, such that a set was formed, with $T_{i+1}$ longer than

$T_i$. As with phase invariance, results for uniform time scaling invariance and warping invariance were not compared against the Manhattan distance. The Manhattan distance and other lock-step distance measures are unable to handle time series with different lengths and therefore by default are not invariant to uniform time scaling or warping. Thus, elastic distance measures were tested and designated as either "Inv" if *all* returned dissimilarities were zero, "Ins" if *all* returned dissimilarities were *identical* and *non-zero*, "Sens" if the returned value depended on the degree of time scaling or warping, or "Unp" if returned values differed but did not depend on the degree of time scaling or warping. All lock-step distance measures were designated as "n/a".

Antiparallelism bias was tested by comparing pairs of time series that differed by the same relative amount in different directions. Distance measures were designated as having "Positive" bias if they gave a greater dissimilarity value to pairs of time series differing in opposite directions than to pairs differing in the same direction, "Negative" bias if they gave a greater dissimilarity value to those differing in the same direction, or "Neutral" if they assigned each pair of time series the same dissimilarity value.

#### 6. Uncontrolled testing

We created a function for each property to be tested, which applies a transformation to one or more time points of a real-world time series given as input. Each function accepts a value q, the purpose of which varies depending on the function (see supplementary materials for details). For example, the translation function adds a real number q to every value $t_i$ of a time series T. The transformed time series is returned as output and compared against its unaltered counterpart. We applied the functions to a range of q in increments, then graphed the results as response curves (see Fig. S5, S6, S7, and S8 in supplementary materials). We did not compare them against a reference or assign sensitivity ratings, as they were intended only as a confirmatory check against the results of controlled testing.

Functions:

Translation sensitivity: Add q to every data point of a time series T.

White noise sensitivity: Create a normal distribution with mean q and standard deviation 0.3 times q (the latter is arbitrary). Randomly select half of the data points of a time series T and add randomly selected values from the normal distribution to the selected points. Finally, subtract randomly selected values from the normal distribution from the points that were not selected. This function scales q by $\frac{q}{\max(q)}$ to avoid the noise being too large.

Biased noise sensitivity: Proceed exactly as with white noise sensitivity but skip the final step (the points that were not selected remain untransformed). This function scales q by $\frac{q}{0.5 * \max(q)}$ (the 0.5 is because the function is only applied to half of the time points).

Outlier sensitivity: Add q to one randomly selected point of a time series T (excluding the first and final points, which can cause unintended behaviour in some distance measures).

Phase invariance: Shift the first q time points of a time series T to the end of the time series.

Warping invariance: Randomly select a single value from a time series T and extend the time series by repeating the chosen value q times.

Uniform time scaling invariance: Stretch a time series T along the x-axis by a factor of q. The y-axis values of the first and final points remain unchanged, but the final point is shifted along the x-axis and all points in between are recomputed. For this function, q is scaled: $\frac{q}{\max(q)} + 1$. Thus, time series T will be stretched to a maximum of twice its original length.

#### 7. Metric test results

In some cases, results depended on input values or settings. Eight of the lock-step shape-based distance measures passed the triangle inequality test and/or non-negativity test when inputs were constrained to non-negative real numbers, but failed when negative numbers were included. EDR behaved as a metric when the threshold setting, ε, was set near zero, but failed the triangle inequality test when ε was set at five. The Normalized Compression Distance (NCD) and the Compression-based Dissimilarity Measure (CDM) both failed our uniqueness and symmetry tests and thus qualified as non-metrics, although NCD is stated by its authors to be a metric (Cilibrasi and Vitányi, 2005). However, this is qualified with respect to the compression algorithm paired with it, with none quite reaching the definition the metric behaviour depends on. NCD should approach closer to true metric behaviour the longer the time series (Cilibrasi and Vitányi, 2005). We tested it here with very short time series, and therefore it would not be expected to behave as a metric. Additional testing (not included) showed NCD came closer to passing the uniqueness and symmetry tests, although as the time series reached a length of one million, it was still failing. Beyond that length, running the tests was too slow to be practical. CDM, on the other hand, is not considered to be a metric (Keogh *et al.*, 2004), nor did it approach closer to metric behaviour when tested with longer time series.

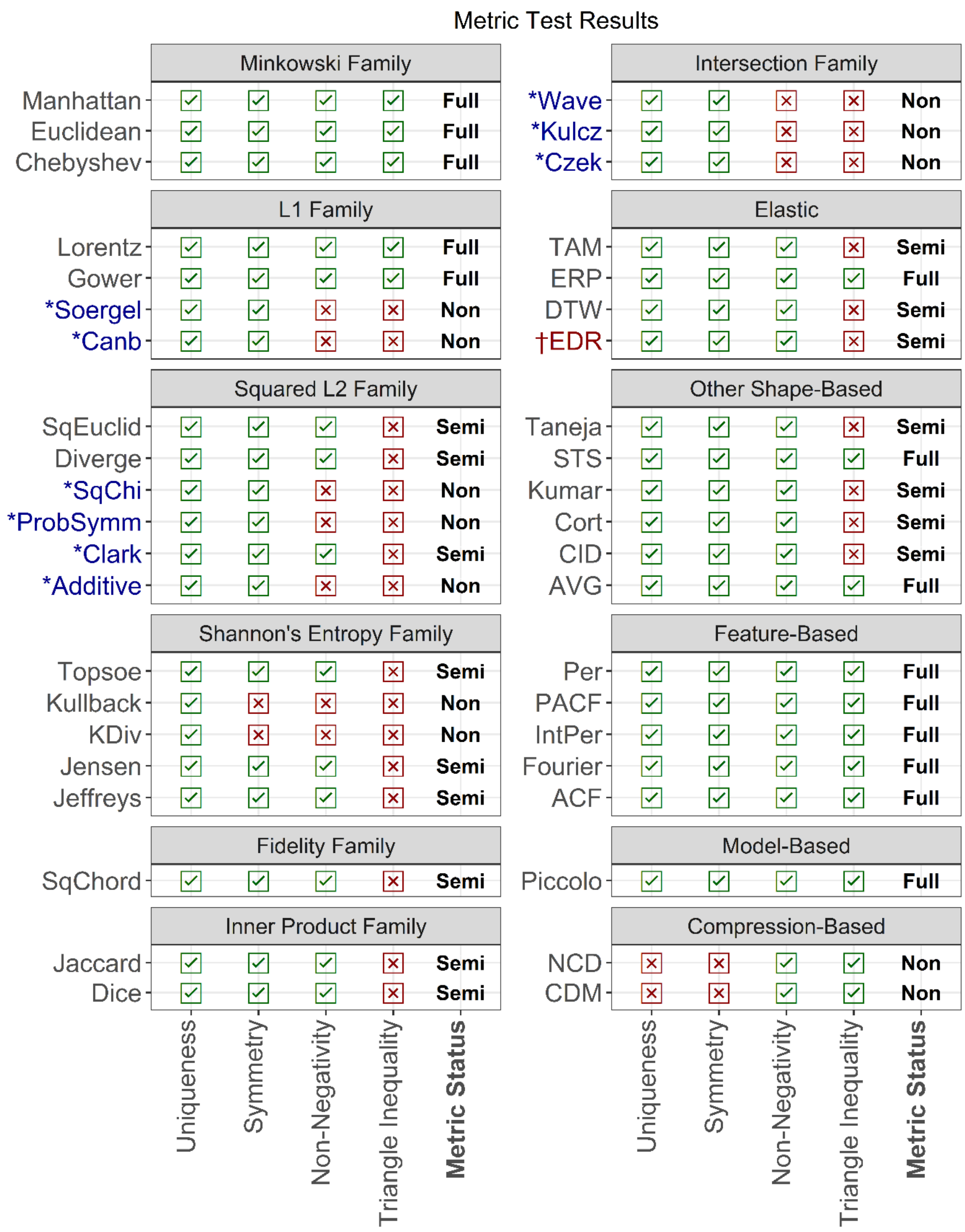


*These distances respond differently when inputs are constrained to non-negative real numbers. As we included negative values in our tests, our results for these measures may differ from others (e.g. Kocher and Savoy, 2017).

†This distance is a full metric when the threshold value (epsilon) is set at 0.

Figure S1. Metric test results for 42 distance measures. Results are arranged by family (for lock-step shape-based measures) or type.

### 8. Controlled test results

#### 8.1. Sensitivity tests

Results for EDR depended on the value of the threshold setting, ε. We reported the results with ε at 0.1. However, when ε was set high, EDR was invariant to all seven of these properties. When ε was set within the range of the input values, results were less predictable.

The two compression-based distances we tested, the Normalized Compression Distance (NCD) and the Compression-based Dissimilarity Measure (CDM), showed insensitivity to translation and outliers. However, our uncontrolled test results did not confirm this. It is not clear why this difference occurred, but keep in mind that compression-based distances may behave differently for short time series than for long ones (e.g., they do not behave as metrics when comparing short time series).

#### 8.2. Time-based invariances and other tests

Again, results for EDR depended on the value of ε. We reported the results with ε at 0.1., but when ε was set high, EDR was invariant to phase, and sensitive to both warping and time scaling. When ε was set within the range of the input values, it responded unpredictably to phase shift, but remained sensitive to both warping and time scaling.

The results for the Autocorrelation-based dissimilarity (ACF) and the Partial Autocorrelation-based dissimilarity (PACF) were "n/a" for both warping and time scaling, suggesting that these distance measures are unable to deal with unequal-length time series. However, this is not the case. The problem is that these measures require an equal number of autocorrelation coefficients, which the short time series we used for controlled testing did not satisfy. However, ACF and PACF did provide results for warping and time scaling in uncontrolled testing (see Fig. S5 and S7).

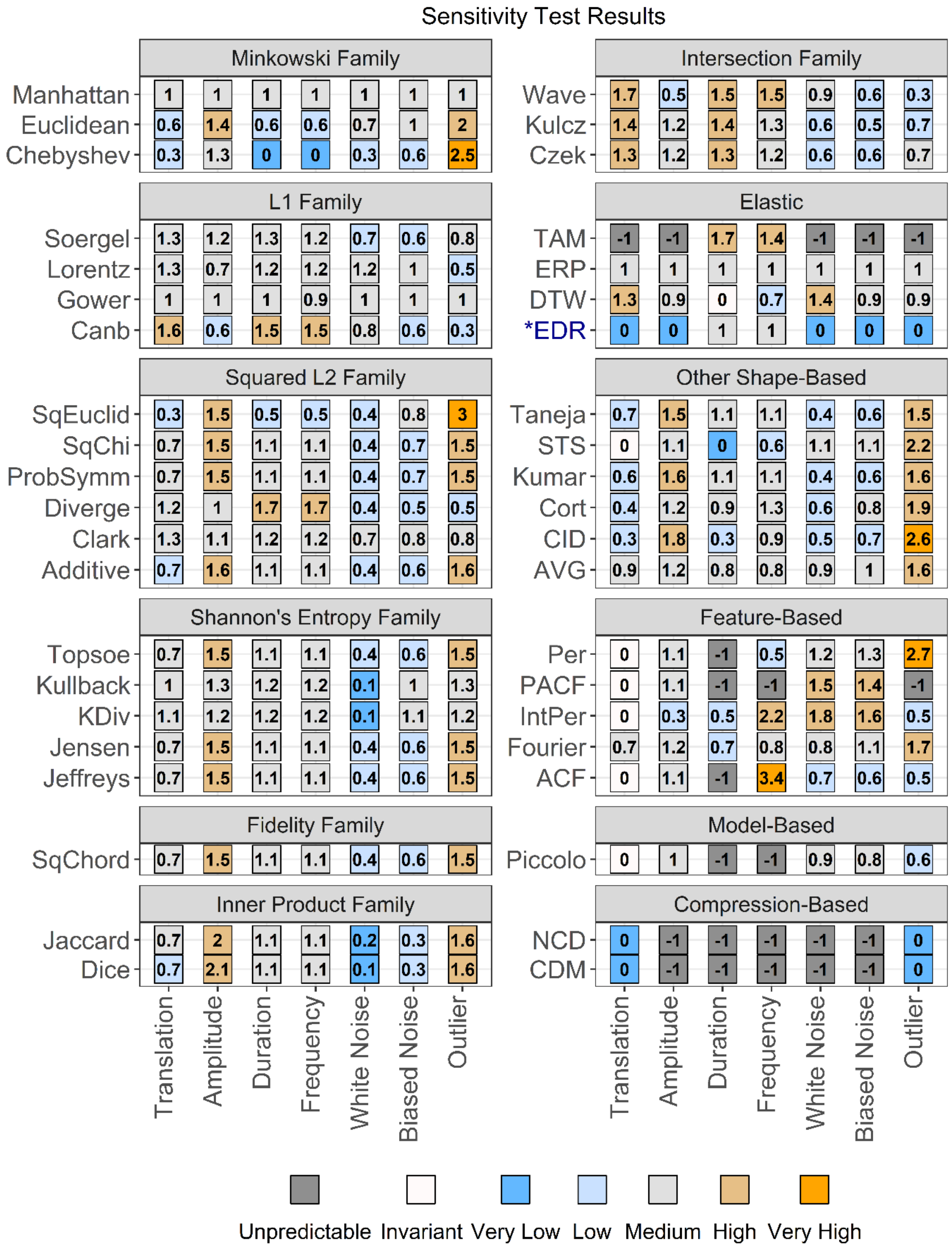


Sensitivity Ranges: Very Low: < 0.2, Low: 0.2 - 0.7, Medium: 0.7 - 1.3, High: 1.3 - 2.5, Very High: > 2.5.

*The results for EDR strongly depended on the threshold setting, epsilon. Here, it was set to 0.1.

Figure S2. Sensitivity test results for 42 distance measures. Results are arranged by family (shape-based measures) or type, and colour-coded according to sensitivity value.

#### Time-Based Invariances & Other Test Results

| Family | Measure | Antiparallelism | Non-Positive Value Handling | Phase Inv. | Uniform Time Scaling Inv. | Warping Inv. |
|---|---|---|---|---|---|---|
| Minkowski Family | Manhattan | ⦶ | All | Unp | n/a | n/a |
| | Euclidean | ⦶ | All | Unp | n/a | n/a |
| | Chebyshev | ⦶ | All | Unp | n/a | n/a |
| L1 Family | Soergel | ⊖ | All | Unp | n/a | n/a |
| | Lorentz | ⦶ | All | Unp | n/a | n/a |
| | Gower | ⦶ | All | Unp | n/a | n/a |
| | Canb | ⊖ | All | Unp | n/a | n/a |
| Squared L2 Family | SqEuclid | ⦶ | All | Unp | n/a | n/a |
| | SqChi | ⊖ | All | Unp | n/a | n/a |
| | ProbSymm | ⊖ | All | Unp | n/a | n/a |
| | Diverge | ⊖ | All | Unp | n/a | n/a |
| | Clark | ⊖ | All | Unp | n/a | n/a |
| | Additive | ⊖ | All | Unp | n/a | n/a |
| Shannon's Entropy Family | Topsoe | ⊖ | None | Unp | n/a | n/a |
| | Kullback | ⊕ | Zeros | Unp | n/a | n/a |
| | KDiv | ⊕ | None | Unp | n/a | n/a |
| | Jensen | ⊖ | Zeros | Unp | n/a | n/a |
| | Jeffreys | ⊖ | Zeros | Unp | n/a | n/a |
| Fidelity Family | SqChord | ⊖ | Zeros | Unp | n/a | n/a |
| Inner Product Family | Jaccard | ⊖ | All | Unp | n/a | n/a |
| | Dice | ⊖ | All | Unp | n/a | n/a |
| Intersection Family | Wave | ⊖ | All | Unp | n/a | n/a |
| | Kulcz | ⊖ | All | Unp | n/a | n/a |
| | Czek | ⊖ | All | Unp | n/a | n/a |
| Elastic | TAM | ⊖ | All | Sens | Unp | Sens |
| | ERP | ⦶ | All | Unp | Sens | Sens |
| | DTW | ⊕ | All | Unp | Unp | Inv |
| | *EDR | ⦶ | All | Unp | Unp | Sens |
| Other Shape-Based | Taneja | ⊖ | Zeros | Unp | n/a | n/a |
| | STS | ⦶ | All | Unp | n/a | n/a |
| | Kumar | ⊖ | Zeros | Unp | n/a | n/a |
| | Cort | ⊖ | All | Unp | n/a | n/a |
| | CID | ⊕ | All | Unp | n/a | n/a |
| | AVG | ⦶ | All | Unp | n/a | n/a |
| Feature-Based | Per | ⊕ | All | Unp | n/a | n/a |
| | PACF | ⊕ | All | Unp | n/a | n/a |
| | IntPer | ⊖ | All | Unp | n/a | n/a |
| | Fourier | ⦶ | All | Unp | n/a | n/a |
| | ACF | ⊖ | All | Unp | n/a | n/a |
| Model-Based | Piccolo | ⊕ | All | Unp | Unp | Unp |
| Compression-Based | NCD | ⦶ | All | Unp | Unp | Unp |
| | CDM | ⦶ | All | Unp | Unp | Unp |

Sens = Sensitive, Ins = Insensitive, Inv = Invariant, Unp = Unpredictable

*For this distance measure, results differ depending on the threshold value, epsilon. Here, epsilon was set to 0.1.

Antiparallelism Bias: ⦶ Neutral, ⊕ Positive, ⊖ Negative

Figure S3. Test results for antiparallelism bias, non-positive value handling, and time-related invariances for 42 distance measures. Results of “n/a” for uniform time scaling invariance and warping invariance mean that the distance measure in question is unable to handle unequal length time series and therefore could not be tested for those properties.

#### 9. Uncontrolled test results

Fig. S5, S6, S7 and S8 show the results of uncontrolled testing of distance measure properties using two real-world time series (Fig. S4) from the UCR Time-Series Classification Archive (Dau *et al.,* 2019), an archive of 128 time-series datasets intended for testing of classification algorithms. All dissimilarity values in the test results have been rescaled to a range of [0,1] using Min-Max scaling. This was done to facilitate placing response curves for different types of transformations on the same plot while still allowing the shape of each response curve to be seen regardless of the strength of the response. For controlled testing, time series were carefully constructed to allow comparison of response strength across different properties. That is a far more difficult problem when working with real-world time series, so we opted instead to exchange strength information for better shape resolution.

For those distance measures that are sensitive to a tested property, the dissimilarity value shows a response curve as the size of the transformation value q increases (see Supplementary Materials section 4 for details on the functions used for testing). The sensitivity response curve may be linear or not but should be described by a function. Invariances show as horizontal lines at a dissimilarity value of zero, while insensitivities show as horizontal lines at some non-zero value. The response curves for some properties differ in shape between time series, especially for elastic distance measures and those designed for stationary time series. Despite this, results are largely consistent with the controlled testing results shown in Fig. S2 and S3.

There are a few exceptions, however. Both compression-based distances we tested, NCD and CDM, registered as insensitive to translation and outliers in controlled testing, while showing unpredictability in uncontrolled testing. Two feature-based distances, ACF and PACF, showed unpredictability for warping invariance and uniform time scaling invariance in uncontrolled testing but failed to give results in controlled testing. This was because these distance measures require the time series being compared to have an equal number of autocorrelation coefficients, a requirement which was met when extending the real-world time series, but not when extending the short time series that we created for controlled testing. Finally, the Time Alignment Measurement distance, TAM, showed unpredictability to outliers in controlled testing, but was insensitive in uncontrolled testing. The raw dissimilarity values from the controlled testing showed a sudden increase from a dissimilarity value of 0 to 0.33 as the value of q increased from 2 to 3. Given that we used the same starting value of q (1) and the same increment size (also 1) for both controlled and uncontrolled testing, the threshold is presumably not determined simply by the value of the outlier, q, but by a more complex calculation.

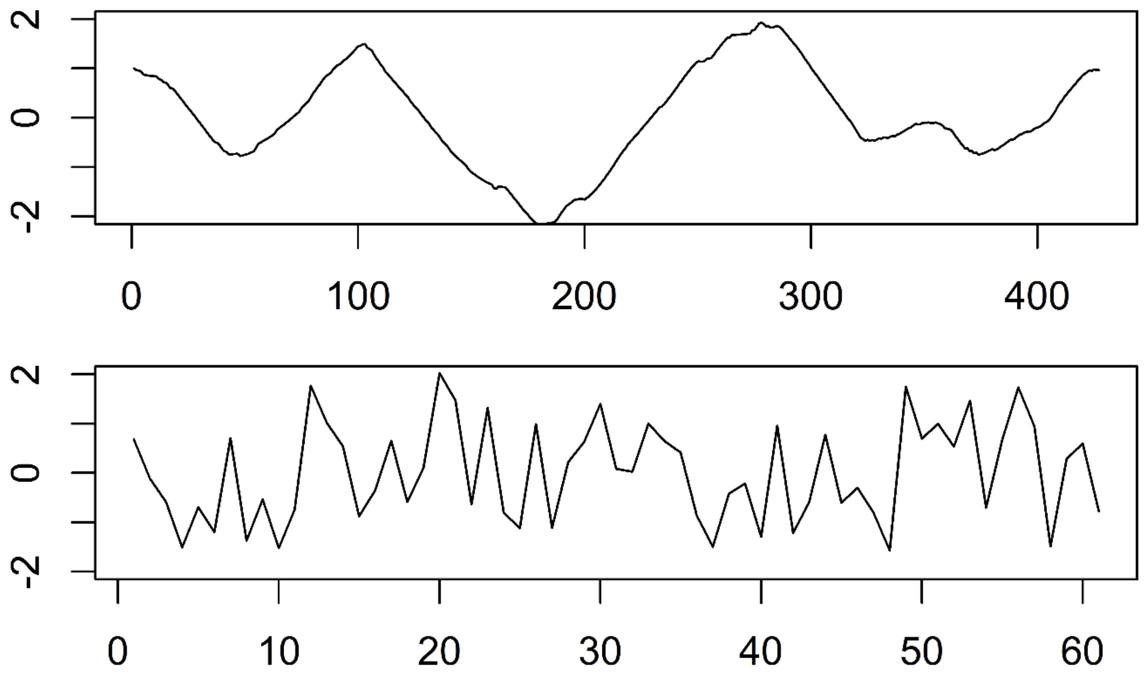


Figure S4. One time series from each of the Yoga (top) and Synthetic Control (bottom) datasets of the UCR Time-Series Archive. Time series in the archive are z-normalized. Therefore, we applied a translation shift before testing to ensure compatibility with distance measures that are unable to handle zeros or negative values.

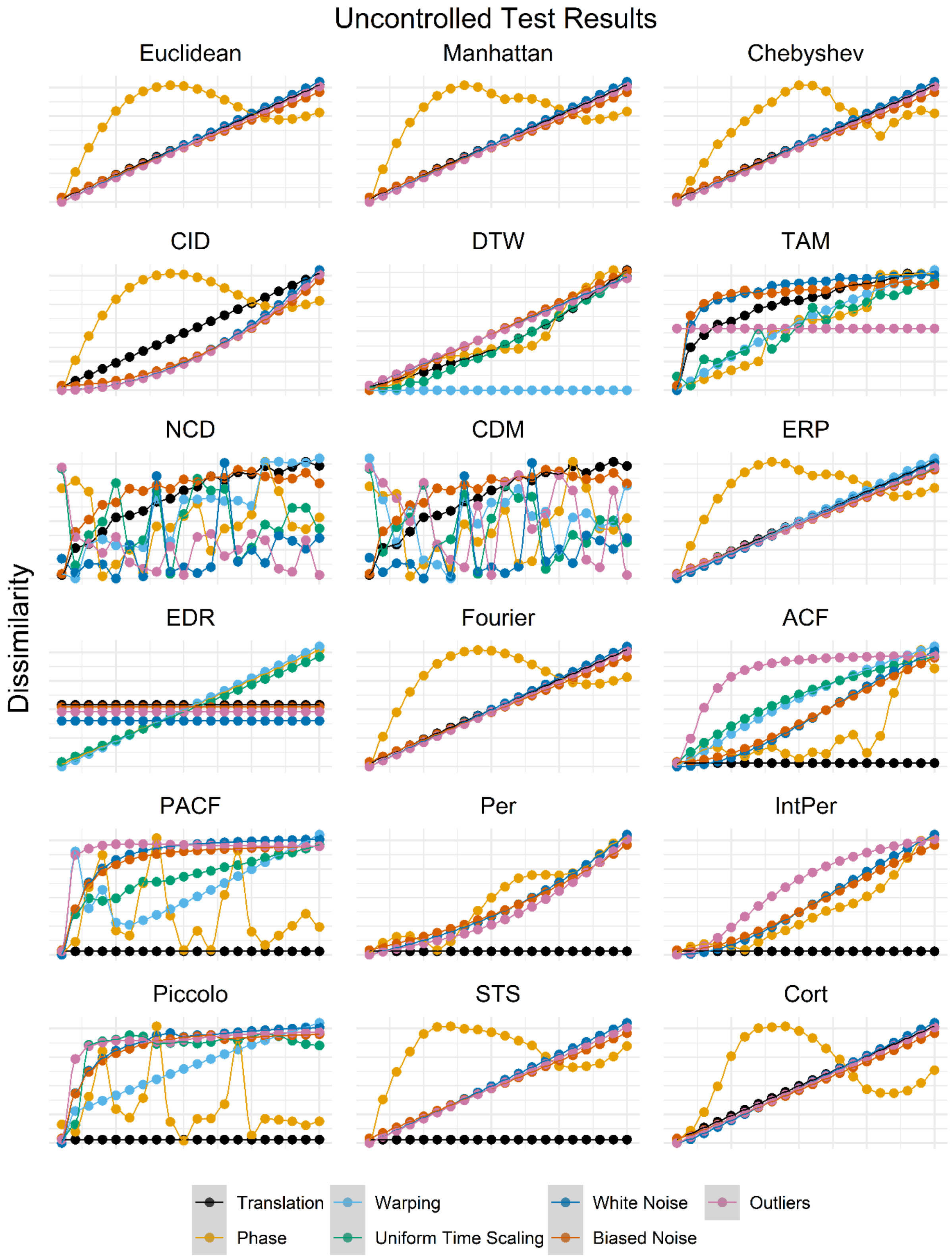


Figure S5. Dissimilarity measurements from 17 distance measures of the TSdist package after applying transformations to a randomly selected time series from the Yoga dataset of the UCR Time-Series Archive. The x-axis depicts the transformation value q across a range of 1 to 200 in increments of 10. Dissimilarity values were rescaled using Min-Max scaling to a range of [0,1] to ensure that the shape of each response curve would be visible regardless of the strength of the response.

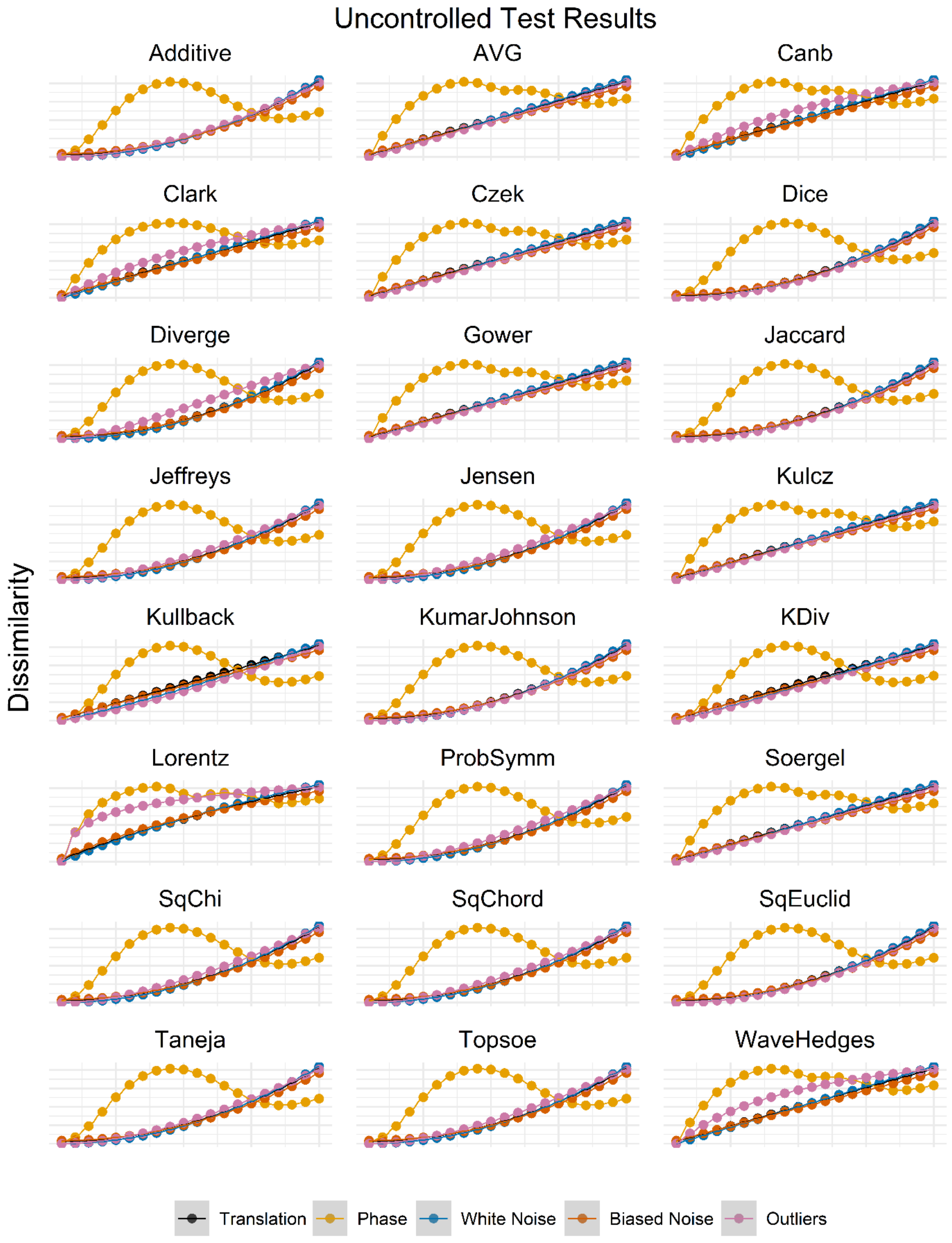


Figure S6. Dissimilarity measurements from 24 distance measures of the philentropy package after applying transformations to a randomly selected time series from the Yoga dataset of the UCR Time-Series Archive. The x-axis depicts the transformation value q across a range of 1 to 200 in increments of 10. Dissimilarity values were rescaled using Min-Max scaling to a range of [0,1] to ensure that the shape of each response curve would be visible regardless of the strength of the response.

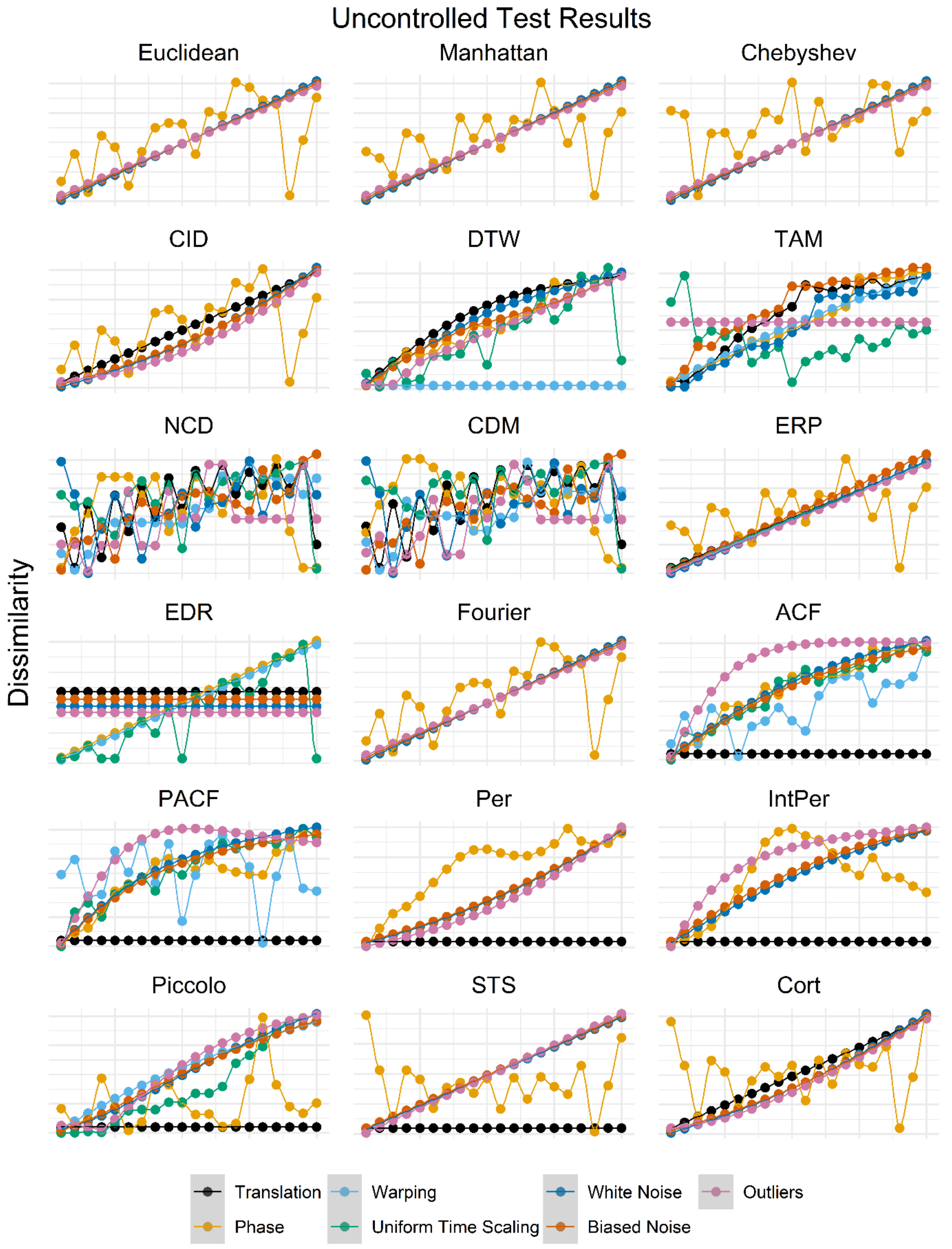


Figure S7. Dissimilarity measurements from 17 distance measures of the TSdist package after applying transformations to a randomly selected time series from the Synthetic Control dataset of the UCR Time-Series Archive. The x-axis depicts the transformation value q across a range of 1 to 20 in increments of 1. Dissimilarity values were rescaled using Min-Max scaling to a range of [0,1] to ensure that the shape of each response curve would be visible regardless of the strength of the response.

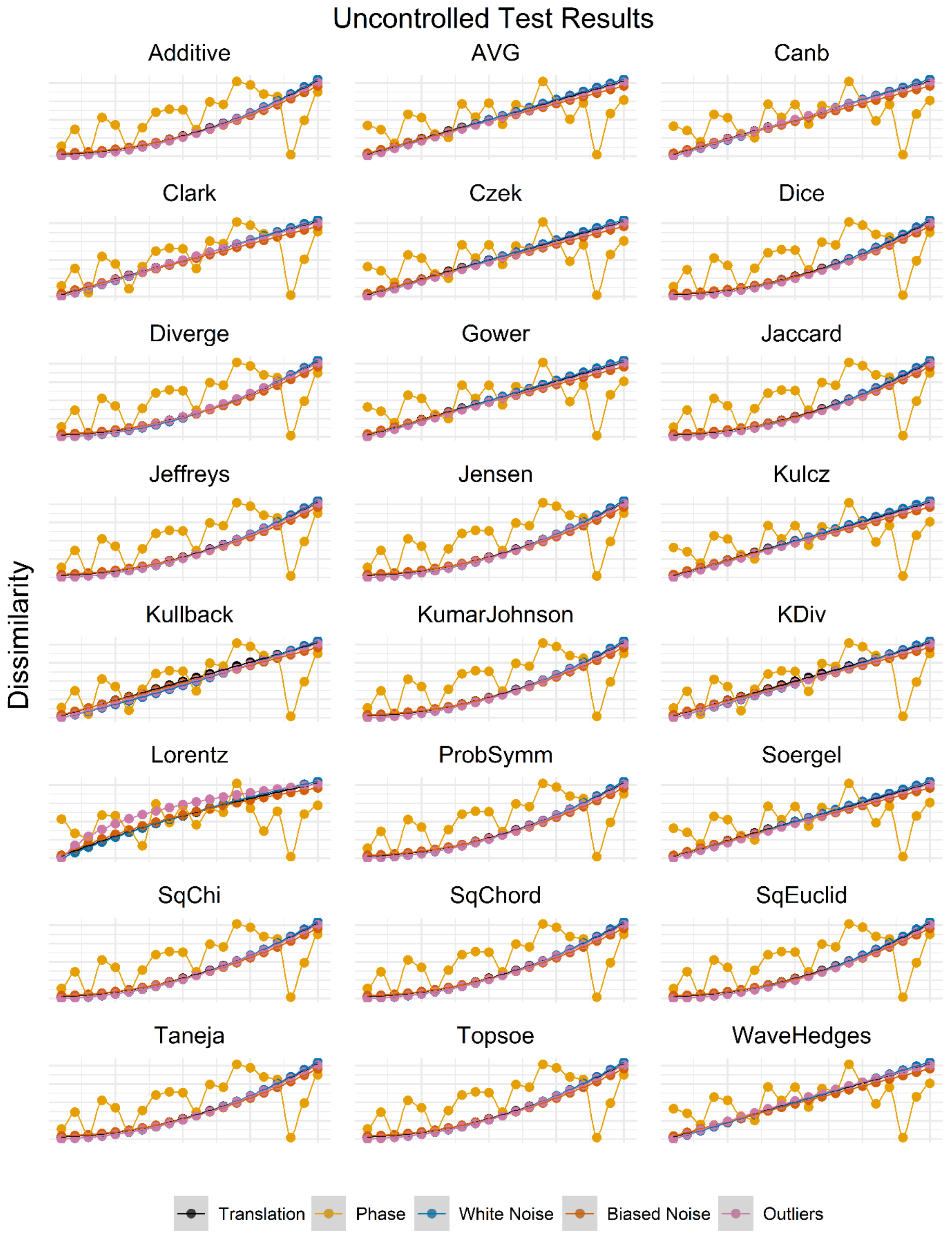


Figure S8. Dissimilarity measurements from 24 distance measures of the philentropy package after applying transformations to a randomly selected time series from the Synthetic Control dataset of the UCR Time-Series Archive. The x-axis depicts the transformation value q across a range of 1 to 20 in increments of 1. Dissimilarity values were rescaled using Min-Max scaling to a range of [0,1] to ensure that the shape of each response curve would be visible regardless of the strength of the response.

#### 10. Example dataset results

When comparing unsmoothed time series, the two selected distance measures gave identical results to percent improvement from Jellesmark *et al.* (2021) (Table S2). Among the 40 unselected distance measures, only one, EDR, gave the same rankings as the selected distance measures and percent improvement (Table S2). Another 30 agreed with Jellesmark *et al.* (2021) in ranking Redshank first, but beyond that the results differed strongly, with 28 ranking Yellow Wagtail second and 23 ranking Curlew last (Table S2). None of the distance measures returned the same results as the t-test.

In the smoothed time series comparison, all seven of the selected distance measures agreed with each other but differed slightly from the percent improvement results of Jellesmark *et al.* (2021) by placing Snipe ahead of Lapwing (Table S3). Of the 35 unselected distance measure, four gave the same rankings as the selected distance measures, while 11 agreed with percent improvement and five agreed with the t-test (Table S3).

Table S2. Comparative rankings of conservation impact on unsmoothed trends of 5 wading bird species according to percent improvement, t-test, and distance measures.

| Method/ Distance Measure | Rank | | | | |
|---|---|---|---|---|---|
| | 1 | 2 | 3 | 4 | 5 |
| % Improvement | Redshank | Lapwing | Snipe | Curlew | Yellow Wagtail |
| t-test | Redshank | Lapwing | Curlew | Snipe | Yellow Wagtail |
| *KDiv | Redshank | Lapwing | Snipe | Curlew | Yellow Wagtail |
| *Kullback | Redshank | Lapwing | Snipe | Curlew | Yellow Wagtail |
| ERP | Redshank | Snipe | Lapwing | Yellow Wagtail | Curlew |
| Euclidean | Redshank | Yellow Wagtail | Snipe | Lapwing | Curlew |
| Manhattan | Redshank | Yellow Wagtail | Snipe | Lapwing | Curlew |
| Gower | Redshank | Yellow Wagtail | Snipe | Lapwing | Curlew |
| Lorentz | Redshank | Lapwing | Yellow Wagtail | Snipe | Curlew |
| Avg | Redshank | Yellow Wagtail | Snipe | Lapwing | Curlew |
| SqEuclid | Redshank | Yellow Wagtail | Snipe | Lapwing | Curlew |
| SDD | Yellow Wagtail | Snipe | Redshank | Curlew | Lapwing |
| ACF | Lapwing | Curlew | Snipe | Redshank | Yellow Wagtail |
| PACF | Yellow Wagtail | Curlew | Snipe | Redshank | Lapwing |
| Additive | Redshank | Yellow Wagtail | Lapwing | Snipe | Curlew |
| Canberra | Redshank | Yellow Wagtail | Lapwing | Curlew | Snipe |
| CDM | Snipe | Yellow Wagtail | Redshank | Curlew | Lapwing |
| Chebyshev | Redshank | Yellow Wagtail | Snipe | Lapwing | Curlew |

| Method/ Distance Measure | Rank 1 | 2 | 3 | 4 | 5 |
|---|---|---|---|---|---|
| CID | Snipe | Yellow Wagtail | Redshank | Lapwing | Curlew |
| Clark | Redshank | Yellow Wagtail | Lapwing | Curlew | Snipe |
| Cort | Redshank | Snipe | Curlew | Yellow Wagtail | Lapwing |
| Czek | Redshank | Yellow Wagtail | Lapwing | Curlew | Snipe |
| Dice | Redshank | Yellow Wagtail | Lapwing | Snipe | Curlew |
| Diverge | Redshank | Yellow Wagtail | Lapwing | Curlew | Snipe |
| DTW | Redshank | Yellow Wagtail | Lapwing | Snipe | Curlew |
| EDR | Redshank | Lapwing | Snipe | Curlew | Yellow Wagtail |
| Fourier | Redshank | Lapwing | Snipe | Yellow Wagtail | Curlew |
| IntPer | Lapwing | Snipe | Yellow Wagtail | Curlew | Redshank |
| Jaccard | Redshank | Yellow Wagtail | Lapwing | Snipe | Curlew |
| Jeffreys | Redshank | Yellow Wagtail | Lapwing | Snipe | Curlew |
| Jensen | Redshank | Yellow Wagtail | Lapwing | Snipe | Curlew |
| Kulcz | Redshank | Yellow Wagtail | Lapwing | Curlew | Snipe |
| KumarJohnson | Redshank | Yellow Wagtail | Lapwing | Snipe | Curlew |
| NCD | Snipe | Yellow Wagtail | Redshank | Curlew | Lapwing |
| Per | Yellow Wagtail | Snipe | Redshank | Lapwing | Curlew |
| Piccolo | Snipe | Curlew | Lapwing | Redshank | Yellow Wagtail |
| ProbSymm | Redshank | Yellow Wagtail | Lapwing | Snipe | Curlew |
| Soergel | Redshank | Yellow Wagtail | Lapwing | Curlew | Snipe |
| SqChi | Redshank | Yellow Wagtail | Lapwing | Snipe | Curlew |
| SqChord | Redshank | Yellow Wagtail | Lapwing | Snipe | Curlew |
| STS | Yellow Wagtail | Snipe | Redshank | Curlew | Lapwing |
| TAM | Redshank | Lapwing | Snipe | Curlew | Yellow Wagtail |
| Taneja | Redshank | Yellow Wagtail | Lapwing | Snipe | Curlew |
| Topsoe | Redshank | Yellow Wagtail | Lapwing | Snipe | Curlew |
| WaveHedges | Redshank | Yellow Wagtail | Lapwing | Curlew | Snipe |
| *Selected distance measures | | | | | |

Table S3. Comparative rankings of conservation impact on trends of 5 wading bird species according to percent improvement, t-test, and distance measures. The distance measures were applied to a LOESS smoothed version of the population trends.

| Method/Distance Measure | Rank 1 | 2 | 3 | 4 | 5 |
|---|---|---|---|---|---|
| % Improvement | Redshank | Lapwing | Snipe | Curlew | Yellow Wagtail |
| t-test | Redshank | Lapwing | Curlew | Snipe | Yellow Wagtail |
| *ERP | Redshank | Snipe | Lapwing | Curlew | Yellow Wagtail |
| *Euclidean | Redshank | Snipe | Lapwing | Curlew | Yellow Wagtail |
| *Manhattan | Redshank | Snipe | Lapwing | Curlew | Yellow Wagtail |
| *Gower | Redshank | Snipe | Lapwing | Curlew | Yellow Wagtail |
| *Lorentz | Redshank | Snipe | Lapwing | Curlew | Yellow Wagtail |
| *Avg | Redshank | Snipe | Lapwing | Curlew | Yellow Wagtail |
| *SqEuclid | Redshank | Snipe | Lapwing | Curlew | Yellow Wagtail |
| ACF | Lapwing | Snipe | Curlew | Redshank | Yellow Wagtail |
| PACF | Snipe | Redshank | Lapwing | Curlew | Yellow Wagtail |

| Method/Distance Measure | Rank | | | | |
|---|---|---|---|---|---|
| | 1 | 2 | 3 | 4 | 5 |
| Additive | Redshank | Lapwing | Snipe | Curlew | Yellow Wagtail |
| Canberra | Redshank | Lapwing | Yellow Wagtail | Curlew | Snipe |
| CDM | Lapwing | Redshank | Snipe | Yellow Wagtail | Curlew |
| Chebyshev | Redshank | Snipe | Lapwing | Curlew | Yellow Wagtail |
| CID | Redshank | Snipe | Curlew | Lapwing | Yellow Wagtail |
| Clark | Redshank | Lapwing | Curlew | Yellow Wagtail | Snipe |
| Cort | Redshank | Lapwing | Snipe | Curlew | Yellow Wagtail |
| Czek | Redshank | Lapwing | Curlew | Snipe | Yellow Wagtail |
| Dice | Redshank | Lapwing | Curlew | Snipe | Yellow Wagtail |
| Diverge | Redshank | Lapwing | Curlew | Yellow Wagtail | Snipe |
| DTW | Redshank | Lapwing | Snipe | Curlew | Yellow Wagtail |
| EDR | Lapwing | Curlew | Snipe | Redshank | Yellow Wagtail |
| Fourier | Redshank | Snipe | Lapwing | Curlew | Yellow Wagtail |
| IntPer | Snipe | Yellow Wagtail | Redshank | Curlew | Lapwing |
| Jaccard | Redshank | Lapwing | Curlew | Snipe | Yellow Wagtail |
| Jeffreys | Redshank | Lapwing | Snipe | Curlew | Yellow Wagtail |
| Jensen | Redshank | Lapwing | Snipe | Curlew | Yellow Wagtail |
| KDiv | Redshank | Snipe | Lapwing | Curlew | Yellow Wagtail |
| Kulcz | Redshank | Lapwing | Curlew | Snipe | Yellow Wagtail |
| Kullback | Redshank | Snipe | Lapwing | Curlew | Yellow Wagtail |
| KumarJohnson | Redshank | Lapwing | Snipe | Curlew | Yellow Wagtail |
| NCD | Lapwing | Redshank | Snipe | Yellow Wagtail | Curlew |
| Per | Snipe | Redshank | Lapwing | Yellow Wagtail | Curlew |
| Piccolo | Snipe | Curlew | Lapwing | Redshank | Yellow Wagtail |
| ProbSymm | Redshank | Lapwing | Snipe | Curlew | Yellow Wagtail |
| Soergel | Redshank | Lapwing | Curlew | Snipe | Yellow Wagtail |
| SqChi | Redshank | Lapwing | Snipe | Curlew | Yellow Wagtail |
| SqChord | Redshank | Lapwing | Snipe | Curlew | Yellow Wagtail |
| STS | Redshank | Snipe | Yellow Wagtail | Lapwing | Curlew |
| TAM | Lapwing | Redshank | Snipe | Curlew | Yellow Wagtail |
| Taneja | Redshank | Lapwing | Snipe | Curlew | Yellow Wagtail |
| Topsoe | Redshank | Lapwing | Snipe | Curlew | Yellow Wagtail |
| WaveHedges | Redshank | Lapwing | Yellow Wagtail | Curlew | Snipe |
| *Selected distance measures | | | | | |

#### 11. Speeding up DTW

For matching problems, such as content queries and classification, the slowness of DTW can be avoided by indexing, which severely reduces the number of time series that need to be compared to find the best match. For the Euclidean Distance, indexing is relatively straightforward to accomplish. However, as DTW does not satisfy the triangle inequality (Fig. 3), it presents more of a challenge. Keogh and Ratanamahatana (2005) solved this problem using a tight 'lower-bounding' measure, which is included in the TSdist package (Mori *et al.,* 2016) as LBKeoghDistance. For an explanation of lower bounding and the indexing process with respect to DTW, refer to Keogh and Ratanamahatana (2005). The lower-bounding technique does not apply to clustering, where some real-world problems can take weeks or even months (Zhu *et al.*, 2012). However, Zhu *et al.* (2012) solved this problem for clustering by creating an interactive 'anytime algorithm', which uses a fast approximation of DTW to give a best available answer that improves over time as exact DTW calculations are performed, and can be paused or terminated at any time.

#### 13. Plots of controlled test results for all distance measures

This section contains plots of controlled testing results for all 42 distance measures we tested. Each figure includes all time-based and values-based properties for which that distance measure gave results. Distance measures are presented in alphabetical order.

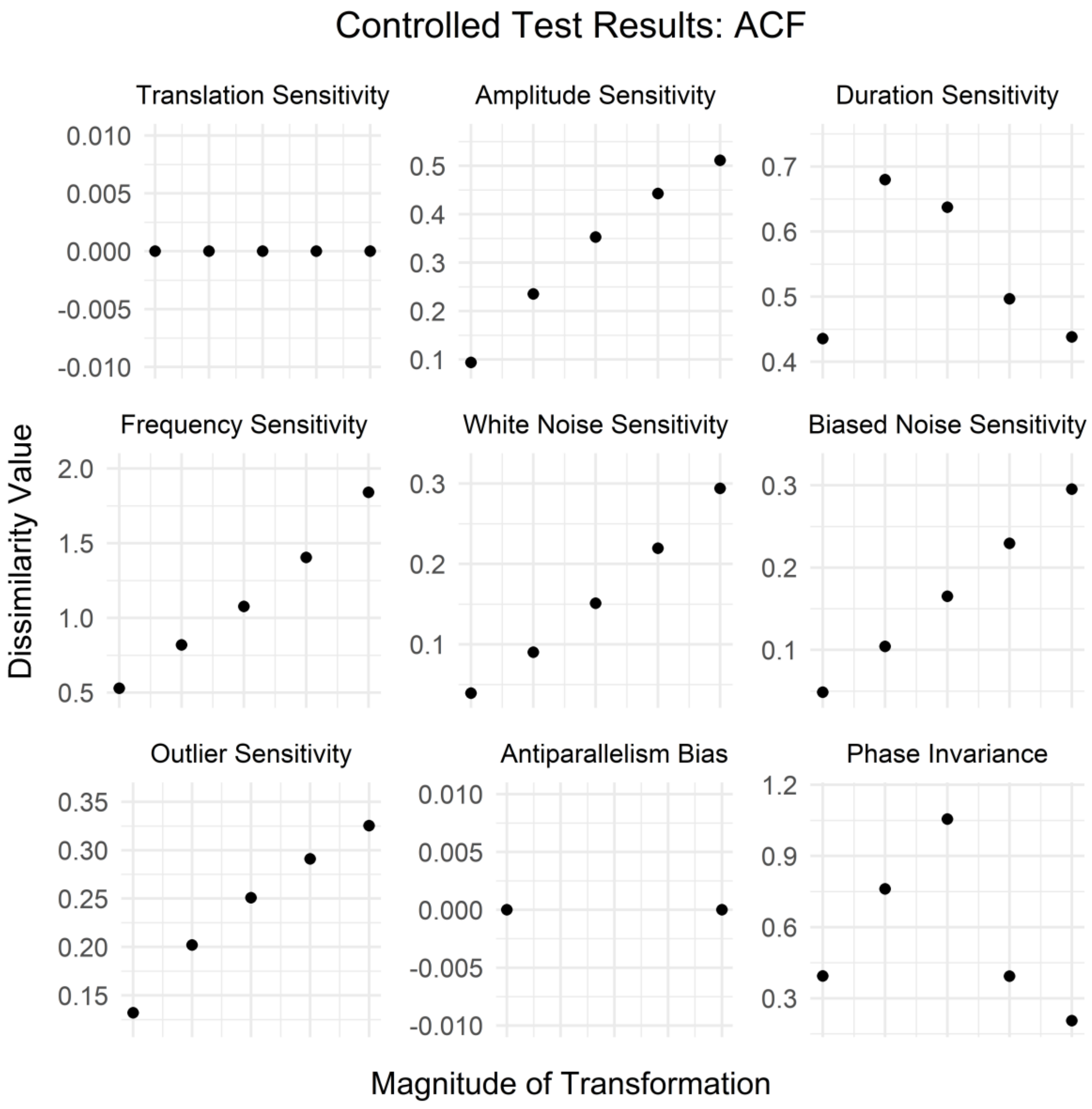

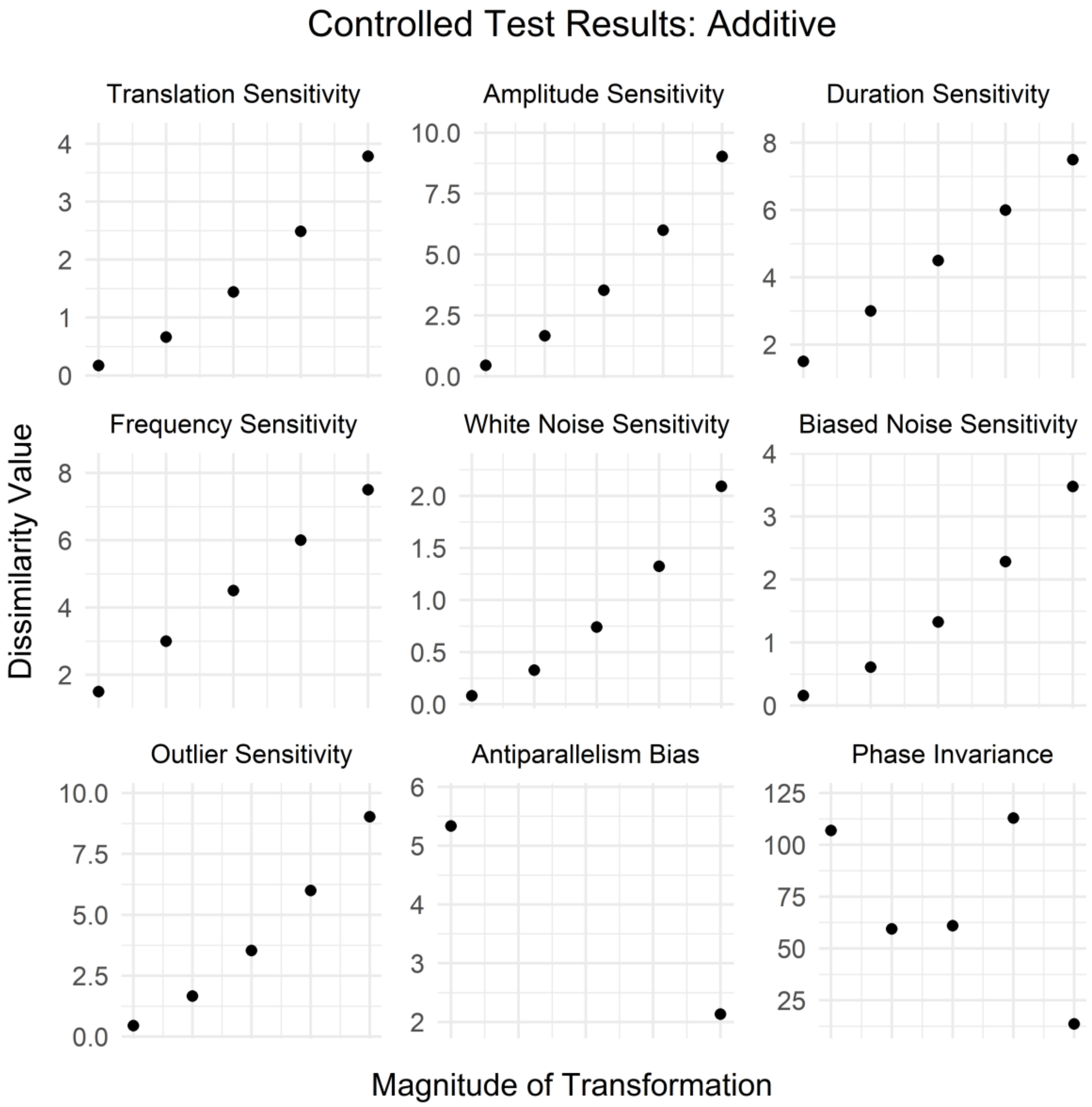
Controlled Test Results: Additive
Translation Sensitivity
Amplitude Sensitivity
Duration Sensitivity
Frequency Sensitivity
White Noise Sensitivity
Biased Noise Sensitivity
Outlier Sensitivity
Antiparallelism Bias
Phase Invariance
Dissimilarity Value
Magnitude of Transformation

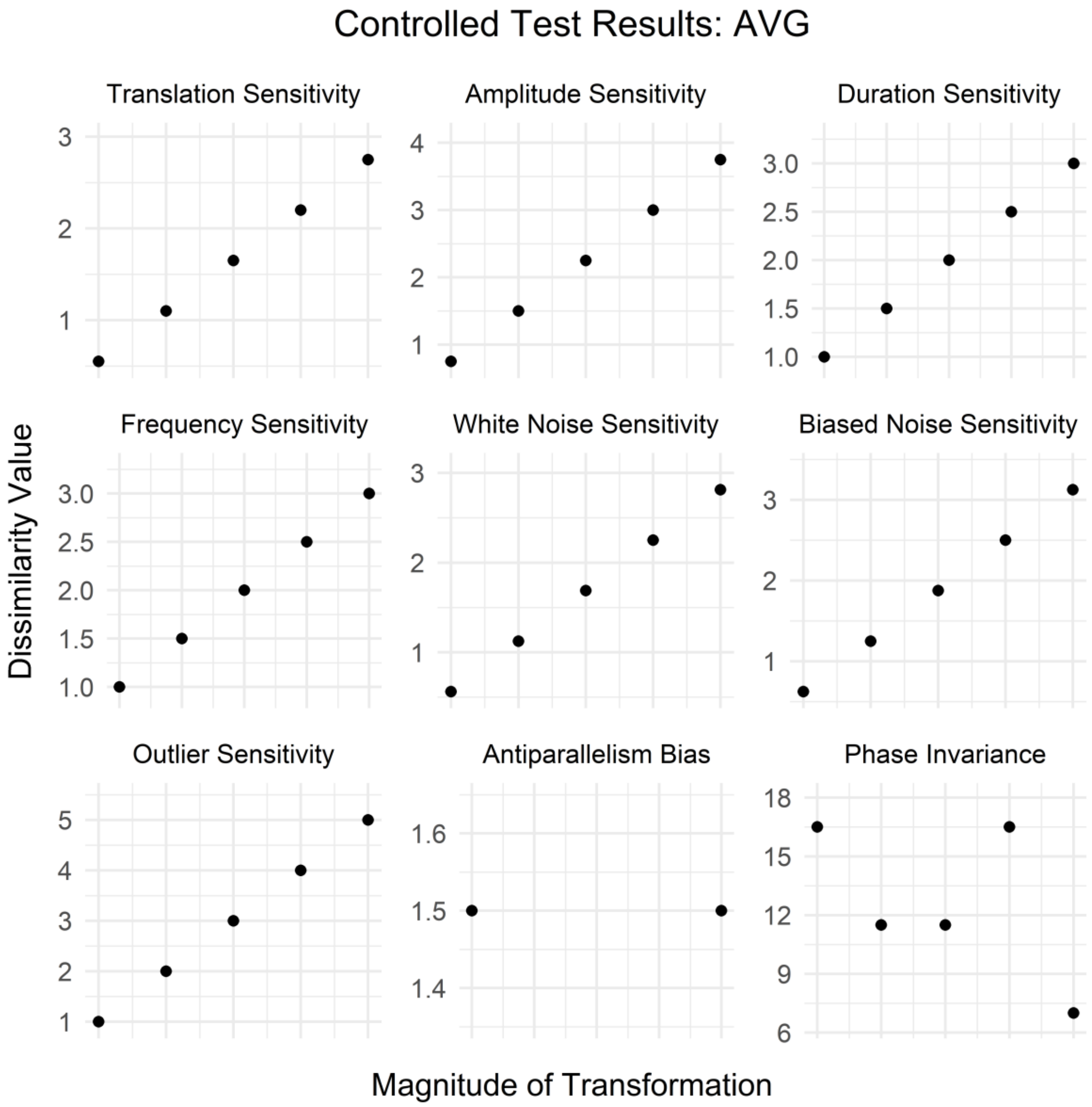
Controlled Test Results: AVG
Translation Sensitivity
Amplitude Sensitivity
Duration Sensitivity
Frequency Sensitivity
White Noise Sensitivity
Biased Noise Sensitivity
Outlier Sensitivity
Antiparallelism Bias
Phase Invariance
Dissimilarity Value
Magnitude of Transformation

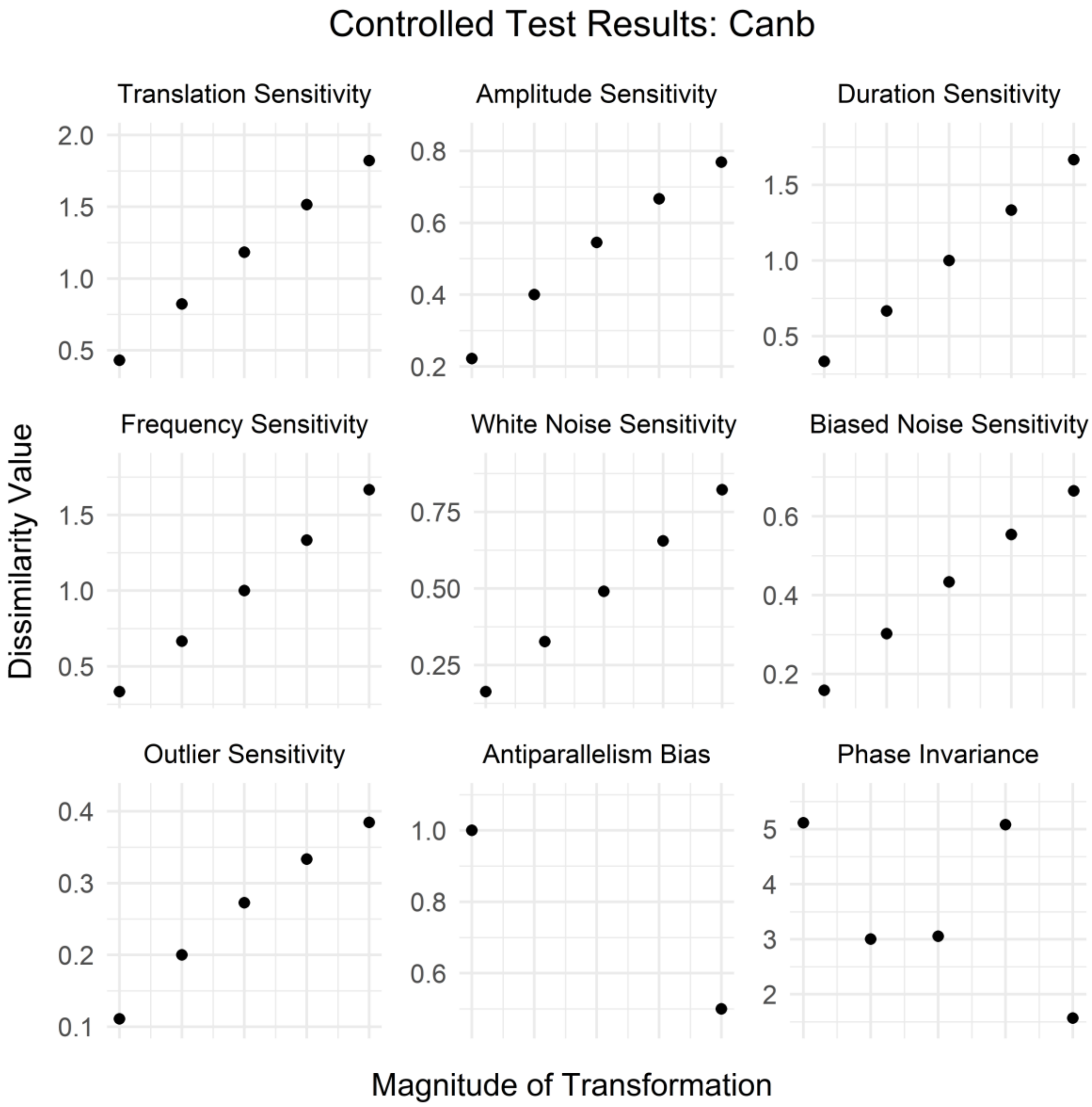
Controlled Test Results: Canb
Translation Sensitivity
Amplitude Sensitivity
Duration Sensitivity
Frequency Sensitivity
White Noise Sensitivity
Biased Noise Sensitivity
Outlier Sensitivity
Antiparallelism Bias
Phase Invariance
Dissimilarity Value
Magnitude of Transformation

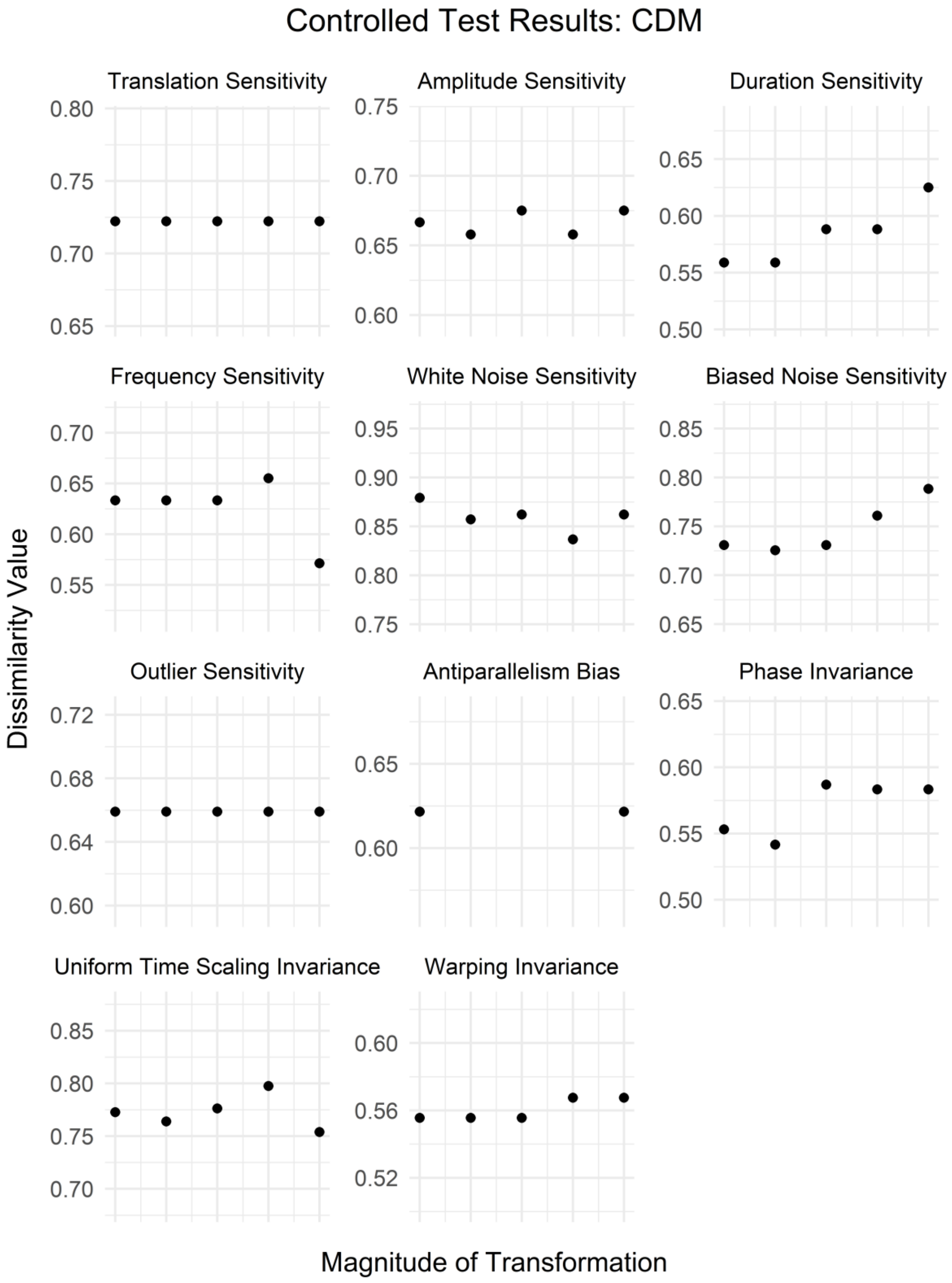
Controlled Test Results: CDM
Translation Sensitivity
Amplitude Sensitivity
Duration Sensitivity
Frequency Sensitivity
White Noise Sensitivity
Biased Noise Sensitivity
Outlier Sensitivity
Antiparallelism Bias
Phase Invariance
Uniform Time Scaling Invariance
Warping Invariance
Dissimilarity Value
Magnitude of Transformation

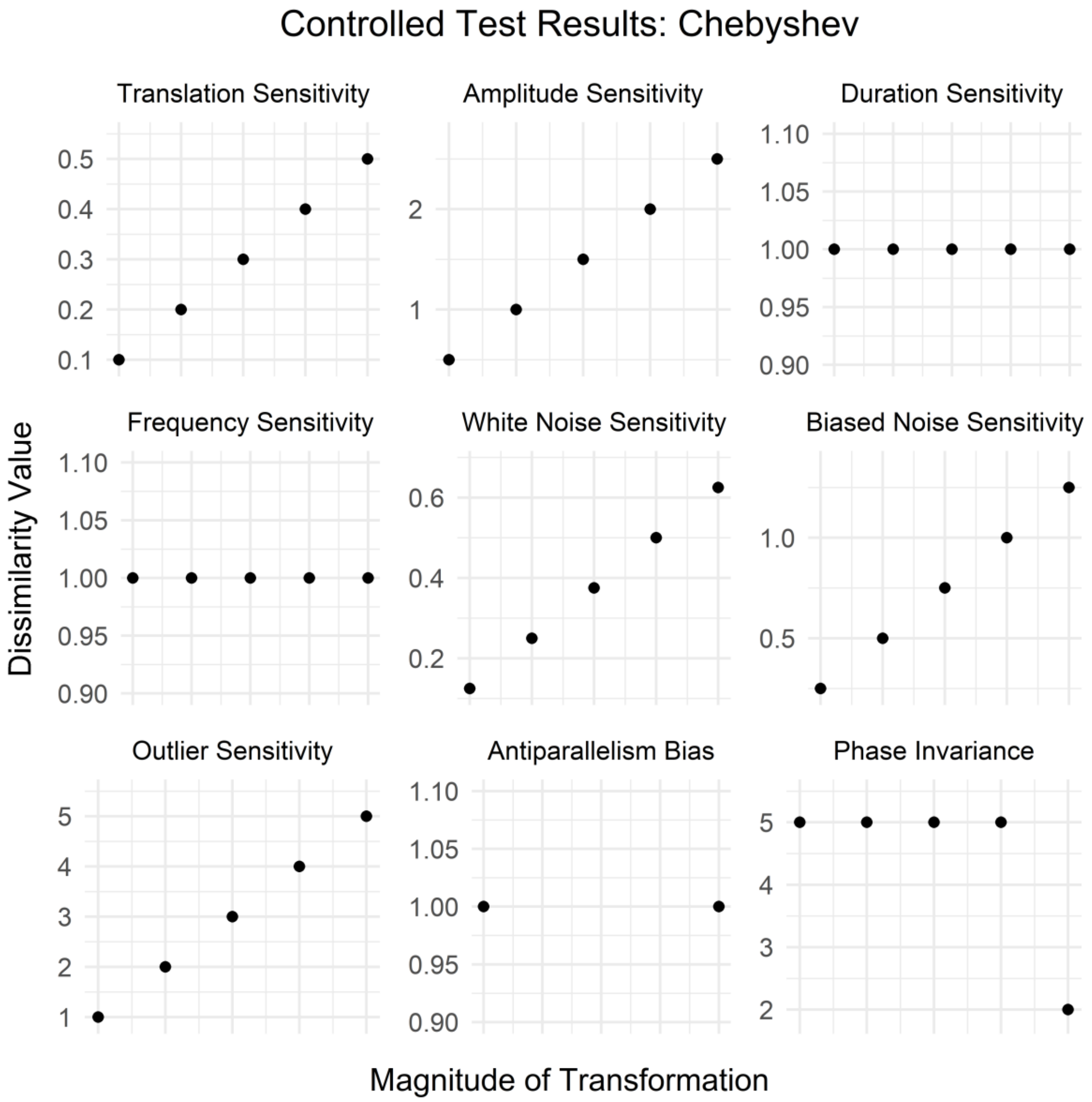
Controlled Test Results: Chebyshev
Translation Sensitivity
Amplitude Sensitivity
Duration Sensitivity
Frequency Sensitivity
White Noise Sensitivity
Biased Noise Sensitivity
Outlier Sensitivity
Antiparallelism Bias
Phase Invariance
Dissimilarity Value
Magnitude of Transformation

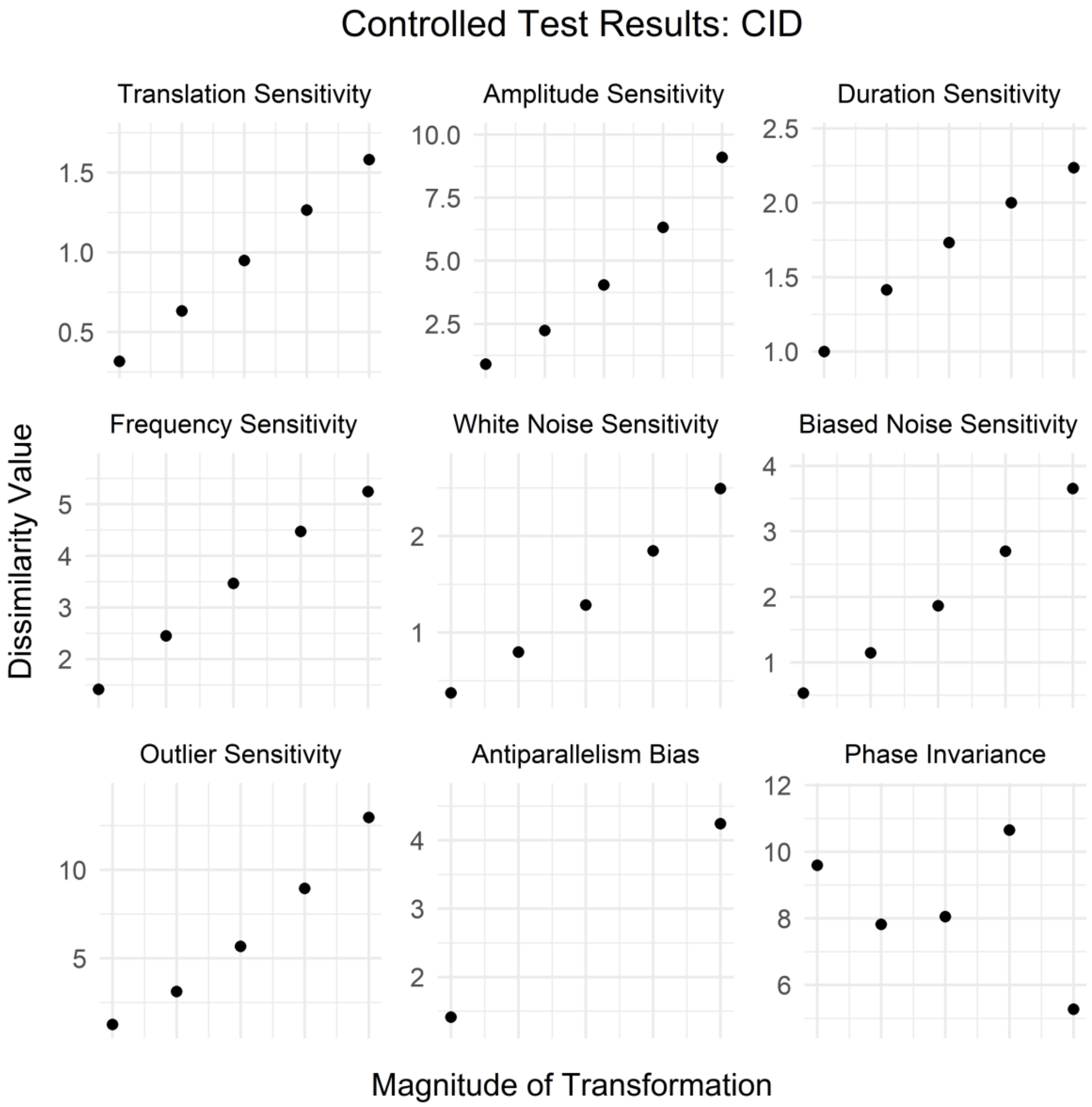
Controlled Test Results: CID
Translation Sensitivity
Amplitude Sensitivity
Duration Sensitivity
Frequency Sensitivity
White Noise Sensitivity
Biased Noise Sensitivity
Outlier Sensitivity
Antiparallelism Bias
Phase Invariance
Dissimilarity Value
Magnitude of Transformation

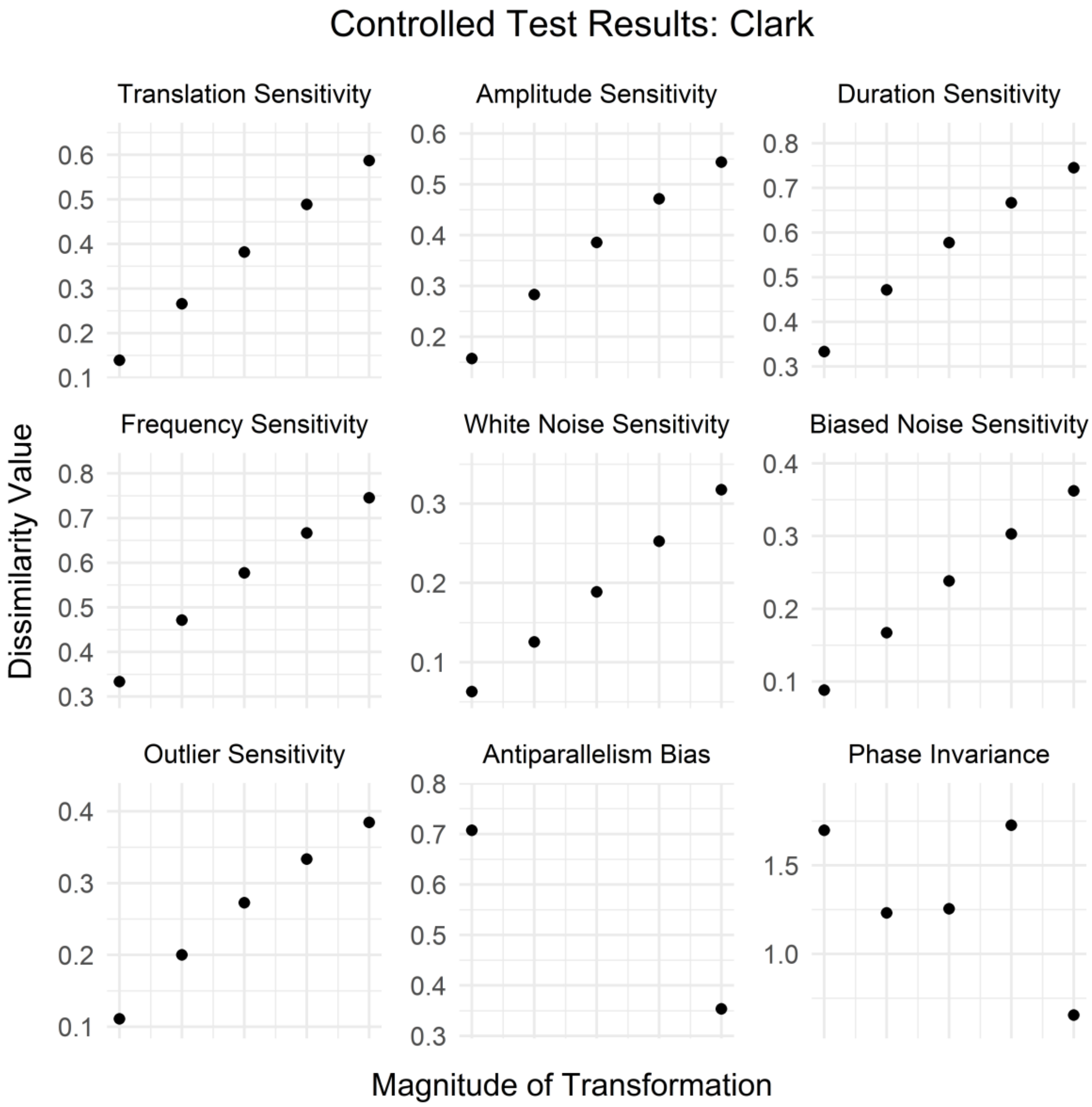
Controlled Test Results: Clark
Translation Sensitivity
Amplitude Sensitivity
Duration Sensitivity
Frequency Sensitivity
White Noise Sensitivity
Biased Noise Sensitivity
Outlier Sensitivity
Antiparallelism Bias
Phase Invariance
Dissimilarity Value
Magnitude of Transformation

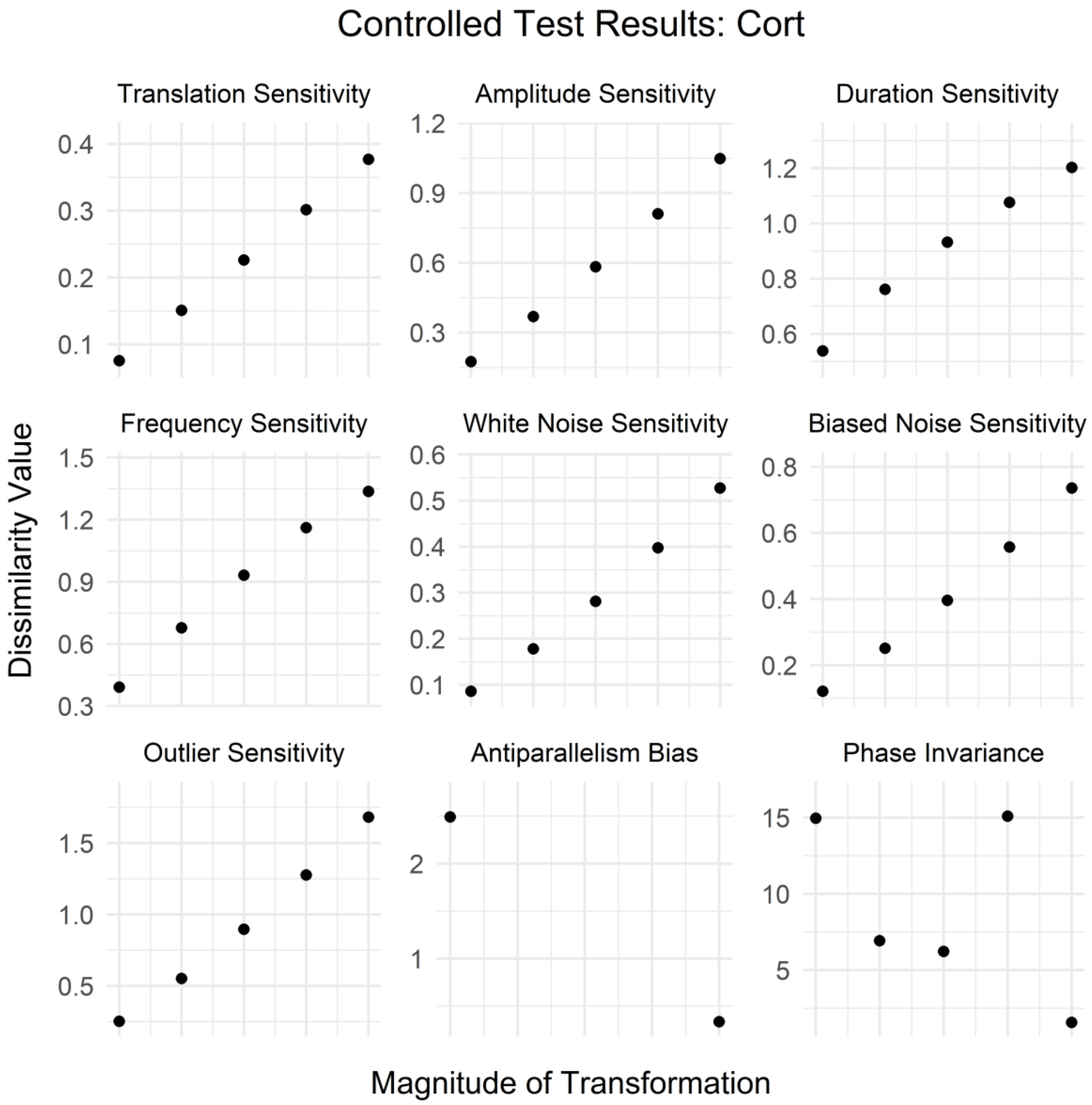
Controlled Test Results: Cort
Translation Sensitivity
Amplitude Sensitivity
Duration Sensitivity
Frequency Sensitivity
White Noise Sensitivity
Biased Noise Sensitivity
Outlier Sensitivity
Antiparallelism Bias
Phase Invariance
Dissimilarity Value
Magnitude of Transformation

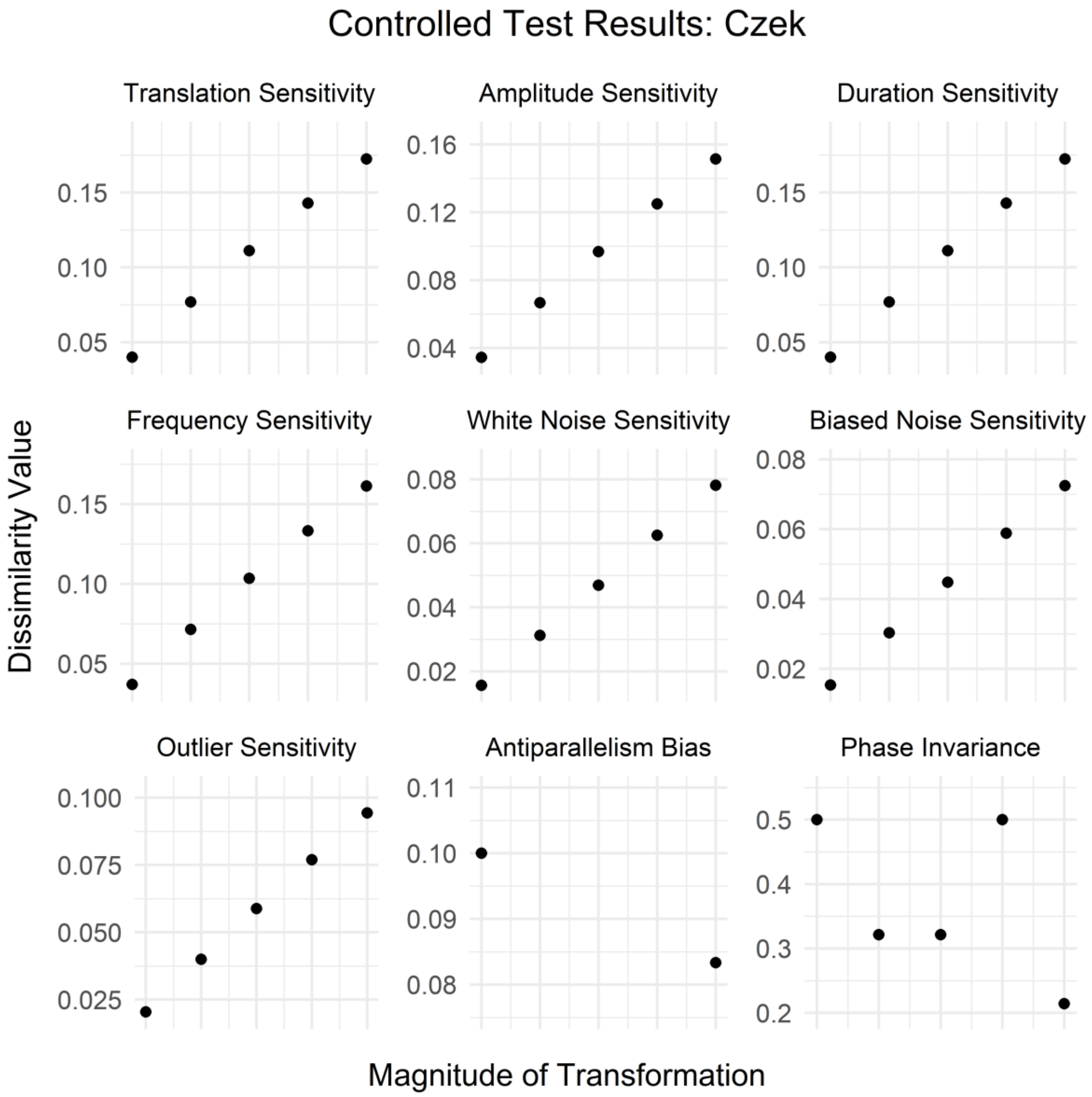

Controlled Test Results: Czek
Translation Sensitivity
Amplitude Sensitivity
Duration Sensitivity
Frequency Sensitivity
White Noise Sensitivity
Biased Noise Sensitivity
Outlier Sensitivity
Antiparallelism Bias
Phase Invariance
Dissimilarity Value
Magnitude of Transformation

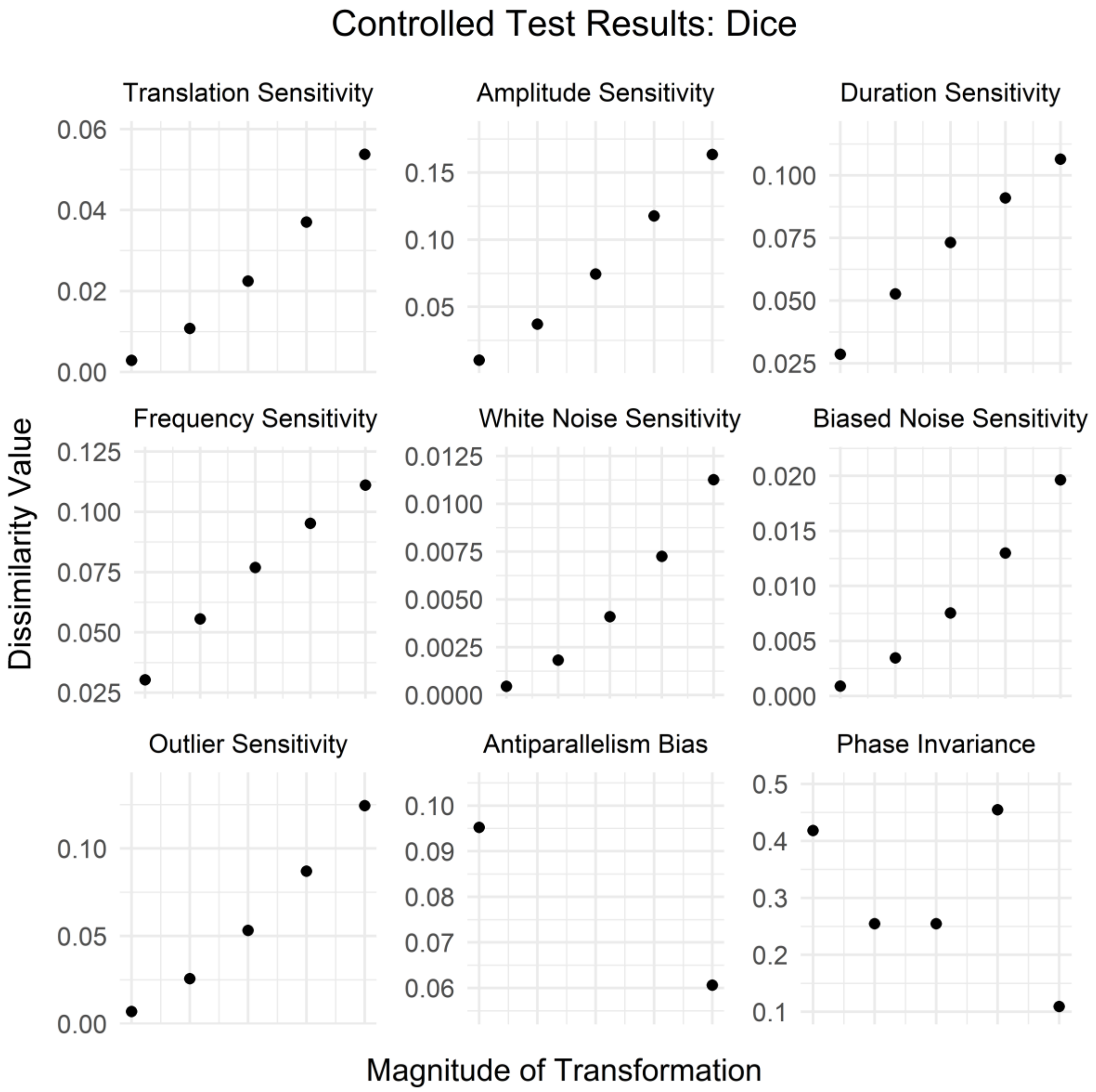
Controlled Test Results: Dice
Translation Sensitivity
Amplitude Sensitivity
Duration Sensitivity
Frequency Sensitivity
White Noise Sensitivity
Biased Noise Sensitivity
Outlier Sensitivity
Antiparallelism Bias
Phase Invariance
Dissimilarity Value
Magnitude of Transformation

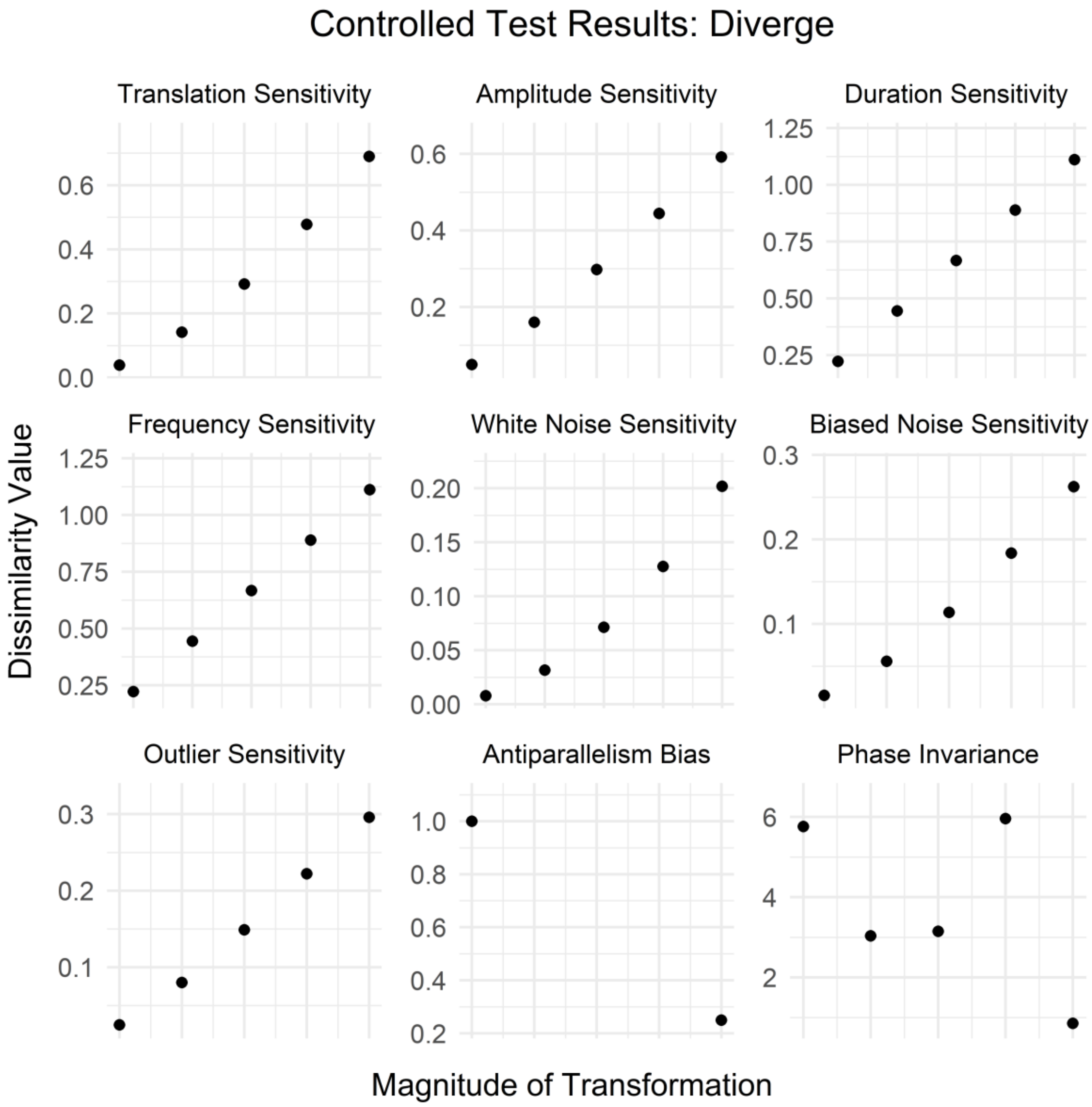
Controlled Test Results: Diverge
Translation Sensitivity
Amplitude Sensitivity
Duration Sensitivity
Frequency Sensitivity
White Noise Sensitivity
Biased Noise Sensitivity
Outlier Sensitivity
Antiparallelism Bias
Phase Invariance
Dissimilarity Value
Magnitude of Transformation

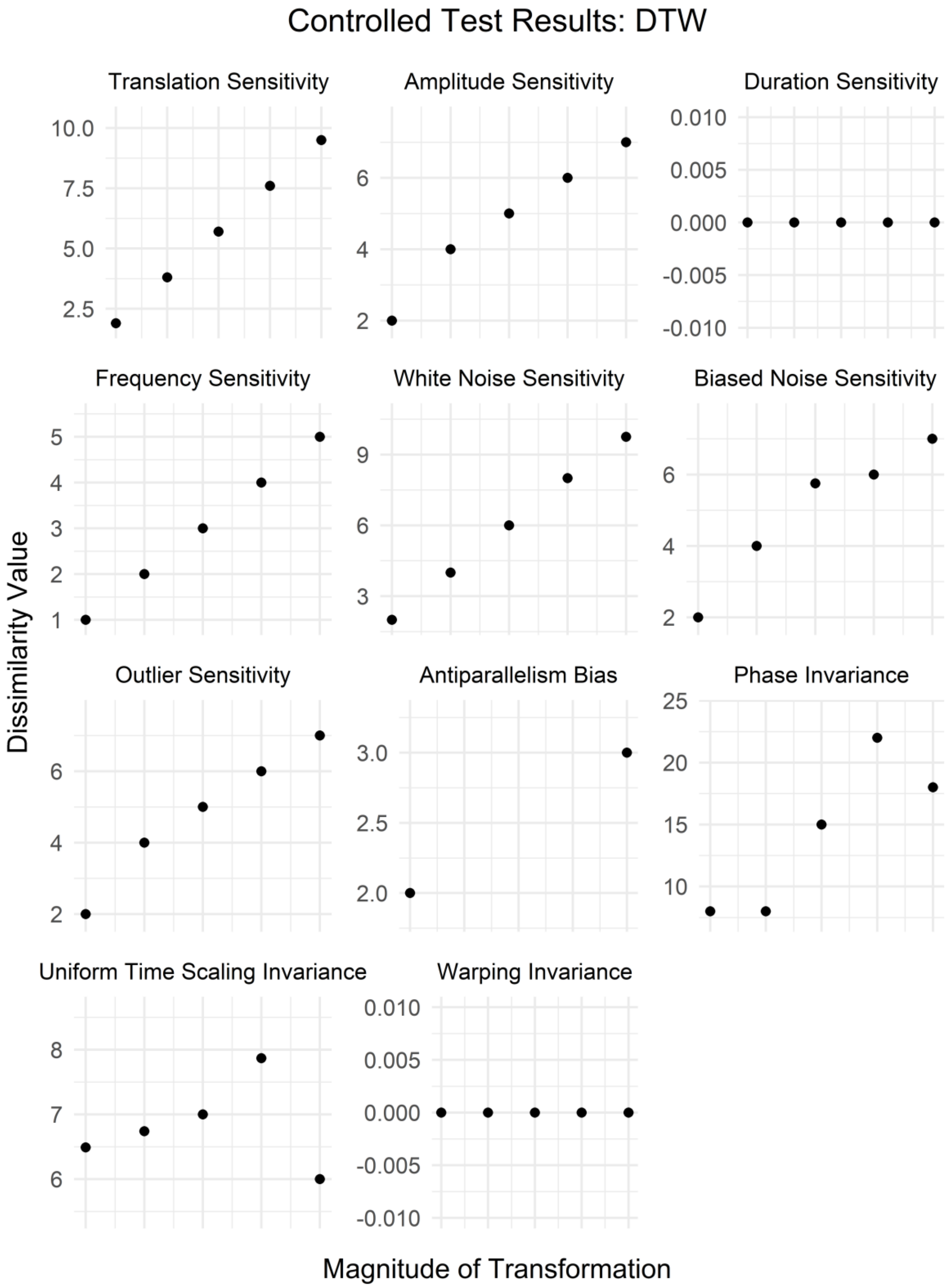
Controlled Test Results: DTW
Translation Sensitivity
Amplitude Sensitivity
Duration Sensitivity
Frequency Sensitivity
White Noise Sensitivity
Biased Noise Sensitivity
Outlier Sensitivity
Antiparallelism Bias
Phase Invariance
Uniform Time Scaling Invariance
Warping Invariance
Dissimilarity Value
Magnitude of Transformation

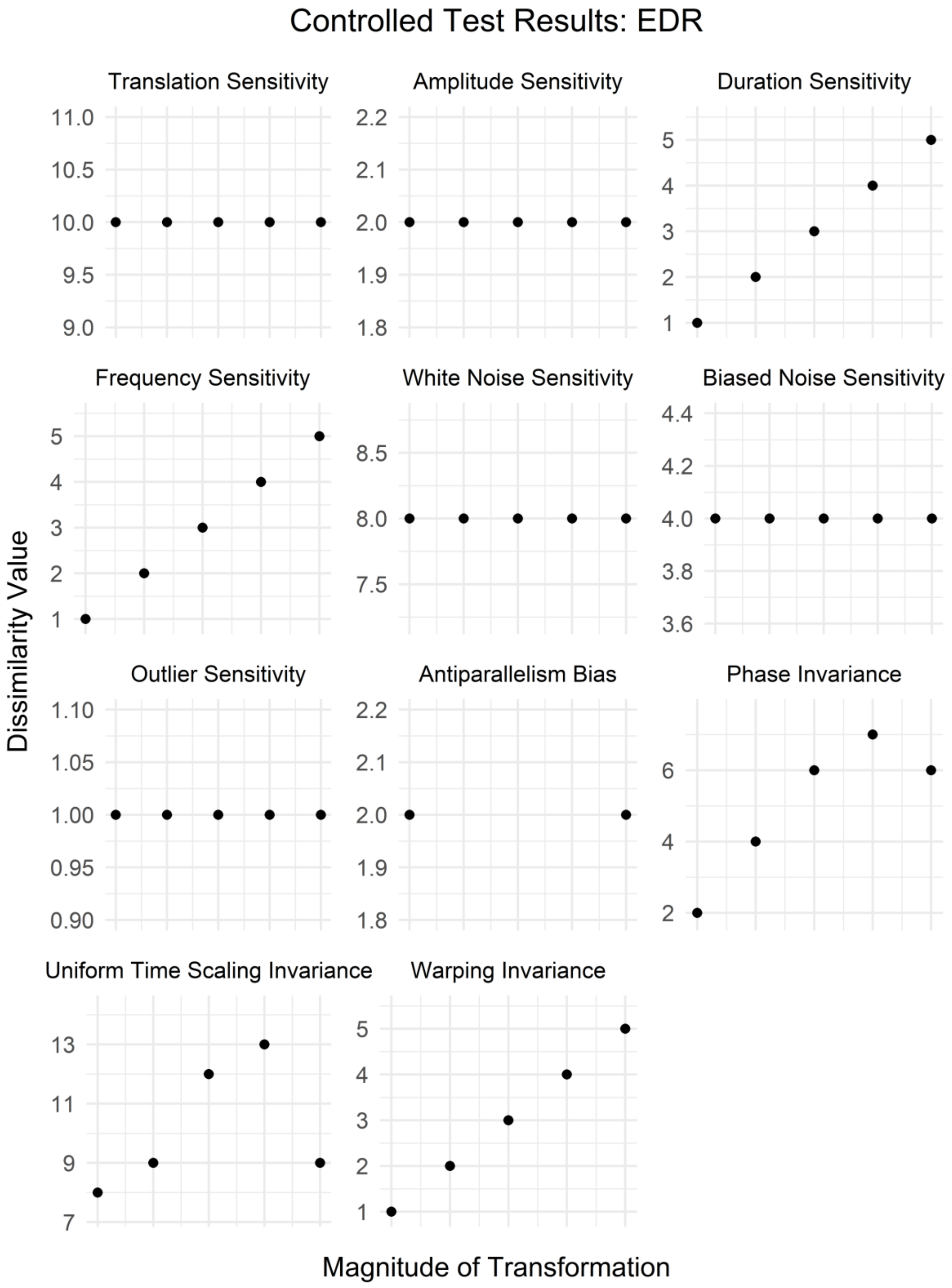
Controlled Test Results: EDR
Translation Sensitivity
Amplitude Sensitivity
Duration Sensitivity
Frequency Sensitivity
White Noise Sensitivity
Biased Noise Sensitivity
Outlier Sensitivity
Antiparallelism Bias
Phase Invariance
Uniform Time Scaling Invariance
Warping Invariance
Dissimilarity Value
Magnitude of Transformation

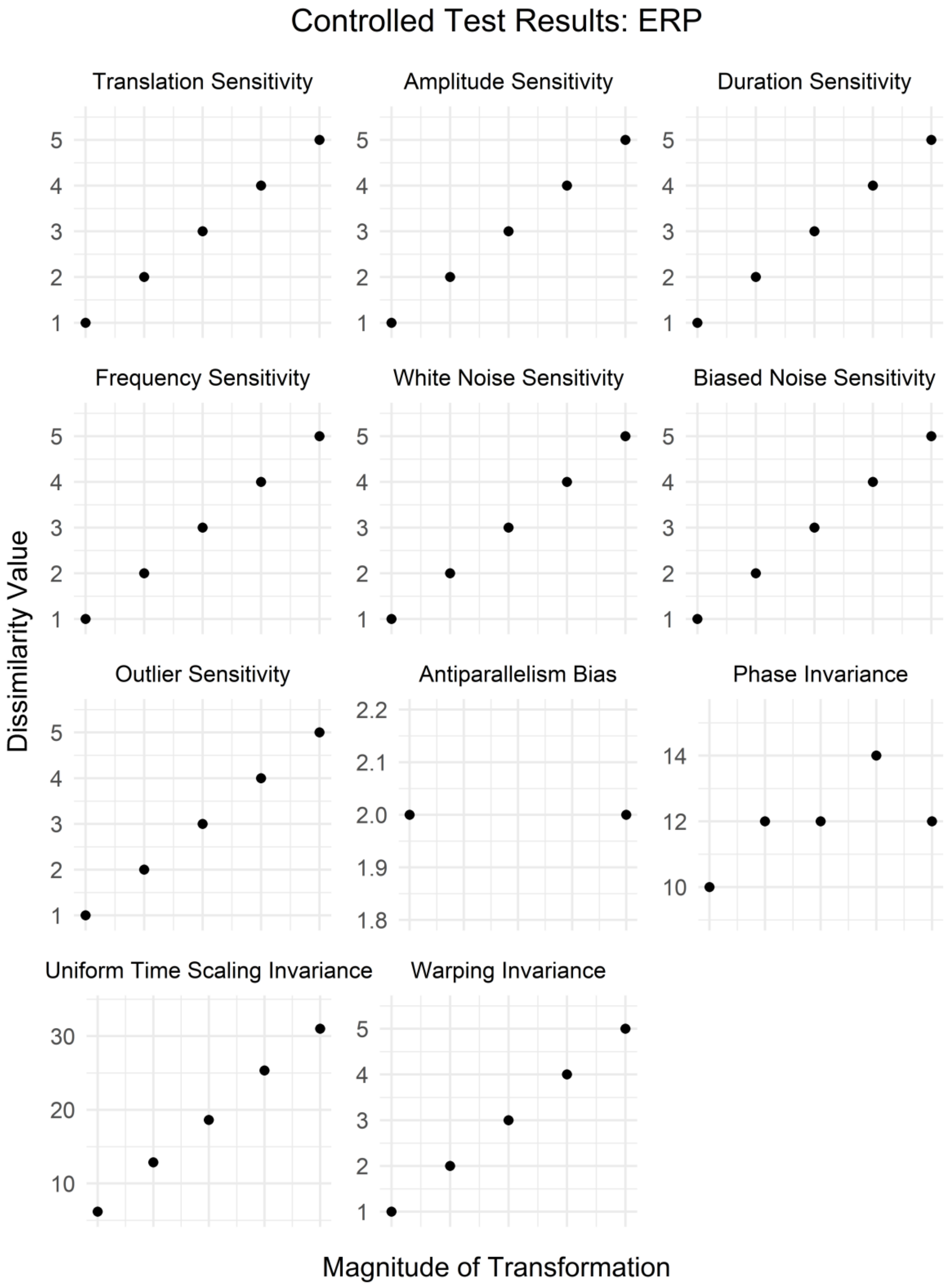
Controlled Test Results: ERP
Translation Sensitivity
Amplitude Sensitivity
Duration Sensitivity
Frequency Sensitivity
White Noise Sensitivity
Biased Noise Sensitivity
Outlier Sensitivity
Antiparallelism Bias
Phase Invariance
Uniform Time Scaling Invariance
Warping Invariance
Dissimilarity Value
Magnitude of Transformation

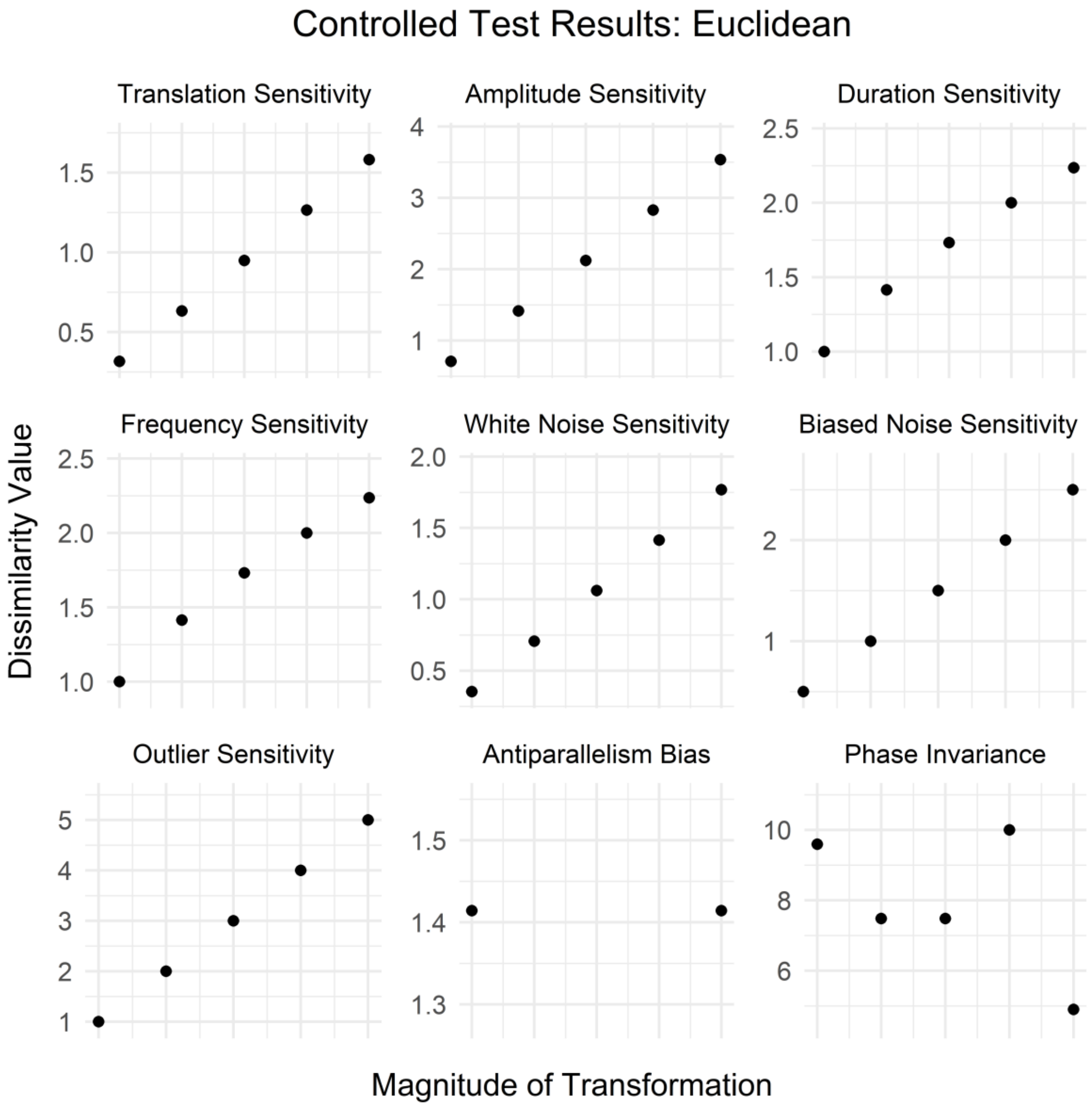
Controlled Test Results: Euclidean
Translation Sensitivity
Amplitude Sensitivity
Duration Sensitivity
Frequency Sensitivity
White Noise Sensitivity
Biased Noise Sensitivity
Outlier Sensitivity
Antiparallelism Bias
Phase Invariance
Dissimilarity Value
Magnitude of Transformation

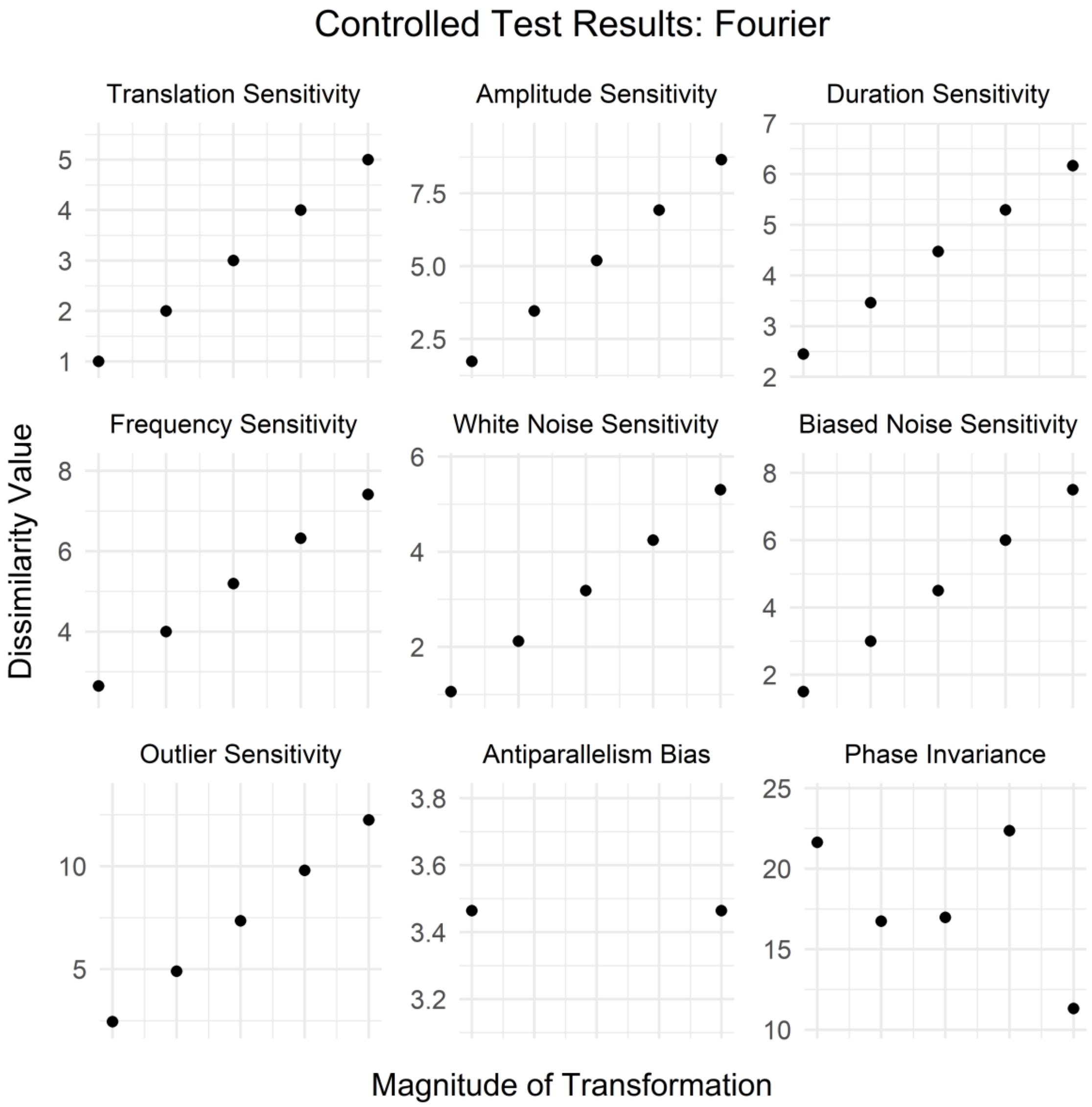
Controlled Test Results: Fourier
Translation Sensitivity
Amplitude Sensitivity
Duration Sensitivity
Frequency Sensitivity
White Noise Sensitivity
Biased Noise Sensitivity
Outlier Sensitivity
Antiparallelism Bias
Phase Invariance
Dissimilarity Value
Magnitude of Transformation

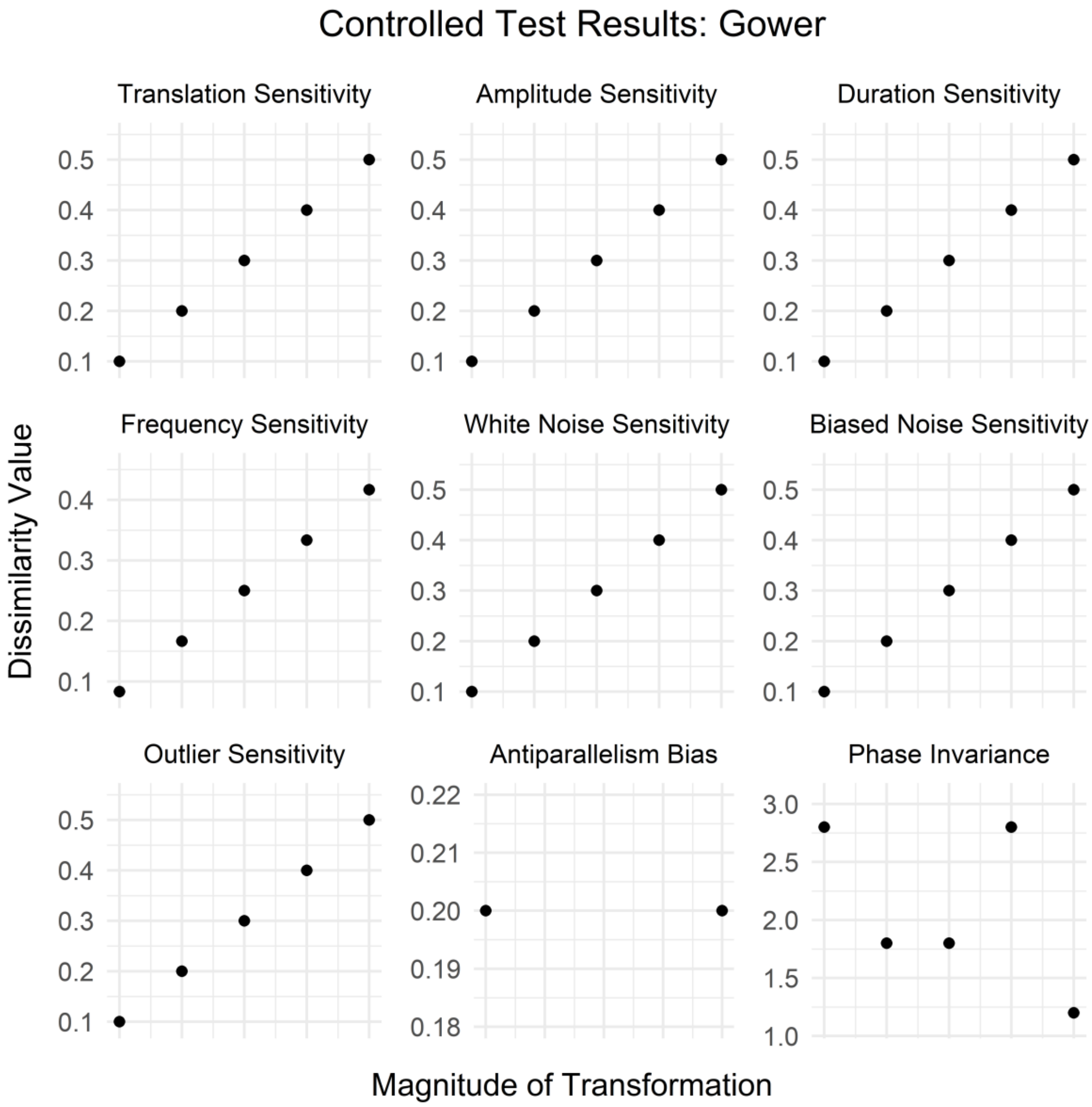
Controlled Test Results: Gower
Translation Sensitivity
Amplitude Sensitivity
Duration Sensitivity
Frequency Sensitivity
White Noise Sensitivity
Biased Noise Sensitivity
Outlier Sensitivity
Antiparallelism Bias
Phase Invariance
Dissimilarity Value
Magnitude of Transformation
0.1
0.2
0.3
0.4
0.5
0.18
0.19
0.20
0.21
0.22
1.0
1.5
2.0
2.5
3.0

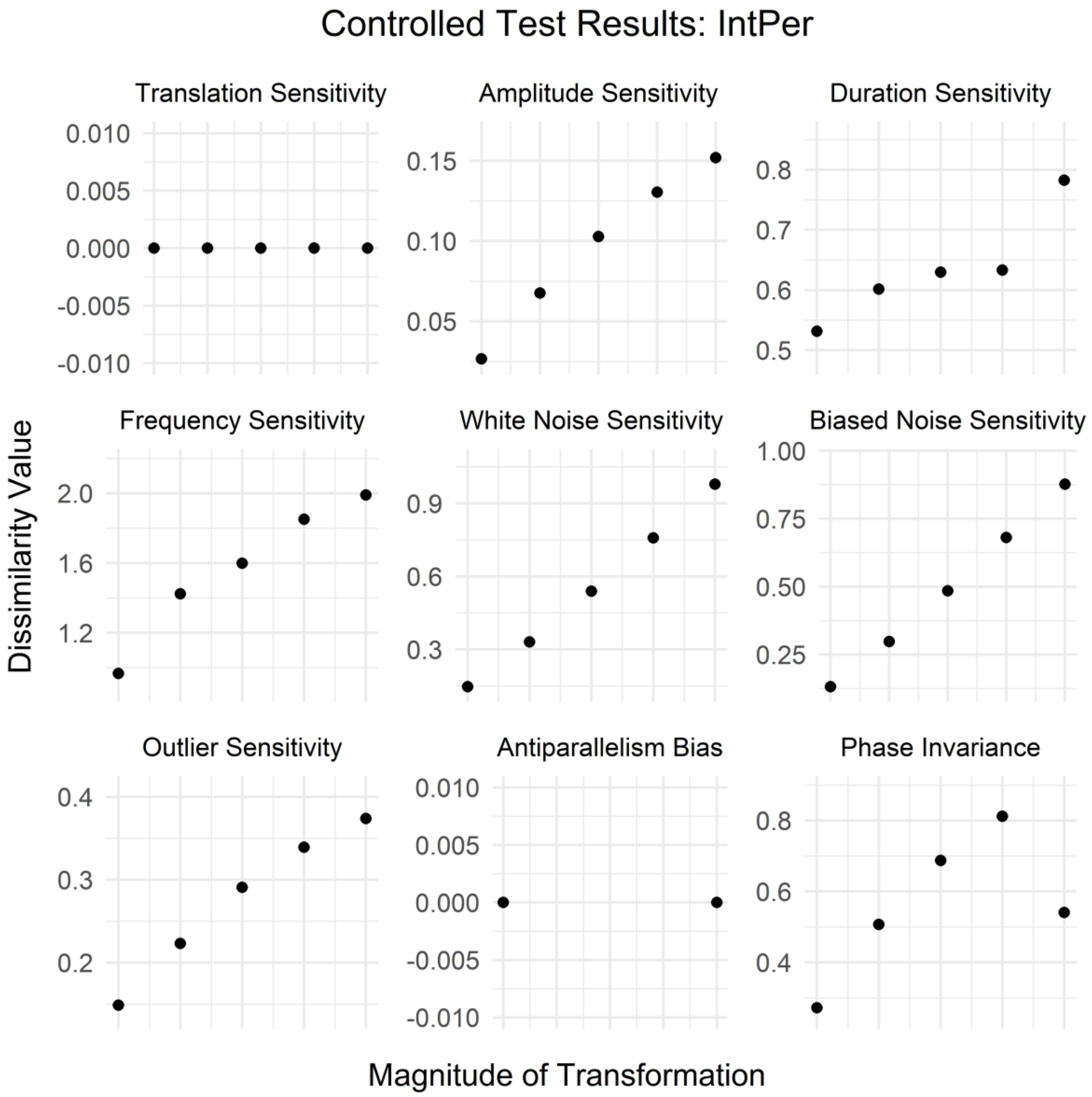

Controlled Test Results: IntPer
Translation Sensitivity
Amplitude Sensitivity
Duration Sensitivity
Frequency Sensitivity
White Noise Sensitivity
Biased Noise Sensitivity
Outlier Sensitivity
Antiparallelism Bias
Phase Invariance
Dissimilarity Value
Magnitude of Transformation
0.010
0.005
0.000
-0.005
-0.010
0.15
0.10
0.05
0.8
0.7
0.6
0.5
2.0
1.6
1.2
0.9
0.6
0.3
1.00
0.75
0.50
0.25
0.4
0.3
0.2
0.8
0.6
0.4

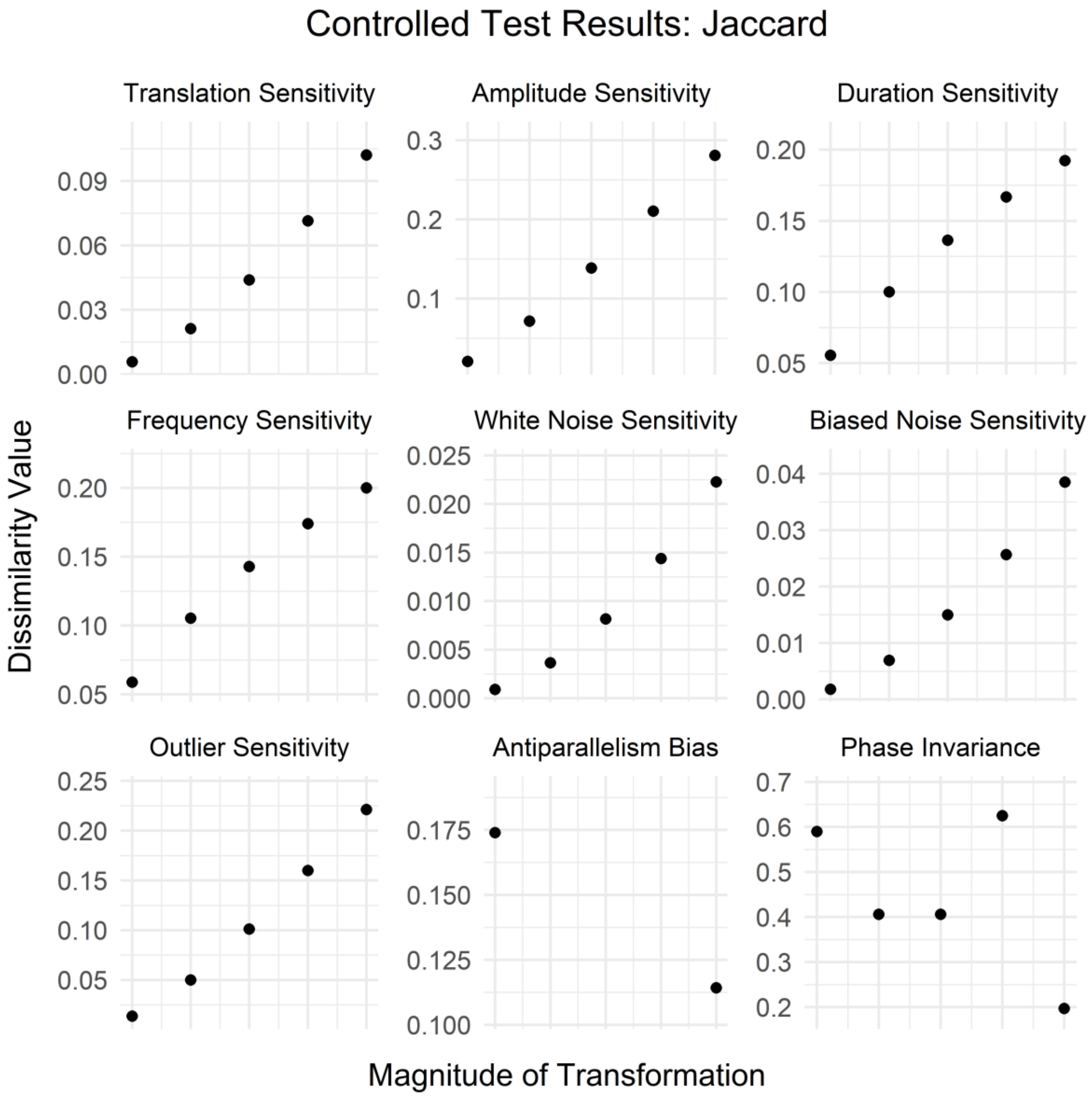
Controlled Test Results: Jaccard
Translation Sensitivity
Amplitude Sensitivity
Duration Sensitivity
Frequency Sensitivity
White Noise Sensitivity
Biased Noise Sensitivity
Outlier Sensitivity
Antiparallelism Bias
Phase Invariance
Dissimilarity Value
Magnitude of Transformation

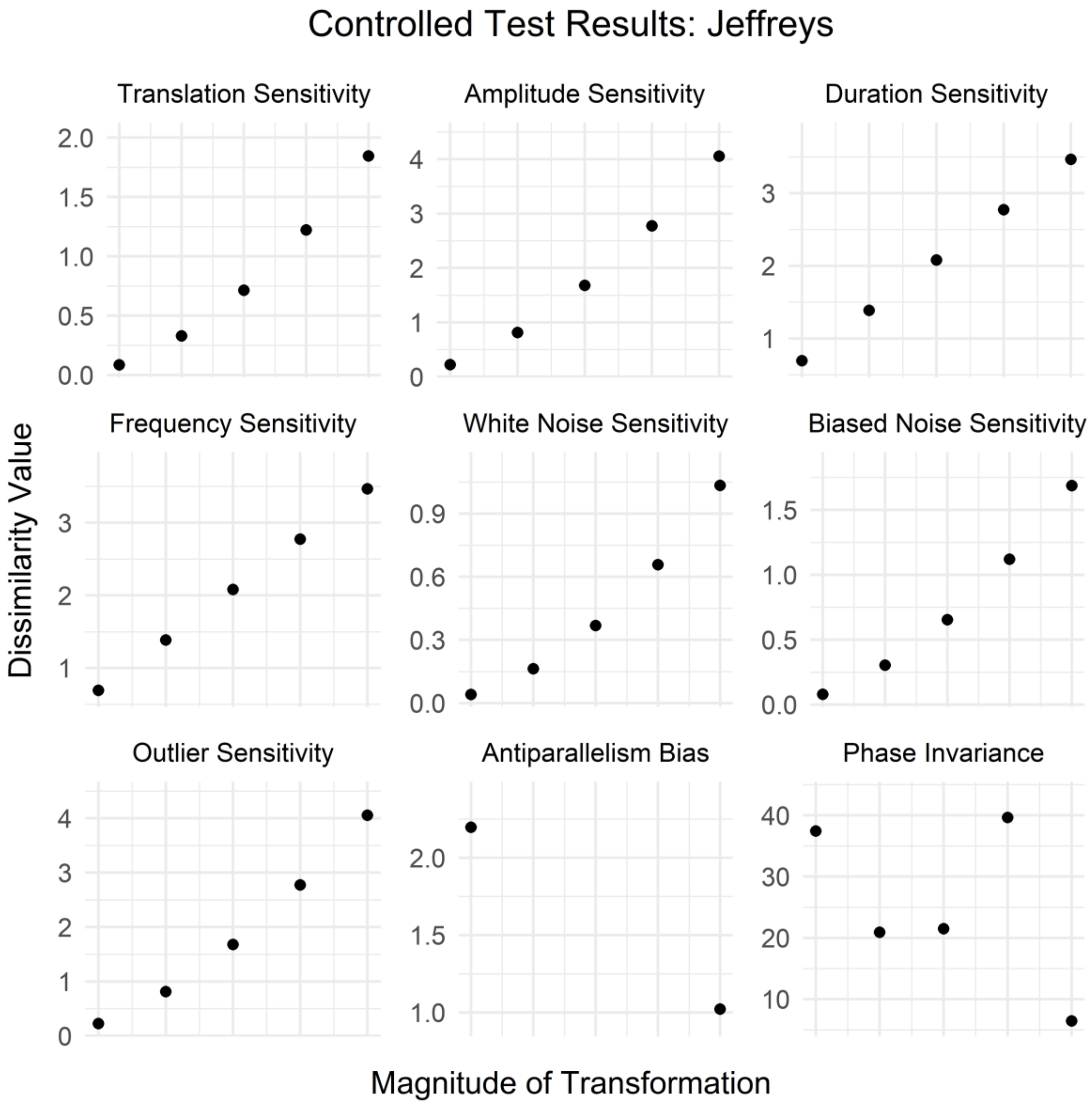
Controlled Test Results: Jeffreys
Translation Sensitivity
Amplitude Sensitivity
Duration Sensitivity
Frequency Sensitivity
White Noise Sensitivity
Biased Noise Sensitivity
Outlier Sensitivity
Antiparallelism Bias
Phase Invariance
Dissimilarity Value
Magnitude of Transformation

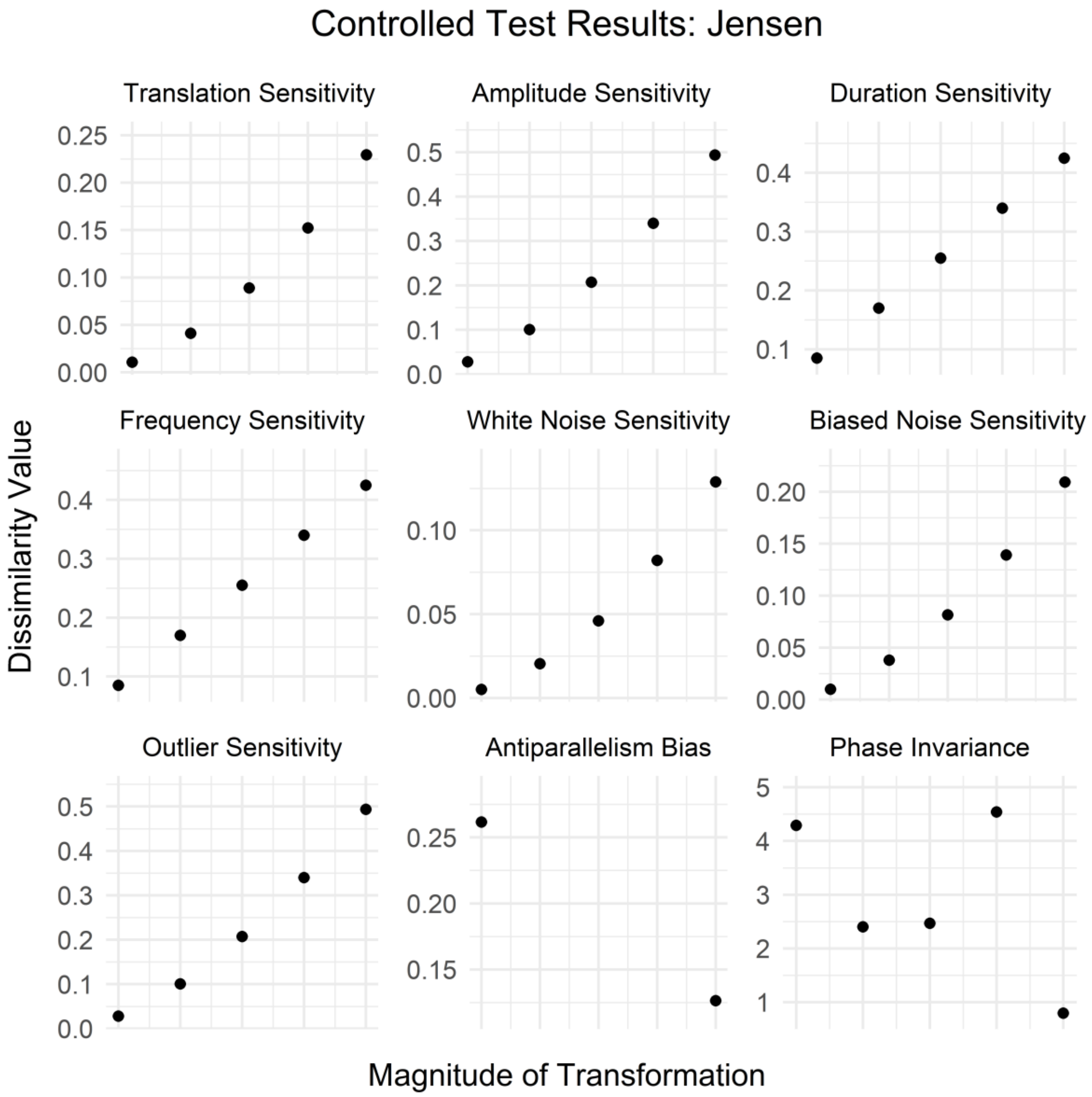
Controlled Test Results: Jensen
Translation Sensitivity
Amplitude Sensitivity
Duration Sensitivity
Frequency Sensitivity
White Noise Sensitivity
Biased Noise Sensitivity
Outlier Sensitivity
Antiparallelism Bias
Phase Invariance
Dissimilarity Value
Magnitude of Transformation

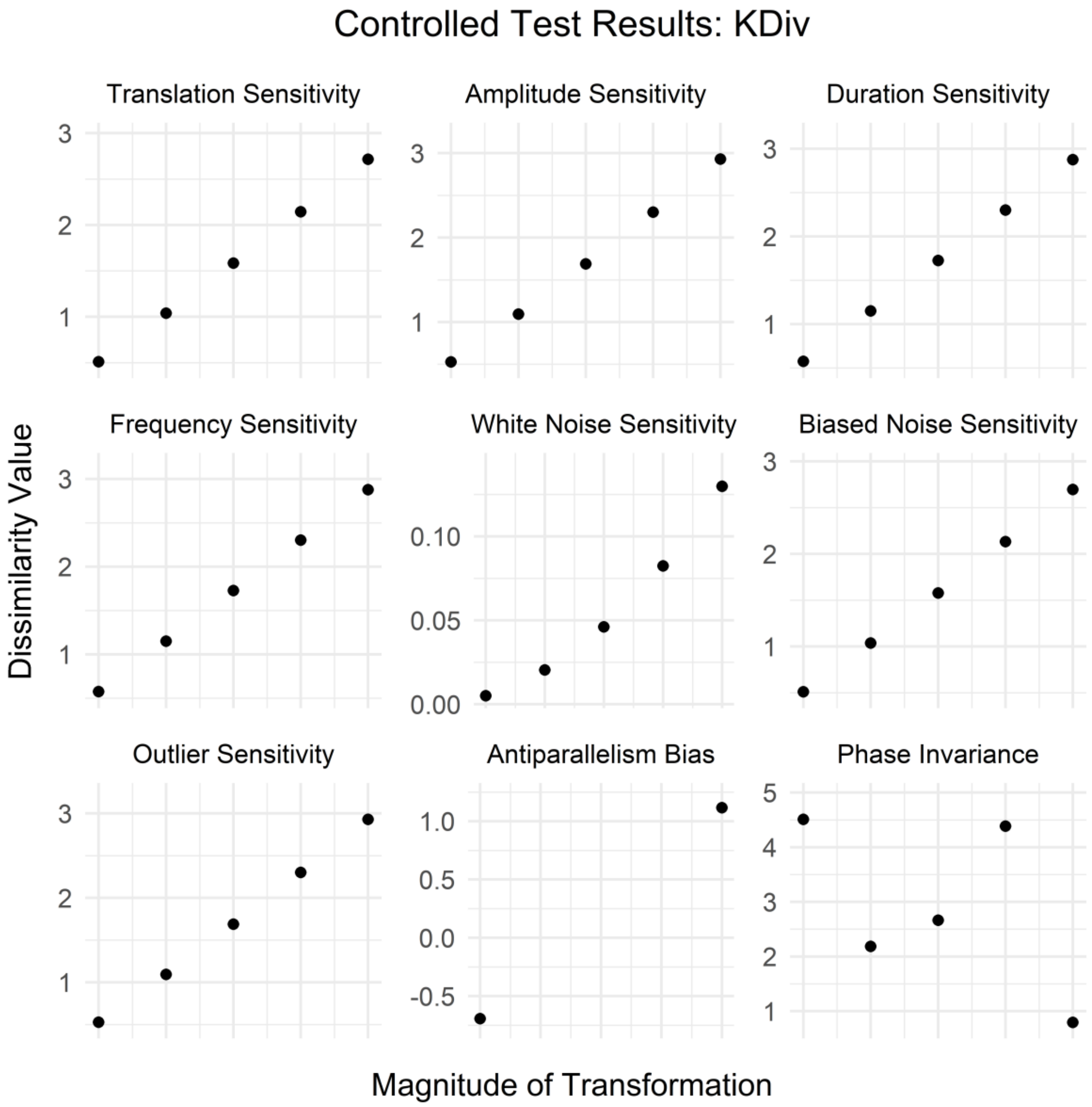
Controlled Test Results: KDiv
Translation Sensitivity
Amplitude Sensitivity
Duration Sensitivity
Frequency Sensitivity
White Noise Sensitivity
Biased Noise Sensitivity
Outlier Sensitivity
Antiparallelism Bias
Phase Invariance
Dissimilarity Value
Magnitude of Transformation
1
2
3
0.00
0.05
0.10
-0.5
0.0
0.5
1.0
4
5

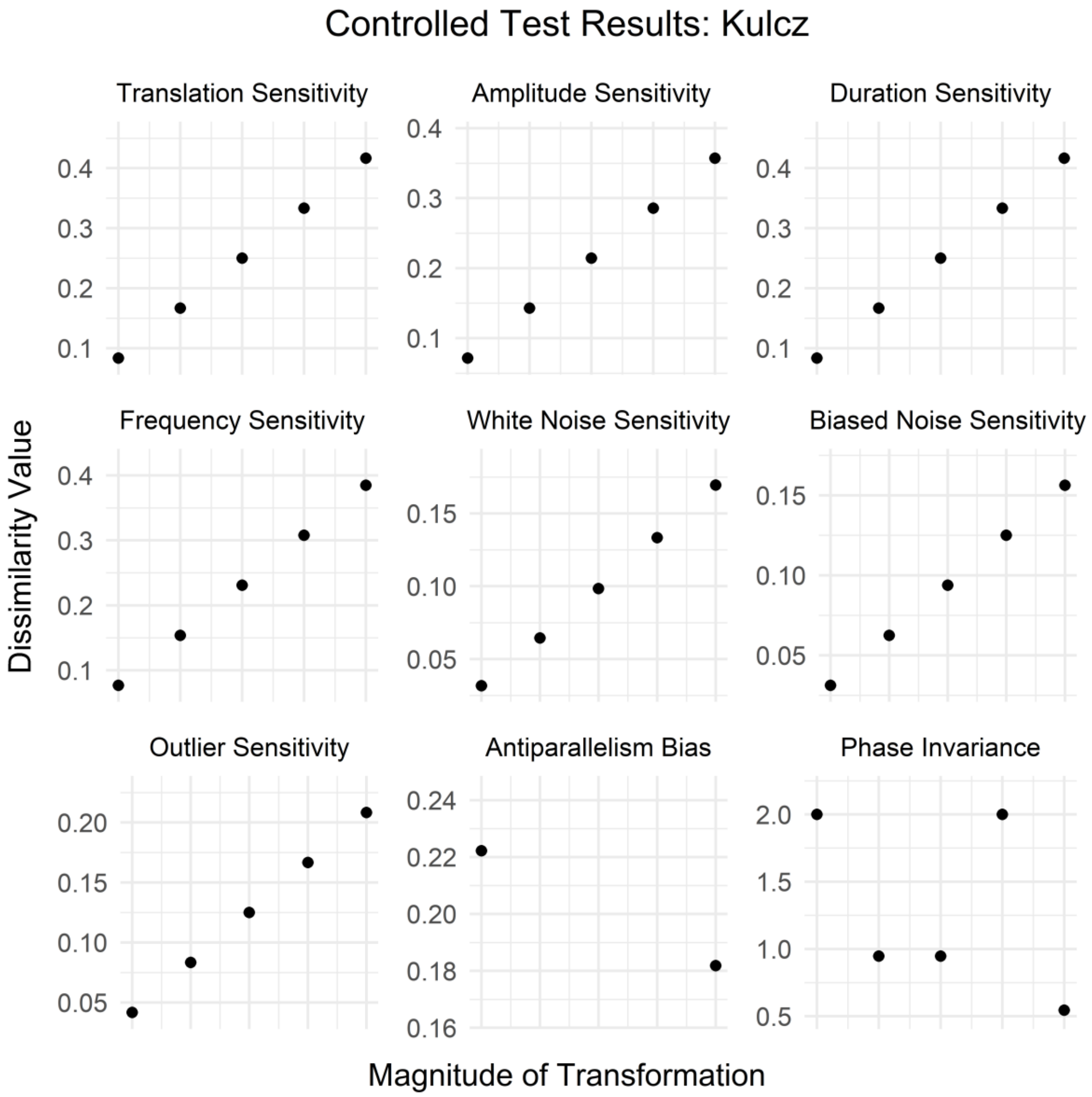

Controlled Test Results: Kulcz
Translation Sensitivity
Amplitude Sensitivity
Duration Sensitivity
Frequency Sensitivity
White Noise Sensitivity
Biased Noise Sensitivity
Outlier Sensitivity
Antiparallelism Bias
Phase Invariance
Dissimilarity Value
Magnitude of Transformation

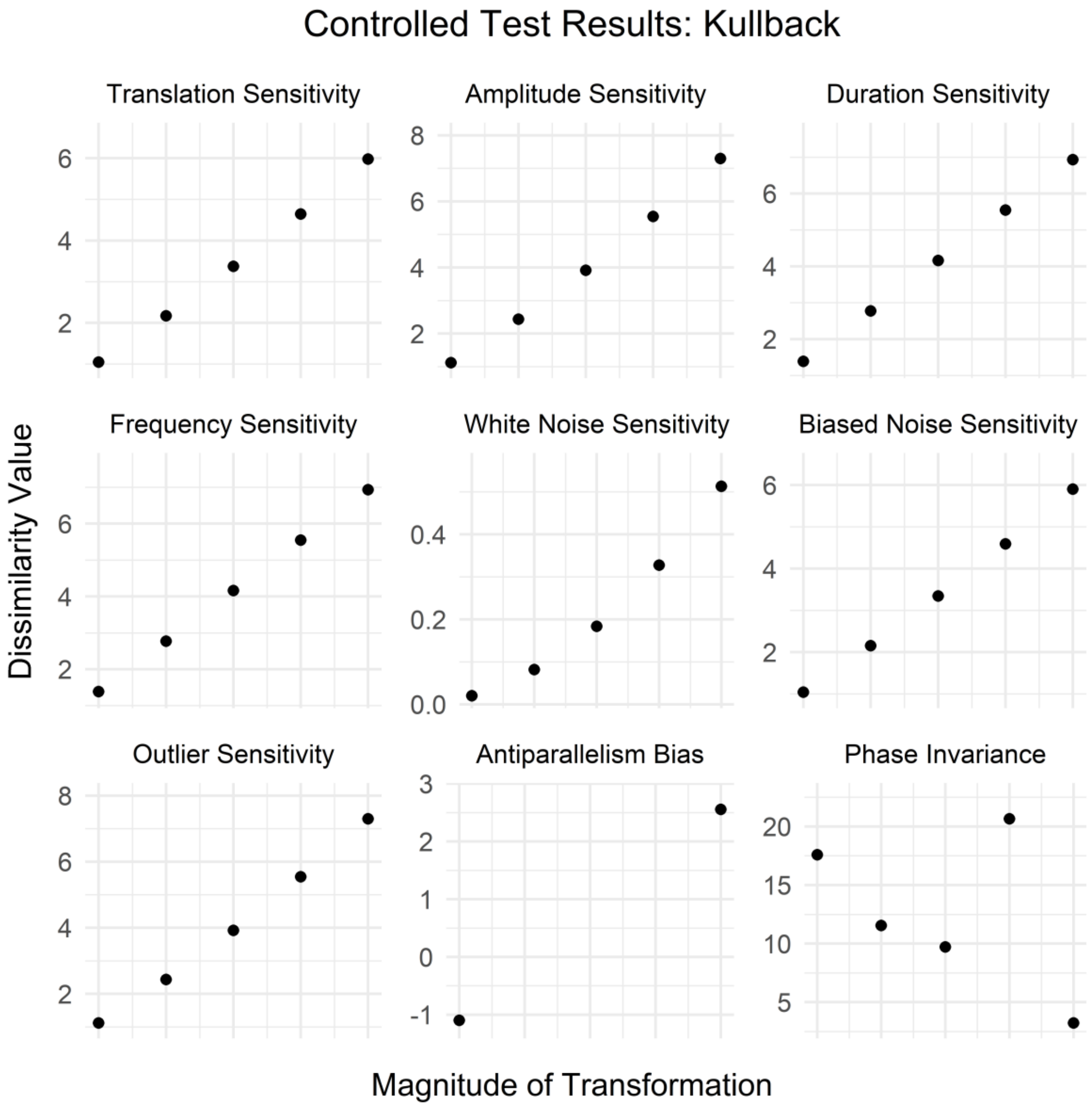
Controlled Test Results: Kullback
Translation Sensitivity
Amplitude Sensitivity
Duration Sensitivity
Frequency Sensitivity
White Noise Sensitivity
Biased Noise Sensitivity
Outlier Sensitivity
Antiparallelism Bias
Phase Invariance
Dissimilarity Value
Magnitude of Transformation

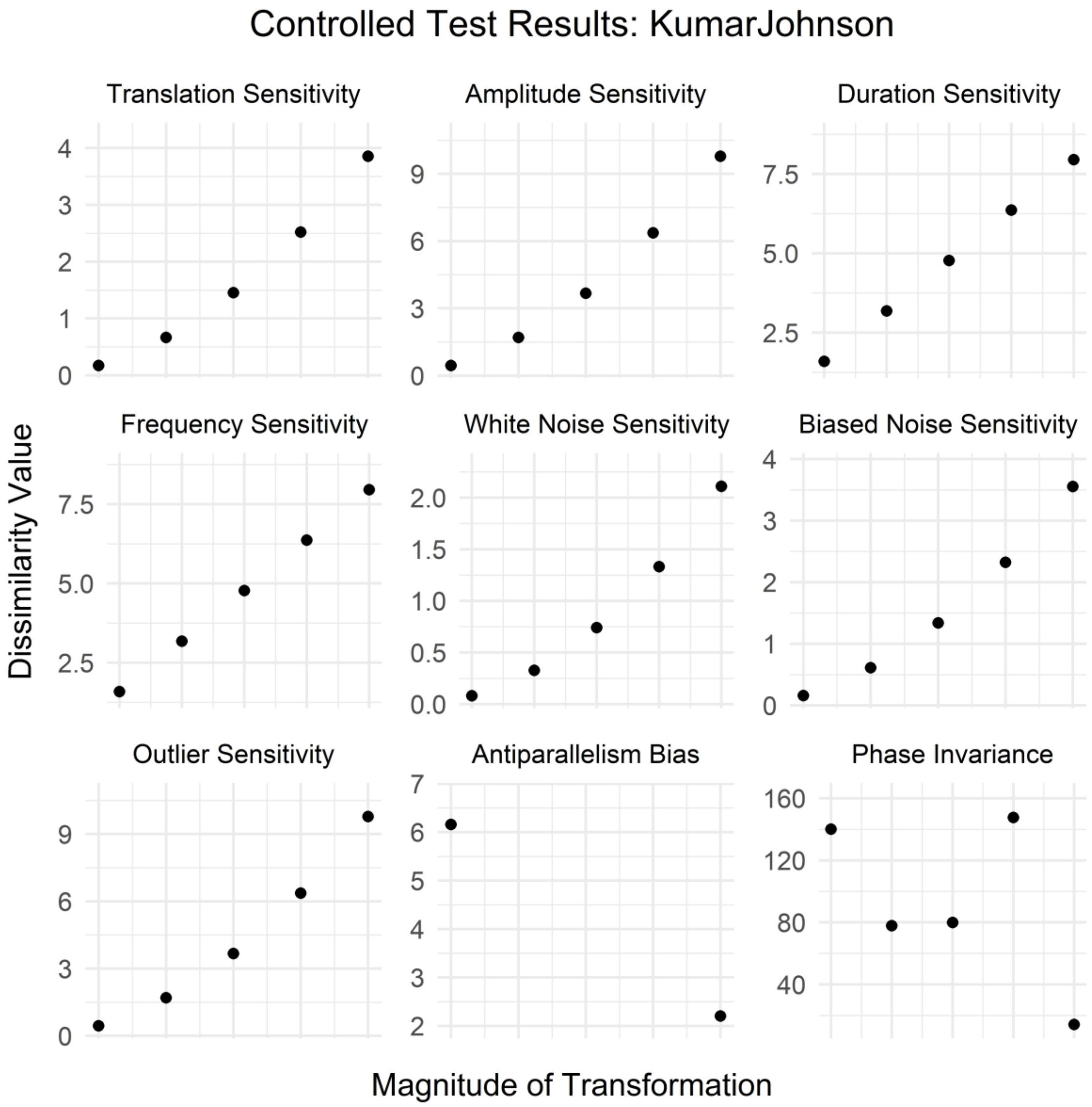
Controlled Test Results: KumarJohnson
Translation Sensitivity
Amplitude Sensitivity
Duration Sensitivity
Frequency Sensitivity
White Noise Sensitivity
Biased Noise Sensitivity
Outlier Sensitivity
Antiparallelism Bias
Phase Invariance
Dissimilarity Value
Magnitude of Transformation

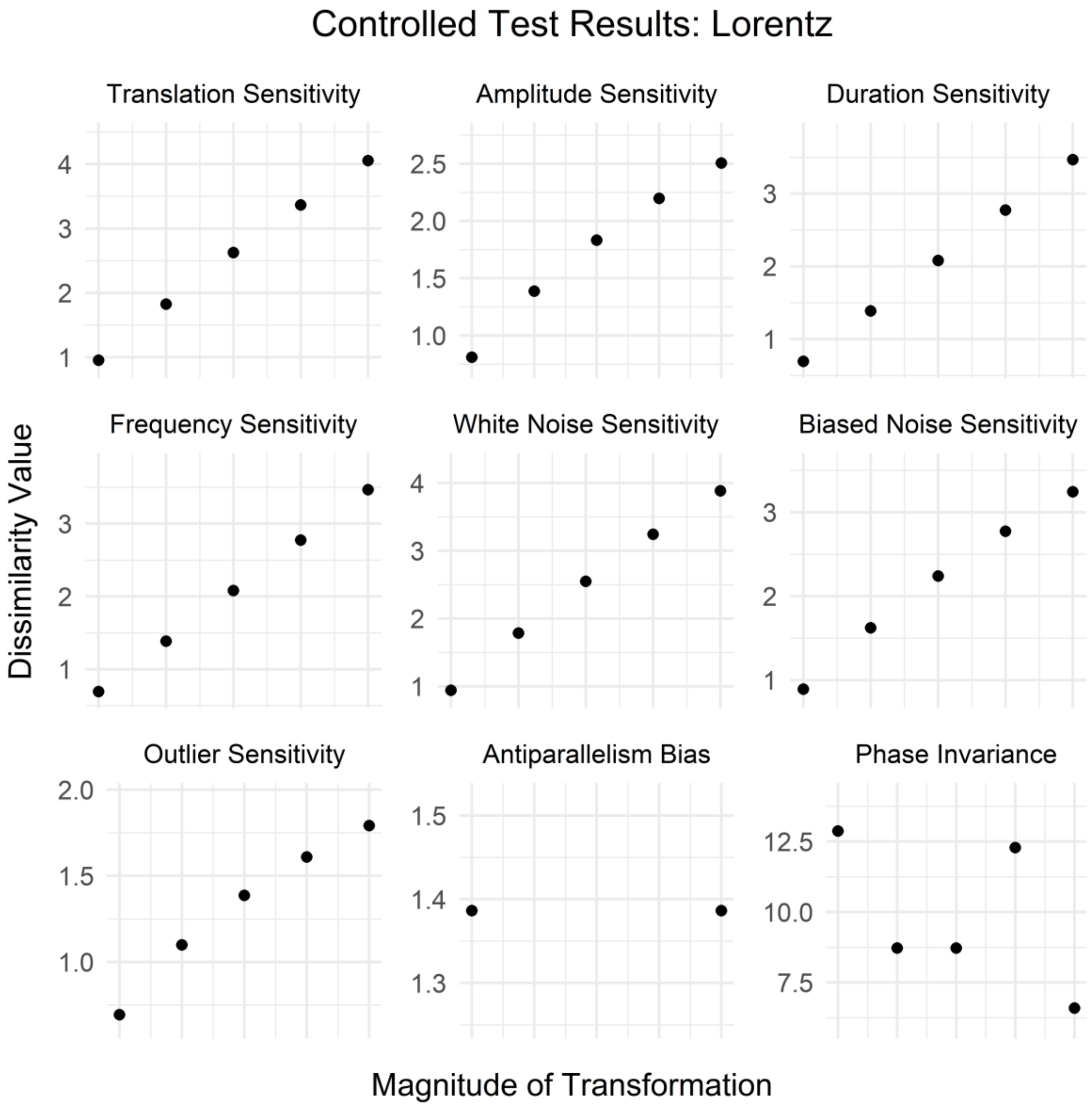

Controlled Test Results: Lorentz
Translation Sensitivity
Amplitude Sensitivity
Duration Sensitivity
Frequency Sensitivity
White Noise Sensitivity
Biased Noise Sensitivity
Outlier Sensitivity
Antiparallelism Bias
Phase Invariance
Dissimilarity Value
Magnitude of Transformation

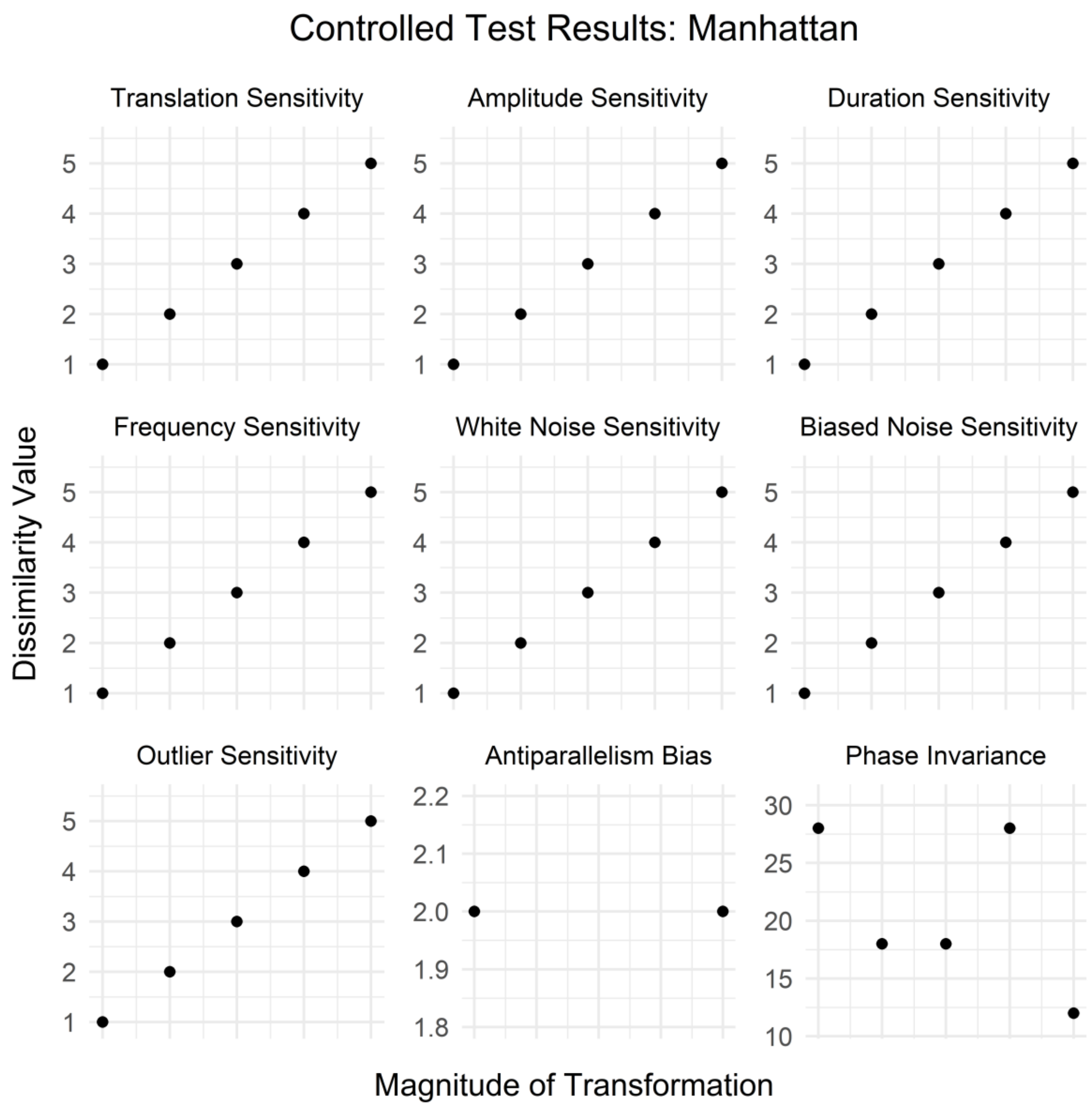
Controlled Test Results: Manhattan
Translation Sensitivity
Amplitude Sensitivity
Duration Sensitivity
Frequency Sensitivity
White Noise Sensitivity
Biased Noise Sensitivity
Outlier Sensitivity
Antiparallelism Bias
Phase Invariance
Dissimilarity Value
Magnitude of Transformation

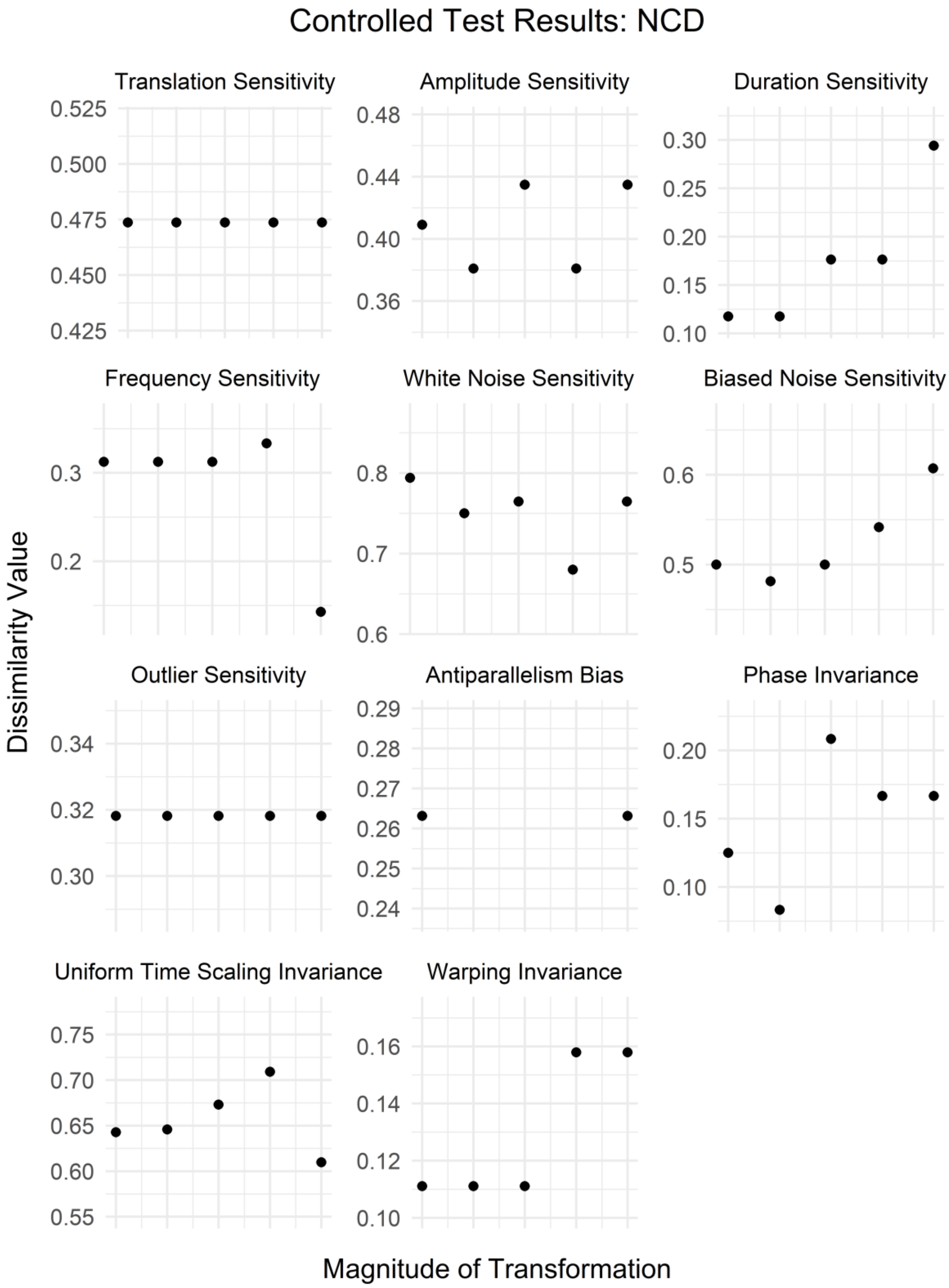
Controlled Test Results: NCD
Translation Sensitivity
0.525
0.500
0.475
0.450
0.425
Amplitude Sensitivity
0.48
0.44
0.40
0.36
Duration Sensitivity
0.30
0.25
0.20
0.15
0.10
Frequency Sensitivity
0.3
0.2
White Noise Sensitivity
0.8
0.7
0.6
Biased Noise Sensitivity
0.6
0.5
Outlier Sensitivity
0.34
0.32
0.30
Antiparallelism Bias
0.29
0.28
0.27
0.26
0.25
0.24
Phase Invariance
0.20
0.15
0.10
Uniform Time Scaling Invariance
0.75
0.70
0.65
0.60
0.55
Warping Invariance
0.16
0.14
0.12
0.10
Dissimilarity Value
Magnitude of Transformation

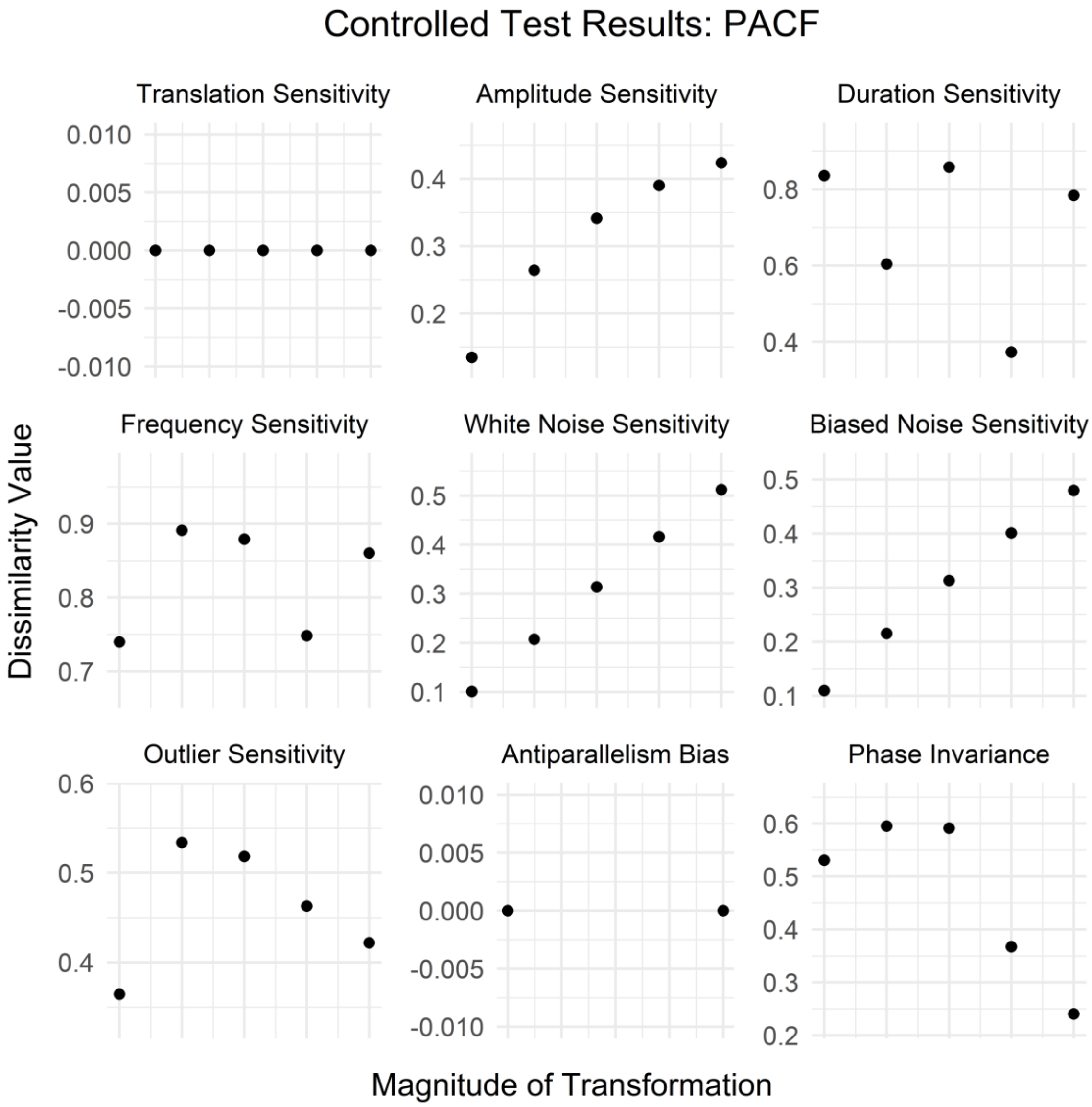
Controlled Test Results: PACF
Translation Sensitivity
Amplitude Sensitivity
Duration Sensitivity
Frequency Sensitivity
White Noise Sensitivity
Biased Noise Sensitivity
Outlier Sensitivity
Antiparallelism Bias
Phase Invariance
Dissimilarity Value
Magnitude of Transformation

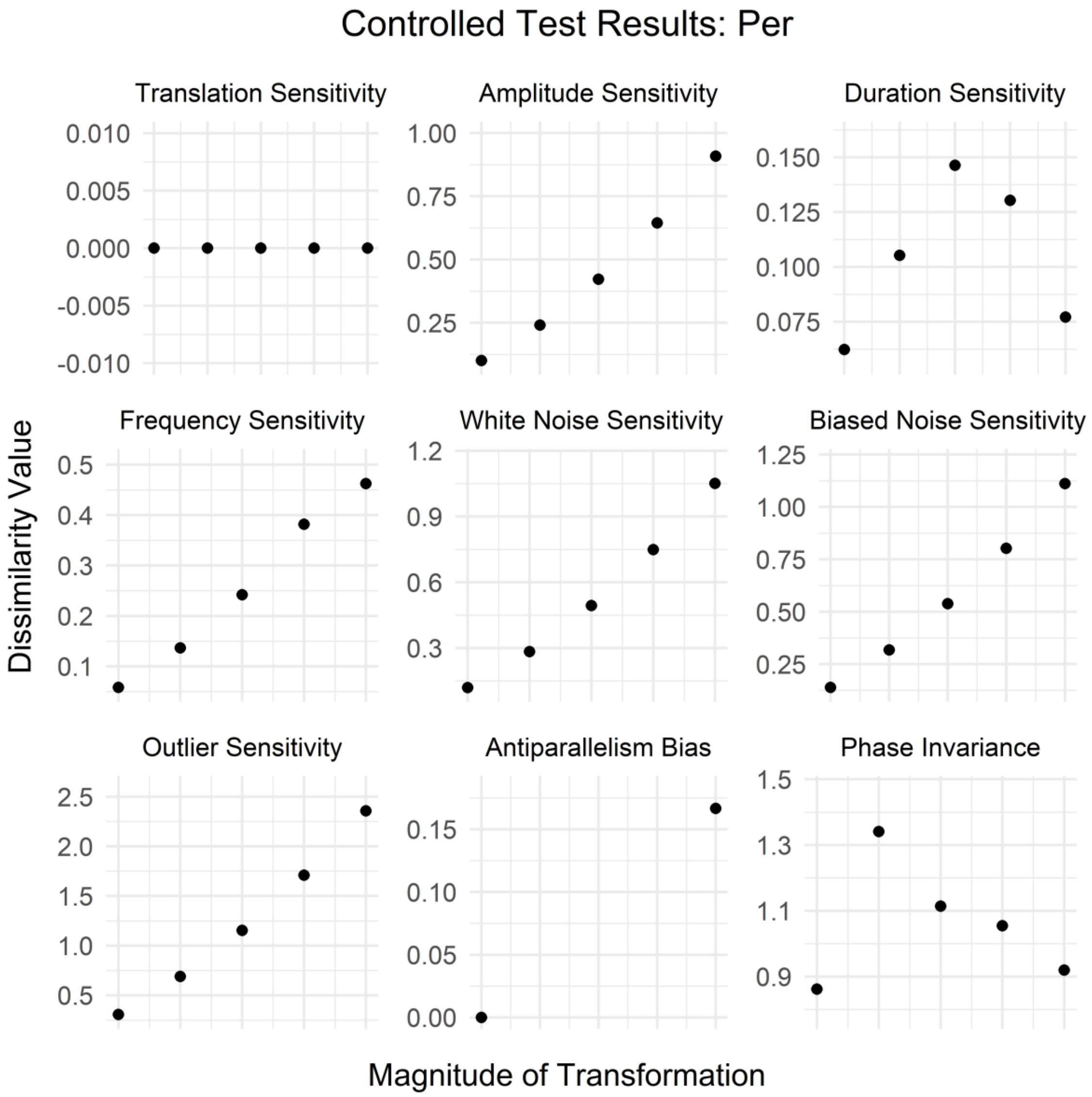
Controlled Test Results: Per
Translation Sensitivity
Amplitude Sensitivity
Duration Sensitivity
Frequency Sensitivity
White Noise Sensitivity
Biased Noise Sensitivity
Outlier Sensitivity
Antiparallelism Bias
Phase Invariance
Dissimilarity Value
Magnitude of Transformation

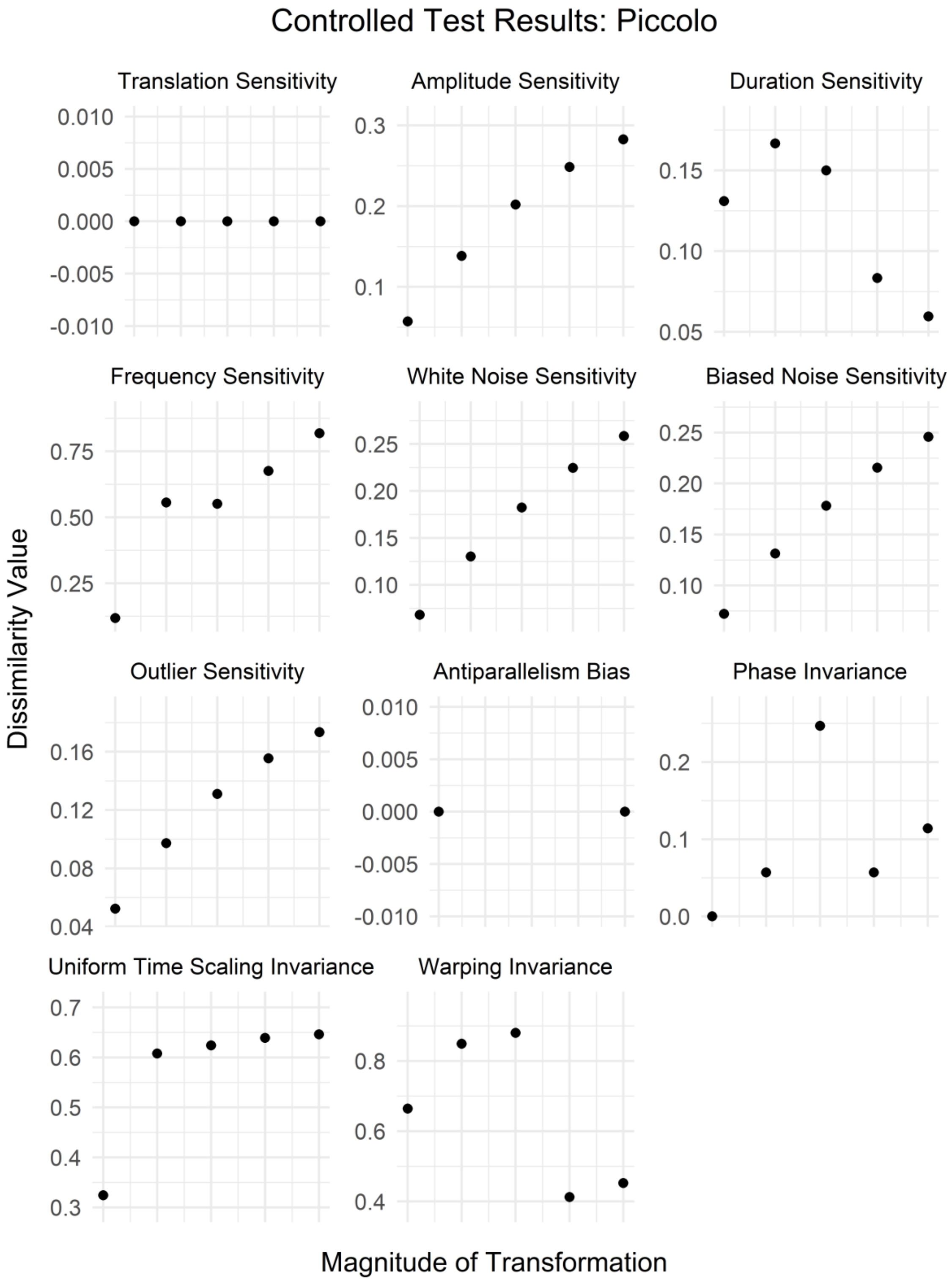
Controlled Test Results: Piccolo
Translation Sensitivity
Amplitude Sensitivity
Duration Sensitivity
Frequency Sensitivity
White Noise Sensitivity
Biased Noise Sensitivity
Outlier Sensitivity
Antiparallelism Bias
Phase Invariance
Uniform Time Scaling Invariance
Warping Invariance
Dissimilarity Value
Magnitude of Transformation

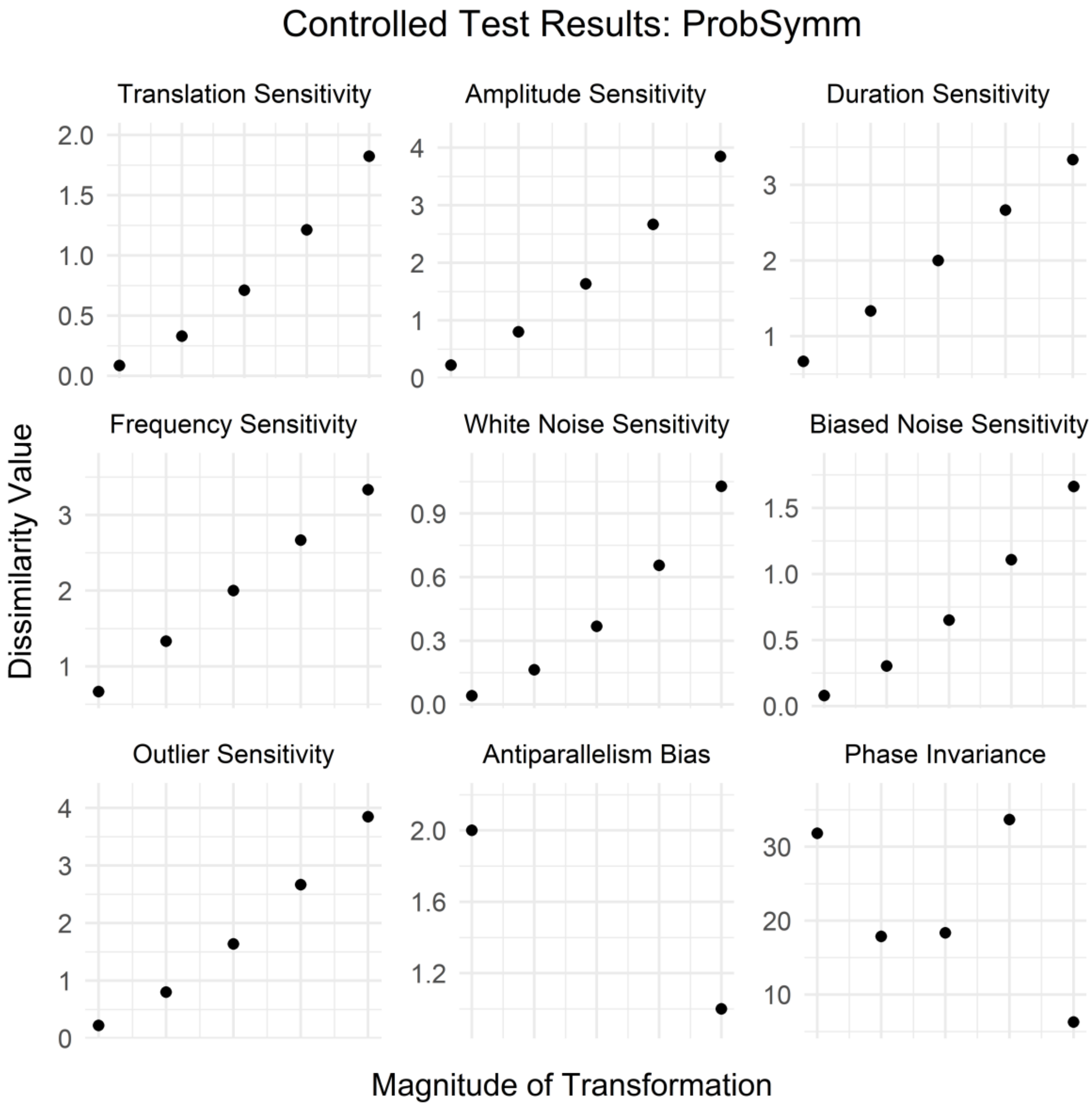
Controlled Test Results: ProbSymm
Translation Sensitivity
Amplitude Sensitivity
Duration Sensitivity
Frequency Sensitivity
White Noise Sensitivity
Biased Noise Sensitivity
Outlier Sensitivity
Antiparallelism Bias
Phase Invariance
Dissimilarity Value
Magnitude of Transformation

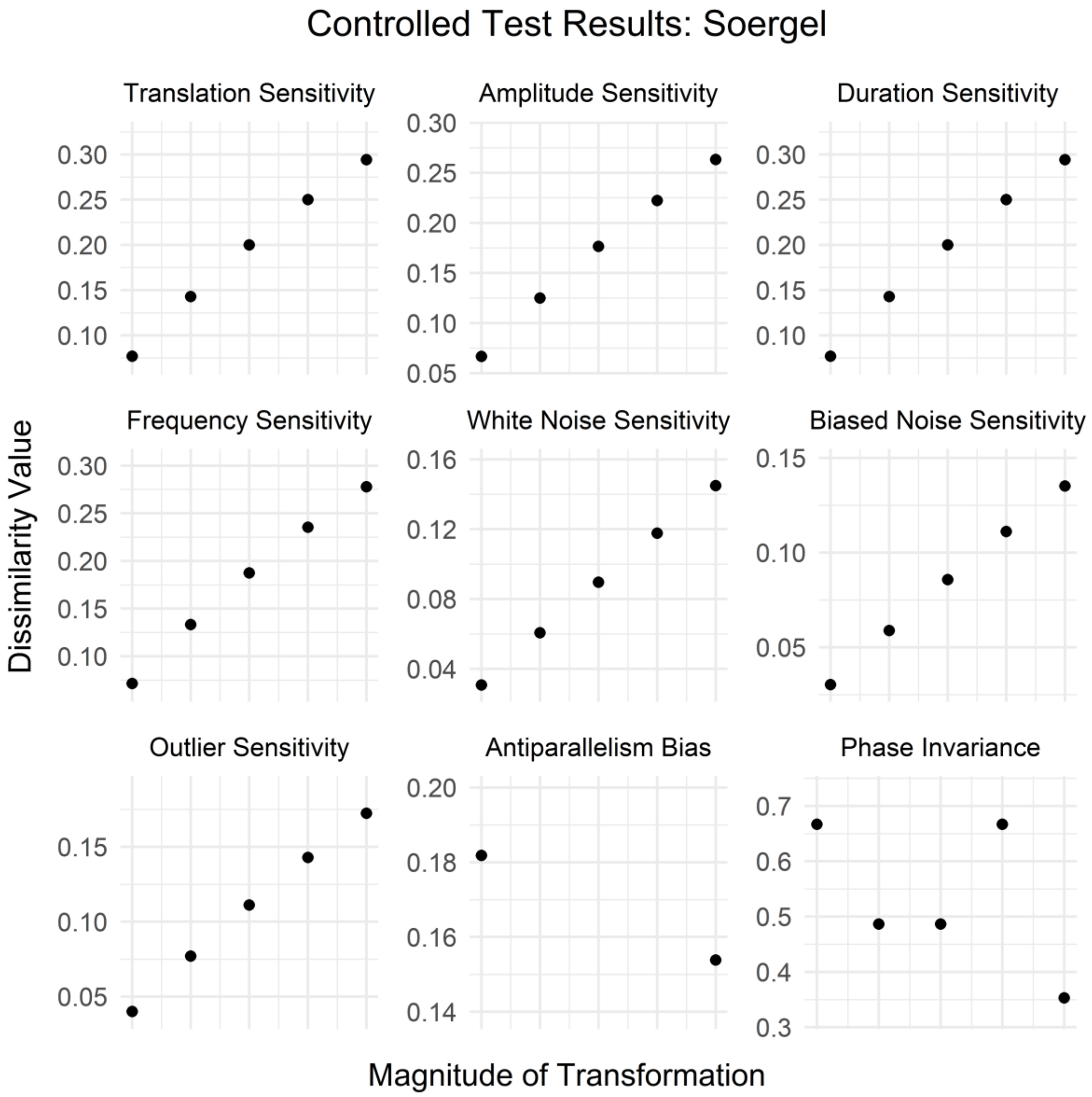
Controlled Test Results: Soergel
Translation Sensitivity
Amplitude Sensitivity
Duration Sensitivity
Frequency Sensitivity
White Noise Sensitivity
Biased Noise Sensitivity
Outlier Sensitivity
Antiparallelism Bias
Phase Invariance
Dissimilarity Value
Magnitude of Transformation

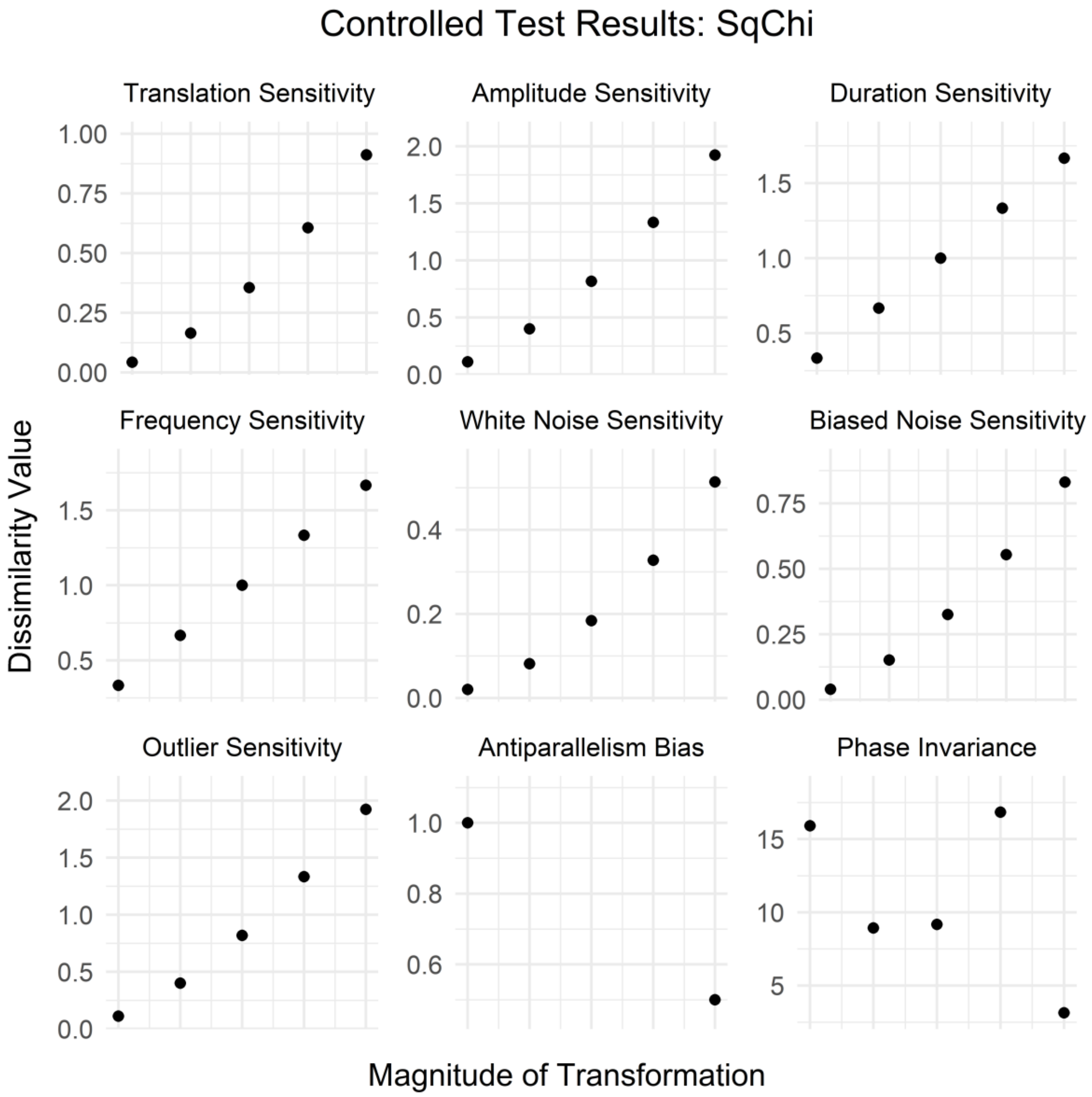
Controlled Test Results: SqChi
Translation Sensitivity
Amplitude Sensitivity
Duration Sensitivity
Frequency Sensitivity
White Noise Sensitivity
Biased Noise Sensitivity
Outlier Sensitivity
Antiparallelism Bias
Phase Invariance
Dissimilarity Value
Magnitude of Transformation
1.00
0.75
0.50
0.25
0.00
2.0
1.5
1.0
0.5
0.0
0.4
0.2
0.8
0.6
15
10
5

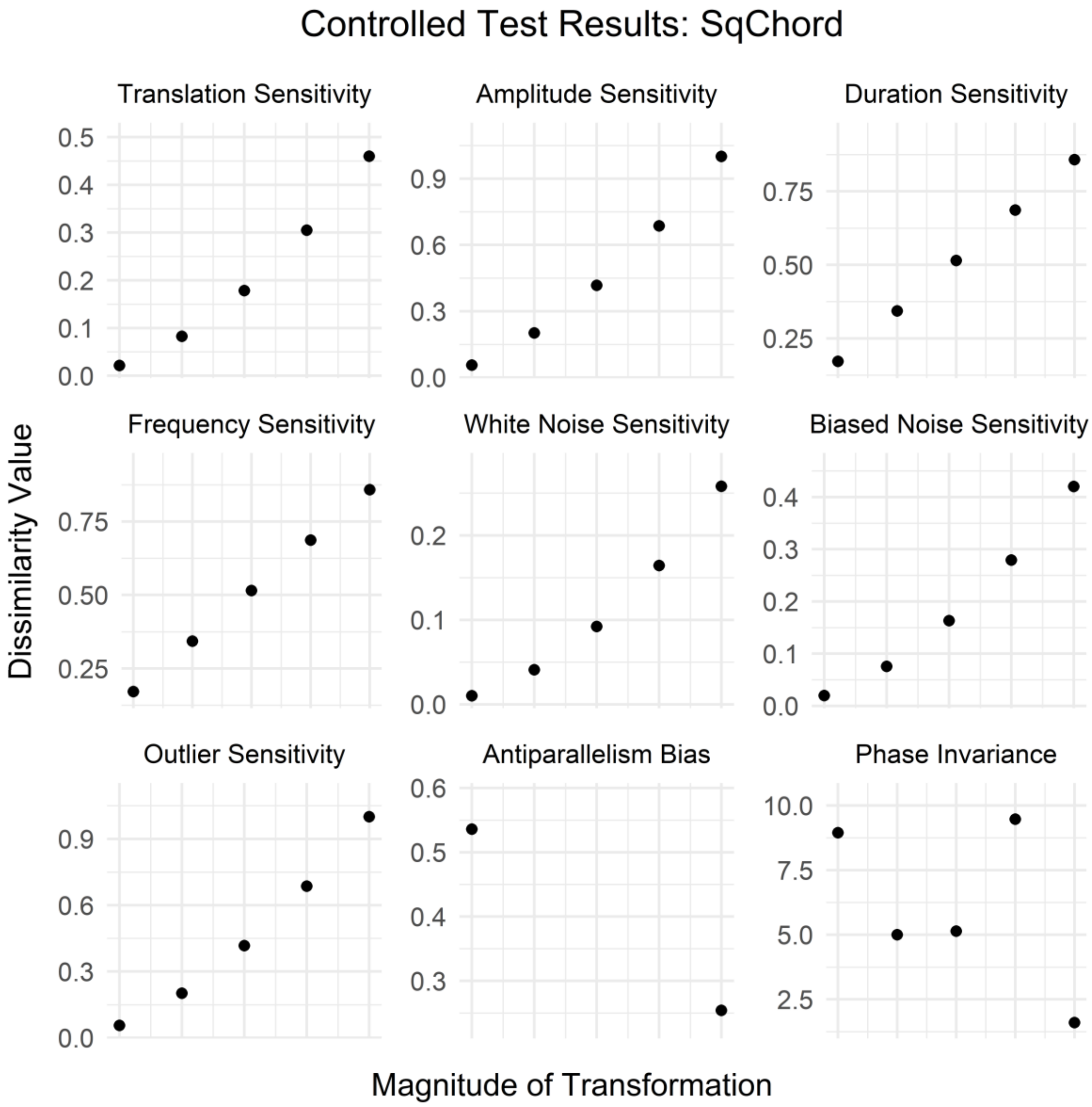
Controlled Test Results: SqChord
Translation Sensitivity
Amplitude Sensitivity
Duration Sensitivity
Frequency Sensitivity
White Noise Sensitivity
Biased Noise Sensitivity
Outlier Sensitivity
Antiparallelism Bias
Phase Invariance
Dissimilarity Value
Magnitude of Transformation
0.0
0.1
0.2
0.3
0.4
0.5
0.6
0.9
0.25
0.50
0.75
2.5
5.0
7.5
10.0

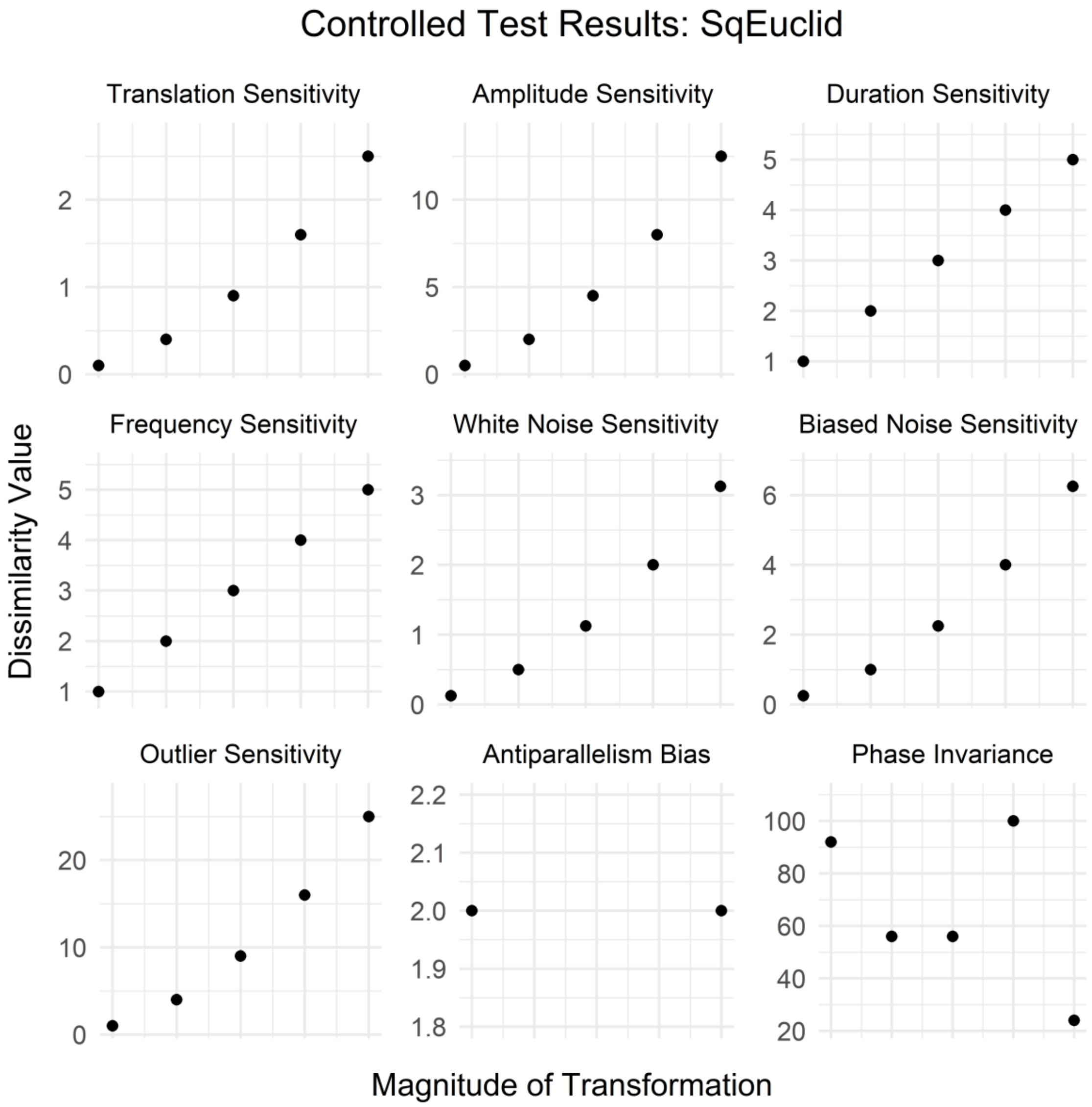
Controlled Test Results: SqEuclid
Translation Sensitivity
Amplitude Sensitivity
Duration Sensitivity
Frequency Sensitivity
White Noise Sensitivity
Biased Noise Sensitivity
Outlier Sensitivity
Antiparallelism Bias
Phase Invariance
Dissimilarity Value
Magnitude of Transformation

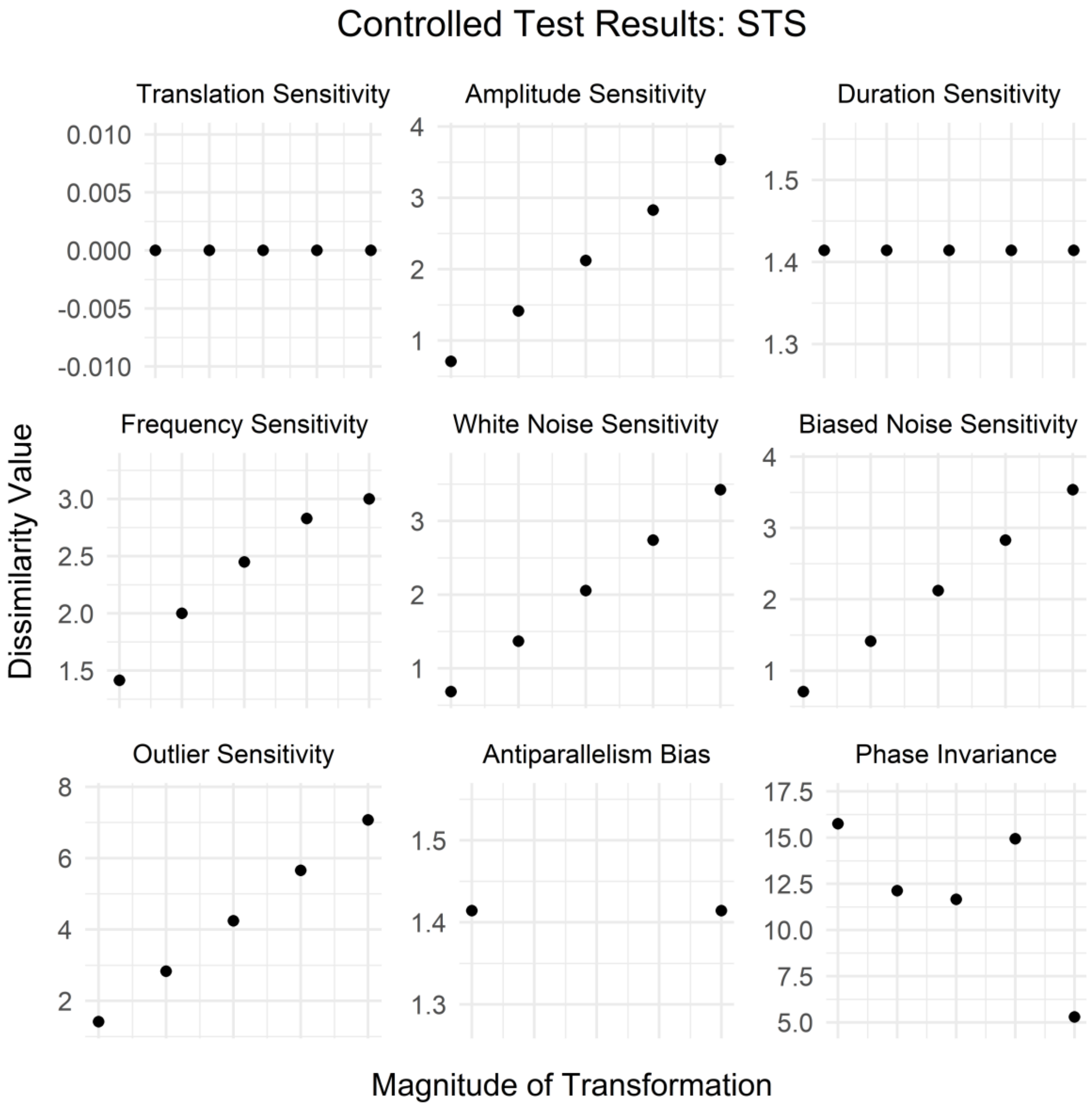
Controlled Test Results: STS
Translation Sensitivity
Amplitude Sensitivity
Duration Sensitivity
Frequency Sensitivity
White Noise Sensitivity
Biased Noise Sensitivity
Outlier Sensitivity
Antiparallelism Bias
Phase Invariance
Dissimilarity Value
Magnitude of Transformation
0.010
0.005
0.000
-0.005
-0.010
1.5
1.4
1.3
3.0
2.5
2.0
17.5
15.0
12.5
10.0
7.5
5.0

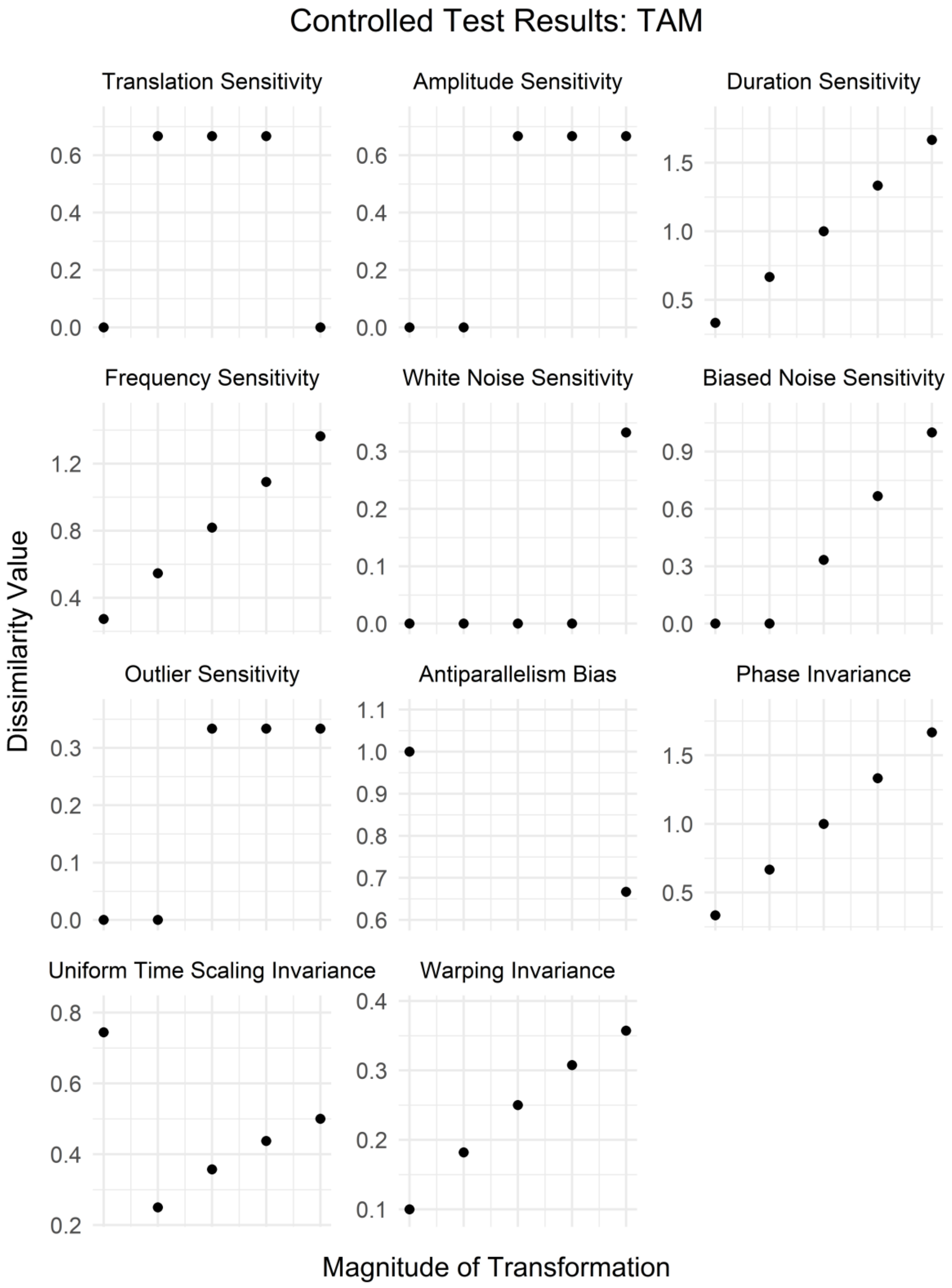
Controlled Test Results: TAM
Translation Sensitivity
Amplitude Sensitivity
Duration Sensitivity
Frequency Sensitivity
White Noise Sensitivity
Biased Noise Sensitivity
Outlier Sensitivity
Antiparallelism Bias
Phase Invariance
Uniform Time Scaling Invariance
Warping Invariance
Dissimilarity Value
Magnitude of Transformation

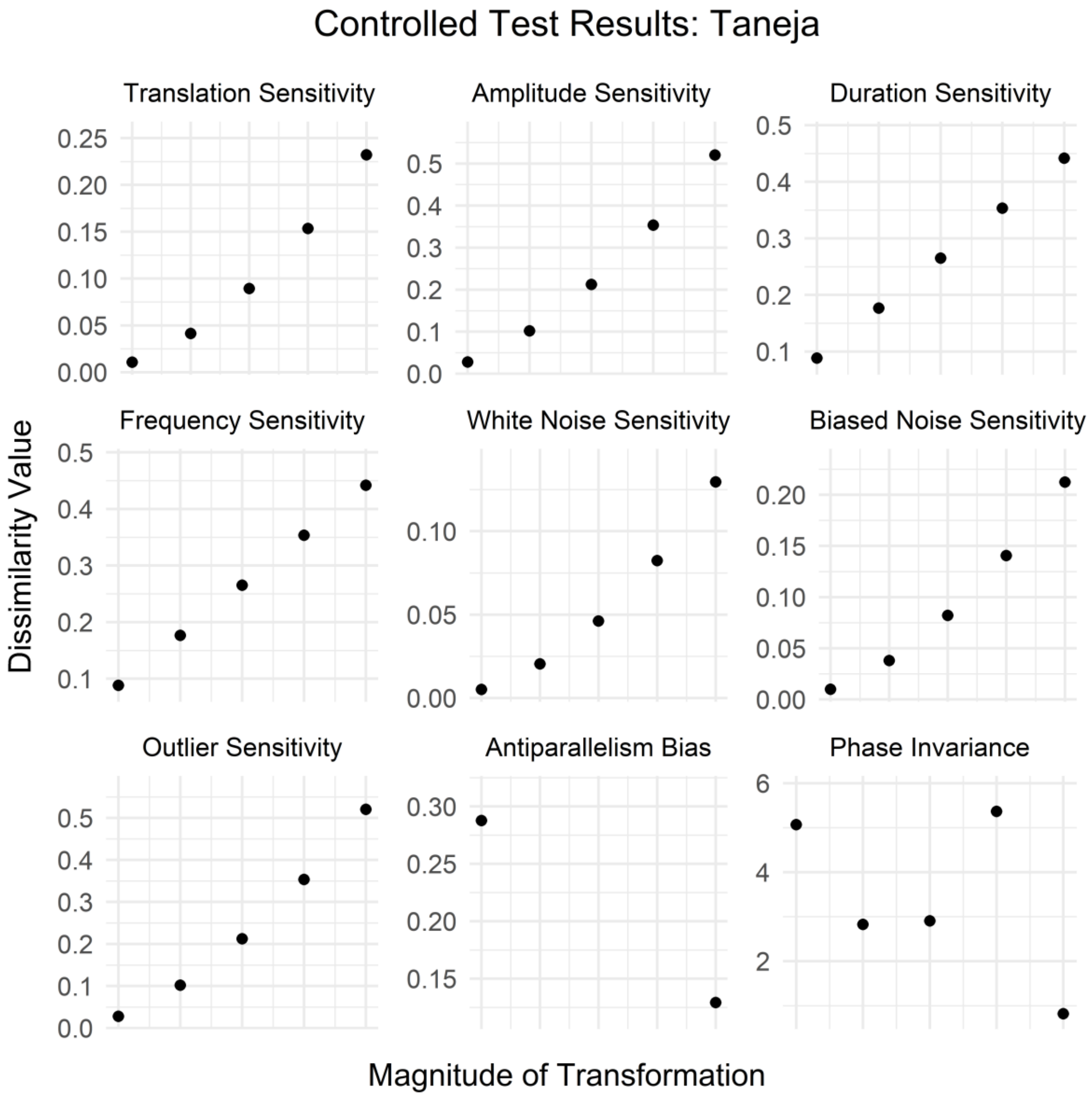
Controlled Test Results: Taneja
Translation Sensitivity
Amplitude Sensitivity
Duration Sensitivity
Frequency Sensitivity
White Noise Sensitivity
Biased Noise Sensitivity
Outlier Sensitivity
Antiparallelism Bias
Phase Invariance
Dissimilarity Value
Magnitude of Transformation

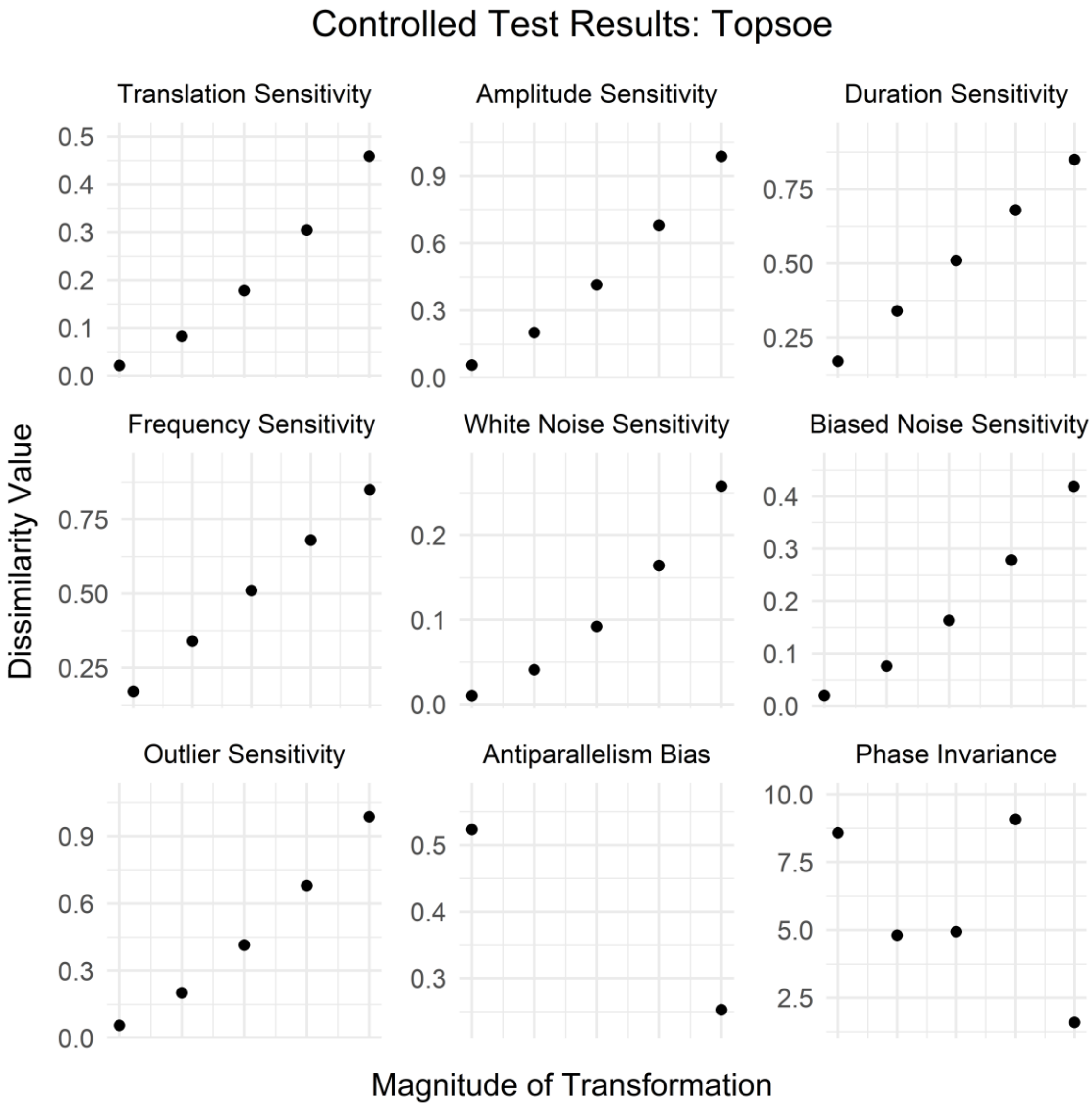
Controlled Test Results: Topsoe
Translation Sensitivity
Amplitude Sensitivity
Duration Sensitivity
Frequency Sensitivity
White Noise Sensitivity
Biased Noise Sensitivity
Outlier Sensitivity
Antiparallelism Bias
Phase Invariance
Dissimilarity Value
Magnitude of Transformation

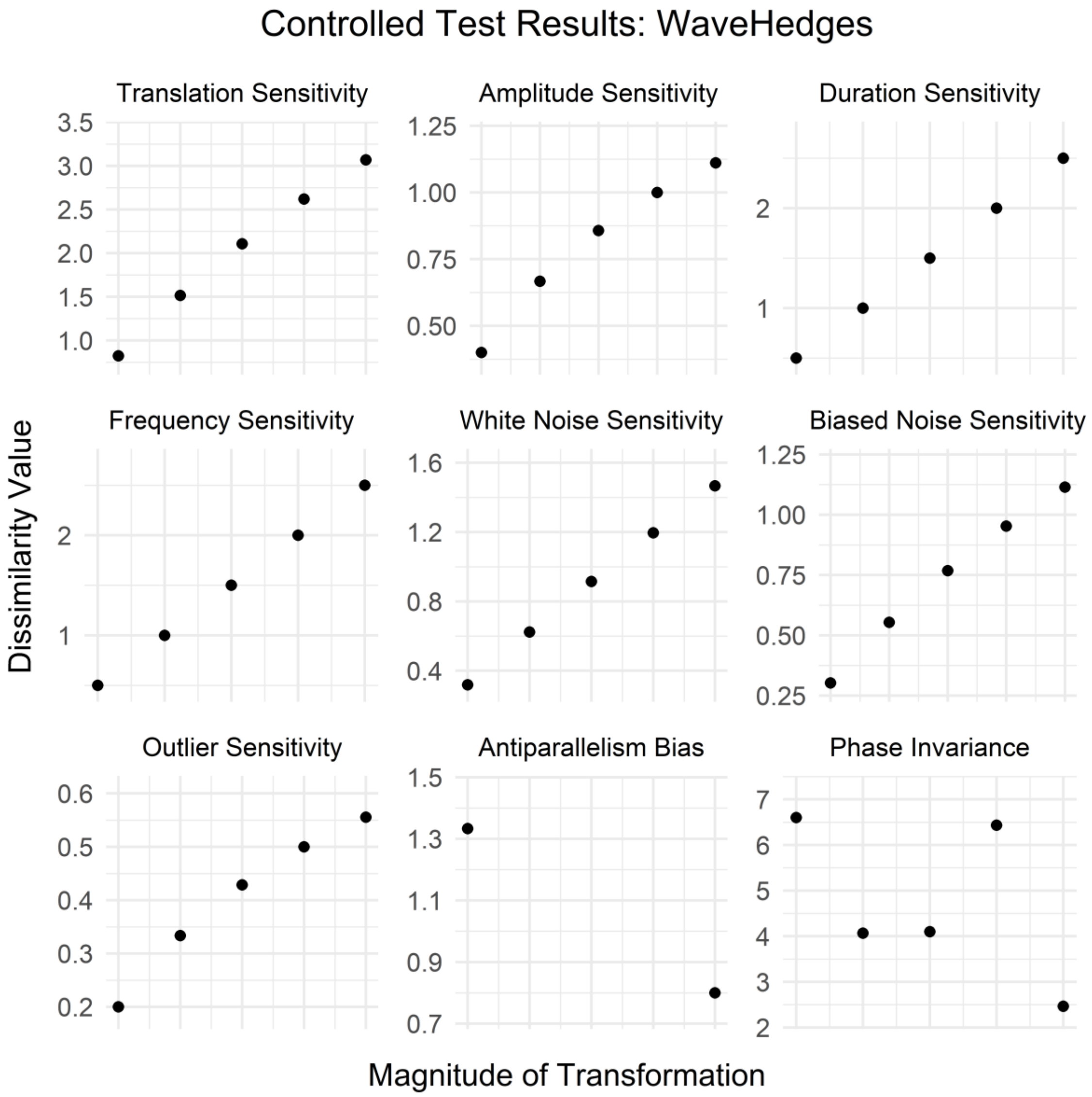
Controlled Test Results: WaveHedges
Translation Sensitivity
Amplitude Sensitivity
Duration Sensitivity
Frequency Sensitivity
White Noise Sensitivity
Biased Noise Sensitivity
Outlier Sensitivity
Antiparallelism Bias
Phase Invariance
Dissimilarity Value
Magnitude of Transformation

### 14. Tables of controlled test results for all distance measures

This section contains tables of controlled testing results for all 42 distance measures we tested. Each figure includes all time-based and values-based properties for which that distance measure gave results. Distance measures are presented in alphabetical order.

Controlled Test Results: ACF

| Test | Res.1 | Res.2 | Res.3 | Res.4 | Res.5 | Res.6 | Res.7 |
|---|---|---|---|---|---|---|---|
| Uniqueness | 0.0000 | | | | | | |
| Symmetry | 0.9611 | 0.9611 | | | | | |
| Translation Sensitivity | 0.0000 | 0.0000 | 0.0000 | 0.0000 | 0.0000 | | |
| Amplitude Sensitivity | 0.0941 | 0.2352 | 0.3529 | 0.4428 | 0.5114 | | |
| Duration Sensitivity | 0.4355 | 0.6796 | 0.6374 | 0.4965 | 0.4381 | | |
| Frequency Sensitivity | 0.5277 | 0.8189 | 1.0761 | 1.4041 | 1.8403 | | |
| White Noise Sensitivity | 0.0393 | 0.0902 | 0.1510 | 0.2196 | 0.2939 | | |
| Biased Noise Sensitivity | 0.0485 | 0.1042 | 0.1651 | 0.2294 | 0.2953 | | |
| Outlier Sensitivity | 0.1320 | 0.2021 | 0.2508 | 0.2910 | 0.3254 | | |
| Antiparallelism Bias | 0.0000 | 0.0000 | | | | | |
| Phase Invariance | 0.3938 | 0.7610 | 1.0553 | 0.3934 | 0.2057 | | |
| Uniform Time Scaling Invariance | | | | | | | |
| Warping Invariance | | | | | | | |
| Non-positive Value Handling | 0.6900 | 0.6900 | 0.8552 | | | | |
| Non-negativity | 1.0000 | | | | | | |
| Triangle Inequality | 1.0000 | | | | | | |
| Relative Sensitivity Ranges | 0.0000 | 1.0946 | 0.6401 | 3.4429 | 0.6678 | 0.6473 | 0.5073 |

Controlled Test Results: Additive

| Test | Res.1 | Res.2 | Res.3 | Res.4 | Res.5 | Res.6 | Res.7 |
|---|---|---|---|---|---|---|---|
| Uniqueness | 0.00 | | | | | | |
| Symmetry | 42.74 | 42.74 | | | | | |
| Translation Sensitivity | 0.17 | 0.66 | 1.44 | 2.49 | 3.78 | | |
| Amplitude Sensitivity | 0.45 | 1.67 | 3.54 | 6.00 | 9.03 | | |
| Duration Sensitivity | 1.50 | 3.00 | 4.50 | 6.00 | 7.50 | | |
| Frequency Sensitivity | 1.50 | 3.00 | 4.50 | 6.00 | 7.50 | | |
| White Noise Sensitivity | 0.08 | 0.33 | 0.74 | 1.32 | 2.09 | | |
| Biased Noise Sensitivity | 0.16 | 0.61 | 1.33 | 2.29 | 3.48 | | |
| Outlier Sensitivity | 0.45 | 1.67 | 3.54 | 6.00 | 9.03 | | |
| Antiparallelism Bias | 5.33 | 2.13 | | | | | |
| Phase Invariance | 106.90 | 59.40 | 61.00 | 112.90 | 13.60 | | |
| Uniform Time Scaling Invariance | | | | | | | |
| Warping Invariance | | | | | | | |
| Non-positive Value Handling | 3,999,998.00 | 0.00 | -4.50 | | | | |
| Non-negativity | 0.00 | | | | | | |
| Triangle Inequality | 0.00 | | | | | | |
| Relative Sensitivity Ranges | 0.66 | 1.58 | 1.10 | 1.10 | 0.37 | 0.61 | 1.58 |

Controlled Test Results: Piccolo

| Test | Res.1 | Res.2 | Res.3 | Res.4 | Res.5 | Res.6 | Res.7 |
|---|---|---|---|---|---|---|---|
| Uniqueness | 0.0000 | | | | | | |
| Symmetry | 0.5655 | 0.5655 | | | | | |
| Translation Sensitivity | 0.0000 | 0.0000 | 0.0000 | 0.0000 | 0.0000 | | |
| Amplitude Sensitivity | 0.0571 | 0.1382 | 0.2019 | 0.2484 | 0.2827 | | |
| Duration Sensitivity | 0.1310 | 0.1667 | 0.1500 | 0.0833 | 0.0595 | | |
| Frequency Sensitivity | 0.1182 | 0.5564 | 0.5511 | 0.6753 | 0.8182 | | |
| White Noise Sensitivity | 0.0680 | 0.1302 | 0.1824 | 0.2246 | 0.2586 | | |
| Biased Noise Sensitivity | 0.0721 | 0.1312 | 0.1782 | 0.2155 | 0.2459 | | |
| Outlier Sensitivity | 0.0523 | 0.0973 | 0.1310 | 0.1555 | 0.1734 | | |
| Antiparallelism Bias | 0.0000 | 0.0000 | | | | | |
| Phase Invariance | 0.0000 | 0.0570 | 0.2468 | 0.0570 | 0.1139 | | |
| Uniform Time Scaling Invariance | 0.3241 | 0.6077 | 0.6239 | 0.6389 | 0.6461 | | |
| Warping Invariance | 0.6642 | 0.8488 | 0.8799 | 0.4125 | 0.4522 | | |
| Non-positive Value Handling | 0.9883 | 0.9883 | 1.1271 | | | | |
| Non-negativity | 1.0000 | | | | | | |
| Triangle Inequality | 1.0000 | | | | | | |
| Relative Sensitivity Ranges | 0.0000 | 1.0399 | 0.4940 | 3.2275 | 0.8785 | 0.8014 | 0.5587 |

Controlled Test Results: AVG

| Test | Res.1 | Res.2 | Res.3 | Res.4 | Res.5 | Res.6 | Res.7 |
|---|---|---|---|---|---|---|---|
| Uniqueness | 0.000 | | | | | | |
| Symmetry | 7.900 | 7.900 | | | | | |
| Translation Sensitivity | 0.550 | 1.100 | 1.650 | 2.200 | 2.750 | | |
| Amplitude Sensitivity | 0.750 | 1.500 | 2.250 | 3.000 | 3.750 | | |
| Duration Sensitivity | 1.000 | 1.500 | 2.000 | 2.500 | 3.000 | | |
| Frequency Sensitivity | 1.000 | 1.500 | 2.000 | 2.500 | 3.000 | | |
| White Noise Sensitivity | 0.562 | 1.125 | 1.688 | 2.250 | 2.812 | | |
| Biased Noise Sensitivity | 0.625 | 1.250 | 1.875 | 2.500 | 3.125 | | |
| Outlier Sensitivity | 1.000 | 2.000 | 3.000 | 4.000 | 5.000 | | |
| Antiparallelism Bias | 1.500 | 1.500 | | | | | |
| Phase Invariance | 16.500 | 11.500 | 11.500 | 16.500 | 7.000 | | |
| Uniform Time Scaling Invariance | | | | | | | |
| Warping Invariance | | | | | | | |
| Non-positive Value Handling | 2.000 | 2.000 | 3.000 | | | | |
| Non-negativity | 1.000 | | | | | | |
| Triangle Inequality | 1.000 | | | | | | |
| Relative Sensitivity Ranges | 0.858 | 1.170 | 0.780 | 0.780 | 0.877 | 0.975 | 1.560 |

Controlled Test Results: Canb

| Test | Res.1 | Res.2 | Res.3 | Res.4 | Res.5 | Res.6 | Res.7 |
|---|---|---|---|---|---|---|---|
| Uniqueness | 0.0000 | | | | | | |
| Symmetry | 4.6030 | 4.6030 | | | | | |
| Translation Sensitivity | 0.4297 | 0.8225 | 1.1830 | 1.5152 | 1.8222 | | |
| Amplitude Sensitivity | 0.2222 | 0.4000 | 0.5455 | 0.6667 | 0.7692 | | |
| Duration Sensitivity | 0.3333 | 0.6667 | 1.0000 | 1.3333 | 1.6667 | | |
| Frequency Sensitivity | 0.3333 | 0.6667 | 1.0000 | 1.3333 | 1.6667 | | |
| White Noise Sensitivity | 0.1633 | 0.3265 | 0.4903 | 0.6553 | 0.8222 | | |
| Biased Noise Sensitivity | 0.1585 | 0.3022 | 0.4334 | 0.5538 | 0.6649 | | |
| Outlier Sensitivity | 0.1111 | 0.2000 | 0.2727 | 0.3333 | 0.3846 | | |
| Antiparallelism Bias | 1.0000 | 0.5000 | | | | | |
| Phase Invariance | 5.1143 | 3.0032 | 3.0528 | 5.0810 | 1.5667 | | |
| Uniform Time Scaling Invariance | | | | | | | |
| Warping Invariance | | | | | | | |
| Non-positive Value Handling | 1.0000 | 1.0000 | 3.0000 | | | | |
| Non-negativity | 0.0000 | | | | | | |
| Triangle Inequality | 0.0000 | | | | | | |
| Relative Sensitivity Ranges | 1.6125 | 0.6334 | 1.5440 | 1.5440 | 0.7630 | 0.5864 | 0.3167 |

Controlled Test Results: CDM

| Test | Res.1 | Res.2 | Res.3 | Res.4 | Res.5 | Res.6 | Res.7 |
|---|---|---|---|---|---|---|---|
| Uniqueness | 0.5588 | | | | | | |
| Symmetry | 0.7073 | 0.6829 | | | | | |
| Translation Sensitivity | 0.7222 | 0.7222 | 0.7222 | 0.7222 | 0.7222 | | |
| Amplitude Sensitivity | 0.6667 | 0.6579 | 0.6750 | 0.6579 | 0.6750 | | |
| Duration Sensitivity | 0.5588 | 0.5588 | 0.5882 | 0.5882 | 0.6250 | | |
| Frequency Sensitivity | 0.6333 | 0.6333 | 0.6333 | 0.6552 | 0.5714 | | |
| White Noise Sensitivity | 0.8793 | 0.8571 | 0.8621 | 0.8367 | 0.8621 | | |
| Biased Noise Sensitivity | 0.7308 | 0.7255 | 0.7308 | 0.7609 | 0.7885 | | |
| Outlier Sensitivity | 0.6591 | 0.6591 | 0.6591 | 0.6591 | 0.6591 | | |
| Antiparallelism Bias | 0.6216 | 0.6216 | | | | | |
| Phase Invariance | 0.5532 | 0.5417 | 0.5870 | 0.5833 | 0.5833 | | |
| Uniform Time Scaling Invariance | 0.7727 | 0.7639 | 0.7763 | 0.7975 | 0.7538 | | |
| Warping Invariance | 0.5556 | 0.5556 | 0.5556 | 0.5676 | 0.5676 | | |
| Non-positive Value Handling | 0.6923 | 0.6571 | 0.6571 | | | | |
| Non-negativity | 1.0000 | | | | | | |
| Triangle Inequality | 1.0000 | | | | | | |
| Relative Sensitivity Ranges | 0.0000 | 0.3138 | 1.2139 | 1.5362 | 0.7810 | 1.1551 | 0.0000 |

Controlled Test Results: CID

| Test | Res.1 | Res.2 | Res.3 | Res.4 | Res.5 | Res.6 | Res.7 |
|---|---|---|---|---|---|---|---|
| Uniqueness | 0.000 | | | | | | |
| Symmetry | 18.512 | 18.512 | | | | | |
| Translation Sensitivity | 0.316 | 0.632 | 0.949 | 1.265 | 1.581 | | |
| Amplitude Sensitivity | 0.901 | 2.236 | 4.039 | 6.325 | 9.100 | | |
| Duration Sensitivity | 1.000 | 1.414 | 1.732 | 2.000 | 2.236 | | |
| Frequency Sensitivity | 1.414 | 2.449 | 3.464 | 4.472 | 5.244 | | |
| White Noise Sensitivity | 0.373 | 0.796 | 1.284 | 1.846 | 2.492 | | |
| Biased Noise Sensitivity | 0.533 | 1.148 | 1.865 | 2.697 | 3.654 | | |
| Outlier Sensitivity | 1.254 | 3.117 | 5.669 | 8.944 | 12.956 | | |
| Antiparallelism Bias | 1.414 | 4.243 | | | | | |
| Phase Invariance | 9.592 | 7.820 | 8.052 | 10.650 | 5.272 | | |
| Uniform Time Scaling Invariance | | | | | | | |
| Warping Invariance | | | | | | | |
| Non-positive Value Handling | 3.464 | 3.464 | 7.036 | | | | |
| Non-negativity | 1.000 | | | | | | |
| Triangle Inequality | 0.000 | | | | | | |
| Relative Sensitivity Ranges | 0.281 | 1.824 | 0.275 | 0.852 | 0.471 | 0.694 | 2.603 |

Controlled Test Results: Clark

| Test | Res.1 | Res.2 | Res.3 | Res.4 | Res.5 | Res.6 | Res.7 |
|---|---|---|---|---|---|---|---|
| Uniqueness | 0.0000 | | | | | | |
| Symmetry | 1.5951 | 1.5951 | | | | | |
| Translation Sensitivity | 0.1390 | 0.2658 | 0.3819 | 0.4886 | 0.5871 | | |
| Amplitude Sensitivity | 0.1571 | 0.2828 | 0.3857 | 0.4714 | 0.5439 | | |
| Duration Sensitivity | 0.3333 | 0.4714 | 0.5774 | 0.6667 | 0.7454 | | |
| Frequency Sensitivity | 0.3333 | 0.4714 | 0.5774 | 0.6667 | 0.7454 | | |
| White Noise Sensitivity | 0.0629 | 0.1256 | 0.1887 | 0.2525 | 0.3176 | | |
| Biased Noise Sensitivity | 0.0881 | 0.1670 | 0.2383 | 0.3030 | 0.3622 | | |
| Outlier Sensitivity | 0.1111 | 0.2000 | 0.2727 | 0.3333 | 0.3846 | | |
| Antiparallelism Bias | 0.7071 | 0.3536 | | | | | |
| Phase Invariance | 1.6974 | 1.2317 | 1.2546 | 1.7263 | 0.6536 | | |
| Uniform Time Scaling Invariance | | | | | | | |
| Warping Invariance | | | | | | | |
| Non-positive Value Handling | 1.0000 | 1.0000 | 3.0000 | | | | |
| Non-negativity | 1.0000 | | | | | | |
| Triangle Inequality | 0.0000 | | | | | | |
| Relative Sensitivity Ranges | 1.2743 | 1.1001 | 1.1718 | 1.1718 | 0.7244 | 0.7797 | 0.7779 |

Controlled Test Results: Cort

| Test | Res.1 | Res.2 | Res.3 | Res.4 | Res.5 | Res.6 | Res.7 |
|---|---|---|---|---|---|---|---|
| Uniqueness | 0.000 | | | | | | |
| Symmetry | 2.566 | 2.566 | | | | | |
| Translation Sensitivity | 0.075 | 0.151 | 0.226 | 0.302 | 0.377 | | |
| Amplitude Sensitivity | 0.174 | 0.369 | 0.582 | 0.810 | 1.048 | | |
| Duration Sensitivity | 0.538 | 0.761 | 0.932 | 1.076 | 1.203 | | |
| Frequency Sensitivity | 0.391 | 0.678 | 0.932 | 1.161 | 1.336 | | |
| White Noise Sensitivity | 0.086 | 0.178 | 0.281 | 0.398 | 0.527 | | |
| Biased Noise Sensitivity | 0.121 | 0.251 | 0.396 | 0.558 | 0.736 | | |
| Outlier Sensitivity | 0.252 | 0.551 | 0.897 | 1.277 | 1.681 | | |
| Antiparallelism Bias | 2.491 | 0.337 | | | | | |
| Phase Invariance | 14.955 | 6.919 | 6.215 | 15.084 | 1.561 | | |
| Uniform Time Scaling Invariance | | | | | | | |
| Warping Invariance | | | | | | | |
| Non-positive Value Handling | 0.959 | 0.959 | 1.793 | | | | |
| Non-negativity | 1.000 | | | | | | |
| Triangle Inequality | 0.000 | | | | | | |
| Relative Sensitivity Ranges | 0.400 | 1.160 | 0.883 | 1.255 | 0.587 | 0.816 | 1.898 |

Controlled Test Results: Czek

| Test | Res.1 | Res.2 | Res.3 | Res.4 | Res.5 | Res.6 | Res.7 |
|---|---|---|---|---|---|---|---|
| Uniqueness | 0.0000 | | | | | | |
| Symmetry | 0.4103 | 0.4103 | | | | | |
| Translation Sensitivity | 0.0400 | 0.0769 | 0.1111 | 0.1429 | 0.1724 | | |
| Amplitude Sensitivity | 0.0345 | 0.0667 | 0.0968 | 0.1250 | 0.1515 | | |
| Duration Sensitivity | 0.0400 | 0.0769 | 0.1111 | 0.1429 | 0.1724 | | |
| Frequency Sensitivity | 0.0370 | 0.0714 | 0.1034 | 0.1333 | 0.1613 | | |
| White Noise Sensitivity | 0.0156 | 0.0312 | 0.0469 | 0.0625 | 0.0781 | | |
| Biased Noise Sensitivity | 0.0154 | 0.0303 | 0.0448 | 0.0588 | 0.0725 | | |
| Outlier Sensitivity | 0.0204 | 0.0400 | 0.0588 | 0.0769 | 0.0943 | | |
| Antiparallelism Bias | 0.1000 | 0.0833 | | | | | |
| Phase Invariance | 0.5000 | 0.3214 | 0.3214 | 0.5000 | 0.2143 | | |
| Uniform Time Scaling Invariance | | | | | | | |
| Warping Invariance | | | | | | | |
| Non-positive Value Handling | 0.0769 | 0.0769 | 0.1200 | | | | |
| Non-negativity | 0.0000 | | | | | | |
| Triangle Inequality | 0.0000 | | | | | | |
| Relative Sensitivity Ranges | 1.3248 | 1.1710 | 1.3248 | 1.2432 | 0.6253 | 0.5711 | 0.7397 |

Controlled Test Results: Dice

| Test | Res.1 | Res.2 | Res.3 | Res.4 | Res.5 | Res.6 | Res.7 |
|---|---|---|---|---|---|---|---|
| Uniqueness | 0.0000 | | | | | | |
| Symmetry | 0.3077 | 0.3077 | | | | | |
| Translation Sensitivity | 0.0029 | 0.0108 | 0.0224 | 0.0370 | 0.0538 | | |
| Amplitude Sensitivity | 0.0103 | 0.0370 | 0.0744 | 0.1176 | 0.1634 | | |
| Duration Sensitivity | 0.0286 | 0.0526 | 0.0732 | 0.0909 | 0.1064 | | |
| Frequency Sensitivity | 0.0303 | 0.0556 | 0.0769 | 0.0952 | 0.1111 | | |
| White Noise Sensitivity | 0.0005 | 0.0018 | 0.0041 | 0.0072 | 0.0113 | | |
| Biased Noise Sensitivity | 0.0009 | 0.0035 | 0.0075 | 0.0130 | 0.0196 | | |
| Outlier Sensitivity | 0.0069 | 0.0256 | 0.0533 | 0.0870 | 0.1244 | | |
| Antiparallelism Bias | 0.0952 | 0.0606 | | | | | |
| Phase Invariance | 0.4182 | 0.2545 | 0.2545 | 0.4545 | 0.1091 | | |
| Uniform Time Scaling Invariance | | | | | | | |
| Warping Invariance | | | | | | | |
| Non-positive Value Handling | 0.0909 | 0.0909 | 0.2000 | | | | |
| Non-negativity | 1.0000 | | | | | | |
| Triangle Inequality | 0.0000 | | | | | | |
| Relative Sensitivity Ranges | 0.6987 | 2.1029 | 1.0688 | 1.1100 | 0.1483 | 0.2575 | 1.6138 |

Controlled Test Results: Diverge

| Test | Res.1 | Res.2 | Res.3 | Res.4 | Res.5 | Res.6 | Res.7 |
|---|---|---|---|---|---|---|---|
| Uniqueness | 0.000 | | | | | | |
| Symmetry | 5.089 | 5.089 | | | | | |
| Translation Sensitivity | 0.039 | 0.141 | 0.292 | 0.478 | 0.689 | | |
| Amplitude Sensitivity | 0.049 | 0.160 | 0.298 | 0.444 | 0.592 | | |
| Duration Sensitivity | 0.222 | 0.444 | 0.667 | 0.889 | 1.111 | | |
| Frequency Sensitivity | 0.222 | 0.444 | 0.667 | 0.889 | 1.111 | | |
| White Noise Sensitivity | 0.008 | 0.032 | 0.071 | 0.128 | 0.202 | | |
| Biased Noise Sensitivity | 0.016 | 0.056 | 0.114 | 0.184 | 0.262 | | |
| Outlier Sensitivity | 0.025 | 0.080 | 0.149 | 0.222 | 0.296 | | |
| Antiparallelism Bias | 1.000 | 0.250 | | | | | |
| Phase Invariance | 5.763 | 3.034 | 3.148 | 5.960 | 0.854 | | |
| Uniform Time Scaling Invariance | | | | | | | |
| Warping Invariance | | | | | | | |
| Non-positive Value Handling | 2.000 | 2.000 | 18.000 | | | | |
| Non-negativity | 1.000 | | | | | | |
| Triangle Inequality | 0.000 | | | | | | |
| Relative Sensitivity Ranges | 1.237 | 1.031 | 1.690 | 1.690 | 0.368 | 0.469 | 0.515 |

Controlled Test Results: DTW

| Test | Res.1 | Res.2 | Res.3 | Res.4 | Res.5 | Res.6 | Res.7 |
|---|---|---|---|---|---|---|---|
| Uniqueness | 0.000 | | | | | | |
| Symmetry | 13.800 | 13.800 | | | | | |
| Translation Sensitivity | 1.900 | 3.800 | 5.700 | 7.600 | 9.500 | | |
| Amplitude Sensitivity | 2.000 | 4.000 | 5.000 | 6.000 | 7.000 | | |
| Duration Sensitivity | 0.000 | 0.000 | 0.000 | 0.000 | 0.000 | | |
| Frequency Sensitivity | 1.000 | 2.000 | 3.000 | 4.000 | 5.000 | | |
| White Noise Sensitivity | 2.000 | 4.000 | 6.000 | 8.000 | 9.750 | | |
| Biased Noise Sensitivity | 2.000 | 4.000 | 5.750 | 6.000 | 7.000 | | |
| Outlier Sensitivity | 2.000 | 4.000 | 5.000 | 6.000 | 7.000 | | |
| Antiparallelism Bias | 2.000 | 3.000 | | | | | |
| Phase Invariance | 8.000 | 8.000 | 15.000 | 22.000 | 18.000 | | |
| Uniform Time Scaling Invariance | 6.490 | 6.740 | 7.000 | 7.870 | 6.000 | | |
| Warping Invariance | 0.000 | 0.000 | 0.000 | 0.000 | 0.000 | | |
| Non-positive Value Handling | 3.000 | 3.000 | 4.000 | | | | |
| Non-negativity | 1.000 | | | | | | |
| Triangle Inequality | 0.000 | | | | | | |
| Relative Sensitivity Ranges | 1.328 | 0.873 | 0.000 | 0.699 | 1.354 | 0.873 | 0.873 |

Controlled Test Results: EDR

| Test | Res.1 | Res.2 | Res.3 | Res.4 | Res.5 | Res.6 | Res.7 |
|---|---|---|---|---|---|---|---|
| Uniqueness | 0.000 | | | | | | |
| Symmetry | 10.000 | 10.000 | | | | | |
| Translation Sensitivity | 10.000 | 10.000 | 10.000 | 10.000 | 10.000 | | |
| Amplitude Sensitivity | 2.000 | 2.000 | 2.000 | 2.000 | 2.000 | | |
| Duration Sensitivity | 1.000 | 2.000 | 3.000 | 4.000 | 5.000 | | |
| Frequency Sensitivity | 1.000 | 2.000 | 3.000 | 4.000 | 5.000 | | |
| White Noise Sensitivity | 8.000 | 8.000 | 8.000 | 8.000 | 8.000 | | |
| Biased Noise Sensitivity | 4.000 | 4.000 | 4.000 | 4.000 | 4.000 | | |
| Outlier Sensitivity | 1.000 | 1.000 | 1.000 | 1.000 | 1.000 | | |
| Antiparallelism Bias | 2.000 | 2.000 | | | | | |
| Phase Invariance | 2.000 | 4.000 | 6.000 | 7.000 | 6.000 | | |
| Uniform Time Scaling Invariance | 8.000 | 9.000 | 12.000 | 13.000 | 9.000 | | |
| Warping Invariance | 1.000 | 2.000 | 3.000 | 4.000 | 5.000 | | |
| Non-positive Value Handling | 1.000 | 1.000 | 1.000 | | | | |
| Non-negativity | 1.000 | | | | | | |
| Triangle Inequality | 0.000 | | | | | | |
| Relative Sensitivity Ranges | 0.000 | 0.000 | 1.000 | 1.000 | 0.000 | 0.000 | 0.000 |

Controlled Test Results: ERP

| Test | Res.1 | Res.2 | Res.3 | Res.4 | Res.5 | Res.6 | Res.7 |
|---|---|---|---|---|---|---|---|
| Uniqueness | 0.000 | | | | | | |
| Symmetry | 11.200 | 11.200 | | | | | |
| Translation Sensitivity | 1.000 | 2.000 | 3.000 | 4.000 | 5.000 | | |
| Amplitude Sensitivity | 1.000 | 2.000 | 3.000 | 4.000 | 5.000 | | |
| Duration Sensitivity | 1.000 | 2.000 | 3.000 | 4.000 | 5.000 | | |
| Frequency Sensitivity | 1.000 | 2.000 | 3.000 | 4.000 | 5.000 | | |
| White Noise Sensitivity | 1.000 | 2.000 | 3.000 | 4.000 | 5.000 | | |
| Biased Noise Sensitivity | 1.000 | 2.000 | 3.000 | 4.000 | 5.000 | | |
| Outlier Sensitivity | 1.000 | 2.000 | 3.000 | 4.000 | 5.000 | | |
| Antiparallelism Bias | 2.000 | 2.000 | | | | | |
| Phase Invariance | 10.000 | 12.000 | 12.000 | 14.000 | 12.000 | | |
| Uniform Time Scaling Invariance | 6.170 | 12.860 | 18.625 | 25.340 | 31.000 | | |
| Warping Invariance | 1.000 | 2.000 | 3.000 | 4.000 | 5.000 | | |
| Non-positive Value Handling | 2.000 | 2.000 | 3.000 | | | | |
| Non-negativity | 1.000 | | | | | | |
| Triangle Inequality | 1.000 | | | | | | |
| Relative Sensitivity Ranges | 1.000 | 1.000 | 1.000 | 1.000 | 1.000 | 1.000 | 1.000 |

Controlled Test Results: Euclidean

| Test | Res.1 | Res.2 | Res.3 | Res.4 | Res.5 | Res.6 | Res.7 |
|---|---|---|---|---|---|---|---|
| Uniqueness | 0.000 | | | | | | |
| Symmetry | 4.541 | 4.541 | | | | | |
| Translation Sensitivity | 0.316 | 0.632 | 0.949 | 1.265 | 1.581 | | |
| Amplitude Sensitivity | 0.707 | 1.414 | 2.121 | 2.828 | 3.536 | | |
| Duration Sensitivity | 1.000 | 1.414 | 1.732 | 2.000 | 2.236 | | |
| Frequency Sensitivity | 1.000 | 1.414 | 1.732 | 2.000 | 2.236 | | |
| White Noise Sensitivity | 0.354 | 0.707 | 1.061 | 1.414 | 1.768 | | |
| Biased Noise Sensitivity | 0.500 | 1.000 | 1.500 | 2.000 | 2.500 | | |
| Outlier Sensitivity | 1.000 | 2.000 | 3.000 | 4.000 | 5.000 | | |
| Antiparallelism Bias | 1.414 | 1.414 | | | | | |
| Phase Invariance | 9.592 | 7.483 | 7.483 | 10.000 | 4.899 | | |
| Uniform Time Scaling Invariance | | | | | | | |
| Warping Invariance | | | | | | | |
| Non-positive Value Handling | 2.000 | 2.000 | 3.000 | | | | |
| Non-negativity | 1.000 | | | | | | |
| Triangle Inequality | 1.000 | | | | | | |
| Relative Sensitivity Ranges | 0.633 | 1.416 | 0.619 | 0.619 | 0.708 | 1.001 | 2.003 |

Controlled Test Results: Fourier

| Test | Res.1 | Res.2 | Res.3 | Res.4 | Res.5 | Res.6 | Res.7 |
|---|---|---|---|---|---|---|---|
| Uniqueness | 0.000 | | | | | | |
| Symmetry | 11.381 | 11.381 | | | | | |
| Translation Sensitivity | 1.000 | 2.000 | 3.000 | 4.000 | 5.000 | | |
| Amplitude Sensitivity | 1.732 | 3.464 | 5.196 | 6.928 | 8.660 | | |
| Duration Sensitivity | 2.449 | 3.464 | 4.472 | 5.292 | 6.164 | | |
| Frequency Sensitivity | 2.646 | 4.000 | 5.196 | 6.325 | 7.416 | | |
| White Noise Sensitivity | 1.061 | 2.121 | 3.182 | 4.243 | 5.303 | | |
| Biased Noise Sensitivity | 1.500 | 3.000 | 4.500 | 6.000 | 7.500 | | |
| Outlier Sensitivity | 2.449 | 4.899 | 7.348 | 9.798 | 12.247 | | |
| Antiparallelism Bias | 3.464 | 3.464 | | | | | |
| Phase Invariance | 21.633 | 16.733 | 16.971 | 22.361 | 11.314 | | |
| Uniform Time Scaling Invariance | | | | | | | |
| Warping Invariance | | | | | | | |
| Non-positive Value Handling | 4.899 | 4.899 | 7.348 | | | | |
| Non-negativity | 1.000 | | | | | | |
| Triangle Inequality | 1.000 | | | | | | |
| Relative Sensitivity Ranges | 0.710 | 1.229 | 0.659 | 0.846 | 0.753 | 1.065 | 1.738 |

Controlled Test Results: Gower

| Test | Res.1 | Res.2 | Res.3 | Res.4 | Res.5 | Res.6 | Res.7 |
|---|---|---|---|---|---|---|---|
| Uniqueness | 0.0000 | | | | | | |
| Symmetry | 1.2800 | 1.2800 | | | | | |
| Translation Sensitivity | 0.1000 | 0.2000 | 0.3000 | 0.4000 | 0.5000 | | |
| Amplitude Sensitivity | 0.1000 | 0.2000 | 0.3000 | 0.4000 | 0.5000 | | |
| Duration Sensitivity | 0.1000 | 0.2000 | 0.3000 | 0.4000 | 0.5000 | | |
| Frequency Sensitivity | 0.0833 | 0.1667 | 0.2500 | 0.3333 | 0.4167 | | |
| White Noise Sensitivity | 0.1000 | 0.2000 | 0.3000 | 0.4000 | 0.5000 | | |
| Biased Noise Sensitivity | 0.1000 | 0.2000 | 0.3000 | 0.4000 | 0.5000 | | |
| Outlier Sensitivity | 0.1000 | 0.2000 | 0.3000 | 0.4000 | 0.5000 | | |
| Antiparallelism Bias | 0.2000 | 0.2000 | | | | | |
| Phase Invariance | 2.8000 | 1.8000 | 1.8000 | 2.8000 | 1.2000 | | |
| Uniform Time Scaling Invariance | | | | | | | |
| Warping Invariance | | | | | | | |
| Non-positive Value Handling | 0.2000 | 0.2000 | 0.3000 | | | | |
| Non-negativity | 1.0000 | | | | | | |
| Triangle Inequality | 1.0000 | | | | | | |
| Relative Sensitivity Ranges | 1.0244 | 1.0244 | 1.0244 | 0.8537 | 1.0244 | 1.0244 | 1.0244 |

Controlled Test Results: Chebyshev

| Test | Res.1 | Res.2 | Res.3 | Res.4 | Res.5 | Res.6 | Res.7 |
|---|---|---|---|---|---|---|---|
| Uniqueness | 0.0000 | | | | | | |
| Symmetry | 3.0000 | 3.0000 | | | | | |
| Translation Sensitivity | 0.1000 | 0.2000 | 0.3000 | 0.4000 | 0.5000 | | |
| Amplitude Sensitivity | 0.5000 | 1.0000 | 1.5000 | 2.0000 | 2.5000 | | |
| Duration Sensitivity | 1.0000 | 1.0000 | 1.0000 | 1.0000 | 1.0000 | | |
| Frequency Sensitivity | 1.0000 | 1.0000 | 1.0000 | 1.0000 | 1.0000 | | |
| White Noise Sensitivity | 0.1250 | 0.2500 | 0.3750 | 0.5000 | 0.6250 | | |
| Biased Noise Sensitivity | 0.2500 | 0.5000 | 0.7500 | 1.0000 | 1.2500 | | |
| Outlier Sensitivity | 1.0000 | 2.0000 | 3.0000 | 4.0000 | 5.0000 | | |
| Antiparallelism Bias | 1.0000 | 1.0000 | | | | | |
| Phase Invariance | 5.0000 | 5.0000 | 5.0000 | 5.0000 | 2.0000 | | |
| Uniform Time Scaling Invariance | | | | | | | |
| Warping Invariance | | | | | | | |
| Non-positive Value Handling | 2.0000 | 2.0000 | 3.0000 | | | | |
| Non-negativity | 1.0000 | | | | | | |
| Triangle Inequality | 1.0000 | | | | | | |
| Relative Sensitivity Ranges | 0.2532 | 1.2658 | 0.0000 | 0.0000 | 0.3165 | 0.6329 | 2.5316 |

Controlled Test Results: IntPer

| Test | Res.1 | Res.2 | Res.3 | Res.4 | Res.5 | Res.6 | Res.7 |
|---|---|---|---|---|---|---|---|
| Uniqueness | 0.0000 | | | | | | |
| Symmetry | 0.8095 | 0.8095 | | | | | |
| Translation Sensitivity | 0.0000 | 0.0000 | 0.0000 | 0.0000 | 0.0000 | | |
| Amplitude Sensitivity | 0.0266 | 0.0676 | 0.1029 | 0.1305 | 0.1520 | | |
| Duration Sensitivity | 0.5316 | 0.6016 | 0.6298 | 0.6331 | 0.7826 | | |
| Frequency Sensitivity | 0.9671 | 1.4235 | 1.5994 | 1.8517 | 1.9900 | | |
| White Noise Sensitivity | 0.1465 | 0.3306 | 0.5389 | 0.7586 | 0.9791 | | |
| Biased Noise Sensitivity | 0.1315 | 0.2976 | 0.4847 | 0.6810 | 0.8772 | | |
| Outlier Sensitivity | 0.1486 | 0.2231 | 0.2909 | 0.3391 | 0.3739 | | |
| Antiparallelism Bias | 0.0000 | 0.0000 | | | | | |
| Phase Invariance | 0.2718 | 0.5069 | 0.6876 | 0.8120 | 0.5408 | | |
| Uniform Time Scaling Invariance | | | | | | | |
| Warping Invariance | | | | | | | |
| Non-positive Value Handling | 1.1307 | 1.1307 | 1.2898 | | | | |
| Non-negativity | 1.0000 | | | | | | |
| Triangle Inequality | 1.0000 | | | | | | |
| Relative Sensitivity Ranges | 0.0000 | 0.2742 | 0.5485 | 2.2354 | 1.8197 | 1.6298 | 0.4924 |

Controlled Test Results: Jaccard

| Test | Res.1 | Res.2 | Res.3 | Res.4 | Res.5 | Res.6 | Res.7 |
|---|---|---|---|---|---|---|---|
| Uniqueness | 0.0000 | | | | | | |
| Symmetry | 0.4706 | 0.4706 | | | | | |
| Translation Sensitivity | 0.0058 | 0.0213 | 0.0439 | 0.0714 | 0.1020 | | |
| Amplitude Sensitivity | 0.0204 | 0.0714 | 0.1385 | 0.2105 | 0.2809 | | |
| Duration Sensitivity | 0.0556 | 0.1000 | 0.1364 | 0.1667 | 0.1923 | | |
| Frequency Sensitivity | 0.0588 | 0.1053 | 0.1429 | 0.1739 | 0.2000 | | |
| White Noise Sensitivity | 0.0009 | 0.0036 | 0.0082 | 0.0144 | 0.0223 | | |
| Biased Noise Sensitivity | 0.0018 | 0.0069 | 0.0150 | 0.0256 | 0.0385 | | |
| Outlier Sensitivity | 0.0137 | 0.0500 | 0.1011 | 0.1600 | 0.2212 | | |
| Antiparallelism Bias | 0.1739 | 0.1143 | | | | | |
| Phase Invariance | 0.5897 | 0.4058 | 0.4058 | 0.6250 | 0.1967 | | |
| Uniform Time Scaling Invariance | | | | | | | |
| Warping Invariance | | | | | | | |
| Non-positive Value Handling | 0.1667 | 0.1667 | 0.3333 | | | | |
| Non-negativity | 1.0000 | | | | | | |
| Triangle Inequality | 0.0000 | | | | | | |
| Relative Sensitivity Ranges | 0.7484 | 2.0254 | 1.0633 | 1.0977 | 0.1660 | 0.2856 | 1.6137 |

Controlled Test Results: Jeffreys

| Test | Res.1 | Res.2 | Res.3 | Res.4 | Res.5 | Res.6 | Res.7 |
|---|---|---|---|---|---|---|---|
| Uniqueness | 0.000 | | | | | | |
| Symmetry | 14.372 | 14.372 | | | | | |
| Translation Sensitivity | 0.086 | 0.330 | 0.714 | 1.223 | 1.845 | | |
| Amplitude Sensitivity | 0.223 | 0.811 | 1.679 | 2.773 | 4.055 | | |
| Duration Sensitivity | 0.693 | 1.386 | 2.079 | 2.773 | 3.466 | | |
| Frequency Sensitivity | 0.693 | 1.386 | 2.079 | 2.773 | 3.466 | | |
| White Noise Sensitivity | 0.041 | 0.163 | 0.368 | 0.658 | 1.034 | | |
| Biased Noise Sensitivity | 0.079 | 0.303 | 0.654 | 1.119 | 1.687 | | |
| Outlier Sensitivity | 0.223 | 0.811 | 1.679 | 2.773 | 4.055 | | |
| Antiparallelism Bias | 2.197 | 1.022 | | | | | |
| Phase Invariance | 37.437 | 20.901 | 21.476 | 39.634 | 6.438 | | |
| Uniform Time Scaling Invariance | | | | | | | |
| Warping Invariance | | | | | | | |
| Non-positive Value Handling | 29.017 | 23.026 | | | | | |
| Non-negativity | 1.000 | | | | | | |
| Triangle Inequality | 0.000 | | | | | | |
| Relative Sensitivity Ranges | 0.701 | 1.527 | 1.105 | 1.105 | 0.396 | 0.641 | 1.527 |

Controlled Test Results: Jensen

| Test | Res.1 | Res.2 | Res.3 | Res.4 | Res.5 | Res.6 | Res.7 |
|---|---|---|---|---|---|---|---|
| Uniqueness | 0.0000 | | | | | | |
| Symmetry | 1.6459 | 1.6459 | | | | | |
| Translation Sensitivity | 0.0107 | 0.0412 | 0.0890 | 0.1522 | 0.2292 | | |
| Amplitude Sensitivity | 0.0278 | 0.1007 | 0.2072 | 0.3398 | 0.4934 | | |
| Duration Sensitivity | 0.0849 | 0.1699 | 0.2548 | 0.3398 | 0.4247 | | |
| Frequency Sensitivity | 0.0849 | 0.1699 | 0.2548 | 0.3398 | 0.4247 | | |
| White Noise Sensitivity | 0.0051 | 0.0204 | 0.0460 | 0.0821 | 0.1288 | | |
| Biased Noise Sensitivity | 0.0099 | 0.0378 | 0.0815 | 0.1392 | 0.2093 | | |
| Outlier Sensitivity | 0.0278 | 0.1007 | 0.2072 | 0.3398 | 0.4934 | | |
| Antiparallelism Bias | 0.2616 | 0.1263 | | | | | |
| Phase Invariance | 4.2886 | 2.4001 | 2.4658 | 4.5413 | 0.7938 | | |
| Uniform Time Scaling Invariance | | | | | | | |
| Warping Invariance | | | | | | | |
| Non-positive Value Handling | 0.6931 | 0.6931 | | | | | |
| Non-negativity | 1.0000 | | | | | | |
| Triangle Inequality | 0.0000 | | | | | | |
| Relative Sensitivity Ranges | 0.7105 | 1.5142 | 1.1052 | 1.1052 | 0.4024 | 0.6485 | 1.5142 |

Controlled Test Results: Kulcz

| Test | Res.1 | Res.2 | Res.3 | Res.4 | Res.5 | Res.6 | Res.7 |
|---|---|---|---|---|---|---|---|
| Uniqueness | 0.0000 | | | | | | |
| Symmetry | 1.3913 | 1.3913 | | | | | |
| Translation Sensitivity | 0.0833 | 0.1667 | 0.2500 | 0.3333 | 0.4167 | | |
| Amplitude Sensitivity | 0.0714 | 0.1429 | 0.2143 | 0.2857 | 0.3571 | | |
| Duration Sensitivity | 0.0833 | 0.1667 | 0.2500 | 0.3333 | 0.4167 | | |
| Frequency Sensitivity | 0.0769 | 0.1538 | 0.2308 | 0.3077 | 0.3846 | | |
| White Noise Sensitivity | 0.0317 | 0.0645 | 0.0984 | 0.1333 | 0.1695 | | |
| Biased Noise Sensitivity | 0.0312 | 0.0625 | 0.0938 | 0.1250 | 0.1562 | | |
| Outlier Sensitivity | 0.0417 | 0.0833 | 0.1250 | 0.1667 | 0.2083 | | |
| Antiparallelism Bias | 0.2222 | 0.1818 | | | | | |
| Phase Invariance | 2.0000 | 0.9474 | 0.9474 | 2.0000 | 0.5455 | | |
| Uniform Time Scaling Invariance | | | | | | | |
| Warping Invariance | | | | | | | |
| Non-positive Value Handling | 0.1667 | 0.1667 | 0.2727 | | | | |
| Non-negativity | 0.0000 | | | | | | |
| Triangle Inequality | 0.0000 | | | | | | |
| Relative Sensitivity Ranges | 1.3811 | 1.1838 | 1.3811 | 1.2749 | 0.5707 | 0.5179 | 0.6905 |

Controlled Test Results: Kullback

| Test | Res.1 | Res.2 | Res.3 | Res.4 | Res.5 | Res.6 | Res.7 |
|---|---|---|---|---|---|---|---|
| Uniqueness | 0.000 | | | | | | |
| Symmetry | 14.449 | -0.077 | | | | | |
| Translation Sensitivity | 1.044 | 2.170 | 3.371 | 4.644 | 5.981 | | |
| Amplitude Sensitivity | 1.116 | 2.433 | 3.917 | 5.545 | 7.298 | | |
| Duration Sensitivity | 1.386 | 2.773 | 4.159 | 5.545 | 6.931 | | |
| Frequency Sensitivity | 1.386 | 2.773 | 4.159 | 5.545 | 6.931 | | |
| White Noise Sensitivity | 0.020 | 0.082 | 0.184 | 0.327 | 0.513 | | |
| Biased Noise Sensitivity | 1.040 | 2.156 | 3.342 | 4.591 | 5.900 | | |
| Outlier Sensitivity | 1.116 | 2.433 | 3.917 | 5.545 | 7.298 | | |
| Antiparallelism Bias | -1.099 | 2.554 | | | | | |
| Phase Invariance | 17.594 | 11.549 | 9.716 | 20.654 | 3.219 | | |
| Uniform Time Scaling Invariance | | | | | | | |
| Warping Invariance | | | | | | | |
| Non-positive Value Handling | -0.000 | 0.000 | | | | | |
| Non-negativity | 0.000 | | | | | | |
| Triangle Inequality | 0.000 | | | | | | |
| Relative Sensitivity Ranges | 1.024 | 1.283 | 1.150 | 1.150 | 0.102 | 1.008 | 1.283 |

Controlled Test Results: KumarJohnson

| Test | Res.1 | Res.2 | Res.3 | Res.4 | Res.5 | Res.6 | Res.7 |
|---|---|---|---|---|---|---|---|
| Uniqueness | 0.00 | | | | | | |
| Symmetry | 60.02 | 60.02 | | | | | |
| Translation Sensitivity | 0.17 | 0.67 | 1.45 | 2.52 | 3.85 | | |
| Amplitude Sensitivity | 0.45 | 1.70 | 3.68 | 6.36 | 9.78 | | |
| Duration Sensitivity | 1.59 | 3.18 | 4.77 | 6.36 | 7.95 | | |
| Frequency Sensitivity | 1.59 | 3.18 | 4.77 | 6.36 | 7.95 | | |
| White Noise Sensitivity | 0.08 | 0.33 | 0.74 | 1.33 | 2.11 | | |
| Biased Noise Sensitivity | 0.16 | 0.61 | 1.34 | 2.32 | 3.55 | | |
| Outlier Sensitivity | 0.45 | 1.70 | 3.68 | 6.36 | 9.78 | | |
| Antiparallelism Bias | 6.16 | 2.20 | | | | | |
| Phase Invariance | 140.11 | 77.92 | 79.91 | 147.54 | 14.17 | | |
| Uniform Time Scaling Invariance | | | | | | | |
| Warping Invariance | | | | | | | |
| Non-positive Value Handling | 2,828,427,124.74 | 1,600,000.00 | | | | | |
| Non-negativity | 1.00 | | | | | | |
| Triangle Inequality | 0.00 | | | | | | |
| Relative Sensitivity Ranges | 0.64 | 1.61 | 1.10 | 1.10 | 0.35 | 0.59 | 1.61 |

Controlled Test Results: KDiv

| Test | Res.1 | Res.2 | Res.3 | Res.4 | Res.5 | Res.6 | Res.7 |
|---|---|---|---|---|---|---|---|
| Uniqueness | 0.0000 | | | | | | |
| Symmetry | 5.1984 | -1.9065 | | | | | |
| Translation Sensitivity | 0.5106 | 1.0400 | 1.5853 | 2.1441 | 2.7147 | | |
| Amplitude Sensitivity | 0.5268 | 1.0939 | 1.6881 | 2.3015 | 2.9288 | | |
| Duration Sensitivity | 0.5754 | 1.1507 | 1.7261 | 2.3015 | 2.8768 | | |
| Frequency Sensitivity | 0.5754 | 1.1507 | 1.7261 | 2.3015 | 2.8768 | | |
| White Noise Sensitivity | 0.0051 | 0.0204 | 0.0461 | 0.0824 | 0.1298 | | |
| Biased Noise Sensitivity | 0.5097 | 1.0367 | 1.5780 | 2.1314 | 2.6954 | | |
| Outlier Sensitivity | 0.5268 | 1.0939 | 1.6881 | 2.3015 | 2.9288 | | |
| Antiparallelism Bias | -0.6931 | 1.1157 | | | | | |
| Phase Invariance | 4.5097 | 2.1852 | 2.6634 | 4.3847 | 0.7938 | | |
| Uniform Time Scaling Invariance | | | | | | | |
| Warping Invariance | | | | | | | |
| Non-positive Value Handling | -0.0000 | | | | | | |
| Non-negativity | 0.0000 | | | | | | |
| Triangle Inequality | 0.0000 | | | | | | |
| Relative Sensitivity Ranges | 1.1083 | 1.2078 | 1.1572 | 1.1572 | 0.0627 | 1.0990 | 1.2078 |

Controlled Test Results: Lorentz

| Test | Res.1 | Res.2 | Res.3 | Res.4 | Res.5 | Res.6 | Res.7 |
|---|---|---|---|---|---|---|---|
| Uniqueness | 0.000 | | | | | | |
| Symmetry | 7.927 | 7.927 | | | | | |
| Translation Sensitivity | 0.953 | 1.823 | 2.624 | 3.365 | 4.055 | | |
| Amplitude Sensitivity | 0.811 | 1.386 | 1.833 | 2.197 | 2.506 | | |
| Duration Sensitivity | 0.693 | 1.386 | 2.079 | 2.773 | 3.466 | | |
| Frequency Sensitivity | 0.693 | 1.386 | 2.079 | 2.773 | 3.466 | | |
| White Noise Sensitivity | 0.942 | 1.785 | 2.548 | 3.244 | 3.884 | | |
| Biased Noise Sensitivity | 0.893 | 1.622 | 2.238 | 2.773 | 3.244 | | |
| Outlier Sensitivity | 0.693 | 1.099 | 1.386 | 1.609 | 1.792 | | |
| Antiparallelism Bias | 1.386 | 1.386 | | | | | |
| Phase Invariance | 12.871 | 8.723 | 8.723 | 12.283 | 6.592 | | |
| Uniform Time Scaling Invariance | | | | | | | |
| Warping Invariance | | | | | | | |
| Non-positive Value Handling | 1.099 | 1.099 | 1.386 | | | | |
| Non-negativity | 1.000 | | | | | | |
| Triangle Inequality | 1.000 | | | | | | |
| Relative Sensitivity Ranges | 1.297 | 0.709 | 1.160 | 1.160 | 1.231 | 0.984 | 0.460 |

Controlled Test Results: Manhattan

| Test | Res.1 | Res.2 | Res.3 | Res.4 | Res.5 | Res.6 | Res.7 |
|---|---|---|---|---|---|---|---|
| Uniqueness | 0.000 | | | | | | |
| Symmetry | 12.800 | 12.800 | | | | | |
| Translation Sensitivity | 1.000 | 2.000 | 3.000 | 4.000 | 5.000 | | |
| Amplitude Sensitivity | 1.000 | 2.000 | 3.000 | 4.000 | 5.000 | | |
| Duration Sensitivity | 1.000 | 2.000 | 3.000 | 4.000 | 5.000 | | |
| Frequency Sensitivity | 1.000 | 2.000 | 3.000 | 4.000 | 5.000 | | |
| White Noise Sensitivity | 1.000 | 2.000 | 3.000 | 4.000 | 5.000 | | |
| Biased Noise Sensitivity | 1.000 | 2.000 | 3.000 | 4.000 | 5.000 | | |
| Outlier Sensitivity | 1.000 | 2.000 | 3.000 | 4.000 | 5.000 | | |
| Antiparallelism Bias | 2.000 | 2.000 | | | | | |
| Phase Invariance | 28.000 | 18.000 | 18.000 | 28.000 | 12.000 | | |
| Uniform Time Scaling Invariance | | | | | | | |
| Warping Invariance | | | | | | | |
| Non-positive Value Handling | 2.000 | 2.000 | 3.000 | | | | |
| Non-negativity | 1.000 | | | | | | |
| Triangle Inequality | 1.000 | | | | | | |
| Relative Sensitivity Ranges | 1.000 | 1.000 | 1.000 | 1.000 | 1.000 | 1.000 | 1.000 |

Controlled Test Results: NCD

| Test | Res.1 | Res.2 | Res.3 | Res.4 | Res.5 | Res.6 | Res.7 |
|---|---|---|---|---|---|---|---|
| Uniqueness | 0.1176 | | | | | | |
| Symmetry | 0.5000 | 0.4583 | | | | | |
| Translation Sensitivity | 0.4737 | 0.4737 | 0.4737 | 0.4737 | 0.4737 | | |
| Amplitude Sensitivity | 0.4091 | 0.3810 | 0.4348 | 0.3810 | 0.4348 | | |
| Duration Sensitivity | 0.1176 | 0.1176 | 0.1765 | 0.1765 | 0.2941 | | |
| Frequency Sensitivity | 0.3125 | 0.3125 | 0.3125 | 0.3333 | 0.1429 | | |
| White Noise Sensitivity | 0.7941 | 0.7500 | 0.7647 | 0.6800 | 0.7647 | | |
| Biased Noise Sensitivity | 0.5000 | 0.4815 | 0.5000 | 0.5417 | 0.6071 | | |
| Outlier Sensitivity | 0.3182 | 0.3182 | 0.3182 | 0.3182 | 0.3182 | | |
| Antiparallelism Bias | 0.2632 | 0.2632 | | | | | |
| Phase Invariance | 0.1250 | 0.0833 | 0.2083 | 0.1667 | 0.1667 | | |
| Uniform Time Scaling Invariance | 0.6429 | 0.6458 | 0.6731 | 0.7091 | 0.6098 | | |
| Warping Invariance | 0.1111 | 0.1111 | 0.1111 | 0.1579 | 0.1579 | | |
| Non-positive Value Handling | 0.4545 | 0.3333 | 0.3333 | | | | |
| Non-negativity | 1.0000 | | | | | | |
| Triangle Inequality | 1.0000 | | | | | | |
| Relative Sensitivity Ranges | 0.0000 | 0.4075 | 1.3358 | 1.4418 | 0.8638 | 0.9512 | 0.0000 |

Controlled Test Results: PACF

| Test | Res.1 | Res.2 | Res.3 | Res.4 | Res.5 | Res.6 | Res.7 |
|---|---|---|---|---|---|---|---|
| Uniqueness | 0.0000 | | | | | | |
| Symmetry | 0.6472 | 0.6472 | | | | | |
| Translation Sensitivity | 0.0000 | 0.0000 | 0.0000 | 0.0000 | 0.0000 | | |
| Amplitude Sensitivity | 0.1346 | 0.2640 | 0.3414 | 0.3904 | 0.4238 | | |
| Duration Sensitivity | 0.8358 | 0.6037 | 0.8581 | 0.3727 | 0.7841 | | |
| Frequency Sensitivity | 0.7398 | 0.8911 | 0.8791 | 0.7482 | 0.8603 | | |
| White Noise Sensitivity | 0.1006 | 0.2073 | 0.3138 | 0.4162 | 0.5122 | | |
| Biased Noise Sensitivity | 0.1096 | 0.2155 | 0.3131 | 0.4011 | 0.4796 | | |
| Outlier Sensitivity | 0.3645 | 0.5340 | 0.5184 | 0.4628 | 0.4217 | | |
| Antiparallelism Bias | 0.0000 | 0.0000 | | | | | |
| Phase Invariance | 0.5308 | 0.5950 | 0.5913 | 0.3674 | 0.2403 | | |
| Uniform Time Scaling Invariance | | | | | | | |
| Warping Invariance | | | | | | | |
| Non-positive Value Handling | 0.8325 | 0.8325 | 0.9658 | | | | |
| Non-negativity | 1.0000 | | | | | | |
| Triangle Inequality | 1.0000 | | | | | | |
| Relative Sensitivity Ranges | 0.0000 | 1.0786 | 1.8102 | 0.5642 | 1.5350 | 1.3798 | 0.6321 |

Controlled Test Results: Per

| Test | Res.1 | Res.2 | Res.3 | Res.4 | Res.5 | Res.6 | Res.7 |
|---|---|---|---|---|---|---|---|
| Uniqueness | 0.0000 | | | | | | |
| Symmetry | 0.8701 | 0.8701 | | | | | |
| Translation Sensitivity | 0.0000 | 0.0000 | 0.0000 | 0.0000 | 0.0000 | | |
| Amplitude Sensitivity | 0.1000 | 0.2404 | 0.4218 | 0.6445 | 0.9086 | | |
| Duration Sensitivity | 0.0623 | 0.1053 | 0.1463 | 0.1303 | 0.0771 | | |
| Frequency Sensitivity | 0.0582 | 0.1368 | 0.2421 | 0.3819 | 0.4624 | | |
| White Noise Sensitivity | 0.1191 | 0.2836 | 0.4936 | 0.7494 | 1.0511 | | |
| Biased Noise Sensitivity | 0.1389 | 0.3176 | 0.5385 | 0.8029 | 1.1115 | | |
| Outlier Sensitivity | 0.3066 | 0.6883 | 1.1539 | 1.7090 | 2.3572 | | |
| Antiparallelism Bias | 0.0000 | 0.1666 | | | | | |
| Phase Invariance | 0.8618 | 1.3408 | 1.1143 | 1.0548 | 0.9195 | | |
| Uniform Time Scaling Invariance | | | | | | | |
| Warping Invariance | | | | | | | |
| Non-positive Value Handling | 0.1963 | 0.1963 | 0.4163 | | | | |
| Non-negativity | 1.0000 | | | | | | |
| Triangle Inequality | 1.0000 | | | | | | |
| Relative Sensitivity Ranges | 0.0000 | 1.0778 | 0.1120 | 0.5386 | 1.2421 | 1.2963 | 2.7332 |

Controlled Test Results: ProbSymm

| Test | Res.1 | Res.2 | Res.3 | Res.4 | Res.5 | Res.6 | Res.7 |
|---|---|---|---|---|---|---|---|
| Uniqueness | 0.000 | | | | | | |
| Symmetry | 12.279 | 12.279 | | | | | |
| Translation Sensitivity | 0.086 | 0.329 | 0.710 | 1.212 | 1.822 | | |
| Amplitude Sensitivity | 0.222 | 0.800 | 1.636 | 2.667 | 3.846 | | |
| Duration Sensitivity | 0.667 | 1.333 | 2.000 | 2.667 | 3.333 | | |
| Frequency Sensitivity | 0.667 | 1.333 | 2.000 | 2.667 | 3.333 | | |
| White Noise Sensitivity | 0.041 | 0.163 | 0.368 | 0.655 | 1.028 | | |
| Biased Noise Sensitivity | 0.079 | 0.302 | 0.650 | 1.108 | 1.662 | | |
| Outlier Sensitivity | 0.222 | 0.800 | 1.636 | 2.667 | 3.846 | | |
| Antiparallelism Bias | 2.000 | 1.000 | | | | | |
| Phase Invariance | 31.810 | 17.854 | 18.334 | 33.676 | 6.267 | | |
| Uniform Time Scaling Invariance | | | | | | | |
| Warping Invariance | | | | | | | |
| Non-positive Value Handling | 4.000 | 4.000 | 18.000 | | | | |
| Non-negativity | 0.000 | | | | | | |
| Triangle Inequality | 0.000 | | | | | | |
| Relative Sensitivity Ranges | 0.720 | 1.502 | 1.105 | 1.105 | 0.409 | 0.656 | 1.502 |

Controlled Test Results: Soergel

| Test | Res.1 | Res.2 | Res.3 | Res.4 | Res.5 | Res.6 | Res.7 |
|---|---|---|---|---|---|---|---|
| Uniqueness | 0.0000 | | | | | | |
| Symmetry | 0.5818 | 0.5818 | | | | | |
| Translation Sensitivity | 0.0769 | 0.1429 | 0.2000 | 0.2500 | 0.2941 | | |
| Amplitude Sensitivity | 0.0667 | 0.1250 | 0.1765 | 0.2222 | 0.2632 | | |
| Duration Sensitivity | 0.0769 | 0.1429 | 0.2000 | 0.2500 | 0.2941 | | |
| Frequency Sensitivity | 0.0714 | 0.1333 | 0.1875 | 0.2353 | 0.2778 | | |
| White Noise Sensitivity | 0.0308 | 0.0606 | 0.0896 | 0.1176 | 0.1449 | | |
| Biased Noise Sensitivity | 0.0303 | 0.0588 | 0.0857 | 0.1111 | 0.1351 | | |
| Outlier Sensitivity | 0.0400 | 0.0769 | 0.1111 | 0.1429 | 0.1724 | | |
| Antiparallelism Bias | 0.1818 | 0.1538 | | | | | |
| Phase Invariance | 0.6667 | 0.4865 | 0.4865 | 0.6667 | 0.3529 | | |
| Uniform Time Scaling Invariance | | | | | | | |
| Warping Invariance | | | | | | | |
| Non-positive Value Handling | 0.1429 | 0.1429 | 0.2143 | | | | |
| Non-negativity | 1.0000 | | | | | | |
| Triangle Inequality | 1.0000 | | | | | | |
| Relative Sensitivity Ranges | 1.2791 | 1.1572 | 1.2791 | 1.2152 | 0.6723 | 0.6174 | 0.7798 |

Controlled Test Results: SqChi

| Test | Res.1 | Res.2 | Res.3 | Res.4 | Res.5 | Res.6 | Res.7 |
|---|---|---|---|---|---|---|---|
| Uniqueness | 0.000 | | | | | | |
| Symmetry | 6.139 | 6.139 | | | | | |
| Translation Sensitivity | 0.043 | 0.165 | 0.355 | 0.606 | 0.911 | | |
| Amplitude Sensitivity | 0.111 | 0.400 | 0.818 | 1.333 | 1.923 | | |
| Duration Sensitivity | 0.333 | 0.667 | 1.000 | 1.333 | 1.667 | | |
| Frequency Sensitivity | 0.333 | 0.667 | 1.000 | 1.333 | 1.667 | | |
| White Noise Sensitivity | 0.020 | 0.082 | 0.184 | 0.328 | 0.514 | | |
| Biased Noise Sensitivity | 0.040 | 0.151 | 0.325 | 0.554 | 0.831 | | |
| Outlier Sensitivity | 0.111 | 0.400 | 0.818 | 1.333 | 1.923 | | |
| Antiparallelism Bias | 1.000 | 0.500 | | | | | |
| Phase Invariance | 15.905 | 8.927 | 9.167 | 16.838 | 3.133 | | |
| Uniform Time Scaling Invariance | | | | | | | |
| Warping Invariance | | | | | | | |
| Non-positive Value Handling | 2.000 | 2.000 | 9.000 | | | | |
| Non-negativity | 0.000 | | | | | | |
| Triangle Inequality | 0.000 | | | | | | |
| Relative Sensitivity Ranges | 0.720 | 1.502 | 1.105 | 1.105 | 0.409 | 0.656 | 1.502 |

Controlled Test Results: SqChord

| Test | Res.1 | Res.2 | Res.3 | Res.4 | Res.5 | Res.6 | Res.7 |
|---|---|---|---|---|---|---|---|
| Uniqueness | 0.0000 | | | | | | |
| Symmetry | 3.4299 | 3.4299 | | | | | |
| Translation Sensitivity | 0.0215 | 0.0824 | 0.1781 | 0.3050 | 0.4598 | | |
| Amplitude Sensitivity | 0.0557 | 0.2020 | 0.4170 | 0.6863 | 1.0000 | | |
| Duration Sensitivity | 0.1716 | 0.3431 | 0.5147 | 0.6863 | 0.8579 | | |
| Frequency Sensitivity | 0.1716 | 0.3431 | 0.5147 | 0.6863 | 0.8579 | | |
| White Noise Sensitivity | 0.0102 | 0.0408 | 0.0921 | 0.1643 | 0.2580 | | |
| Biased Noise Sensitivity | 0.0198 | 0.0757 | 0.1633 | 0.2791 | 0.4201 | | |
| Outlier Sensitivity | 0.0557 | 0.2020 | 0.4170 | 0.6863 | 1.0000 | | |
| Antiparallelism Bias | 0.5359 | 0.2540 | | | | | |
| Phase Invariance | 8.9453 | 4.9998 | 5.1372 | 9.4719 | 1.5984 | | |
| Uniform Time Scaling Invariance | | | | | | | |
| Warping Invariance | | | | | | | |
| Non-positive Value Handling | 1.9972 | 2.0000 | | | | | |
| Non-negativity | 1.0000 | | | | | | |
| Triangle Inequality | 0.0000 | | | | | | |
| Relative Sensitivity Ranges | 0.7057 | 1.5204 | 1.1050 | 1.1050 | 0.3990 | 0.6446 | 1.5204 |

Controlled Test Results: SqEuclid

| Test | Res.1 | Res.2 | Res.3 | Res.4 | Res.5 | Res.6 | Res.7 |
|---|---|---|---|---|---|---|---|
| Uniqueness | 0.00 | | | | | | |
| Symmetry | 20.62 | 20.62 | | | | | |
| Translation Sensitivity | 0.10 | 0.40 | 0.90 | 1.60 | 2.50 | | |
| Amplitude Sensitivity | 0.50 | 2.00 | 4.50 | 8.00 | 12.50 | | |
| Duration Sensitivity | 1.00 | 2.00 | 3.00 | 4.00 | 5.00 | | |
| Frequency Sensitivity | 1.00 | 2.00 | 3.00 | 4.00 | 5.00 | | |
| White Noise Sensitivity | 0.12 | 0.50 | 1.12 | 2.00 | 3.12 | | |
| Biased Noise Sensitivity | 0.25 | 1.00 | 2.25 | 4.00 | 6.25 | | |
| Outlier Sensitivity | 1.00 | 4.00 | 9.00 | 16.00 | 25.00 | | |
| Antiparallelism Bias | 2.00 | 2.00 | | | | | |
| Phase Invariance | 92.00 | 56.00 | 56.00 | 100.00 | 24.00 | | |
| Uniform Time Scaling Invariance | | | | | | | |
| Warping Invariance | | | | | | | |
| Non-positive Value Handling | 4.00 | 4.00 | 9.00 | | | | |
| Non-negativity | 1.00 | | | | | | |
| Triangle Inequality | 0.00 | | | | | | |
| Relative Sensitivity Ranges | 0.30 | 1.52 | 0.51 | 0.51 | 0.38 | 0.76 | 3.03 |

Controlled Test Results: STS

| Test | Res.1 | Res.2 | Res.3 | Res.4 | Res.5 | Res.6 | Res.7 |
|---|---|---|---|---|---|---|---|
| Uniqueness | 0.000 | | | | | | |
| Symmetry | 5.257 | 5.257 | | | | | |
| Translation Sensitivity | 0.000 | 0.000 | 0.000 | 0.000 | 0.000 | | |
| Amplitude Sensitivity | 0.707 | 1.414 | 2.121 | 2.828 | 3.536 | | |
| Duration Sensitivity | 1.414 | 1.414 | 1.414 | 1.414 | 1.414 | | |
| Frequency Sensitivity | 1.414 | 2.000 | 2.449 | 2.828 | 3.000 | | |
| White Noise Sensitivity | 0.685 | 1.369 | 2.054 | 2.739 | 3.423 | | |
| Biased Noise Sensitivity | 0.707 | 1.414 | 2.121 | 2.828 | 3.536 | | |
| Outlier Sensitivity | 1.414 | 2.828 | 4.243 | 5.657 | 7.071 | | |
| Antiparallelism Bias | 1.414 | 1.414 | | | | | |
| Phase Invariance | 15.748 | 12.124 | 11.662 | 14.933 | 5.292 | | |
| Uniform Time Scaling Invariance | | | | | | | |
| Warping Invariance | | | | | | | |
| Non-positive Value Handling | 2.828 | 2.828 | 4.243 | | | | |
| Non-negativity | 1.000 | | | | | | |
| Triangle Inequality | 1.000 | | | | | | |
| Relative Sensitivity Ranges | 0.000 | 1.085 | 0.000 | 0.608 | 1.051 | 1.085 | 2.170 |

Controlled Test Results: TAM

| Test | Res.1 | Res.2 | Res.3 | Res.4 | Res.5 | Res.6 | Res.7 |
|---|---|---|---|---|---|---|---|
| Uniqueness | 0.0000 | | | | | | |
| Symmetry | 2.0000 | 2.0000 | | | | | |
| Translation Sensitivity | 0.0000 | 0.6667 | 0.6667 | 0.6667 | 0.0000 | | |
| Amplitude Sensitivity | 0.0000 | 0.0000 | 0.6667 | 0.6667 | 0.6667 | | |
| Duration Sensitivity | 0.3333 | 0.6667 | 1.0000 | 1.3333 | 1.6667 | | |
| Frequency Sensitivity | 0.2727 | 0.5455 | 0.8182 | 1.0909 | 1.3636 | | |
| White Noise Sensitivity | 0.0000 | 0.0000 | 0.0000 | 0.0000 | 0.3333 | | |
| Biased Noise Sensitivity | 0.0000 | 0.0000 | 0.3333 | 0.6667 | 1.0000 | | |
| Outlier Sensitivity | 0.0000 | 0.0000 | 0.3333 | 0.3333 | 0.3333 | | |
| Antiparallelism Bias | 1.0000 | 0.6667 | | | | | |
| Phase Invariance | 0.3333 | 0.6667 | 1.0000 | 1.3333 | 1.6667 | | |
| Uniform Time Scaling Invariance | 0.7444 | 0.2500 | 0.3571 | 0.4375 | 0.5000 | | |
| Warping Invariance | 0.1000 | 0.1818 | 0.2500 | 0.3077 | 0.3571 | | |
| Non-positive Value Handling | 0.3333 | 0.3333 | 0.3333 | | | | |
| Non-negativity | 1.0000 | | | | | | |
| Triangle Inequality | 0.0000 | | | | | | |
| Relative Sensitivity Ranges | 0.8603 | 0.8603 | 1.7207 | 1.4078 | 0.4302 | 1.2905 | 0.4302 |

Controlled Test Results: Taneja

| Test | Res.1 | Res.2 | Res.3 | Res.4 | Res.5 | Res.6 | Res.7 |
|---|---|---|---|---|---|---|---|
| Uniqueness | 0.000 | | | | | | |
| Symmetry | 1.947 | 1.947 | | | | | |
| Translation Sensitivity | 0.011 | 0.041 | 0.089 | 0.153 | 0.232 | | |
| Amplitude Sensitivity | 0.028 | 0.102 | 0.213 | 0.353 | 0.520 | | |
| Duration Sensitivity | 0.088 | 0.177 | 0.265 | 0.353 | 0.442 | | |
| Frequency Sensitivity | 0.088 | 0.177 | 0.265 | 0.353 | 0.442 | | |
| White Noise Sensitivity | 0.005 | 0.020 | 0.046 | 0.082 | 0.130 | | |
| Biased Noise Sensitivity | 0.010 | 0.038 | 0.082 | 0.141 | 0.212 | | |
| Outlier Sensitivity | 0.028 | 0.102 | 0.213 | 0.353 | 0.520 | | |
| Antiparallelism Bias | 0.288 | 0.129 | | | | | |
| Phase Invariance | 5.071 | 2.825 | 2.903 | 5.367 | 0.816 | | |
| Uniform Time Scaling Invariance | | | | | | | |
| Warping Invariance | | | | | | | |
| Non-positive Value Handling | 6.561 | 12.206 | | | | | |
| Non-negativity | 1.000 | | | | | | |
| Triangle Inequality | 0.000 | | | | | | |
| Relative Sensitivity Ranges | 0.692 | 1.539 | 1.104 | 1.104 | 0.389 | 0.633 | 1.539 |

Controlled Test Results: Topsoe

| Test | Res.1 | Res.2 | Res.3 | Res.4 | Res.5 | Res.6 | Res.7 |
|---|---|---|---|---|---|---|---|
| Uniqueness | 0.0000 | | | | | | |
| Symmetry | 3.2919 | 3.2919 | | | | | |
| Translation Sensitivity | 0.0215 | 0.0824 | 0.1779 | 0.3043 | 0.4584 | | |
| Amplitude Sensitivity | 0.0557 | 0.2014 | 0.4143 | 0.6796 | 0.9868 | | |
| Duration Sensitivity | 0.1699 | 0.3398 | 0.5097 | 0.6796 | 0.8495 | | |
| Frequency Sensitivity | 0.1699 | 0.3398 | 0.5097 | 0.6796 | 0.8495 | | |
| White Noise Sensitivity | 0.0102 | 0.0408 | 0.0920 | 0.1641 | 0.2577 | | |
| Biased Noise Sensitivity | 0.0198 | 0.0757 | 0.1631 | 0.2784 | 0.4186 | | |
| Outlier Sensitivity | 0.0557 | 0.2014 | 0.4143 | 0.6796 | 0.9868 | | |
| Antiparallelism Bias | 0.5232 | 0.2527 | | | | | |
| Phase Invariance | 8.5772 | 4.8003 | 4.9316 | 9.0826 | 1.5876 | | |
| Uniform Time Scaling Invariance | | | | | | | |
| Warping Invariance | | | | | | | |
| Non-positive Value Handling | 1.3863 | | | | | | |
| Non-negativity | 1.0000 | | | | | | |
| Triangle Inequality | 0.0000 | | | | | | |
| Relative Sensitivity Ranges | 0.7105 | 1.5142 | 1.1052 | 1.1052 | 0.4024 | 0.6485 | 1.5142 |

Controlled Test Results: WaveHedges

| Test | Res.1 | Res.2 | Res.3 | Res.4 | Res.5 | Res.6 | Res.7 |
|---|---|---|---|---|---|---|---|
| Uniqueness | 0.0000 | | | | | | |
| Symmetry | 6.0500 | 6.0500 | | | | | |
| Translation Sensitivity | 0.8225 | 1.5152 | 2.1070 | 2.6190 | 3.0667 | | |
| Amplitude Sensitivity | 0.4000 | 0.6667 | 0.8571 | 1.0000 | 1.1111 | | |
| Duration Sensitivity | 0.5000 | 1.0000 | 1.5000 | 2.0000 | 2.5000 | | |
| Frequency Sensitivity | 0.5000 | 1.0000 | 1.5000 | 2.0000 | 2.5000 | | |
| White Noise Sensitivity | 0.3189 | 0.6231 | 0.9147 | 1.1955 | 1.4670 | | |
| Biased Noise Sensitivity | 0.3022 | 0.5538 | 0.7677 | 0.9524 | 1.1141 | | |
| Outlier Sensitivity | 0.2000 | 0.3333 | 0.4286 | 0.5000 | 0.5556 | | |
| Antiparallelism Bias | 1.3333 | 0.8000 | | | | | |
| Phase Invariance | 6.6000 | 4.0667 | 4.1000 | 6.4333 | 2.4667 | | |
| Uniform Time Scaling Invariance | | | | | | | |
| Warping Invariance | | | | | | | |
| Non-positive Value Handling | 1.0000 | 1.0000 | 1.5000 | | | | |
| Non-negativity | 0.0000 | | | | | | |
| Triangle Inequality | 0.0000 | | | | | | |
| Relative Sensitivity Ranges | 1.6945 | 0.5369 | 1.5101 | 1.5101 | 0.8669 | 0.6130 | 0.2685 |

#### 15. Plots of wading bird rankings for all distance measures

This section contains plots of wading bird dissimilarity results for all 42 distance measures we tested. Each figure shows dissimilarity results for both smoothed and unsmoothed indices of all five wading birds. Distance measures are presented in alphabetical order.

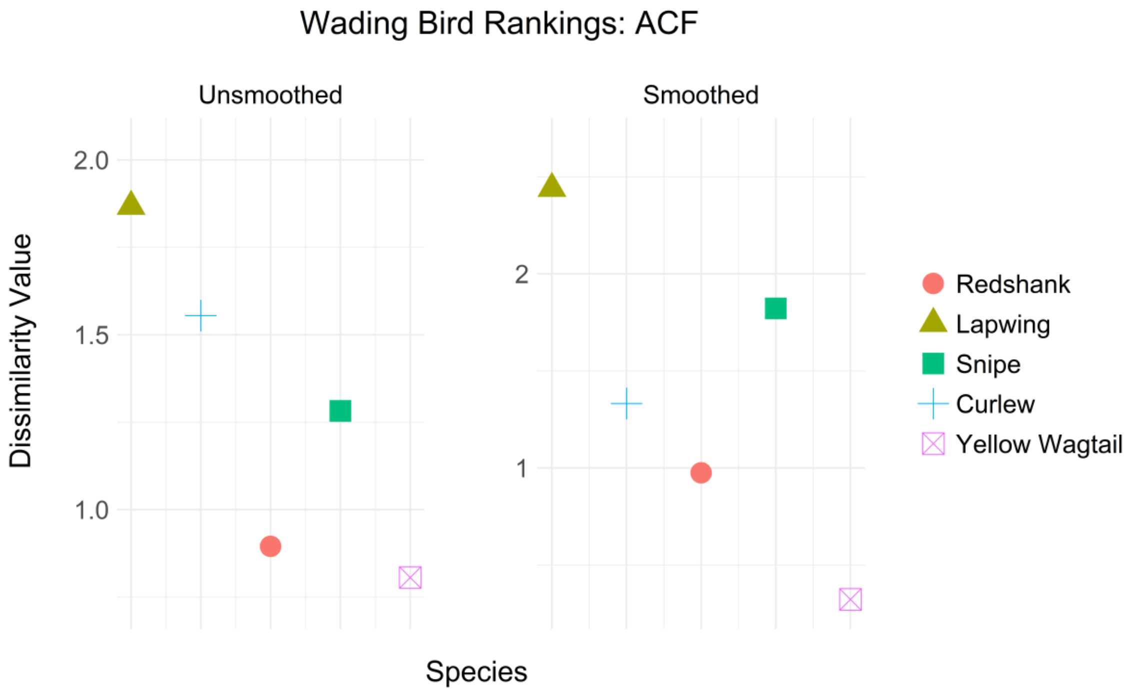


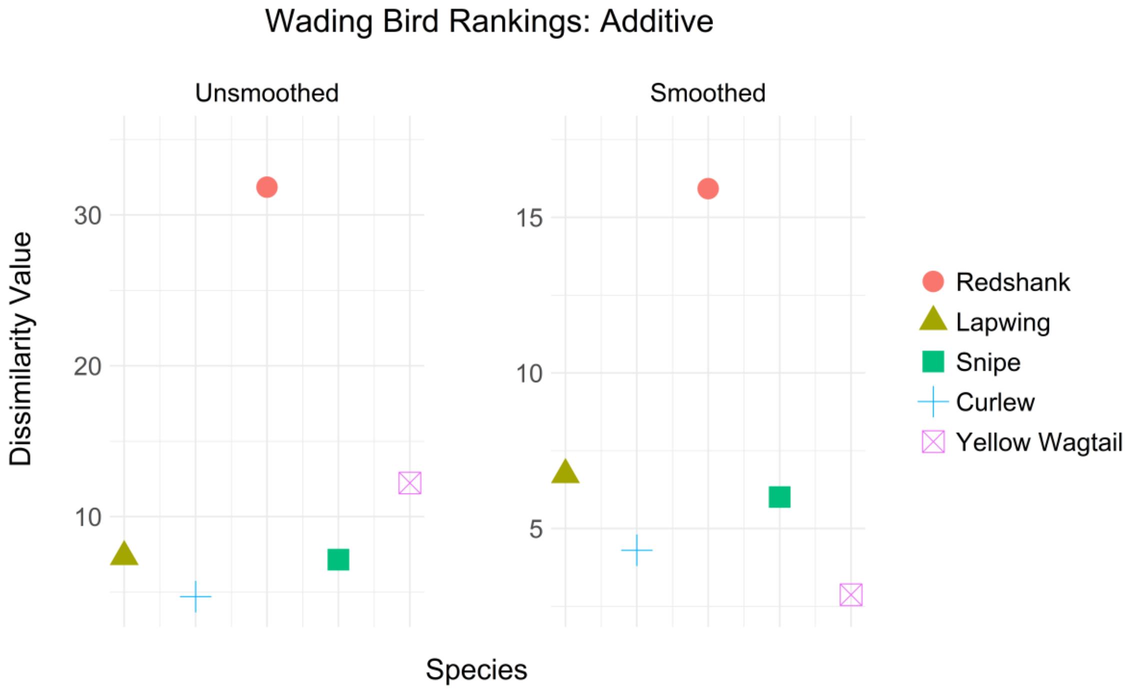

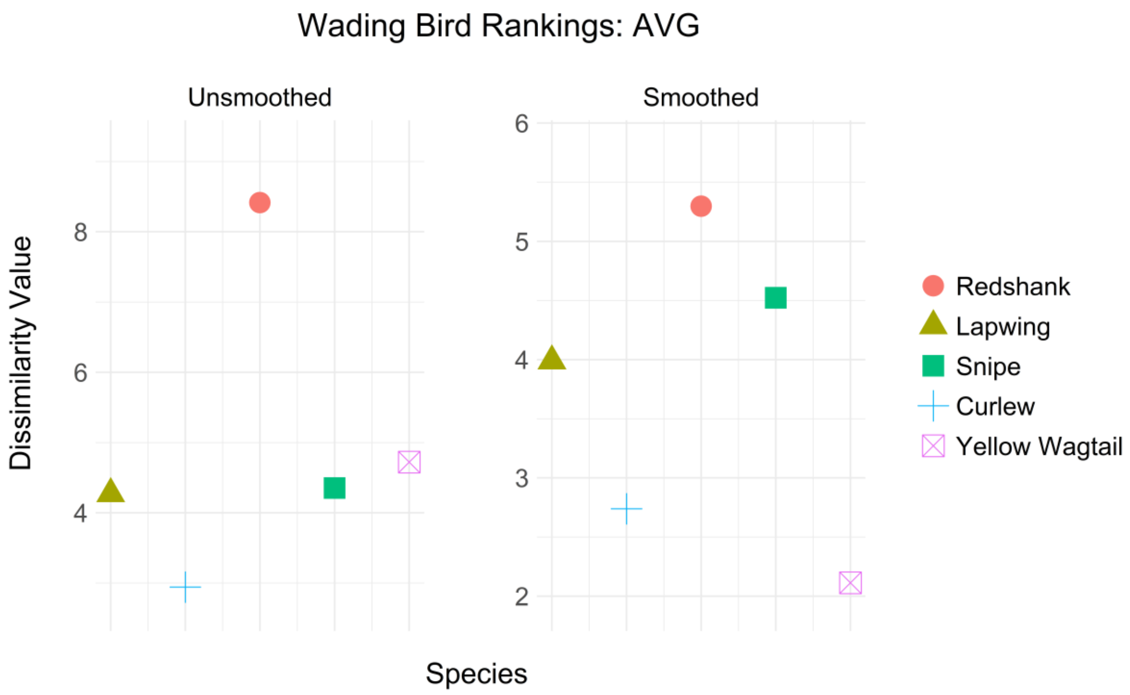
Wading Bird Rankings: AVG
Unsmoothed
Smoothed
Dissimilarity Value
8
6
4
6
5
4
3
2
Species
Redshank
Lapwing
Snipe
Curlew
Yellow Wagtail

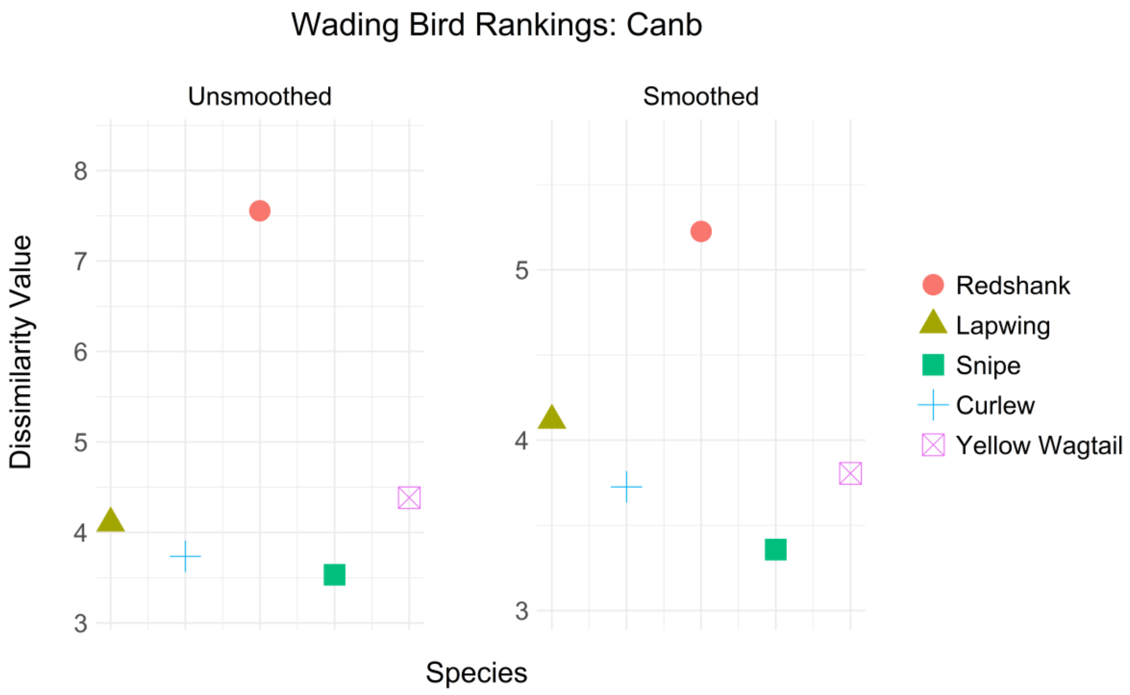
Wading Bird Rankings: Canb
Unsmoothed
Smoothed
Dissimilarity Value
8
7
6
5
4
3
5
4
3
Species
Redshank
Lapwing
Snipe
Curlew
Yellow Wagtail

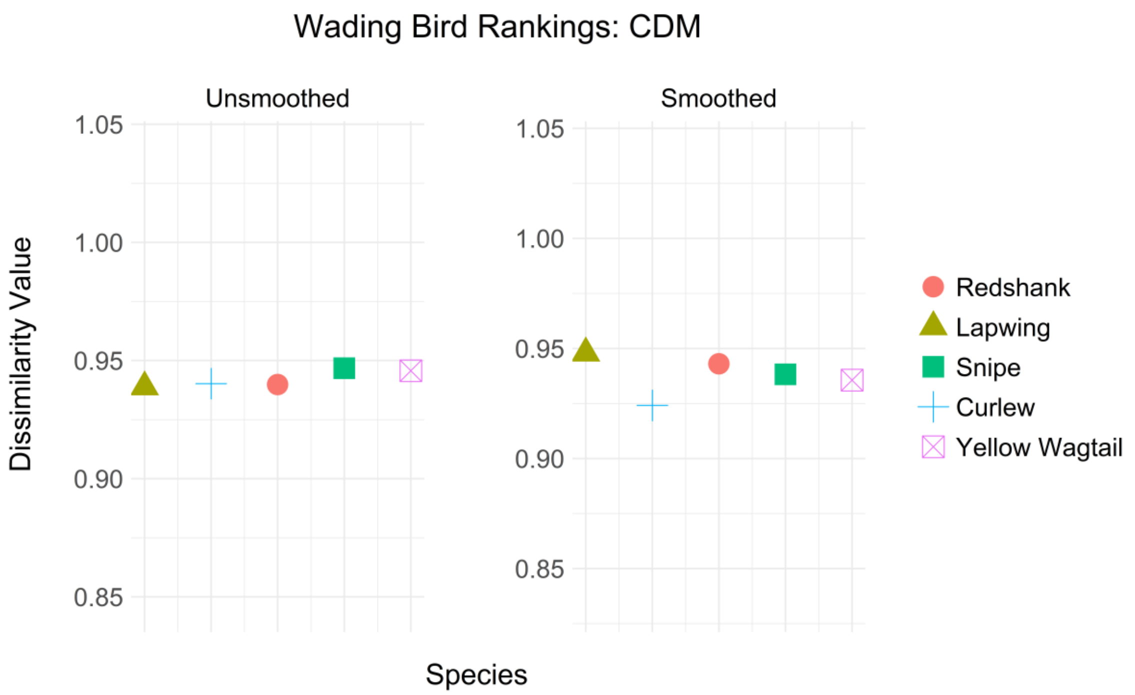

Wading Bird Rankings: CDM
Unsmoothed
Smoothed
Dissimilarity Value
1.05
1.00
0.95
0.90
0.85
Species
Redshank
Lapwing
Snipe
Curlew
Yellow Wagtail


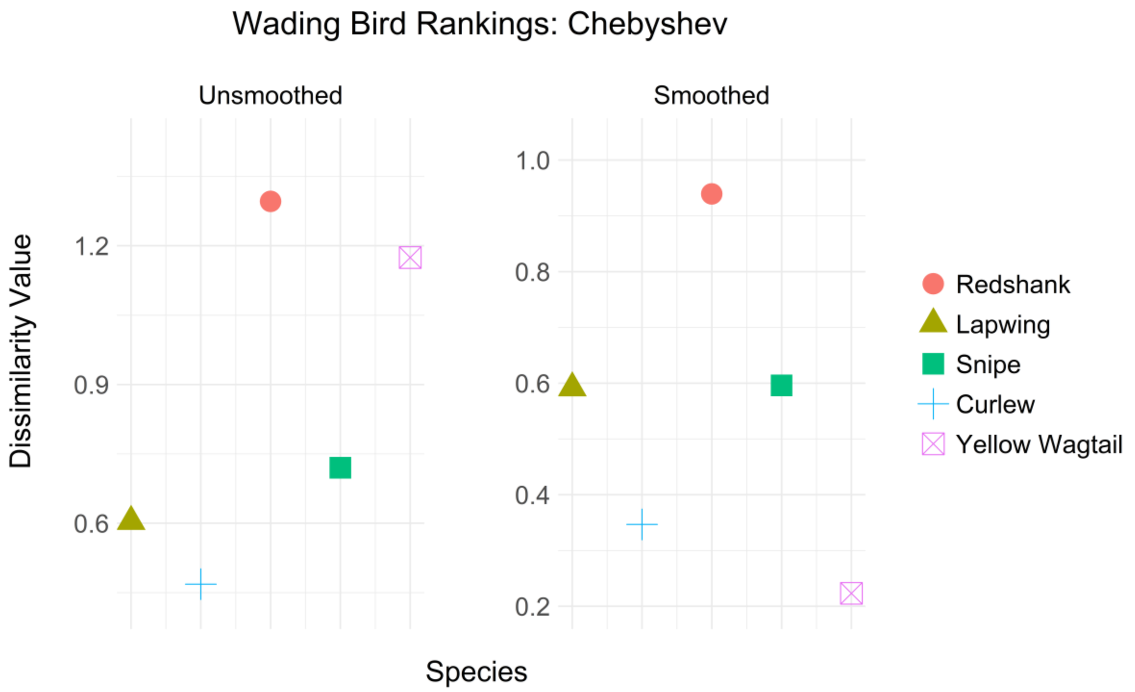

Wading Bird Rankings: Chebyshev
Unsmoothed
Smoothed
Dissimilarity Value
1.2
0.9
0.6
1.0
0.8
0.6
0.4
0.2
Species
Redshank
Lapwing
Snipe
Curlew
Yellow Wagtail

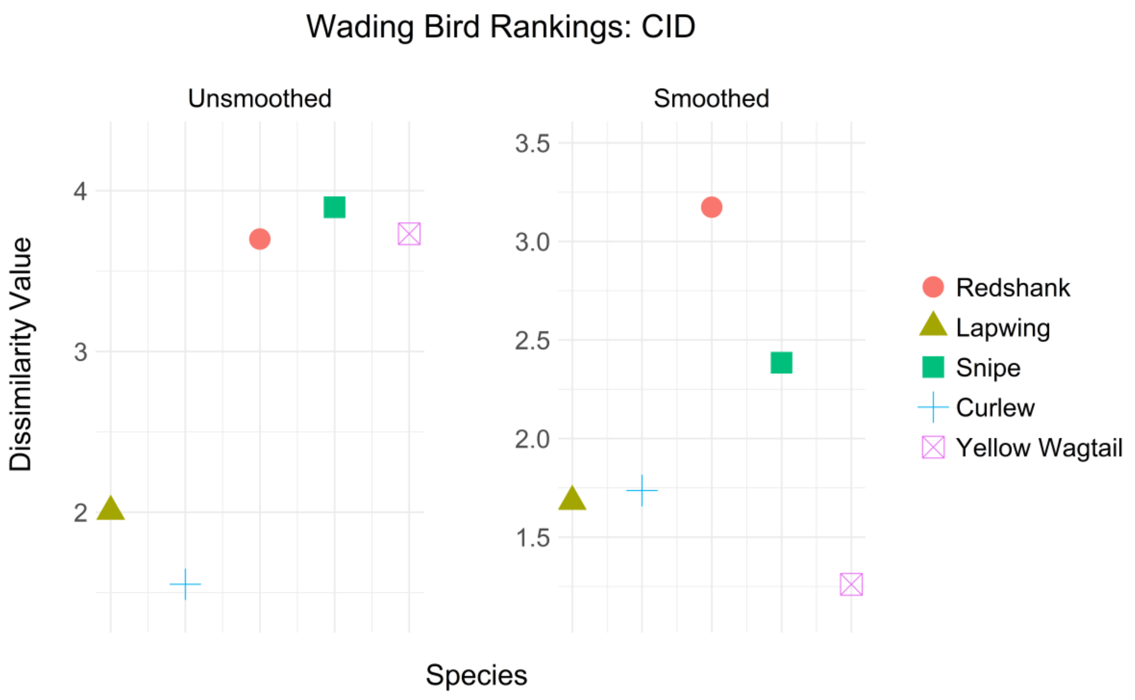
Wading Bird Rankings: CID
Unsmoothed
Smoothed
Dissimilarity Value
4
3
2
3.5
3.0
2.5
2.0
1.5
Species
Redshank
Lapwing
Snipe
Curlew
Yellow Wagtail

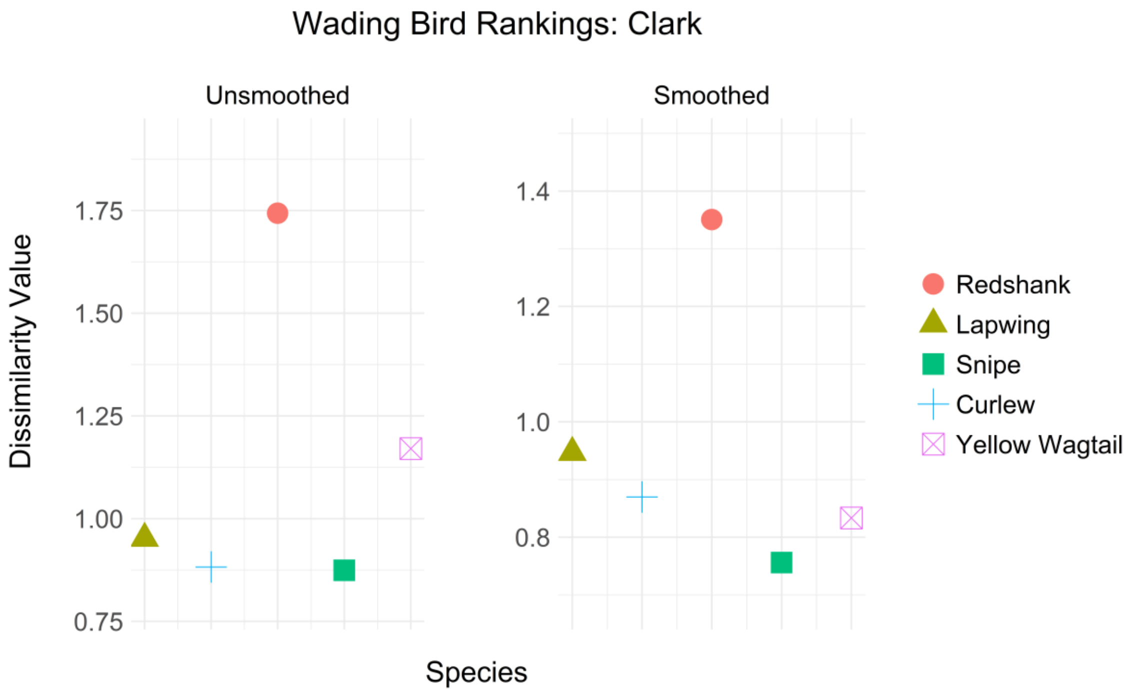
Wading Bird Rankings: Clark
Unsmoothed
Smoothed
Dissimilarity Value
1.75
1.50
1.25
1.00
0.75
1.4
1.2
1.0
0.8
Species
Redshank
Lapwing
Snipe
Curlew
Yellow Wagtail

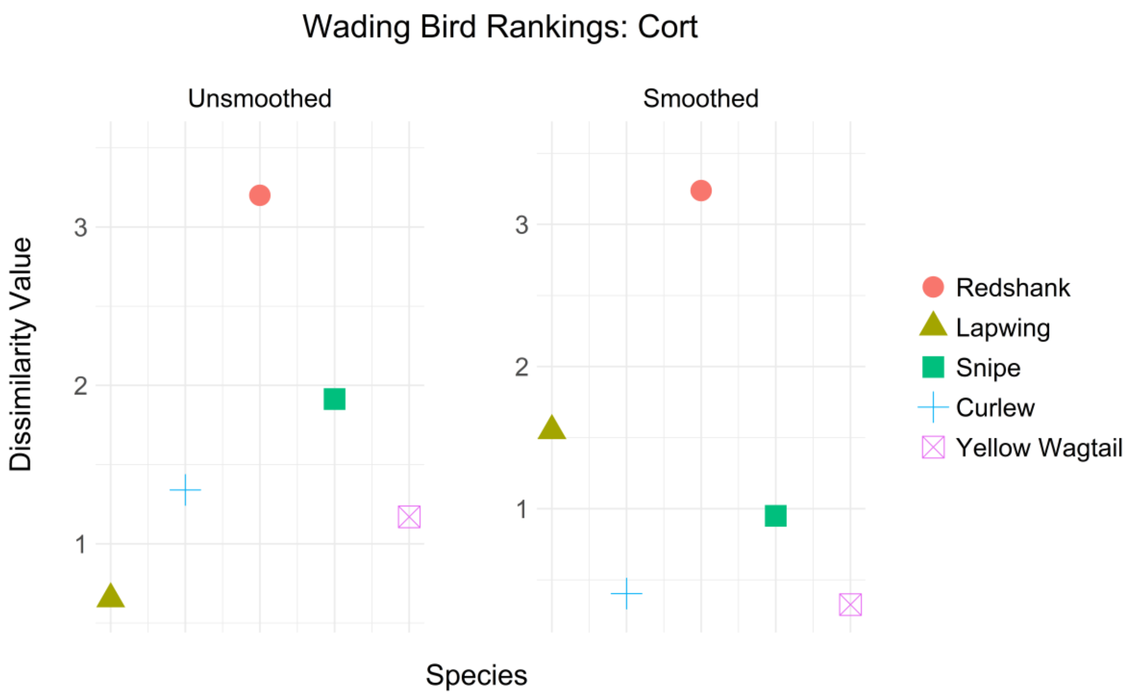

Wading Bird Rankings: Cort
Unsmoothed
Smoothed
Dissimilarity Value
3
2
1
Species
Redshank
Lapwing
Snipe
Curlew
Yellow Wagtail


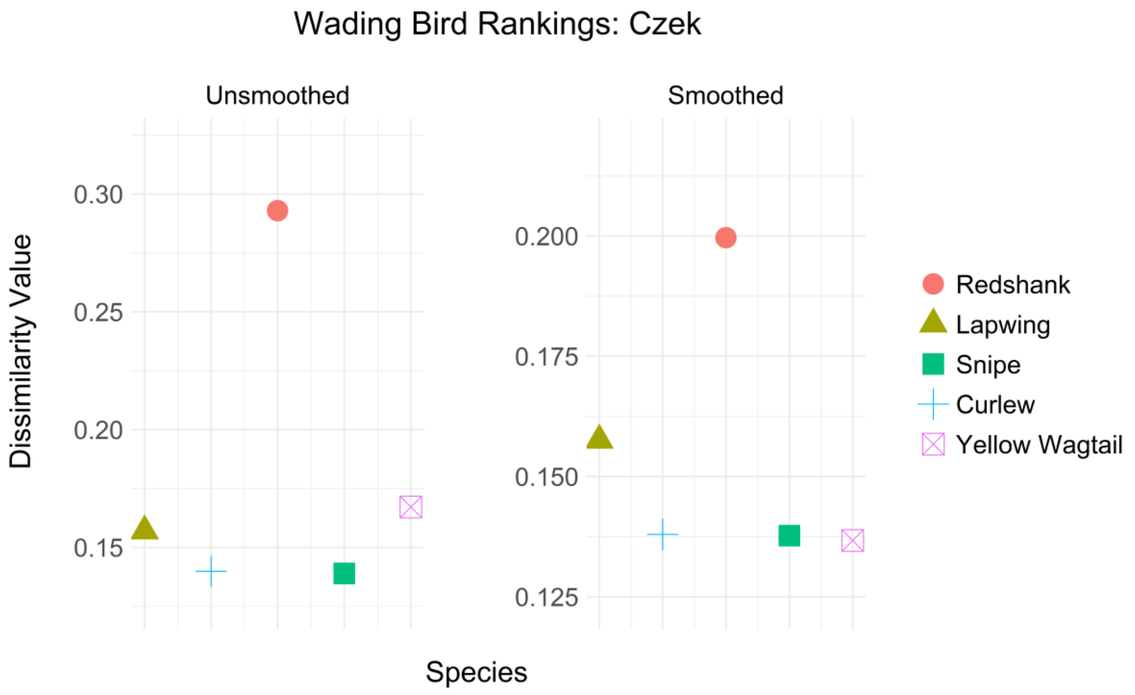

Wading Bird Rankings: Czek
Unsmoothed
Smoothed
Dissimilarity Value
0.30
0.25
0.20
0.15
0.200
0.175
0.150
0.125
Species
Redshank
Lapwing
Snipe
Curlew
Yellow Wagtail

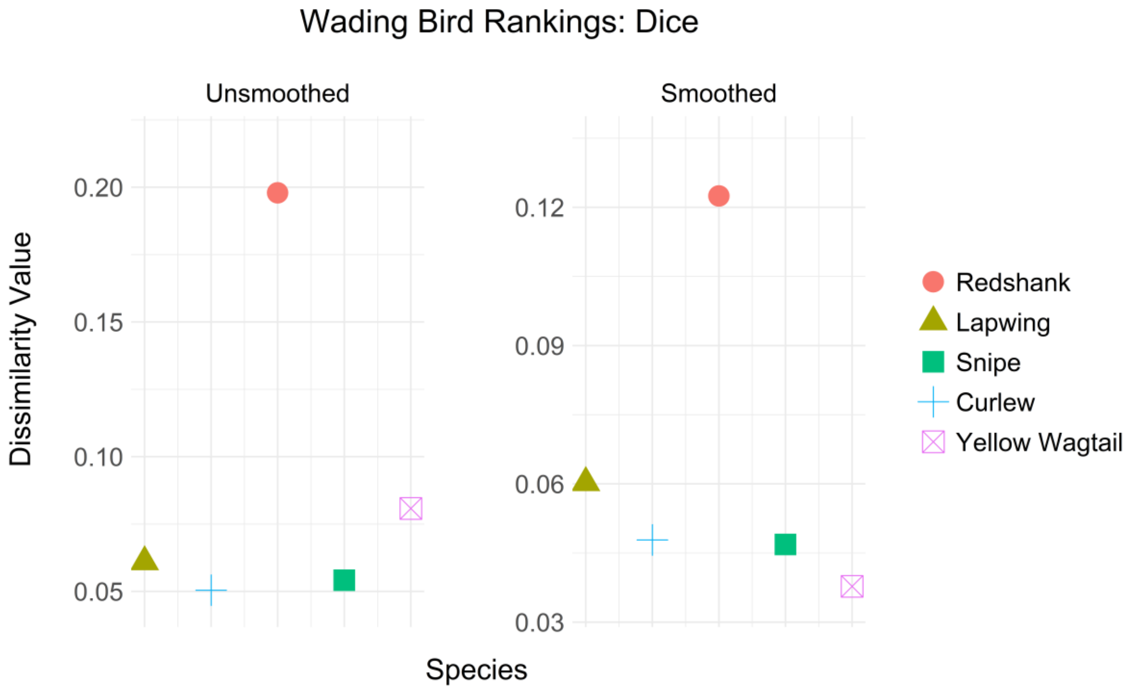
Wading Bird Rankings: Dice
Unsmoothed
Smoothed
Dissimilarity Value
0.20
0.15
0.10
0.05
0.12
0.09
0.06
0.03
Species
Redshank
Lapwing
Snipe
Curlew
Yellow Wagtail

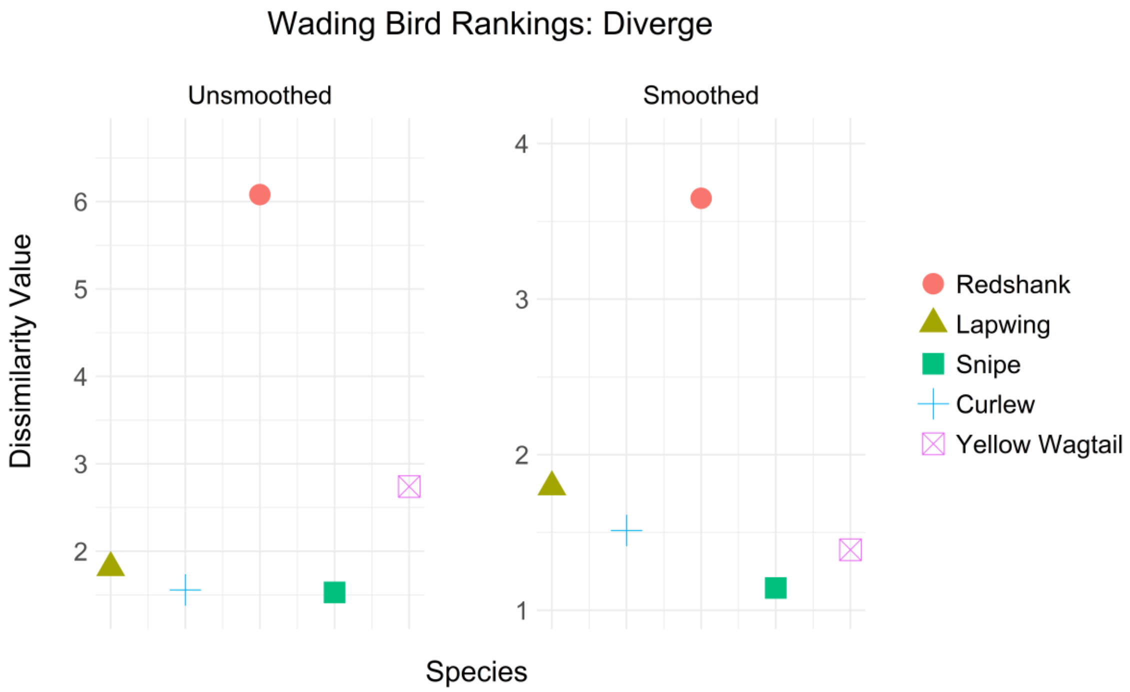
Wading Bird Rankings: Diverge
Unsmoothed
Smoothed
Dissimilarity Value
6
5
4
3
2
4
3
2
1
Species
Redshank
Lapwing
Snipe
Curlew
Yellow Wagtail

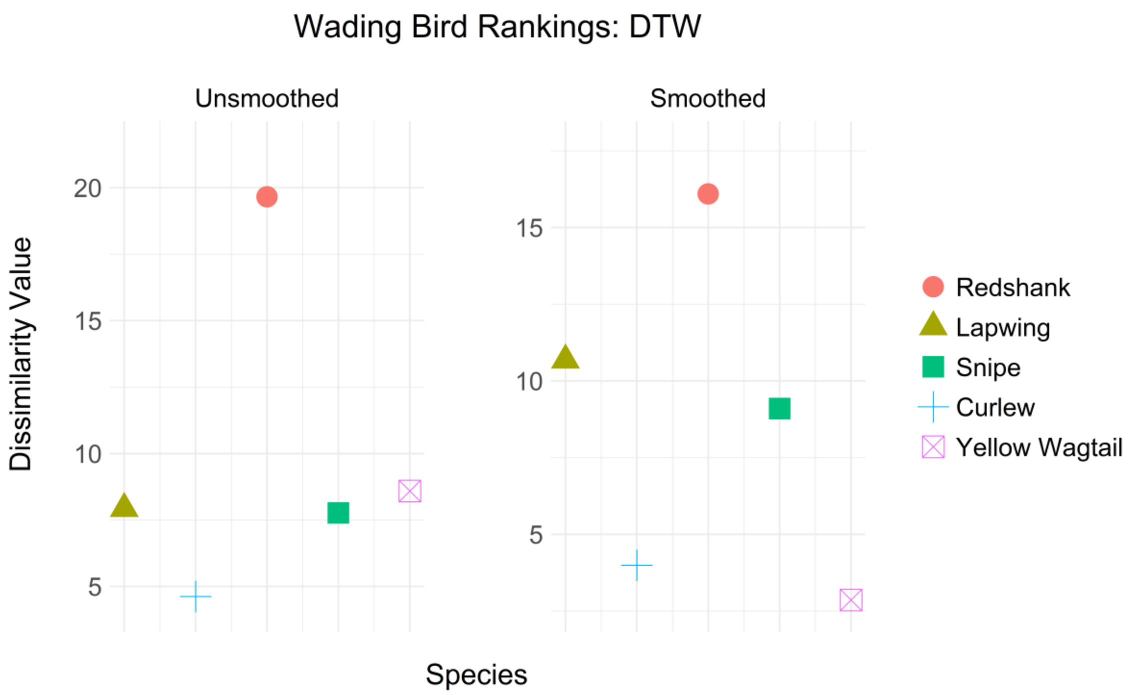
Wading Bird Rankings: DTW
Unsmoothed
Smoothed
Dissimilarity Value
20
15
10
5
Species
Redshank
Lapwing
Snipe
Curlew
Yellow Wagtail

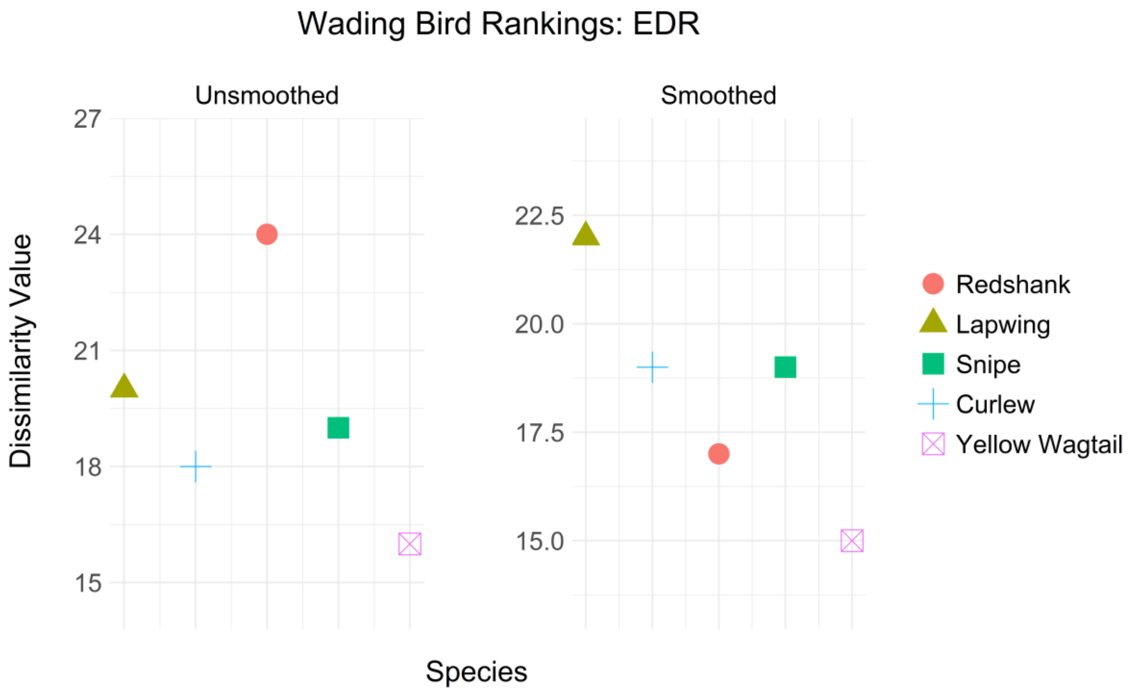
Wading Bird Rankings: EDR
Unsmoothed
Smoothed
Dissimilarity Value
27
24
21
18
15
22.5
20.0
17.5
15.0
Species
Redshank
Lapwing
Snipe
Curlew
Yellow Wagtail

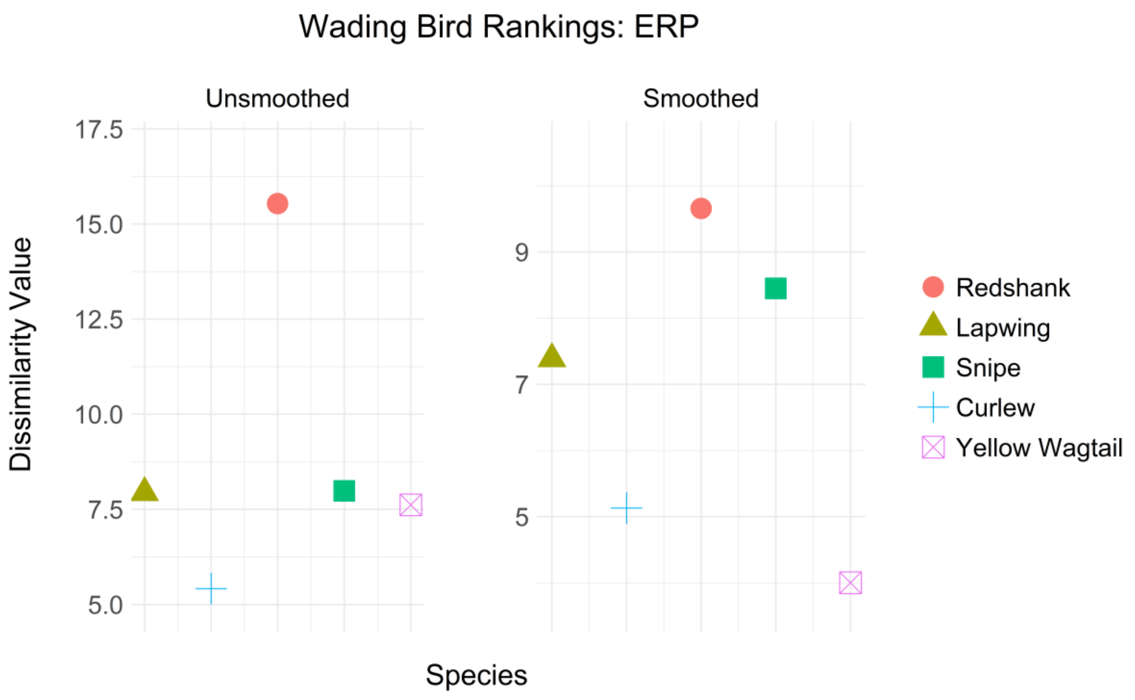
Wading Bird Rankings: ERP
Unsmoothed
Smoothed
17.5
15.0
12.5
10.0
7.5
5.0
9
7
5
Dissimilarity Value
Species
Redshank
Lapwing
Snipe
Curlew
Yellow Wagtail

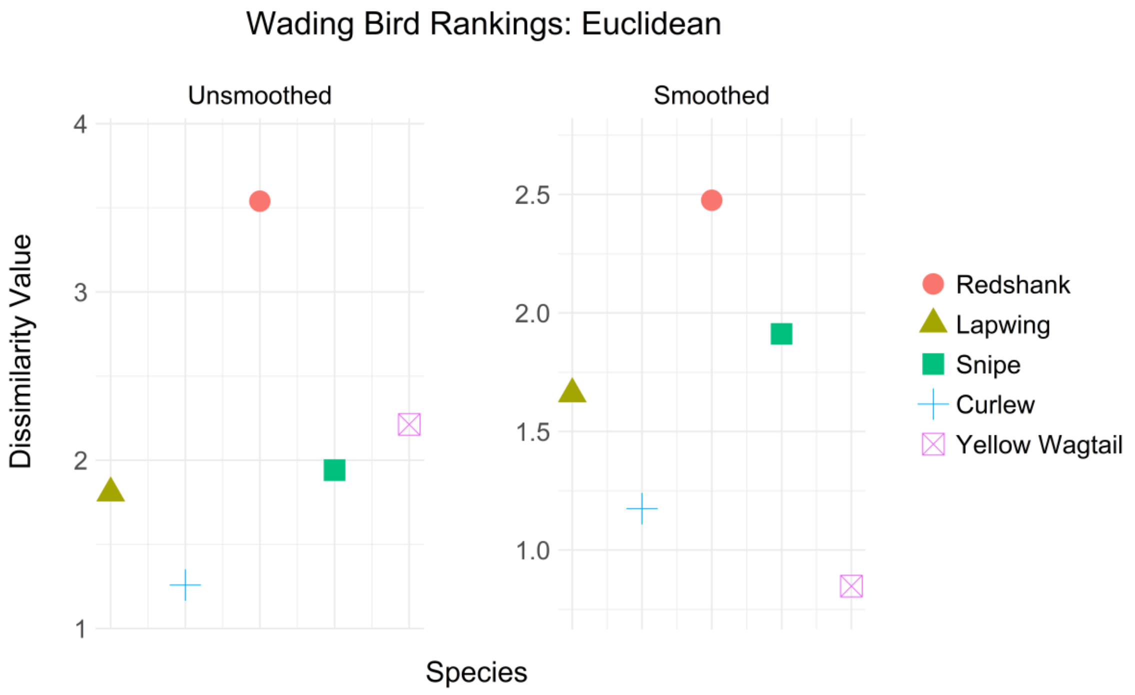
Wading Bird Rankings: Euclidean
Unsmoothed
Smoothed
4
3
2
1
2.5
2.0
1.5
1.0
Dissimilarity Value
Species
Redshank
Lapwing
Snipe
Curlew
Yellow Wagtail

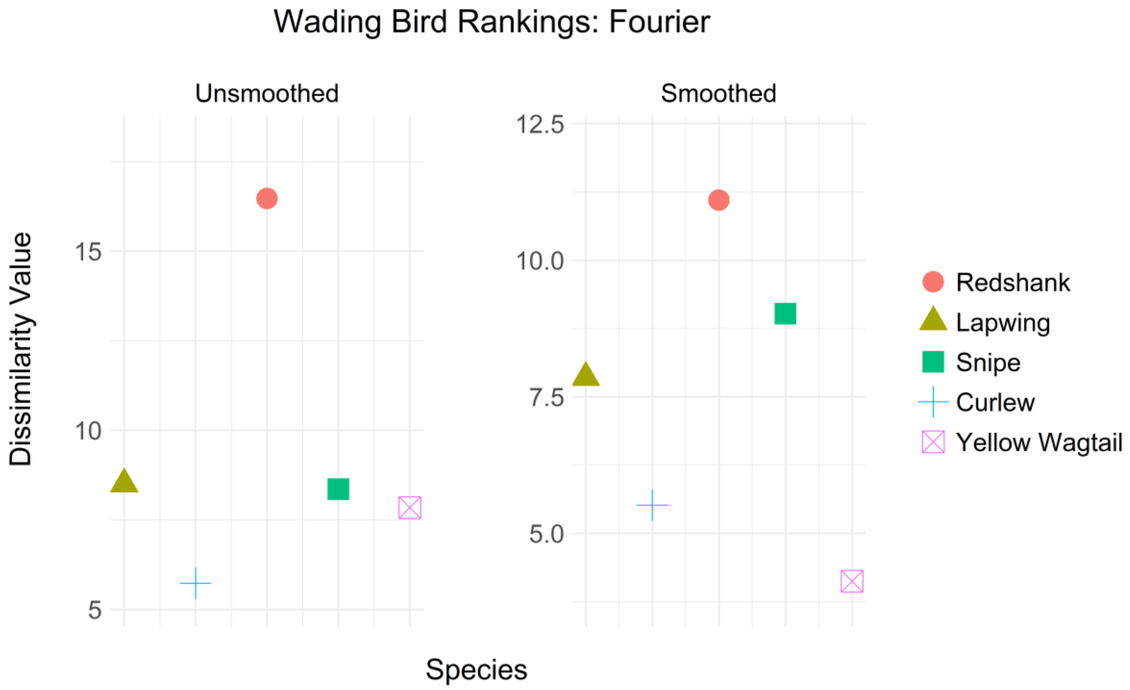
Wading Bird Rankings: Fourier
Unsmoothed
Smoothed
Dissimilarity Value
15
10
5
12.5
10.0
7.5
5.0
Species
Redshank
Lapwing
Snipe
Curlew
Yellow Wagtail

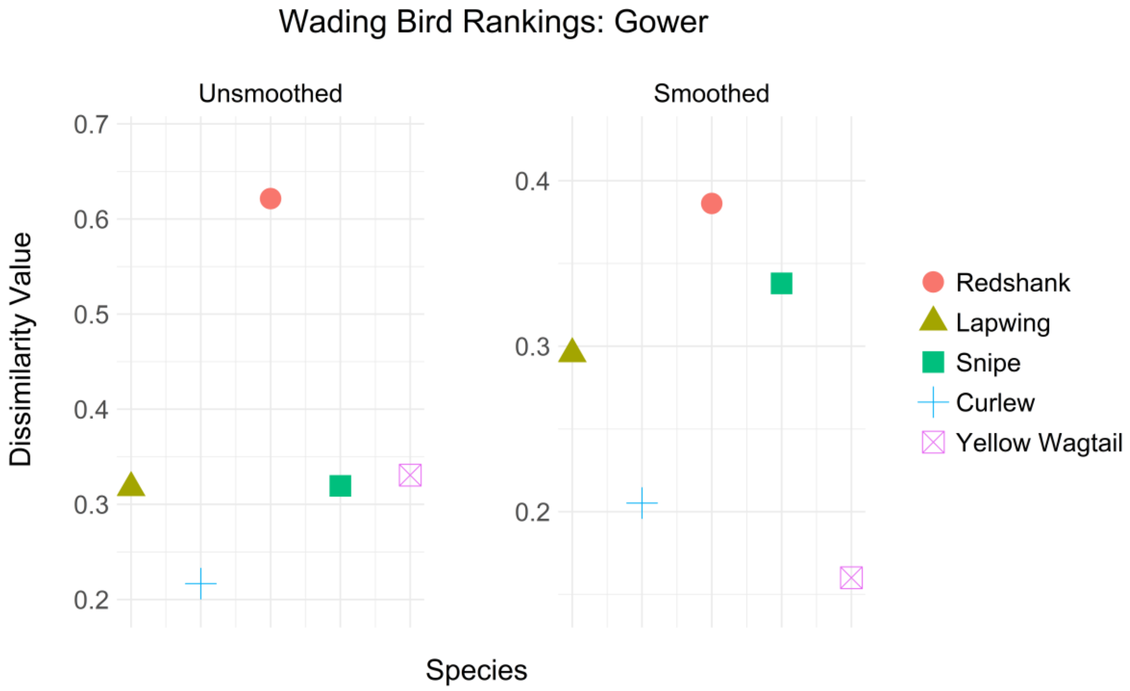
Wading Bird Rankings: Gower
Unsmoothed
Smoothed
Dissimilarity Value
0.7
0.6
0.5
0.4
0.3
0.2
0.4
0.3
0.2
Species
Redshank
Lapwing
Snipe
Curlew
Yellow Wagtail

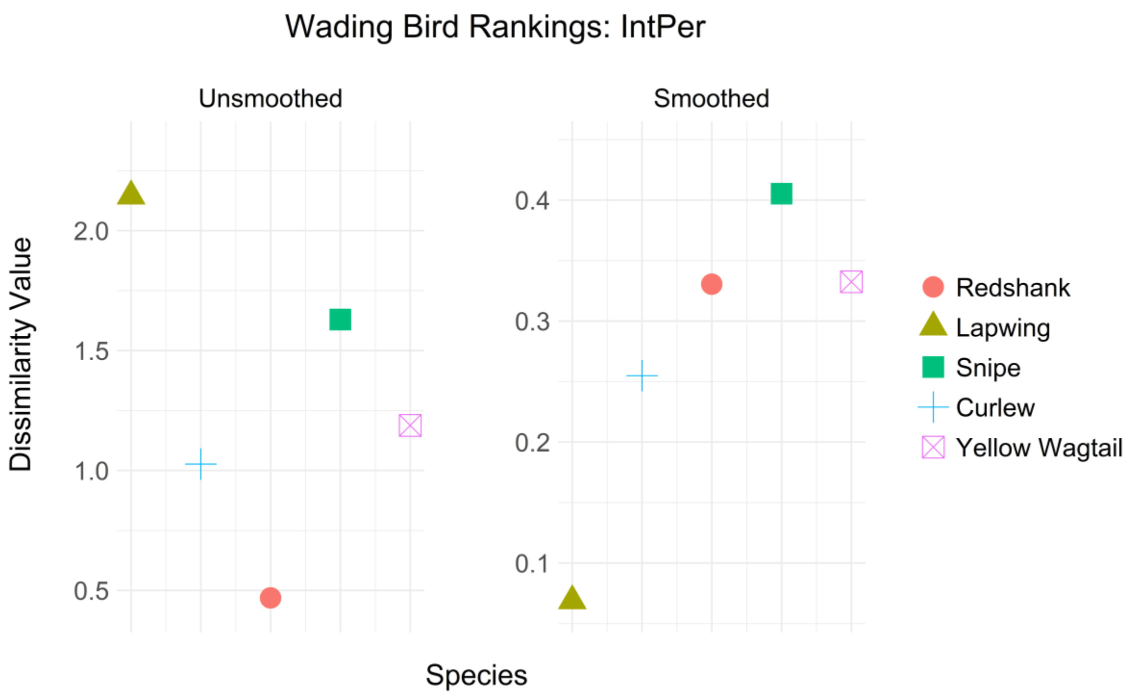
Wading Bird Rankings: IntPer
Unsmoothed
Smoothed
Dissimilarity Value
2.0
1.5
1.0
0.5
0.4
0.3
0.2
0.1
Species
Redshank
Lapwing
Snipe
Curlew
Yellow Wagtail

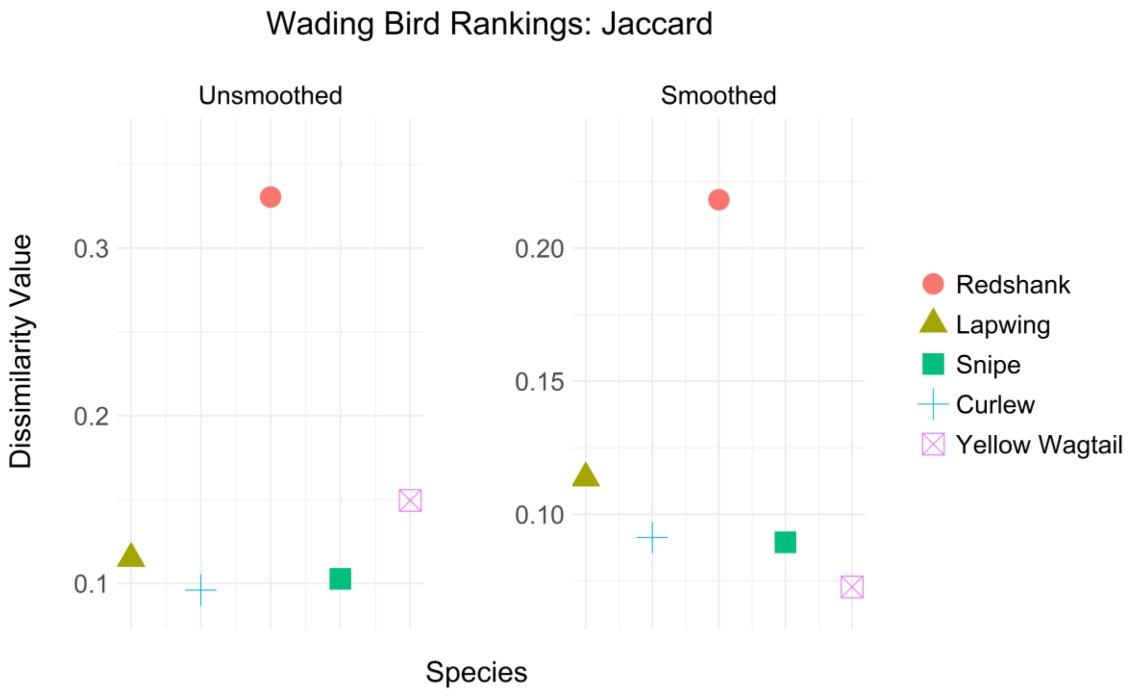
Wading Bird Rankings: Jaccard
Unsmoothed
Smoothed
Dissimilarity Value
0.3
0.2
0.1
0.20
0.15
0.10
Species
Redshank
Lapwing
Snipe
Curlew
Yellow Wagtail

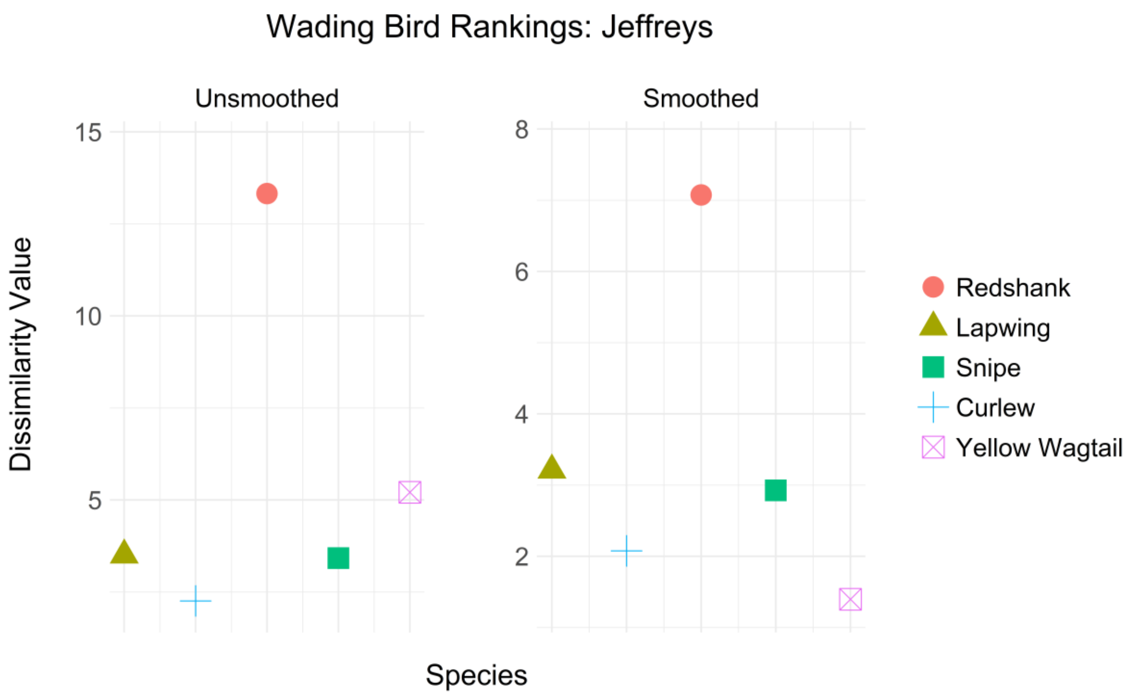
Wading Bird Rankings: Jeffreys
Unsmoothed
Smoothed
Dissimilarity Value
15
10
5
8
6
4
2
Species
Redshank
Lapwing
Snipe
Curlew
Yellow Wagtail

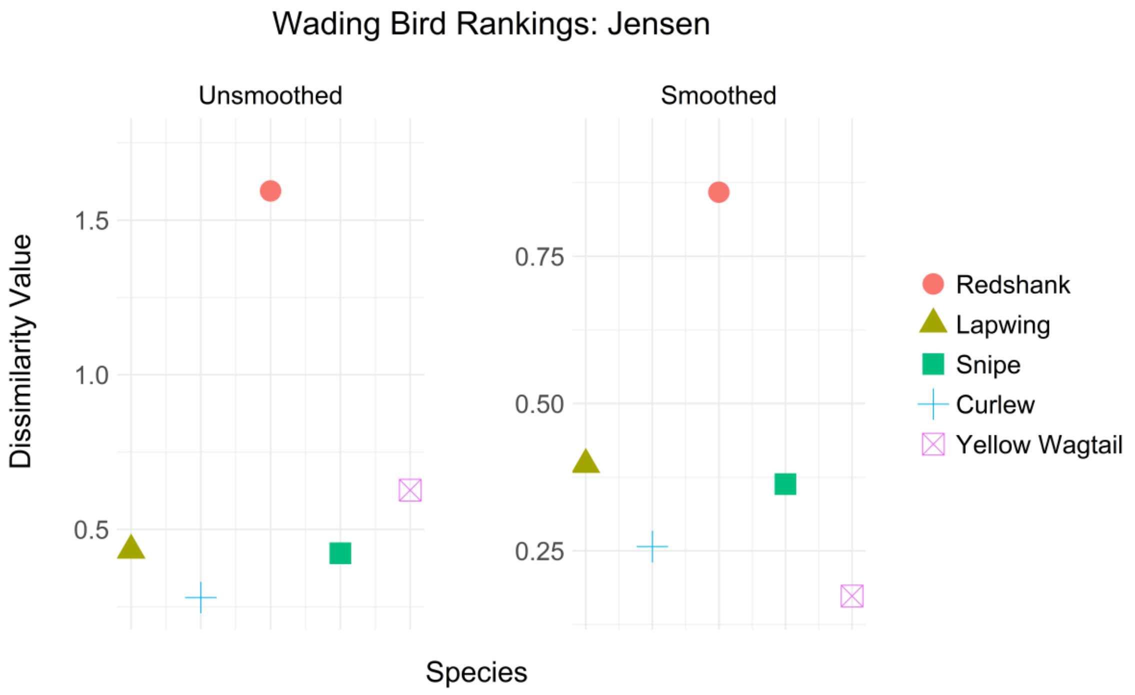
Wading Bird Rankings: Jensen
Unsmoothed
Smoothed
Dissimilarity Value
1.5
1.0
0.5
0.75
0.50
0.25
Species
Redshank
Lapwing
Snipe
Curlew
Yellow Wagtail

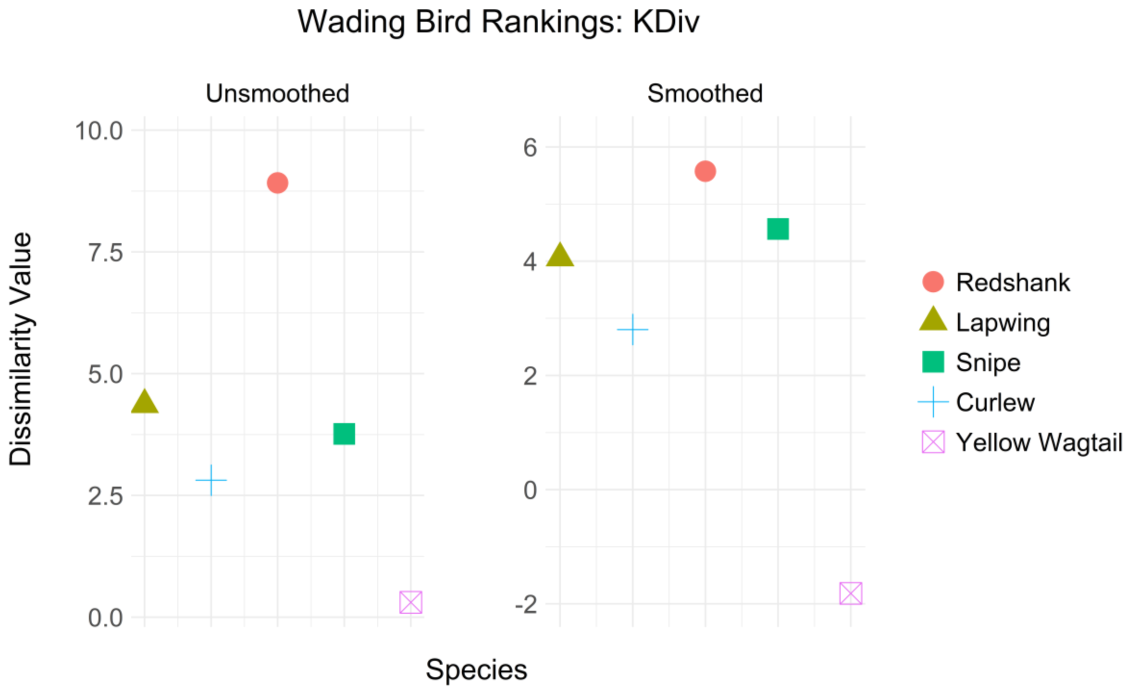
Wading Bird Rankings: KDiv
Unsmoothed
Smoothed
Dissimilarity Value
10.0
7.5
5.0
2.5
0.0
6
4
2
0
-2
Species
Redshank
Lapwing
Snipe
Curlew
Yellow Wagtail

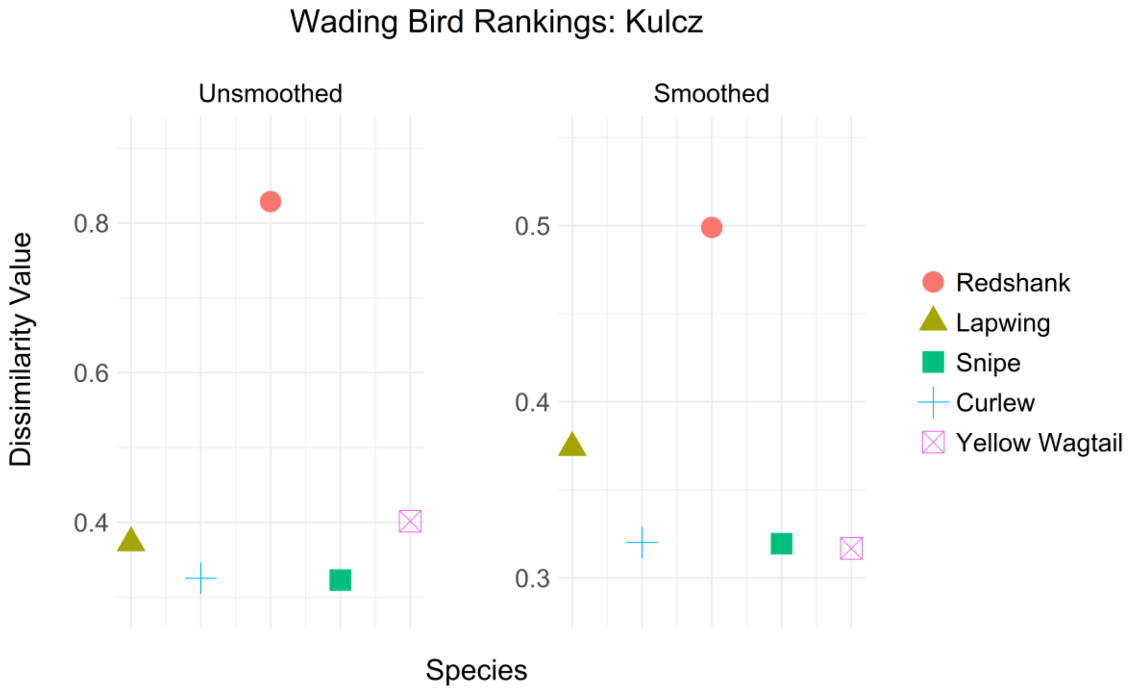
Wading Bird Rankings: Kulcz
Unsmoothed
Smoothed
Dissimilarity Value
0.8
0.6
0.4
0.5
0.4
0.3
Species
Redshank
Lapwing
Snipe
Curlew
Yellow Wagtail

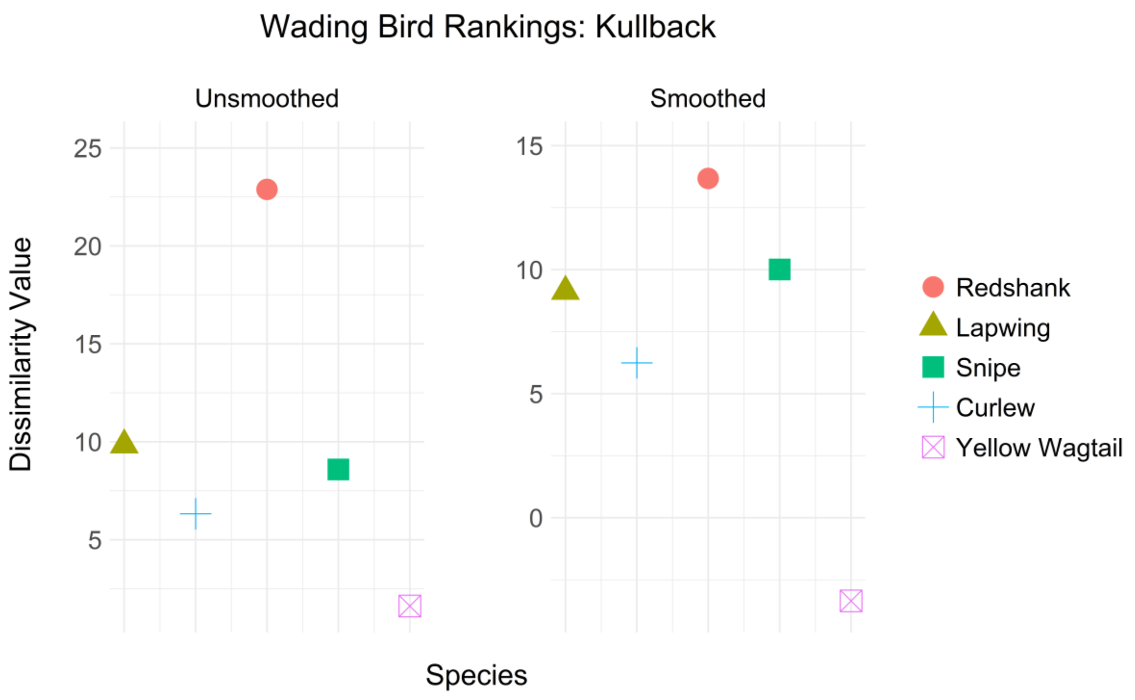
Wading Bird Rankings: Kullback
Unsmoothed
Smoothed
Dissimilarity Value
25
20
15
10
5
15
10
5
0
Species
Redshank
Lapwing
Snipe
Curlew
Yellow Wagtail

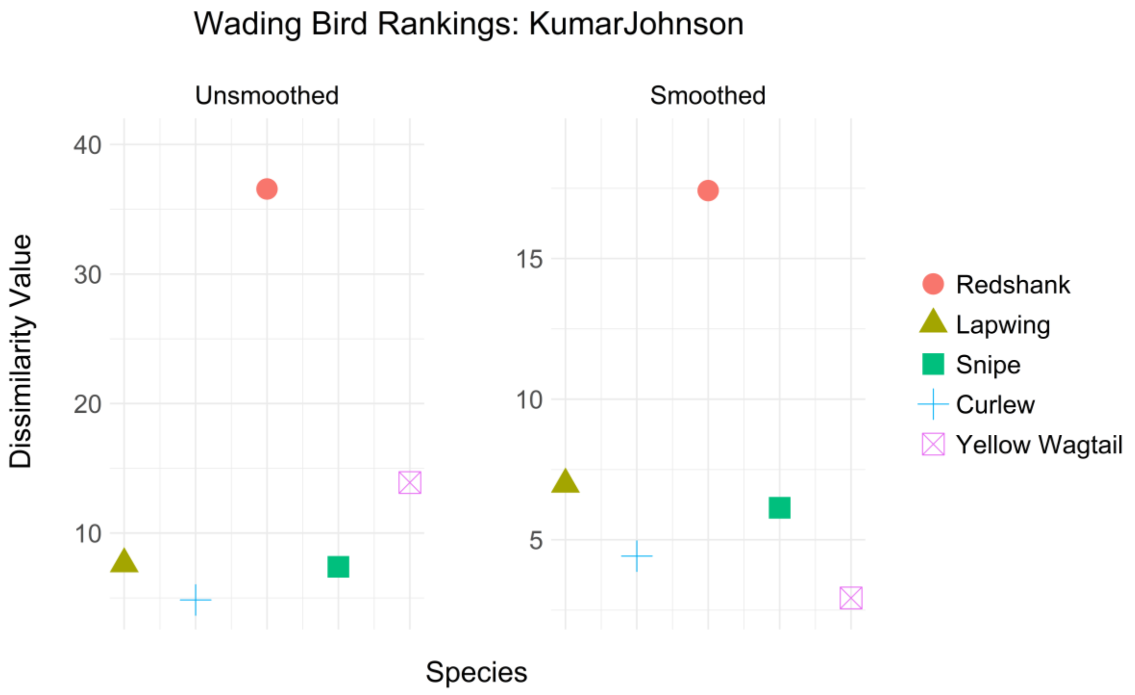
Wading Bird Rankings: KumarJohnson
Unsmoothed
Smoothed
Dissimilarity Value
40
30
20
10
15
10
5
Species
Redshank
Lapwing
Snipe
Curlew
Yellow Wagtail

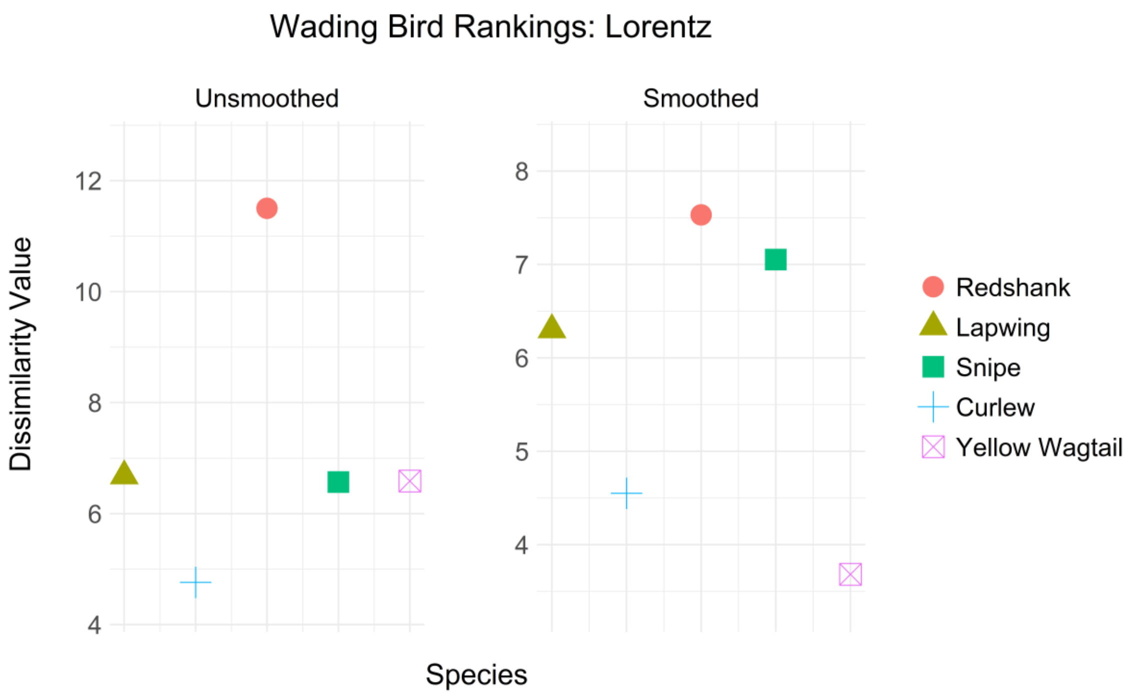

Wading Bird Rankings: Lorentz
Unsmoothed
Smoothed
Dissimilarity Value
12
10
8
6
4
8
7
6
5
4
Species
Redshank
Lapwing
Snipe
Curlew
Yellow Wagtail


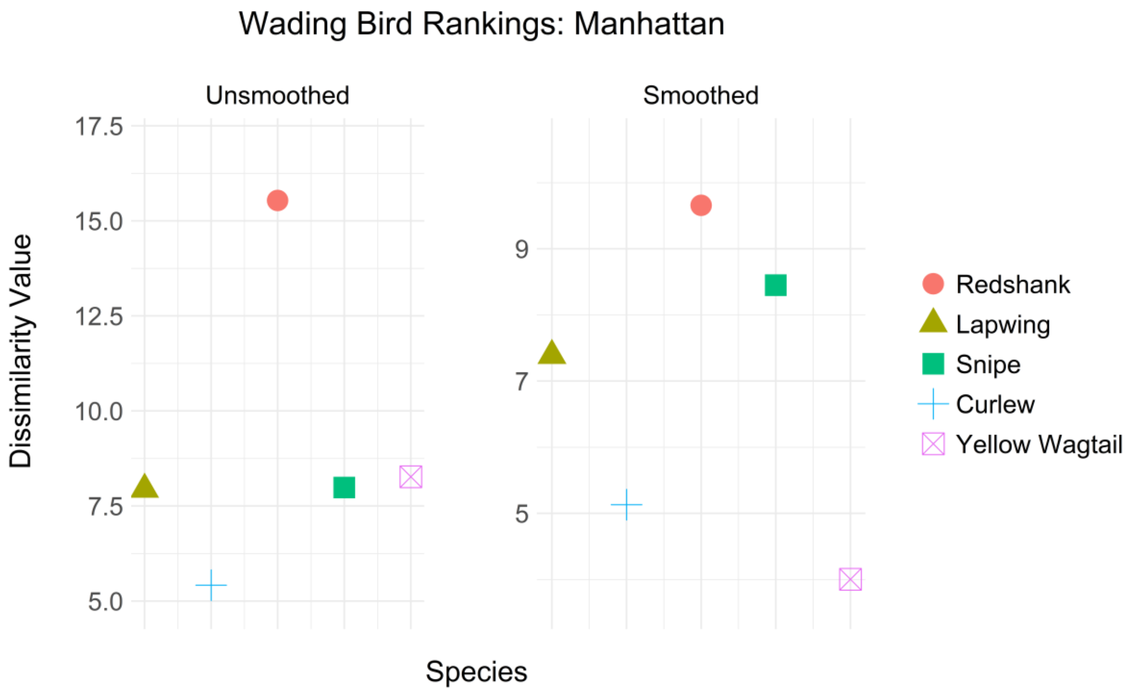

Wading Bird Rankings: Manhattan
Unsmoothed
Smoothed
Dissimilarity Value
17.5
15.0
12.5
10.0
7.5
5.0
9
7
5
Species
Redshank
Lapwing
Snipe
Curlew
Yellow Wagtail

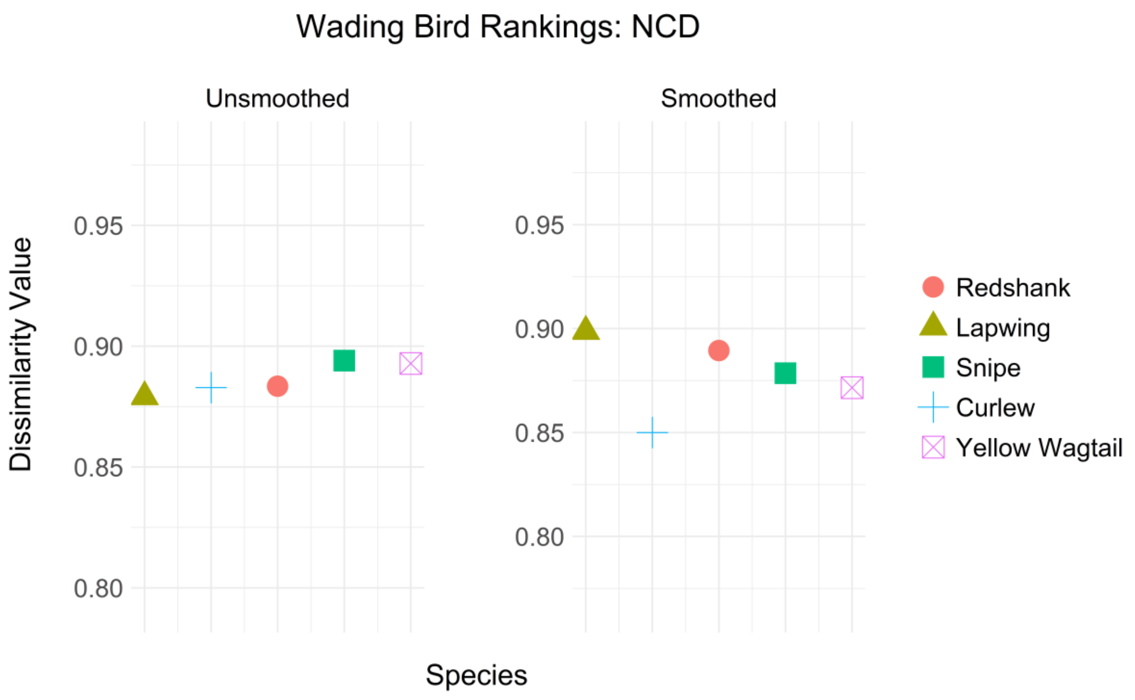
Wading Bird Rankings: NCD
Unsmoothed
Smoothed
Dissimilarity Value
0.95
0.90
0.85
0.80
Species
Redshank
Lapwing
Snipe
Curlew
Yellow Wagtail

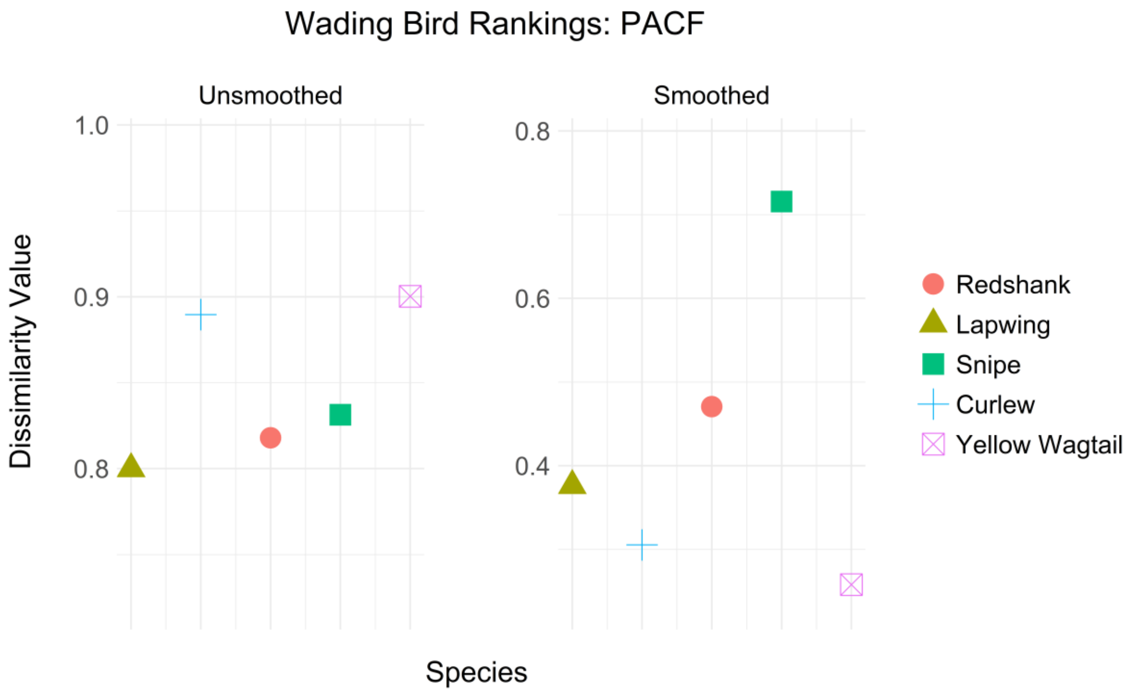
Wading Bird Rankings: PACF
Unsmoothed
Smoothed
Dissimilarity Value
1.0
0.9
0.8
0.6
0.4
Species
Redshank
Lapwing
Snipe
Curlew
Yellow Wagtail

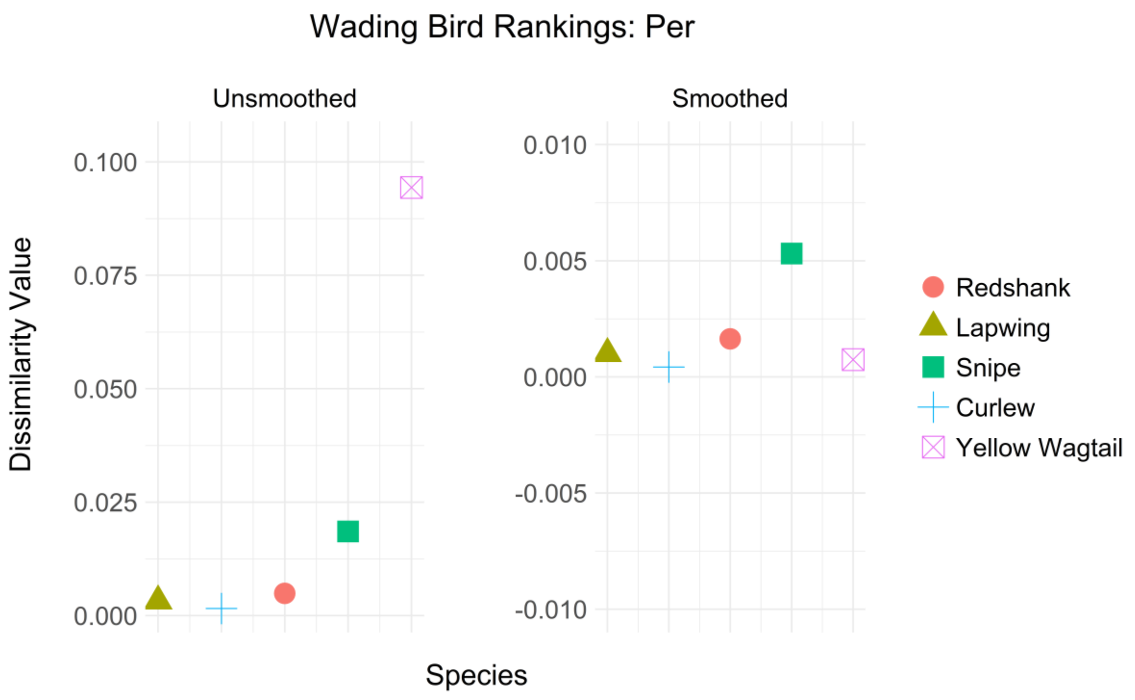
Wading Bird Rankings: Per
Unsmoothed
Smoothed
0.100
0.075
0.050
0.025
0.000
0.010
0.005
0.000
-0.005
-0.010
Dissimilarity Value
Species
Redshank
Lapwing
Snipe
Curlew
Yellow Wagtail

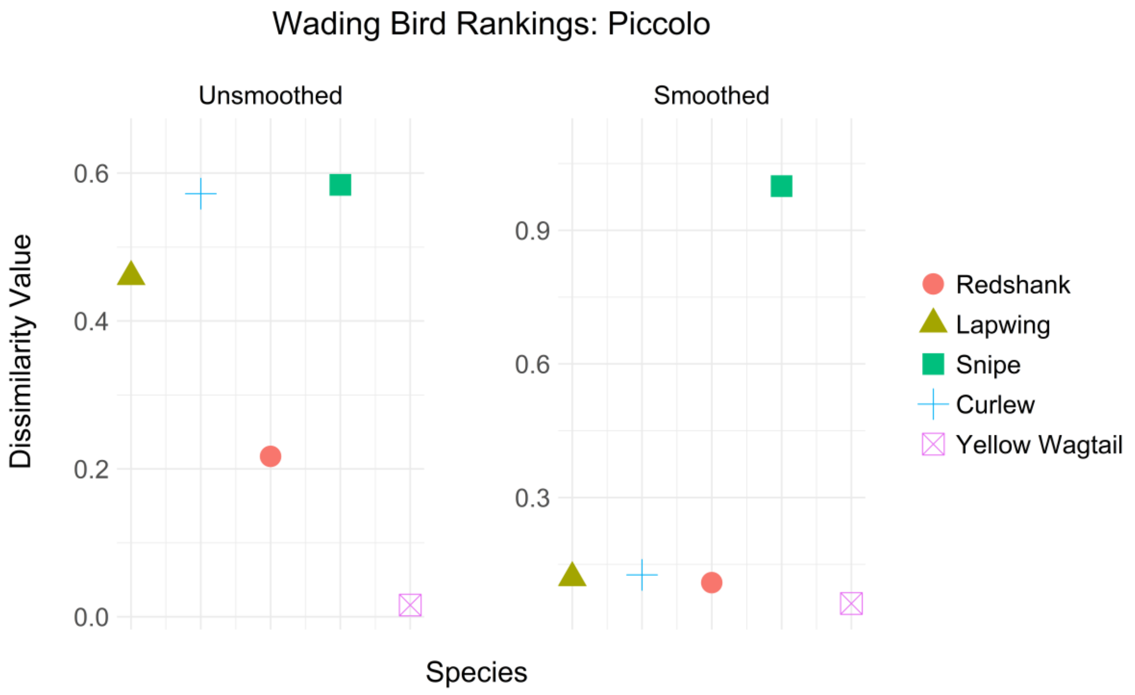
Wading Bird Rankings: Piccolo
Unsmoothed
Smoothed
0.6
0.4
0.2
0.0
0.9
0.6
0.3
Dissimilarity Value
Species
Redshank
Lapwing
Snipe
Curlew
Yellow Wagtail

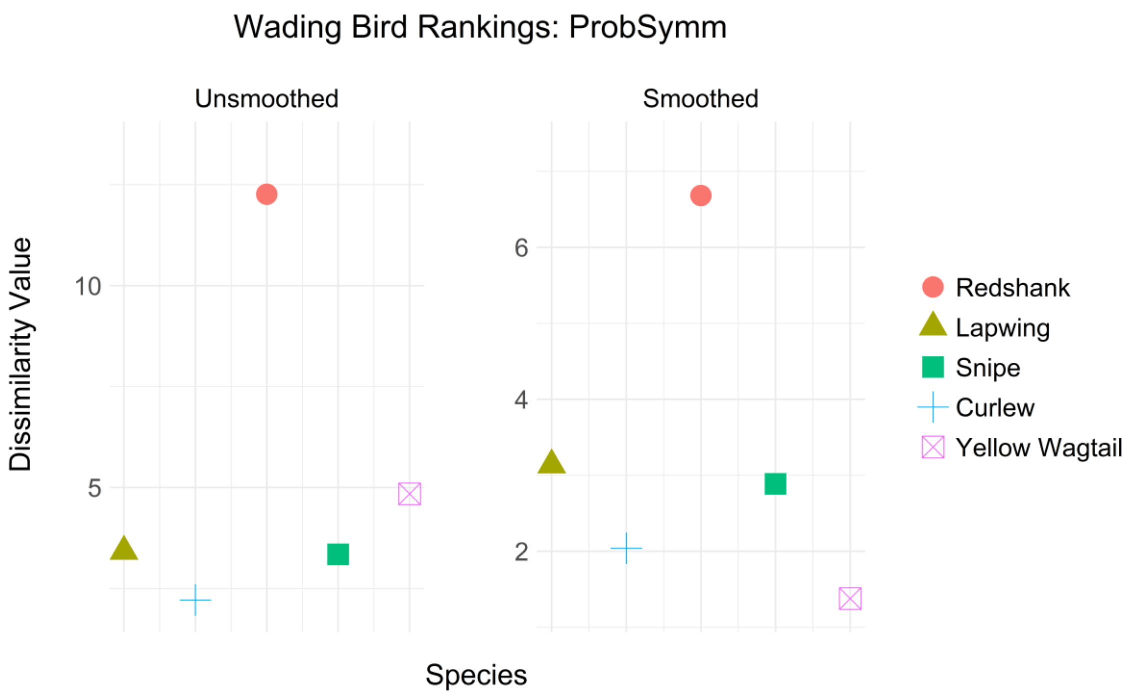
Wading Bird Rankings: ProbSymm
Unsmoothed
Smoothed
Dissimilarity Value
10
5
6
4
2
Species
Redshank
Lapwing
Snipe
Curlew
Yellow Wagtail

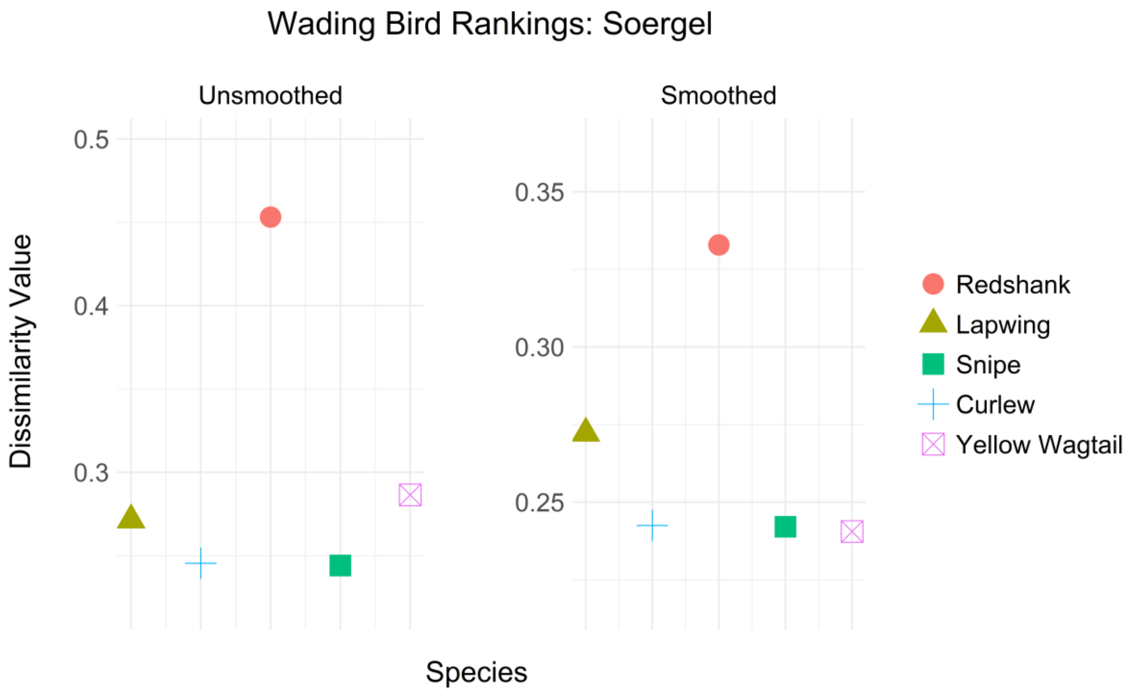
Wading Bird Rankings: Soergel
Unsmoothed
Smoothed
Dissimilarity Value
0.5
0.4
0.3
0.35
0.30
0.25
Species
Redshank
Lapwing
Snipe
Curlew
Yellow Wagtail

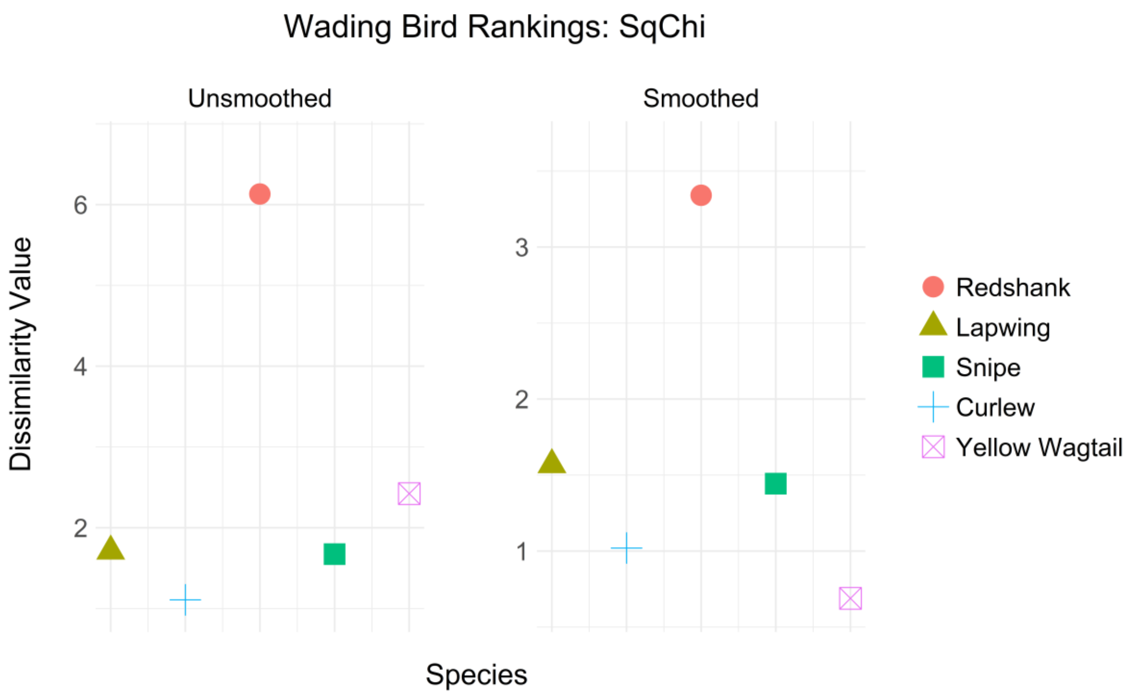
Wading Bird Rankings: SqChi
Unsmoothed
Smoothed
Dissimilarity Value
Species
Redshank
Lapwing
Snipe
Curlew
Yellow Wagtail

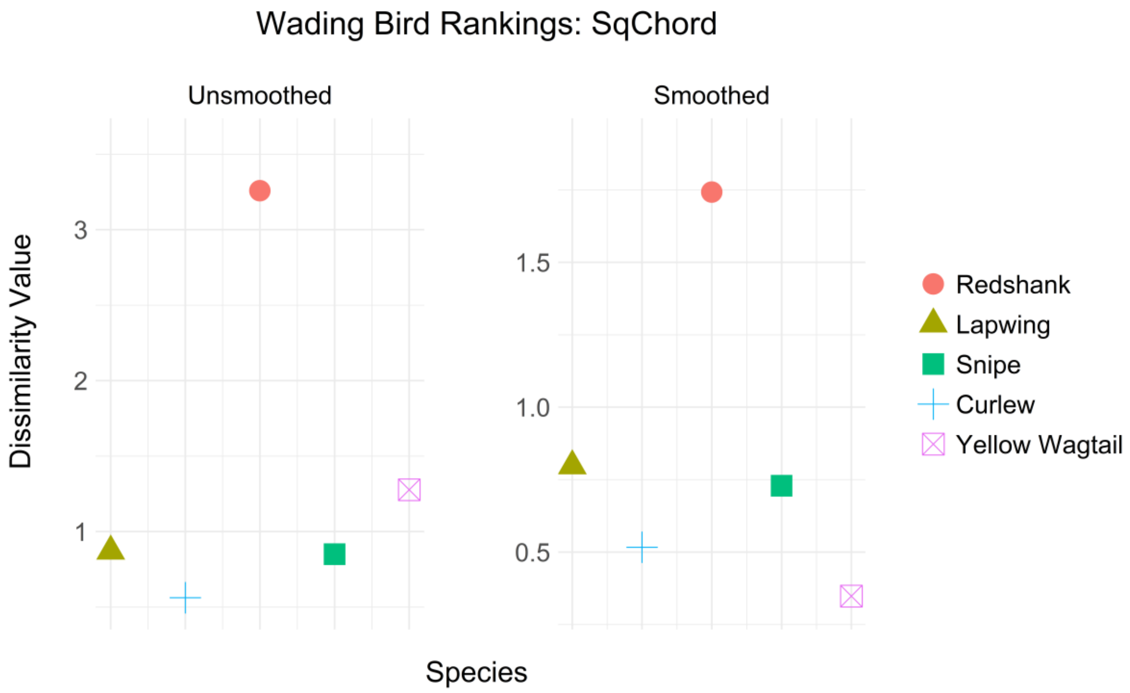
Wading Bird Rankings: SqChord
Unsmoothed
Smoothed
Dissimilarity Value
Species
Redshank
Lapwing
Snipe
Curlew
Yellow Wagtail

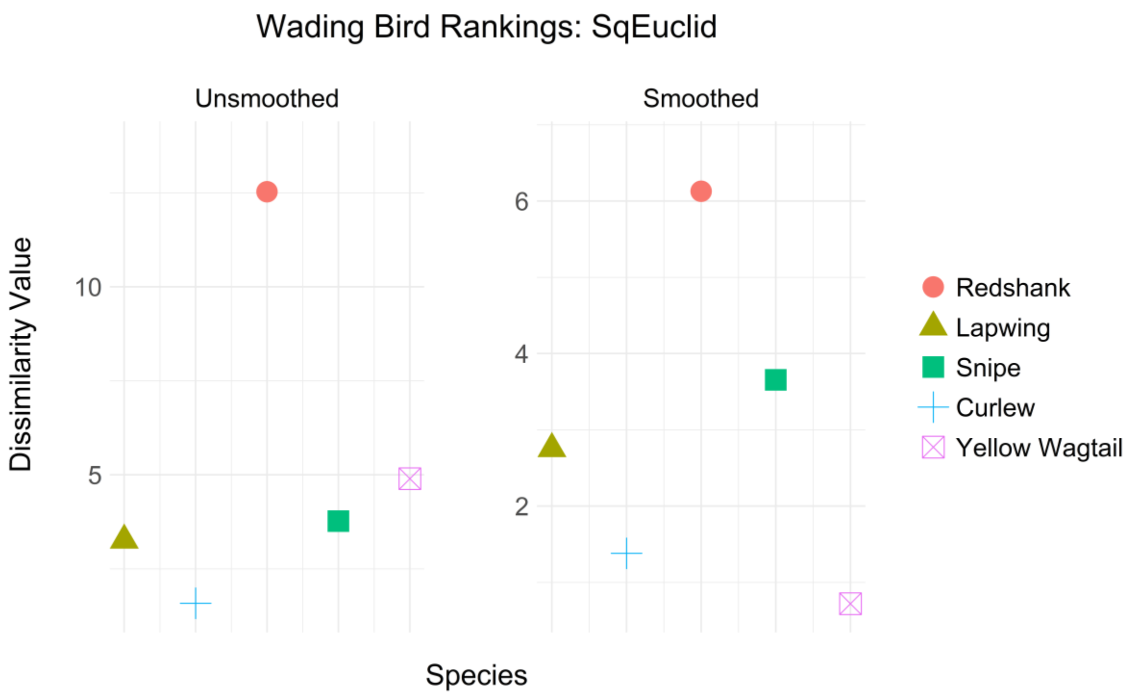

Wading Bird Rankings: SqEuclid
Unsmoothed
Smoothed
Dissimilarity Value
10
5
6
4
2
Species
Redshank
Lapwing
Snipe
Curlew
Yellow Wagtail


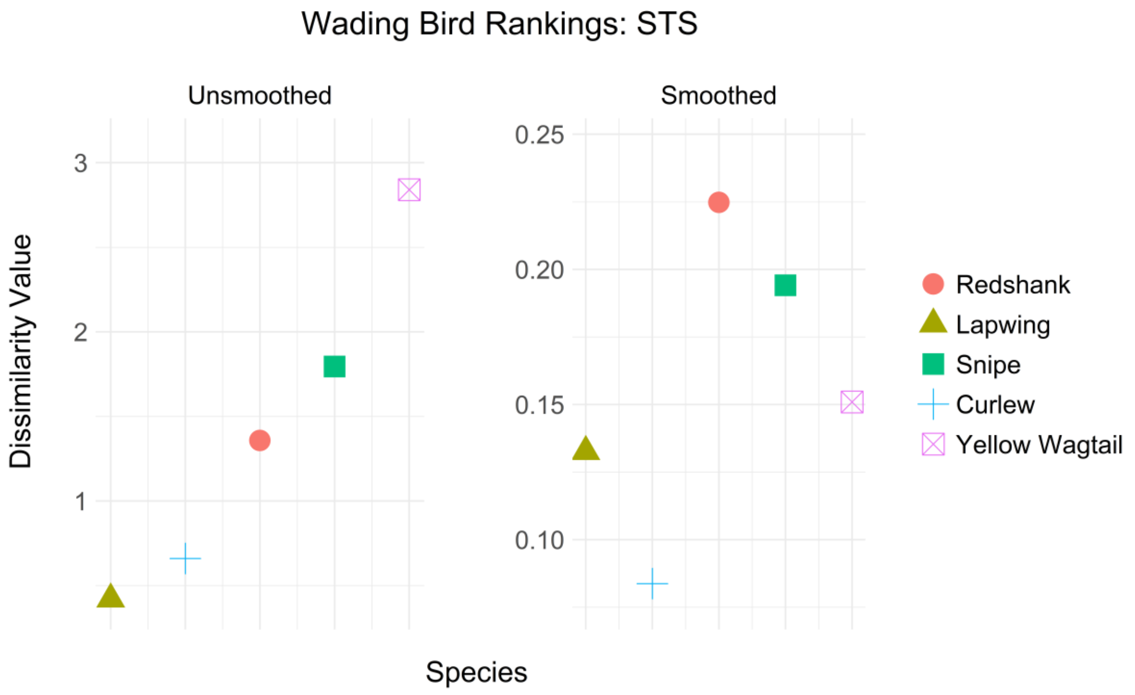

Wading Bird Rankings: STS
Unsmoothed
Smoothed
Dissimilarity Value
3
2
1
0.25
0.20
0.15
0.10
Species
Redshank
Lapwing
Snipe
Curlew
Yellow Wagtail

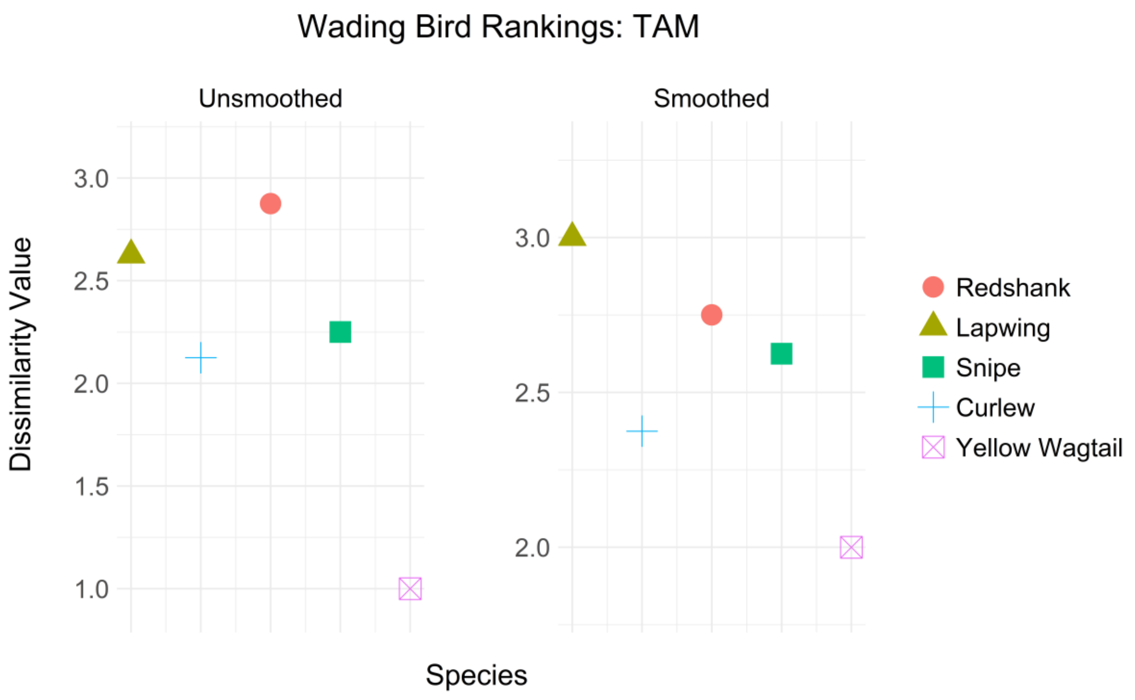

Wading Bird Rankings: TAM
Unsmoothed
Smoothed
Dissimilarity Value
3.0
2.5
2.0
1.5
1.0
3.0
2.5
2.0
Species
Redshank
Lapwing
Snipe
Curlew
Yellow Wagtail


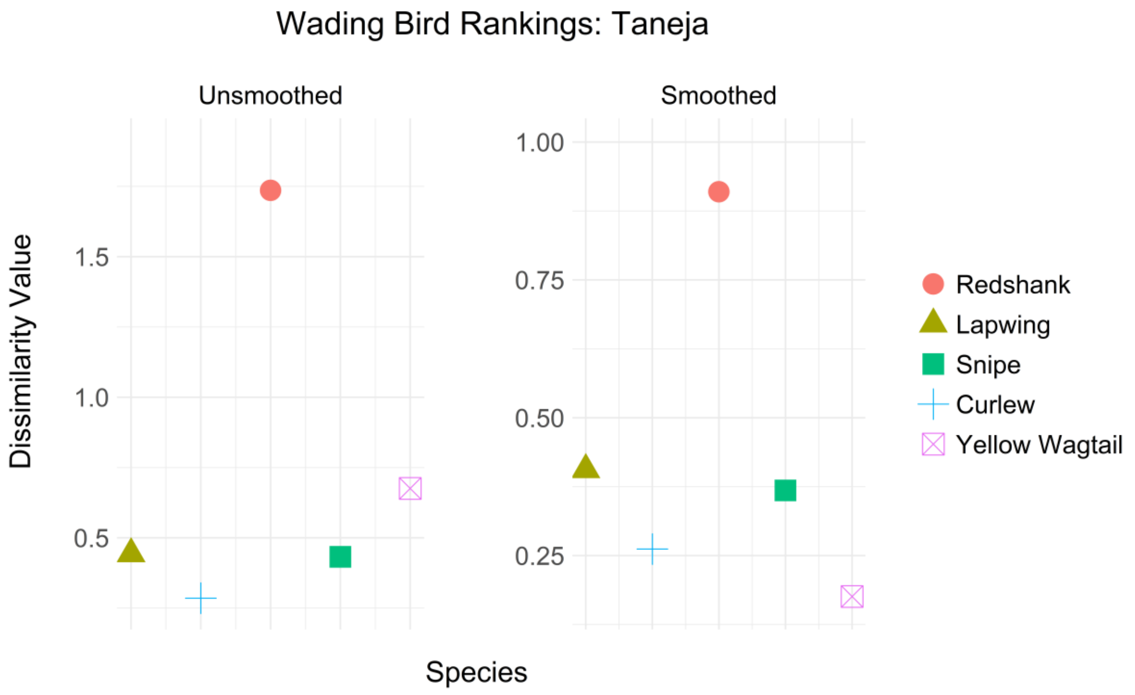

Wading Bird Rankings: Taneja
Unsmoothed
Smoothed
Dissimilarity Value
1.5
1.0
0.5
1.00
0.75
0.50
0.25
Species
Redshank
Lapwing
Snipe
Curlew
Yellow Wagtail

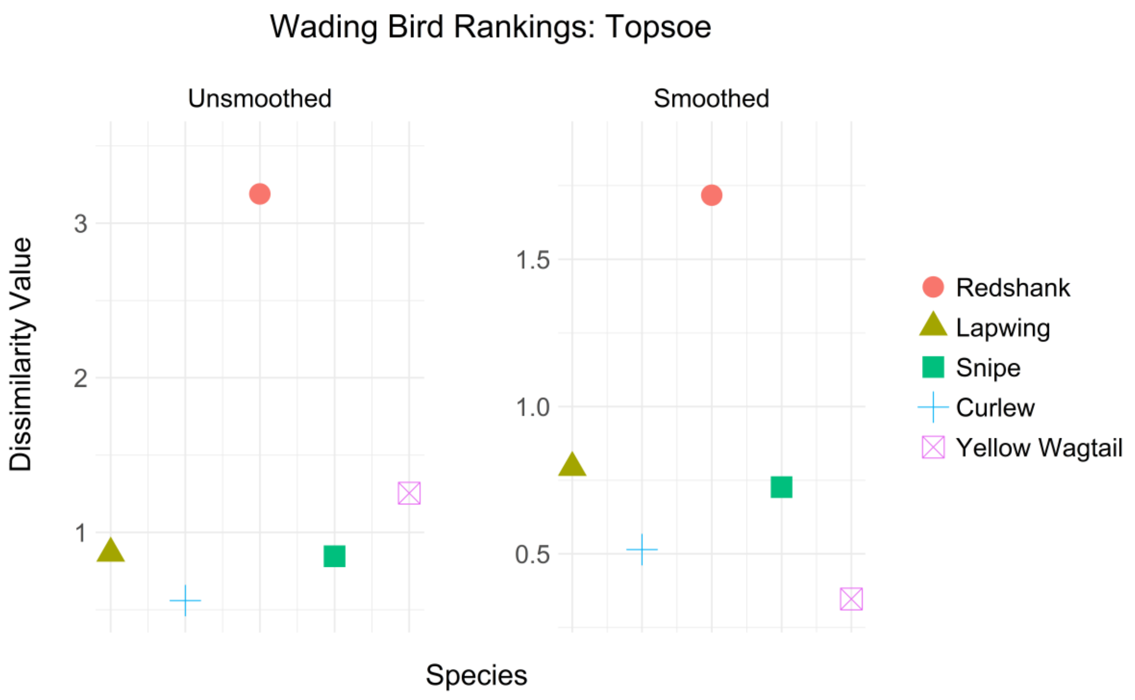

Wading Bird Rankings: Topsoe
Unsmoothed
Smoothed
Dissimilarity Value
3
2
1
1.5
1.0
0.5
Species
Redshank
Lapwing
Snipe
Curlew
Yellow Wagtail


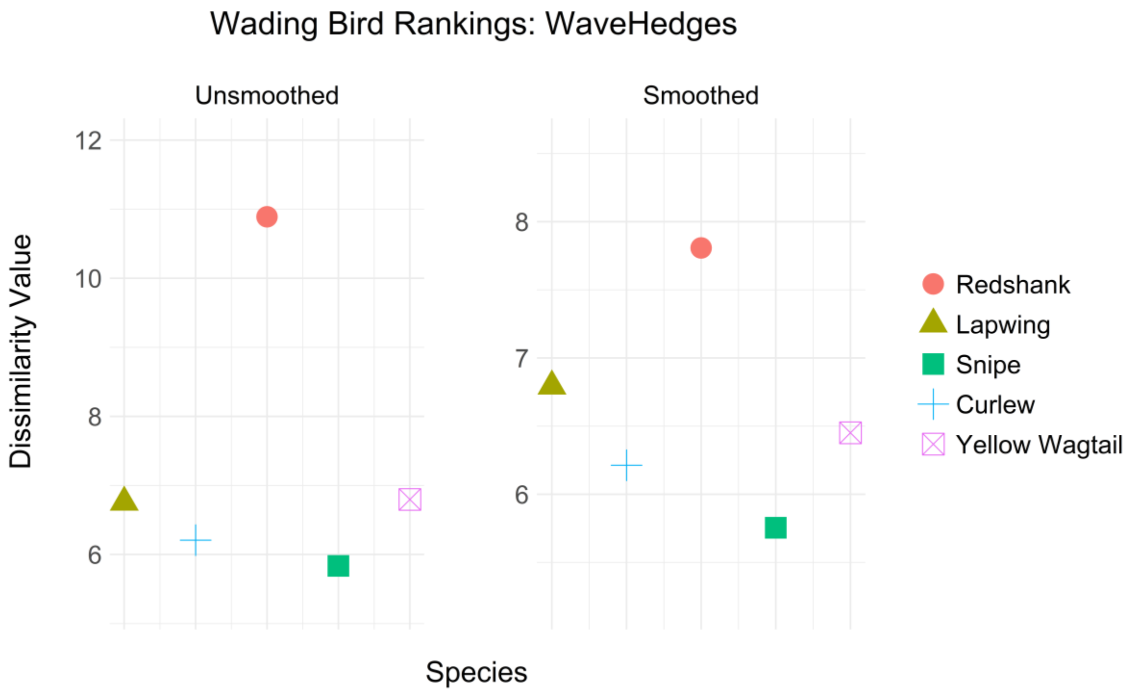

Wading Bird Rankings: WaveHedges
Unsmoothed
Smoothed
Dissimilarity Value
12
10
8
6
8
7
6
Species
Redshank
Lapwing
Snipe
Curlew
Yellow Wagtail
